# Biocompatible sulfonium-based covalent probes for endogenous tubulin fluorescence nanoscopy in live and fixed cells

**DOI:** 10.1101/2025.01.27.635008

**Authors:** Marie Auvray, Tanja Koenen, Olexandr Dybkov, Henning Urlaub, Gražvydas Lukinavičius

## Abstract

Fluorescent probes enable the visualization of dynamic cellular processes with high precision, particularly when coupled with super-resolution imaging techniques that surpass the diffraction limit. Traditional methods include fluorescent protein fusion (e.g., GFP) or organic fluorophores linked to ligands targeting the protein of interest. However, these approaches often introduce functional disruptions or ligand-associated biological effects. Herein, we address these challenges by developing covalent fluorescent probes for endogenous tubulin, a critical cytoskeletal protein involved in processes such as cell movement, division, and biomolecule trafficking. Using well-known tubulin binder cabazitaxel and cell permeable fluorophore silicon-rhodamine—as a basis, we introduce a novel biocompatible cleavable linker containing a sulfonium center. This allowed the construction of the optimized probe**, 6-SiR-*o*-C_9_-CTX**, demonstrating excellent cell permeability, fluorogenic properties, and the ability to covalently label tubulin across various human cell lines. Importantly, the targeting moiety could be washed out while preserving tubulin staining, ensuring minimal disruption of tubulin function. This labeling technique is compatible with STED nanoscopy in both live and fixed cells, offering a powerful high-resolution imaging tool.

## Introduction

Fluorescence microscopy offers a possibility to visualize dynamic processes in living cells. In the recent decades, super-resolution microscopy has become an essential tool to understand these biological processes with an unrivalled precision by overcoming the diffraction limit.^1^ These methods heavily relies on labeling techniques, which show a minimal influence on the function of the studied biomolecules^2,3^. In living cells, two strategies are mostly used for labeling of a protein-of-interest (POI). First, the fusion with a fluorescent protein (for example, the well-known Green Fluorescent Protein – GFP) or with a self-labelling tag that hosts a synthetic fluorophore.^4^ Nevertheless, these strategies add a large extra domain to the POI: for example, GFP (27 kDa) has a length of 4.2 nm and a diameter of 2.4 nm.^5^ This large size mainly causes two problems: mislocalization and function disruption of POI.^6^ The second approach involves using fluorescent probes, which consist of an organic fluorophore conjugated via a linker to a ligand that targets and non-covalently binds the POI. This method ensures the fluorescent tag is as small as possible. However, ligands can often exhibit biological effects and may occupy active sites on the protein, potentially altering its function. In this study, we aimed to explore an alternative labeling strategy for endogenous proteins that minimizes impact on their native function. In 2009, Hamachi introduced the ligand-directed labeling strategy.^7^ This approach relies on proximity induced reaction between a cleavable linker and POI. Once the probe binds to the POI, the nucleophilic side chain of an amino acid nearby the binding site can react with the cleavable linker to create a covalent bond between the tag and the protein, while releasing the ligand. Over the last fifteen years, cleavable linkers that can react with various amino acids have been developed.^8,9,10,11,12^ Despite the tremendous potential of this approach for fluorescence microscopy, it has not been extensively applied. In particular, its application in the field of fluorescence microscopy is mostly restricted to extracellular targets, like membrane receptors.^13–16^ The main reason is that it is challenging to find a compromise between reactivity and stability of the cleavable linker inside cells. In addition, making these probes cell-permeable can be an issue.

In this study, we developed a series of covalent fluorescent probes for endogenous tubulin. This protein is highly abundant in mammalian cells, in which it represents 3-4% of the total proteins and up to 10 % in brain.^17^ Two subunits α- and β-tubulin form heterodimers and polymerize into microtubules, one of the main components of the cytoskeleton. Both proteins exist in numerous isotypes encoded by different genes. In human, there are 9 genes for α-tubulin and 10 for β-tubulin.^18^ In addition, tubulin can undergo post-translational modifications, leading to a huge variety of forms in cells.^19^ It is therefore involved in many different cellular processes like cell movement, division or trafficking of biomolecules. We developed fluorescent probes based on widely used tubulin binders—taxanes, which are well-known anticancer drugs. To enhance their utility, we introduced a novel biocompatible cleavable linker containing a sulfonium center. Through *in vitro* and *in cellulo* evaluations, we identified silicon-rhodamine (SiR) probe **6-SiR-*o*-C_9_-CTX** as an outstanding fluorogenic, cell-membrane-permeable candidate. Importantly, we demonstrated that the ligand can be washed out while retaining robust tubulin staining, thus providing a labeling technique that minimally impacts protein structure and function. This probe enables covalent labeling of endogenous tubulin across various cell lines, making it suitable for stimulated emission depletion (STED) nanoscopy in both live and fixed cells.

## Results

### Design and synthesis of the probes

Tubulin-covalent probes are made of three parts: a dye, a ligand to direct the probe to the POI and a cleavable electrophile to create a covalent bond between the POI and the fluorophore, while releasing the ligand (**Figure 1A**).

**Figure 1.**
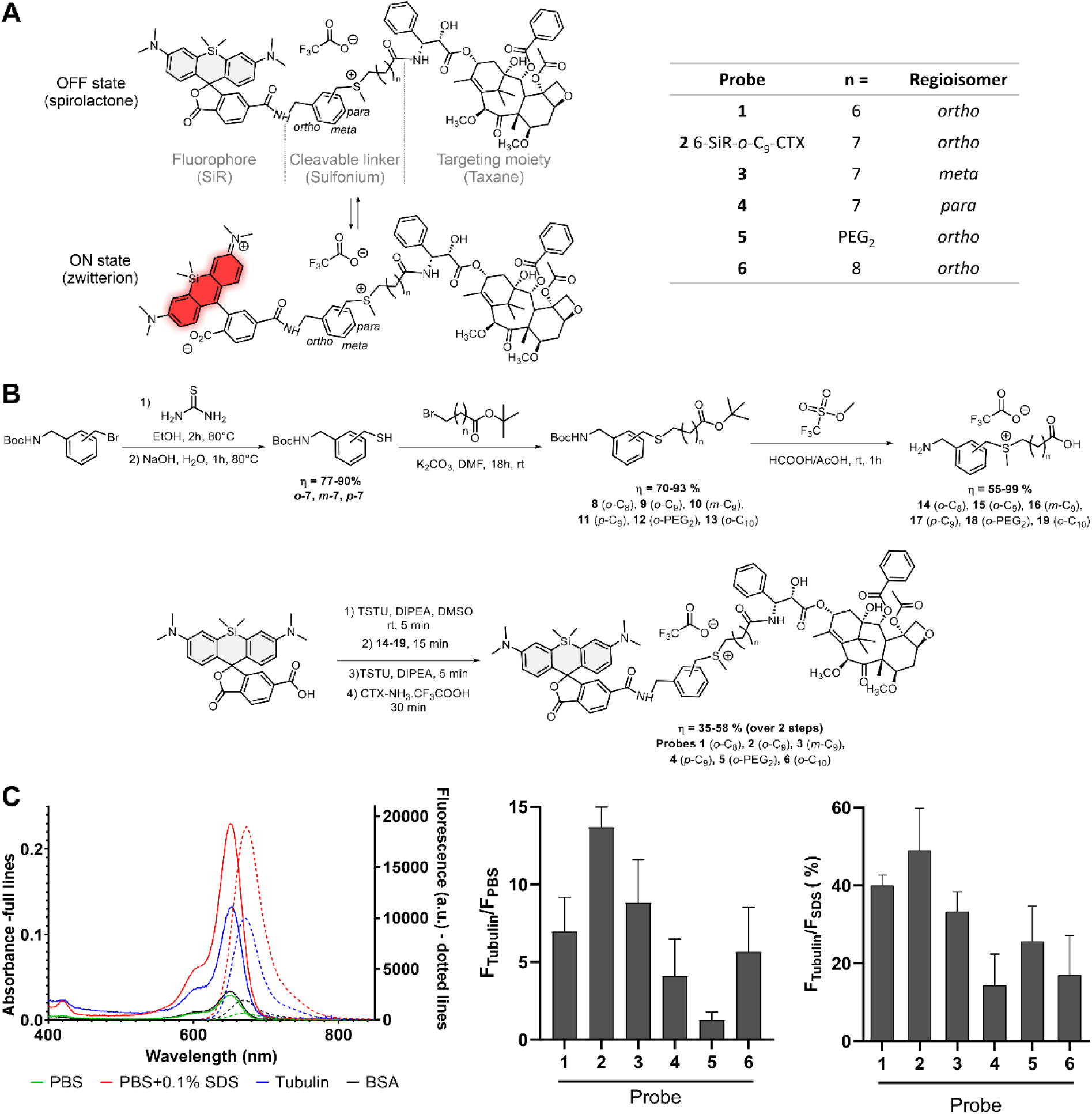
Structure, synthesis and properties of the covalent probes. (A) Structure of tubulin probes featuring a sulfonium-based cleavable linker and silicon-rhodamine (SiR) which can toggle between ON and OFF states. (B) Synthetic path of the covalent probes (C) Photophysical properties of the probes. Left: Absorption and emission spectra of **6-SiR-*o*-C_9_-CTX** in presence of tubulin (blue), PBS (green), in presence of BSA (black) and PBS containing 0.1% SDS (red). Spectra are represented as averages of three independently repeated experiments (N=3). Middle: Fluorescence increase upon tubulin binding, as compared to PBS. Right: Percent of dye open when bound to tubulin. Data are presented as mean of triplicate with standard deviation.

Many cleavable electrophiles have been developed in the past ten years, highlighting the importance of finding a compromise between the reactivity and the stability of probe. We focused our attention on sulfonium, a biocompatible electrophile, that is endogenously present at high concentrations in cells across multiple species.^6–10^ In addition, recent proteomic studies by Z. Li demonstrated promising results for ligand-directed chemistry with *para*-benzyl sulfonium.^20,21^ Rhodamine derivatives are among the most popular dyes to develop probes for live-cell imaging. They are in equilibrium between two forms: a nonfluorescent hydrophobic spirolactone (OFF state) and a fluorescent hydrophilic zwitterion (ON state) (**Figure 1A**). In aqueous media, aggregation drives rhodamine to the spirolactone form, which is cell permeable. Once bound to its target, the interactions with the protein will push equilibrium towards the fluorescent zwitterionic form. One of the most prominent examples is SiR, which displays outstanding fluorogenicity and cell permeability.^22^ In our previous studies, we showed that the best performing ligand for tubulin in combination with SiR isomer-6 derivatives was cabazitaxel.^23,24^ We thus performed a mini screening using silicon-rhodamine as dye and cabazitaxel (CTX) as ligand, and fine-tuned the structure by probing three different regioisomers (*ortho*, *meta* and *para*) of the cleavable electrophile (**Figure 1A** and ESI Figure S1).

The probes were synthesized *via* a convergent 7-step synthetic path with overall yields between 20 and 43 % (**Figure 1B**). First, thiols **7** were synthesized from the corresponding bromine derivatives, performing a nucleophilic substitution with thiourea followed by hydrolysis. Afterwards, thiols reacted with bromine derivatives of various chain lengths to give thioethers **8-13**. Sulfoniums with free amines and carboxylic acids **14**-**19** were obtained with excellent yields *via* a one pot reaction in a mixture of formic and acetic acids in presence of an excess of methyl trifluoromethanesulfonate. The synthesis ended with a four-step, one pot reaction that links the cleavable electrophile to both the ligand and the dye. This last step allowed to get expected compounds with good yields ranging from 35 to 58%. Overall, this synthetic path is highly convergent, which allows a rapid access to structural diversity.

### Photophysical properties

Photophysical properties of the probes were investigated in phosphate-buffered saline (PBS), in PBS containing 0.5 mg/ml bovine serum albumin (BSA), in PBS containing 0.1% sodium dodecyl sulfate (SDS) or in presence of tubulin after 4h at 37°C to ensure complete tubulin polymerization (**Figure 1C**, ESI Figures S2, S3 and Table S1). As mentioned previously, SiR-based probes are in equilibrium between two forms: a spirolactone and a zwitterionic forms. Multiple studies demonstrate that SiR equilibrium is highly sensitive to the environment.^22,25,26^ In aqueous media (e.g. PBS), the SiR is mainly in closed spirolactone form and tends to form aggregates, which explains the low absorbance and emission observed in this media for most of the probes. However, probe **5**, containing a PEG linker, exhibits a fluorescence at least six times higher than other probes in aqueous media (ESI Figure S3). This linker prevents aggregation by increasing water solubility, and therefore push the equilibrium towards the open fluorescent form. In the presence of SDS, aggregation is completely inhibited, with SiR adopting a zwitterionic form. This results in similar photophysical properties across all derivatives. Upon binding to tubulin, a shift in the spirocyclization equilibrium causes a 4- to 14-fold fluorescence enhancement relative to PBS for all probes except probe **5**. For probe **5**, only a modest increase in fluorescence (1.4- fold) is observed, due to its residual fluorescence in aqueous media. Among the probes, probe **2** demonstrates the highest fluorescence enhancement upon tubulin binding compared to PBS (14.3- fold, Table S1). Notably, the non-target protein bovine serum albumin (BSA) induces only minimal changes in absorbance and fluorescence (ESI Figure S3).

We assumed that the probes fluorescence in presence of PBS containing 0.1% SDS corresponds to 100 % of SiR fluorescent zwitterion. This allowed to estimate the equilibrium shift upon tubulin binding. We estimate that probes can reach up to 49 % of fluorescent zwitterion content once bound to tubulin (**Figure 1C**). Overall, probe **2** possesses the best photophysical properties, being the most fluorescent (49% of probe in ON state after binding to tubulin) and the most fluorogenic (14.3-fold fluorescent enhancement in presence of tubulin compared to PBS).

### *In vitro* reaction with purified tubulin

Next, we checked the stability of the probes in aqueous media (PBS) at 37°C as some probes containing cleavable linkers might be stable only for a few hours in solution.^12^ Only minor degradation was observed, as more than 75% of the probes remain after 48h at 37°C (ESI Figure S4). Major degradation pathways were hydrolysis (approx. 5-10% after 48h) and intramolecular reaction (approx. 5% after 48h), as the ligand contains some free nucleophilic groups. Prior to live-cell experiments, reactions between purified tubulin from porcine brain and the probes were conducted *in vitro* at 37°C (**Figure 2**, ESI Figure S5). Probes were used in 4-fold excess compared to tubulin in order to assess the selectivity of the reaction. The reaction was monitored by SDS–polyacrylamide gel electrophoresis (SDS-PAGE) and *in-gel* fluorescence analysis.

**Figure 2.**
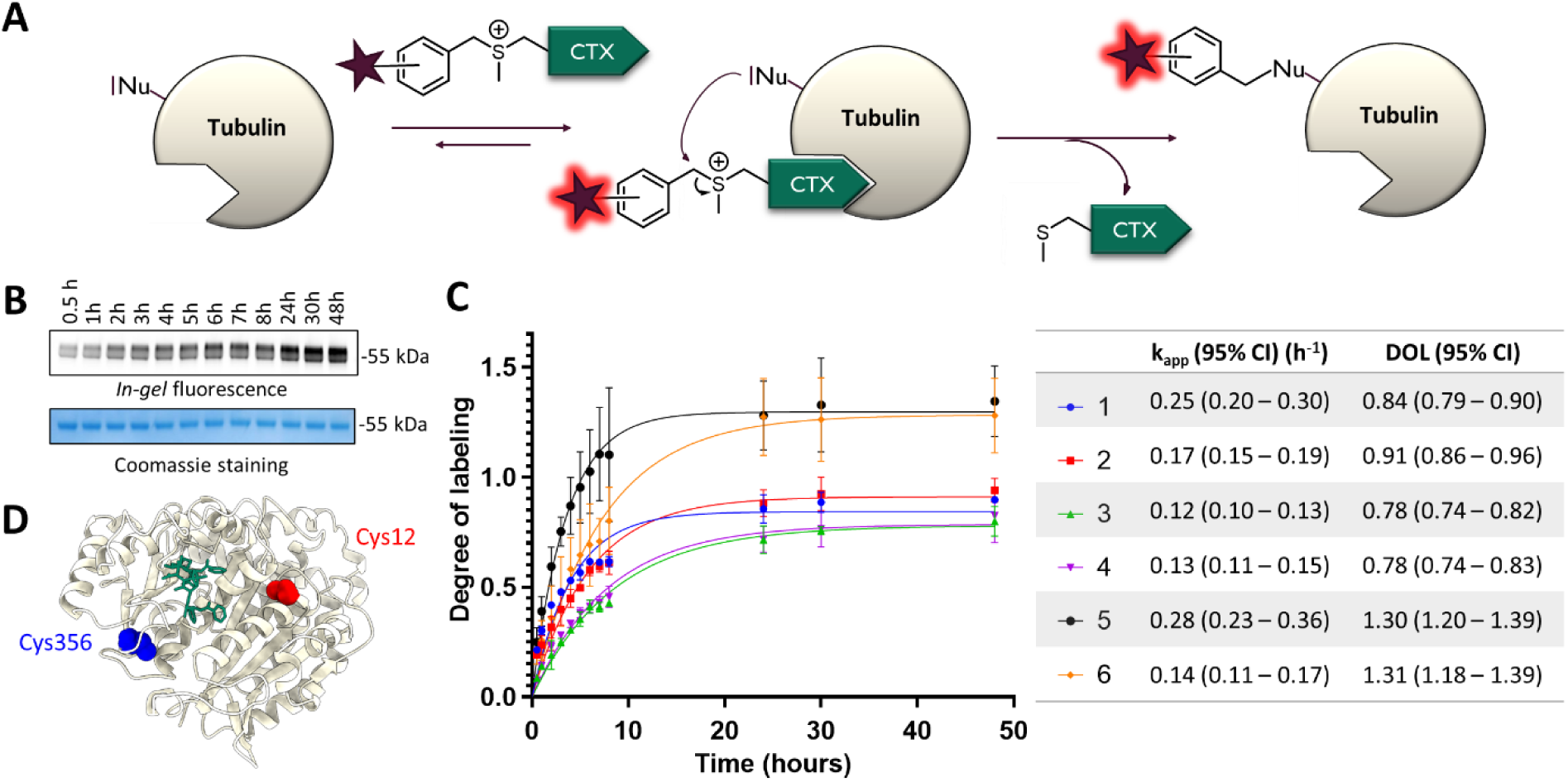
*In vitro* labeling experiments of purified pig (Sus scrofa) tubulin (0.5 mg/mL ̴ 5 µM) with the covalent probes (20 µM) over 48h at 37°C in General Tubulin Buffer. (**A**) Reaction between the probes and tubulin. (**B**) Representative SDS-PAGE analysis of the labeling reaction. The gel was analyzed by *in-gel* fluorescence imaging (up) and stained with Coomassie Brilliant Blue (down). (**C**) Time-course of *in vitro* labeling of tubulin with the covalent probes. The experiments were performed in triplicate to obtain mean and standard deviation values (shown as error bars). Table contains mean values and 95% confidence intervals (Cl) obtained from fitting data points to one phase association model from GraphPad Prism. (**D**) Structure of β-tubulin (from pig – PDB: 5SYF) with Taxol. The main labeling site is highlighted in blue (Cys 356), and the minor labeling site in red (Cys 12).

Apparent reaction rate was ranging from 0.12 to 0.28 h^-1^ and it corresponds to a second order rate constant approx. 2 to 5 L_•_mol^-1^_•_s^-1^, which is comparable to previously reported LDAI (ligand-directed acyl-imidazole chemistry).^27^ On average, reactions with *ortho*-isomers were around 50% faster than with *meta* or *para* isomers. Compounds can be divided into two groups based on the degree-of-labeling (DOL). Probes **1-4** reach DOL between 0.8 and 0.9, which corresponds to an excellent efficiency, close to ideal DOL of 1. Decent labeling yield (around 50%) can be achieved within 2-3 hours for Probe **1** and **6-SiR-*o*-C_9_-CTX** (**2**). Probes **5** and **6** demonstrate DOL > 1 suggesting multiple labeling sites and a lower selectivity.

Afterwards, we also checked that the bond between the dye and tubulin was stable in aqueous conditions (ESI Figure S6). Indeed, no degradation was observed after 48h in PBS (pH=7.4) at 37°C. The labeling site(s) were then determined by proteolytic *in-gel* digestion followed by mass spectrometry. Seven different peptides were observed with a +573 Da modification, corresponding to five different sequences labeled with SiR (ESI Figures S7 & S8, Table S2). For all these peptides, the modified amino acid was a cysteine, suggesting that sulfonium ligand directed chemistry is selective for cysteine. The high number of observed peptides can be explained by the numerous tubulin isotypes and isoforms contained in the sample. Indeed, most of the identified peptides come from different tubulin isotypes but correspond to the same modified amino acid. Overall, a primary labeling site and three secondary labeling sites were identified. The major labeling site was Cys356, located within the TAV<u>C</u>DIPPR/VAV<u>C</u>DIPPR peptide of β-tubulin (ESI Figures S7 & S8, Table S2). Additionally, a secondary labeling site was identified in β-tubulin at Cys12 within the peptide EIVHIQAGQ<u>C</u>GNQIGAK, which MS1 precursor abundance was approximately ten times lower (ESI Figure S7). Modeling demonstrates that both labeling sites are located near the taxane binding pocket inside lumen of microtubule, as illustrated in **Figure 2D**, Figure S11 and Movie 1.

Surprisingly, two minor labeling sites were also identified on α-tubulin (Cys347 and Cys376) despite cabazitaxel is known to bind β-tubulin. It might be explained by the close proximity of α and β subunit in microtubules. Nevertheless, the precursor abundance for the corresponding peptides was 20 times lower than for the major labeling site on β-tubulin, suggesting a really low labeling efficiency at these two positions on α-tubulin (ESI Figure S7).

Despite this experiment was done with porcine tubulin, there is high structural homology of tubulin among species.^28^ The sequences of the different α and β-tubulin isotypes are highly conserved between human and pig as shown by sequence alignments and crystal structures (ESI Figures S9-11). All the detected labeled peptides can be also found in human tubulin, including the main labeled peptide (AV<u>C</u>DIPPRGL), which is highly conserved. This suggests that analogous labeling might occur in other organisms and we can expect similar labeling sites using human cell line for living cells experiments.

Overall, the *in vitro* tubulin labeling study highlighted the great potential of probes **1** and **2** for tubulin labeling in living cells. We have also shown that Cys356, which is nearby the taxane binding site on β-tubulin, is the main labeling site of the probe.

### Tubulin labeling in living cells

Probe cytotoxicity was first evaluated on HeLa CCL cells (ESI Figure S12, Table S3). After 24h, most of the probes results in toxicity in the nanomolar range, with cytotoxicity thresholds ranging from 31.25 to 250 nM. These values are consistent with the toxicity of previously reported tubulin probe **SiR-CTX** (cytotoxicity threshold : 62.5 nM).^24^ Taxane derivatives like cabazitaxel are known to stabilize microtubules and inhibit cell proliferation by blocking the cell cycle just before mitosis.^29^ This leads to apoptosis and an increase in SubG1 phase population of cells. We hypothesize that this measurement could be useful to assess and compare the cell permeability of the probes. According to the obtained values, *ortho* isomers are more permeable compared to *meta* and *para* isomers. Once more, probe **5** is an exception and seems to be less cell-permeable than other *ortho* isomers. PEG linker increases the solubility while reducing the cell permeability, by pushing the equilibrium towards the open form. It is consistent with the spectroscopic data showing that for this compound, most of probes are in zwitterionic form in aqueous media.

To find out the best probe for fluorescence microscopy, a mini screening was performed on three different cancerous and non-cancerous human cell lines (dermal fibroblasts, HeLa CCL and U-2 OS) in living cells (**Figure 3 A,B** & ESI Figures S13-14). With U-2 OS cells, verapamil was added to inhibit efflux pumps.^30^ For probes **4** and **5**, the staining was weak, which is in accordance with the lower cell permeability suggested by cytotoxicity experiments. Probes **1**,**3** and **6** stained microtubules in living fibroblasts but only a weak staining was obtained with U-2 OS and HeLa. For all conditions tested, probe **2** (**6-SiR-*o*-C_9_-CTX**) was the best microtubules stain. Next step was to check whether the bond between the protein and the dye is covalent. After staining various cell lines with **6-SiR-*o*-C_9_-CTX**, cells were lysed (**Figure 3C**) and lysates were analyzed by SDS-PAGE. *In-gel* fluorescence image revealed the presence of a strong fluorescent band at 55 kDa for **6-SiR-*o*-C_9_-CTX** in the three cell lines tested, whereas nothing was observed for the non-covalent probe **SiR-CTX**. Western blot analysis confirmed that the fluorescent band corresponds to β-tubulin. Additional minor bands with masses exceeding 55 kDa were also detected, which we hypothesize could represent different β-tubulin isotypes and isoforms, and post-translational modifications of them.^28^

**Figure 3.**
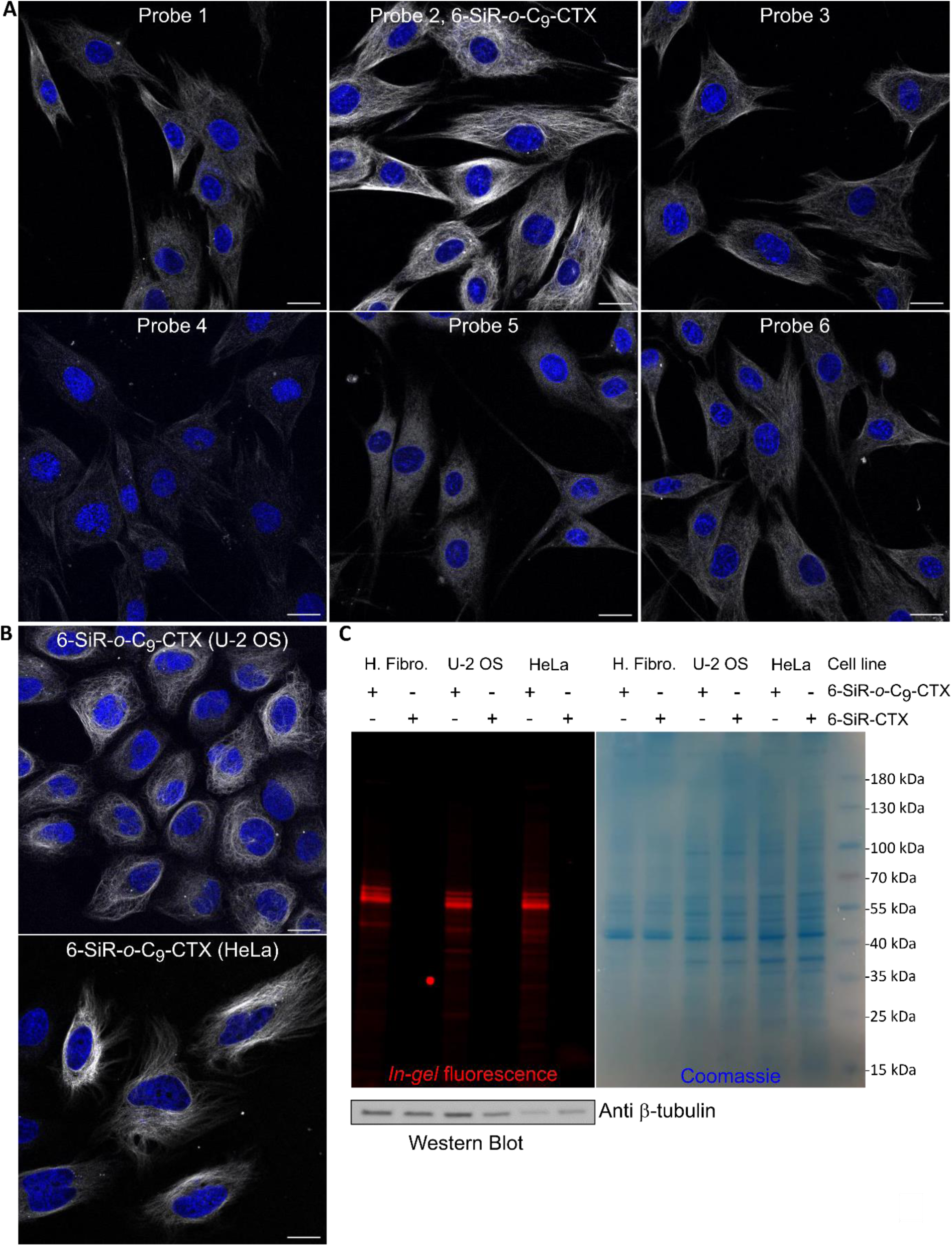
Tubulin labeling in living cells. (**A**) Live human dermal fibroblasts stained with probes (1 µM in OptiMEM) and Hoechst 33342 nucleic acid dye (1 µg/mL) for 4h. Confocal microscopy images were acquired using LEICA SP8. Gray channel (λ_ex_= 633 nm, λ_em_= 650-710 nm) corresponds to probe staining and blue channel (λ_ex_= 405 nm, λ_em_= 415-480 nm) corresponds to Hoechst 33342 staining. Scale bar = 20 μm. (**B**) Live-cell imaging of U-2 OS and HeLa CCL cells stained with **6-SiR-*o*-C_9_-CTX** (1 µM in OptiMEM) and Hoechst 33342 (1 µg/mL) for 4h (in case of U-2 OS, verapamil 10 µM was also added). Cells were washed three times with HBSS and imaged in DMEM+. Confocal microscopy images were acquired using LEICA SP8. Gray channel (λ_ex_= 633 nm, λ_em_= 650-710 nm) corresponds to probe staining and blue channel (λ_ex_= 405 nm, λ_em_= 415-480 nm) corresponds to Hoechst 33342 staining. Scale bar = 20 μm. (**C**) Analysis of cell lysates (human fibroblasts (H. Fibro.), U-2 OS and HeLa CCL) using SDS-PAGE and Western blot. Predominant single fluorescent band demonstrates selective β-tubulin labeling with **6-SiR-*o*-C_9_-CTX** (3 µM for 1h) in living cells.

The confirmation that the bond is covalent for **6-SiR-*o*-C_9_-CTX** and not for **SiR-CTX** was further obtained by washing experiments (**Figure 4**). Living U-2 OS cells were stained with either **6-SiR-*o*-C_9_-CTX** or **SiR-CTX** at the same concentration in presence of verapamil. After 4h of incubation, extensive washing was performed. Images acquired before the washing process show a nice microtubule staining for both probes. After washing, the staining performed with **6-SiR-*o*-C_9_-CTX** was still clearly visible without any significant loss of intensity, whereas for cells treated with **SiR-CTX**, the staining was not visible anymore, meaning the probe was fully washed out.

**Figure 4.**
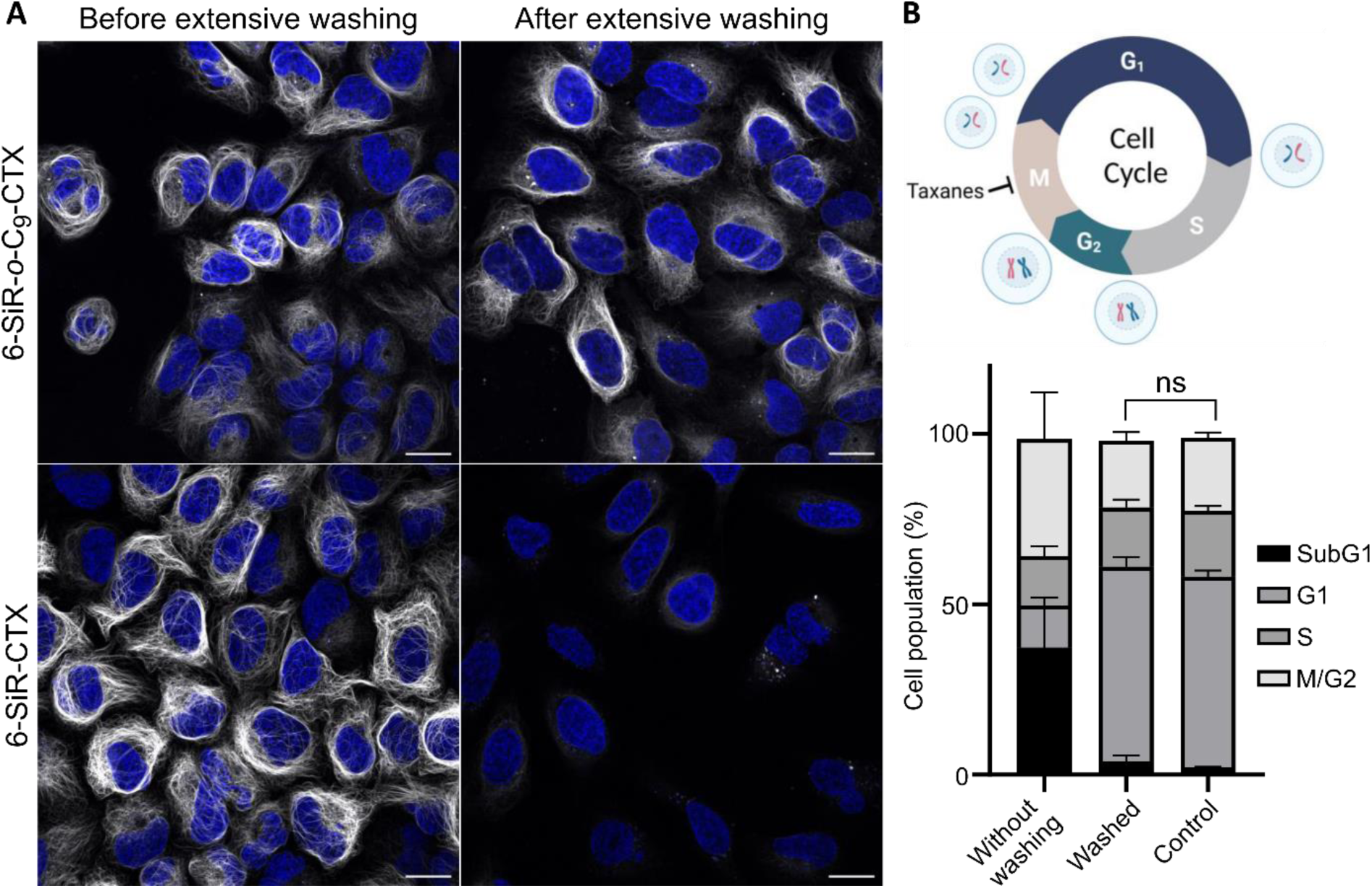
Cleavable linker allows covalent tubulin labeling and prevents taxane cytotoxicity. (**A**) Live-cell imaging of U-2 OS stained with probe (**6-SiR-*o*-C_9_-CTX** – first row, **SiR-CTX** – second row, 1 µM in OptiMEM), Hoechst 33342 (1 µg/mL) and verapamil (10 µM) for 4h before and after extensive washing (every 10 min over 2h). Note, signal intensity is enchanted 2-folds for **6-SiR-*o*-C_9_-CTX** (both before and after washing). Confocal microscopy images were acquired using LEICA SP8. Gray channel (λ_ex_= 633 nm, λ_em_= 650-710 nm) corresponds to probe staining and blue channel (λ_ex_= 405 nm, λ_em_= 415-480 nm) corresponds to Hoechst 33342 staining. Scale bar = 20 μm. The experiment was reproduced three times with similar results. (**B**) Cell cycle perturbation induced by **6-SiR-*o*-C_9_-CTX** (1 µM in OptiMEM) after 24h with or without extensive washing after 4h incubation. Results are averages of four independent experiments (N=4) and presented as means with standard deviations. Cell cycle diagram created with BioRender.com.

We investigated cytotoxicity of **6-SiR-*o*-C_9_-CTX** probe 24 hours after staining cells, with or without washing after 4h incubation. In non-washed cells, a significant cytotoxic effect was observed, with approximately 40% of the SubG1 phase population. However, for extensively washed cells, no significant cytotoxicity was observed compared to the control (**Figure 4B**, ESI Figure S15). These results demonstrate that extensive washing effectively removes the ligand released during proximity-induced labeling, recovering cells from taxane induced phenotype and leaving labeled microtubules inside living cells.

Finally, we wanted to investigate if this new covalent probe for tubulin, **6-SiR-*o*-C_9_-CTX**, was suitable for super-resolution microscopy and in particular stimulated emission depletion (STED) microscopy. The fluorescence of SiR can be inhibited using a 775 nm depletion laser, leading to an enhanced resolution. We stained three different cell lines (human dermal fibroblasts, U-2 OS and HeLa) with **6- SiR-*o*-C_9_-CTX** and successfully acquired STED images of living cells or after fixation with glutaraldehyde (**Figure 5**). We obtained microtubule apparent FWHM 38 ± 9 nm for fixed fibroblasts and 49 ± 13 nm living fibroblasts, which corresponds to a 8-9-fold resolution enhancement compared to confocal (**Figure 5D**, ESI Figure S16). In addition, the probe was photostable enough to acquire a 500 seconds timelapse (20 frames, Movie 2).

**Figure 5.**
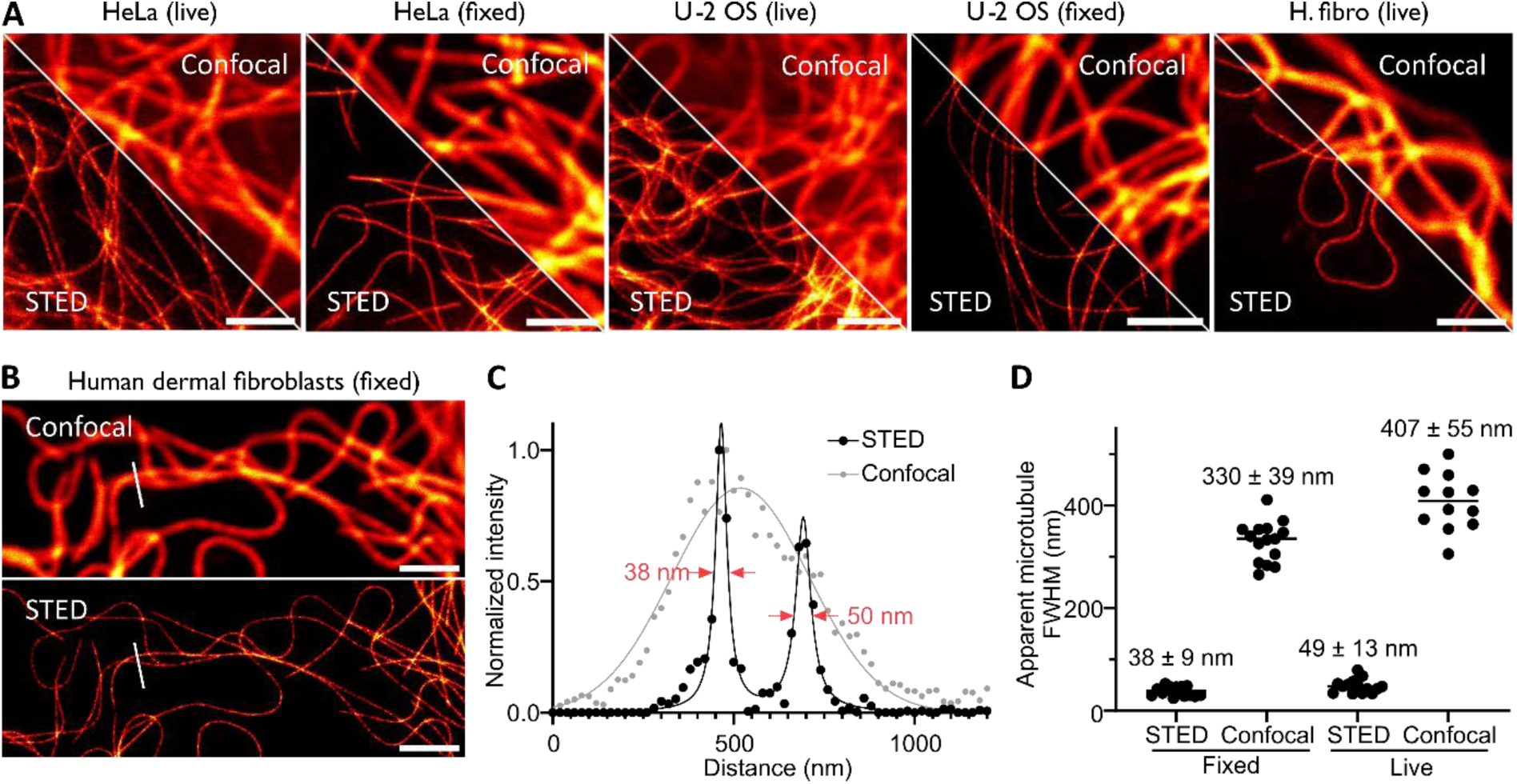
Nanoscopy of covalently labeled tubulin in living and fixed cells. (**A**) Confocal and STED images of microtubules in HeLa, U-2 OS or human dermal fibroblast (H. fibro) cells stained with **6-SiR- *o*-C_9_-CTX** (1 µM in OptiMEM) for 4h. Images show excellent tubulin labeling in live and glutaraldehyde fixed cells. Images acquired with Abberior Expert Line. Scale bar = 2 µm. (**B**) Confocal and STED image of microtubules in glutaraldehyde fixed human dermal fibroblasts stained before fixation with **6-SiR- *o*-C_9_-CTX** (1 µM in OptiMEM) for 4h. (**C**) Line profile plots of fluorescence signal of the area marked in images from (**B**). (**D**) Quantitative analysis of apparent microtubules FWHM in human dermal fibroblasts live or fixed. The line corresponds to the mean and each dot corresponds to the result of one single measurement, N ≥ 12 from three different fields of view.

## Discussion

We developed a series of covalent fluorescent probes for endogenous tubulin based on proximity induced reactivity. To do so, biocompatible benzyl sulfoniums were used as cleavable linkers. Probes were obtained thanks to a highly convergent synthetic path, which will allow this strategy to be easily extended to other dyes and other ligands in the near future. *Ortho* isomers showed enhanced photophysical properties, reactivity and cell permability over *meta* and *para* isomers. *Ortho*-benzyl sulfonium offers a good balance between stability and reactivity: they posses a similar labeling rate as LDAI, but can be used inside living cells because they are stable enough. We demonstrated that **6-SiR- *o*-C_9_-CTX** can be used to covalently label endogenous tubulin in living cells after only few hours and that this probe is suitable for STED nanoscopy. Probe is likely to be useful for labeling of microtubules in multiple species because the labeling site is mapped to highly conserved sequence AV<u>C</u>DIPPR of β- tubulin. Introduction of cleavable linker generated tubulin probe that, after labeling releases cytotoxic taxane, which can be washed out and has a minimal impact on microtubule structure and function. Overall, this work underscores the immense potential of ligand-directed labeling and sulfonium chemistry for the covalent labeling of endogenous proteins. Our methodology offers new opportunities for advanced imaging techniques like nanoscopy and sets the stage for broader applications in cellular biology and beyond.

## Methods

### General procedure for the final step of probe synthesis

In a 1.5 mL Eppendorf, **6-SiR-COOH (1.0 eq.)** was solubilized in **dry DMSO (0.1 M)**, and **DIPEA (10 eq.)** was added. After 5 min**, TSTU (1.05 eq.)** was added. After 10 min, the formation of activated acid was controlled by LC-MS and the solution of **amine-bearing sulfonium in DMSO (1.3 eq.) –** *obtained at the previous step* - was added. The formation of the amide was controlled by LC-MS (it typically takes 15- 20 min). Once the amide bond was formed, **TSTU (1.5 eq.)** was added, followed by **CTX-NH_3_.CF_3_OCO_2_ (3.0 eq.)** and **DIPEA (10 eq.)**. The formation of the final product was controlled by LC-MS. The reaction was quenched by adding **trifluoroacetic acid (50 µL)**. The crude was diluted in **ACN (1 mL)** and purified by reverse-phase HPLC (Device A (see ESI), H_2_O+0.1%TFA/ACN, linear gradient 70:30 to 0:100, 40 mL/min). The fractions containing the product were lyophilized and the yields were determined as followed:

The fluorescent probes were dissolved in a precise amount of DMSO-d_6_ (600 µL) and were transferred to an NMR tube to obtain ^1^H spectra. Afterwards the contents of the NMR tubes were transferred to an Eppendorf and were considered as stock solution. Two 1 µL samples were taken from stock solution and were diluted in Eppendorf with 59 µL of PBS containing 0.1% SDS (60-fold dilution). After 15 min, absorption of 2 µL of the diluted samples were measured on nanodrop (Nanodrop 1000, Peqlab) with 1 mm optical path. The measured absorption intensity values at the dyes absorption maxima value were averaged and concentration of the stock solution was determined according to the equation (considering that the molar absorption coefficient of the probe is comparable to that of 6-SiR-COOH ɛ(λ_MAX_) = 90 000 L.mol^-1^.cm^-1^)^23^:

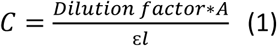

where C is the concentration of stock solution; A the sample absorption, ɛ the extinction coefficient of the dye in PBS containing 0.1% SDS and l the path length.

Once the concentration of the stock solution was measured the amount of the obtained fluorescent conjugate could be calculated by following equation:

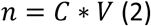

where C is the concentration of stock solution and V the volume of stock solution. Finally, the yield could be determined by the classical equation:

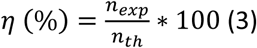

where n_exp_ is the obtained amount of the isolated product (mol) and n_th_ – maximal theoretical amount of the product in the reaction (mol).

Note, the final probes of ortho isomer are not fully stable under the reaction conditions. Therefore, it is better to stop the reaction after 30-45 min even if the conversion is not full.

### Measurement of labeling kinetics

Tubulin from porcine brain (1 mg, Cytoskeleton, #T240, >99% Pure) was solubilized in 100 µL General Tubulin Buffer (Cytoskeleton, #BST01) supplemented with 1 mM of GTP (Thermo Fisher, #R0461) to get a 10 mg/mL solution. This solution was aliquoted, samples were snap-frozen in liquid nitrogen and stored at -80°C prior to use.

Before the experiment, an aliquot was thawed using a water bath at room temperature. Once liquid, the sample was immediately placed on ice. In parallel, a solution of General Tubulin Buffer containing 1 mM of GTP was prepared and placed on ice.

The reaction was conducted mixing tubulin (0.5 mg/mL ̴ 4.5 µM) and the probe (20 µM) in General Tubulin Buffer containing 1mM of GTP at 37°C for 48h. Aliquots were taken at different timepoints, immediately mixed with 1/3 volume of 4x SDS sample buffer (Tris-HCl pH=6.8: 0.2M, SDS: 8.0% (w/v), Bromophenol Blue: 0.6 mM, glycerol: 5.4 M + 50 µL of β-mercaptoethanol per mL of solution prior to use), and boiled for 5 min at 95°C.

The different samples were loaded on 4-15% Mini-PROTEAN® TGX™ Precast Protein Gels (Biorad, #4561086). After electrophoresis in Mini-PROTEAN® Tetra Cell using SDS-PAGE running buffer (0.25 M Tris HCl, 1.92 M glycine and 1% (w/v) sodium dodecyl sulfate (SDS) pH=8.3), fluorescence images were recorded using Amersham Imager 600 RGB. Quantitative data analysis was then performed using the “gel analysis” function of Fiji^7^.

All experiments were performed in triplicate on different days. A gel was performed with the 48h timepoints (3 for each compound) for all the compounds to calibrate the values accordingly. Finally, all data were calibrated with **6-SiR-*o*-C_9_-CTX** for which the DOL was found to be 0.94 (see below).

Considering one reactant was in excess, data obtained were fitted using a single exponential (one-phase association model from GraphPad Prism- Y=Y_0_ + (Plateau-Y_0_)*(1-exp(-K*x)), imposing the constraint Y_0_=0) to obtain the degree-of-labeling and the reaction rate for each compound.

### Sample preparation for determination of DOL (degree-of-labeling)

**6-SiR-*o*-C_9_-CTX** (20 µM) and Tubulin (1 mg, 0.5 mg/mL ̴ 4.5 µM, Cytoskeleton, #T240, >99% Pure) were mixed together in General Tubulin Buffer (Cytoskeleton, #BST01) supplemented with 1 mM of GTP (Thermo Fisher, #R0461) for 24h at 37°C. The solution was cooled at 4°C for 1h to depolymerize tubulin. The probe that did not react was then removed using PD MidiTrap G-25® (Cytivia). Procedure used was the one advised by the supplier, using BRB80 (80 mM PIPES pH=6.8, 1 mM EGTA, 1 mM MgCl_2_) as eluting buffer and conducted at 4°C. The solution containing the protein was concentrated to the desire volume using Vivaspin® Turbo 4 (Sartorius, MWCO = 10 kDa, #VS04T02).

DOL was determined measuring A_280nm_ and A_652nm_ with a NanoDrop ND-1000 spectrophotometer (Peqlab) to determine dye and protein concentrations to calculate DOL according to this formula:

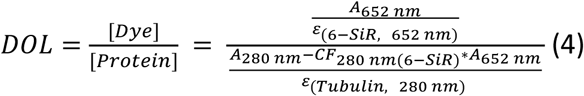

A solution of 2% SDS (0.5 µL) was added to the solution of protein (5 µL) to ensure that the dye was fully open (three different solutions were prepared). After 30 min at room temperature, absorbance was measured at 280 nm and 652 nm. Considering ɛ(6-SiR, 652 nm) = 90 000 L.mol^-1^.cm^-1^, ɛ(Tubulin, 280 nm) = 110 000 L.mol^-1^.cm^-1^ and CF_280_(6-SiR) = 0.147, the DOL for this sample was found to be 0.94.

### Proteomic study

#### Sample preparation

One µg of the **6-SiR-*o*-C_9_-CTX**-labeled tubulin (as described above) was separated on 4-15% Mini- PROTEAN® TGX™ Precast Protein Gels (Biorad, #4561086) and stained with Coomassie brilliant blue. A protein band corresponding to a SiR-labeled tubulin was excised from the gel, washed, reduced with dithiothreitol (DTT), alkylated with iodoacetamide and digested with trypsin (sequencing grade, Promega) overnight. The resulting peptides were extracted, dried in a SpeedVac vacuum concentrators (Thermo Scientific) and dissolved in 2% acetonitrile/0.05% trifluoroacetic acid (v:v).

#### Data acquisition

Peptides were analyzed by electrospray ionization mass spectrometry in a Thermo Orbitrap Exploris 480 mass spectrometer coupled to an UltiMate3000 ultrahigh performance liquid chromatography system (Thermo Scientific). Chromatographic separation was performed with an in-house packed C18 reverse-phase column (75 µm ID × 300 mm, Reprosil-Pur 120 C18-AQ, 3 μm, Dr. Maisch) using 0.1% formic acid as solvent A and 80% acetonitrile / 0.08% formic acid as solvent B. Separating part of the HPLC method included 3 steps of linear gradients: (1) 12-42%B over 40 min, (2) 42-65%B over 27 min and (3) 65-95%B over 7.1 min. Mass spectrometer was equipped with a Nanospray Flex Ion source and controlled by Thermo Scientific Xcalibur 4.4 and Thermo Exploris 480 3.0 software. Data were acquired using an 88-min Top30 data-dependent acquisition method. One full MS scan across the 350–1600 m/z range was acquired at a resolution of 120000, with an AGC target of 300% and a maximum fill time of 25 ms. Precursors with charge states 2–6 above a 1e4 intensity threshold were then sequentially selected using isolation window of 1.6 m/z, fragmented with nitrogen at a normalized collision energy setting of 28%, and the resulting MS2 spectra recorded at a resolution of 30000, AGC targets of 75% and a maximum fill time of 50 ms. Dynamic exclusion of precursors was set to 22 s.

#### Data processing

Proteins and sites of SiR-labeling were identified with Proteome Discoverer 3.1.1.93 using SequestHT as a search engine. For this, Thermo raw files were searched against a database that included sequences of Sus scrofa UniProt proteome (release 24-01-2024) and common contaminants observed in MS experiments. Two missed cleavages were allowed. Protein N-terminal acetylation, M-oxidation, C-carbamidomethylation as well as C/K+573.2442 were set as variable modifications.

### Cell cycle analysis by imaging cytometry

The probes were dissolved in DMSO (Sigma Aldrich, #900645-4x2mL) at 500 – 2000-fold stock concentration and added to culture media of cells at 500 – 2000-fold dilution accordingly. In parallel, the appropriate DMSO control samples were prepared by adding corresponding amount of DMSO volume to the separate well. HeLa cells were grown in 6-well plates (∼250,000 cells per well) in presence of the fluorescent probe in variable concentrations for 24 h at 37°C in humidified incubator with 5% CO_2_. Cells were processed according to the NucleoCounter® NC-3000™ two-step cell cycle analysis protocol for cells attached to T-flasks, cell culture plates or micro-carriers. In particular, the 250 µL lysis solution (Solution 10, Chemometec Cat. No. 910-3010) supplemented with 10 µg/ml DAPI (Solution 12, Chemometec Cat. No. 910-3012) was used per well, incubated at 37 °C for 5 min. Then 250 µL of stabilization solution (Solution 11, Chemometec Cat. No. 910-3011) was added. Cells were counted on a NucleoCounter® NC-3000™ in NC-Slide A2™ slides (Chemometec, Cat. No. 942-0001) loaded with ∼30 µL of each of the cell suspensions into the chambers of the slide. Each time, ∼10,000 cells in total were measured, and the obtained cell cycle histograms were analyzed with ChemoMetec NucleoView NC-3000 software, version 2.1.25.8. All experiments were repeated three times (with cells from different passages) and the results are presented as means with standard deviations.

### Cytotoxicity experiment after washing

*This protocol refers to Figure 4B.* U-2 OS cells were seeded in 12-well plates (∼300,000 cells per well) 24h to 48h prior to the experiment. Cells were incubated in a humidified 5% CO_2_ incubator at 37 °C. Two different conditions were tested: first cells were incubated for 24h in presence of OptiMEM containing 1 µM of **Probe 2** and 10 µM of verapamil. For the second condition, cells were incubated 4h in presence of OptiMEM containing 1 µM of **Probe 2** and 10 µM of verapamil, washed briefly 4 times with HBSS and then washed 10 times over 2h with DMEM+, and then further incubate with DMEM+ for 18h. Cytotoxicity experiments were then proceed as described above. This experiment was reproduced 4 times (N=4) on different days with cells from different passages. The results are presented as means with standard deviations.

### Western-Blot

Confluent cells in a 6-well plate were incubated in presence of OptiMEM (Thermo Fisher, #11058021) supplemented with probe (3 µM) for 1h at 37°C in humidified incubator with 5% CO_2_. The media was removed and cells were washed twice with HBSS. CelLytic™ M (300 µL per well, Sigma-Aldrich #C2978) was added and the plate was put on a shaker for 30 min. Cell lysates were collected in Eppendorf and were centrifugated at 15 000 g and 4°C for 20 min. Supernatants were collected and mixed with 1/3 volume of 4x SDS sample buffer (Tris-HCl pH=6.8: 0.2M, SDS: 8.0% (w/v), Bromophenol blue: 0.6 mM, Glycerol: 5.4 M + 50 µL of β-mercaptoethanol per mL of solution prior to use), and boiled for 5 min at 95°C. The different samples were loaded on 4-15% Mini-PROTEAN® TGX™ Precast Protein Gels (Biorad, #4561086). After electrophoresis in Mini-PROTEAN® Tetra Cell using SDS-Page running buffer (0.25 M Tris HCl, 1.92 M Glycine and 1% (w/v) Sodium Dodecyl Sulfate (SDS) pH=8.3), proteins were transferred from the gel to a PVDF-membrane (iBlot™ 2 Transfer Stacks, PVDF, regular size, # IB24001) using iBlot 2 Gel Transfer Device (P0 method from the supplier was used: 20V for 1min, 23V for 4 min, 25 V for 2 min).

Membrane was then blocked with 1% BSA in PBS containing 0.1% of Tween 20 (Blocking buffer) overnight at 4°C. Primary antibody (Rabbit Recombinant Monoclonal Beta-3-tubulin antibody, Abcam, #ab52623) was added (1/2000, v:v). After 1h at room temperature, the membrane was washed three times 10 min with PBS +0.1% Tween 20. Membrane was incubated for 1h at room temperature in presence of a solution of secondary antibody (Donkey anti-Rabbit IgG (H+L) Highly Cross-Adsorbed Secondary Antibody, Alexa Fluor™ Plus 488, Thermo Fisher, #A32790) in blocking buffer (1/1000, v:v). After a brief washing step with PBS+0.1% Tween 20, fluorescence images were recorded using Amersham Imager 600 RGB.

### Sample preparation for live-cell imaging

Cells were seeded on µ-Slide 8 Well Glass Bottom dishes (Ibidi, #80827) 24 to 48h prior to imaging. Cells were washed four times with HBSS (Gibco, #14025) to remove FBS and incubated with OptiMEM (Thermo Fisher, #11058021) containing 1 µM of probe in a humidified 5% CO_2_ incubator at 37 °C. After 2h or 4h (depending on the experiment), the media supplemented with probe was removed, cells were washed three times with HBSS and imaged in DMEM+ [high-glucose DMEM (Thermo Fisher, #31053044) with 10% FBS (Thermo Fisher, #10082147) supplemented with 1 mM Sodium pyruvate (Sigma, #S8636), 1% GlutaMax (Thermo Fisher, #35050038) and 1% Penicillin-Streptomycin (Sigma, #P0781)].

### Sample preparation for fixed cells imaging

Protocol adapted from R. Gerasimaite *et al*., 2021.^31^ Cells were seeded on µ-Slide 8 Well Glass Bottom dishes (Ibidi, #80827) 24 to 48h prior to imaging. Cells were washed 4 times with HBSS (Gibco, #14025) to remove FBS and incubated with OptiMEM (Thermo Fisher, #11058021) containing 1 µM of probe in a humidified 5% CO_2_ incubator at 37 °C. After 2h or 4h (depending on the experiment), the media supplemented with probe was removed, cells were washed 4 times with 200 µL of PEMP (100 mM PIPES pH 6.8, 1 mM EGTA, 1 mM MgCl_2_ + 4% PEG8000), permeabilized 90s with 0.5% Triton X-100 in PEM (PEMP without PEG8000), and washed again 4 times with 200 µL of PEMP. Then, cells were incubated with 200 μL of 0.2% glutaraldehyde in PEM for 15 min, followed by 200 μL of 2 mg/mL NaBH_4_ in PEM (dissolved immediately before use) for another 15 min. The samples were washed 4× with 200 μL of PEM and imaged in glycerol buffer (GB, 10 mM Na-PO_4_, pH 6.8, 1mM EGTA, 6 mM MgCl_2_, 3.4 M glycerol).

### Live cell imaging with or without extensive washing

*This protocol refers to Figure 4A.* U-2 OS were seeded on µ-Slide 8 Well Glass Bottom dishes (Ibidi, #80827) 24 to 48h prior to imaging. Cells were washed 4 times with HBSS (Gibco, #14025) to remove FBS and incubated with OptiMEM (Thermo Fisher, #11058021) containing 10 µM of verapamil and 1 µM of **6-SiR-*o*-C_9_-CTX** or 1 µM of **SiR-CTX** in a humidified 5% CO_2_ incubator at 37 °C for 4h. Cells were washed briefly 4 times with HBSS, media was replaced by DMEM+ and cells were imaged without further washing (it corresponds to the conditions “Before extensive washing”). The other condition “After extensive washing” refers to cells that were additionally washed 10 times over 2h with DMEM+ after the brief washing with HBSS, and then imaged in DMEM+. This experiment was reproduced three times on different days with cells from different passages with similar results.

### Confocal/STED microscopes and imaging parameters

Confocal and STED images were acquired using Abberior STED Facility Line scanning (Abberior Instruments GmbH) or TCS SP8 (Leica) microscopes. STED images were acquired using Abberior STED Facility Line scanning (Abberior Instruments GmbH) microscopes. Imaging parameters are summarized in Table S4.

TCS SP8 confocal microscope is equipped with 405, 458, 476, 488, 496, 514, 561 and 633 nm excitation lasers as well as HC PL APO CS2 63x/1.40 Oil objective (Leica). Microscope has three Hybrid and two PMT detectors which can be tuned to any detection window in the range 400 – 800 nm. Probes were excited at 633 nm (Laser power: 1%) and emission was collected between 650 and 710 nm.

Abberior STED Facility Line is equipped with 488, 515, 561, 640 and 700 nm 40 MHz pulsed excitation lasers, a pulsed 775 nm 40 MHz 3W STED laser, and an UPlanSApo 60x/1.40 Oil objective. Microscope has two APD and two MATRIX detectors which can be tuned to any detection window in the range 400 – 800 nm. Pixel size was 30 nm in the xy plane was used for 2D STED images and 80 nm in the xy plane for large field of view images. Laser powers were optimized for each sample.

Abberior STED Expert Line equipped with 561 nm and 640 nm 40 MHz pulsed excitation lasers, a pulsed 775 nm 40 MHz 3W STED laser, and an UPlanSApo 100x/1.40 Oil objective. The following detection windows were used: for the SiR channel 685 / 70 nm. Pixel size was 20 nm in the xy plane was used for 2D STED images. Laser powers were optimized for each sample.

### Visualization and modeling of the tubulin cryo-EM structures and models

Cryo-EM structures of pig tubulin complex with Taxol (PDB: 5SYF)^32^ and human (PDB: 6E7C)^33^ tubulin were downloaded from Protein Data Bank repository and visualized using Swiss-PdbViewer (v 4.1) or UCSF ChimeraX (v 1.6)^34^. To identify Taxol binding pocket in human protein, both structures were superimposed using iterative fit function (backbone atoms only) of the program.

### Processing and visualization of the acquired images

All acquired images were processed and visualized using Fiji^35^. Line profiles were measured using the “straight line” tool with the line width set to 3 pixels. To define apparent microtubule FWHM, line profiles were fitted with Gaussian and Lorentzian distributions for confocal and STED images respectively.

### Statistical tests

Comparisons were performed using unpaired t-test (two-tailed) in GraphPad Prism 10.4 (GraphPad Software, Inc., San Diego, CA, USA) and *p*-values ≤ 0.05 were considered statistically significant. All microscopy imaging experiments were repeated at least two times on different non-consecutive days (n ≥ 2). Multiple fields of view (n ≥ 3) were acquired during each imaging session and representative images are shown in the figures.

## Acknowledgements

The authors thank the Max Planck Society for supporting this work. The authors are grateful to Dr. Vladimir Belov, Jan Seikowski, Jens Schimpfhauser and Jürgen Bienert for the NMR measurements of numerous probes and the central analytics’ team (Institute for Organic and Biomolecular Chemistry, Georg-August University, Göttingen) for acquiring HRMS. Figures 2D and S11A were performed with UCSF ChimeraX, developed by the Resource for Biocomputing, Visualization, and Informatics at the University of California, San Francisco, with support from National Institutes of Health R01-GM129325 and the Office of Cyber Infrastructure and Computational Biology, National Institute of Allergy and Infectious Diseases.

## Author contributions

M.A. and G.L. conceived and planned the study. M.A, T.K., O.D. and G.L. performed the experiments. M.A, T.K., O.D., H.U. and G.L. performed the data analysis. M.A. and G.L. wrote the initial draft; all authors contributed to the final version of the manuscript.

## References

1 Sahl, S. J., Hell, S. W. & Jakobs, S. Fluorescence nanoscopy in cell biology. Nat Rev Mol Cell Biol 18, 685–701, doi:10.1038/nrm.2017.71 (2017).

2 Lukinavičius, G. et al. Stimulated emission depletion microscopy. Nature Reviews Methods Primers 4, 56, doi:10.1038/s43586-024-00335-1 (2024).

3 D’Este, E., Lukinavičius, G., Lincoln, R., Opazo, F. & Fornasiero, E. F. Advancing cell biology with nanoscale fluorescence imaging: essential practical considerations. Trends Cell Biol 34, 671–684, doi:10.1016/j.tcb.2023.12.001 (2024).

4 Wilhelm, J. et al. Kinetic and Structural Characterization of the Self-Labeling Protein Tags HaloTag7, SNAP-tag, and CLIP-tag. Biochemistry 60, 2560–2575, doi:10.1021/acs.biochem.1c00258 (2021).

5 Hink, M. A. et al. Structural Dynamics of Green Fluorescent Protein Alone and Fused with a Single Chain Fv Protein*. Journal of Biological Chemistry 275, 17556–17560, 10.1074/jbc.M001348200 (2000).

6. 6 Fatti, E., Khawaja, S. & Weis, K. The dark side of fluorescent protein tagging – the impact of protein tags on biomolecular condensation. bioRxiv, 2024.2011.2023.624970, doi:10.1101/2024.11.23.624970 (2024).

7 Tsukiji, S., Miyagawa, M., Takaoka, Y., Tamura, T. & Hamachi, I. Ligand-directed tosyl chemistry for protein labeling in vivo. Nat Chem Biol 5, 341–343, doi:10.1038/nchembio.157 (2009).

8 Fujishima, S. H., Yasui, R., Miki, T., Ojida, A. & Hamachi, I. Ligand-directed acyl imidazole chemistry for labeling of membrane-bound proteins on live cells. J Am Chem Soc 134, 3961–3964, doi:10.1021/ja2108855 (2012).

9 Takaoka, Y., Nishikawa, Y., Hashimoto, Y., Sasaki, K. & Hamachi, I. Ligand-directed dibromophenyl benzoate chemistry for rapid and selective acylation of intracellular natural proteins. Chem Sci 6, 3217–3224, doi:10.1039/c5sc00190k (2015).

10 Matsuo, K., Nishikawa, Y., Masuda, M. & Hamachi, I. Live-Cell Protein Sulfonylation Based on Proximity-driven N-Sulfonyl Pyridone Chemistry. Angew Chem Int Ed Engl 57, 659–662, doi:10.1002/anie.201707972 (2018).

11 Tamura, T. et al. Rapid labelling and covalent inhibition of intracellular native proteins using ligand-directed N-acyl-N-alkyl sulfonamide. Nat Commun 9, 1870, doi:10.1038/s41467-018-04343-0 (2018).

12 Xin, X. et al. Ultrafast and selective labeling of endogenous proteins using affinity-based benzotriazole chemistry. Chem Sci 13, 7240–7246, doi:10.1039/d1sc05974b (2022).

13 Stoddart, L. A. et al. Ligand-directed covalent labelling of a GPCR with a fluorescent tag in live cells. Commun Biol 3, 722, doi:10.1038/s42003-020-01451-w (2020).

14 Comeo, E. et al. Ligand-Directed Labeling of the Adenosine A1 Receptor in Living Cells. Journal of Medicinal Chemistry 67, 12099–12117, doi:10.1021/acs.jmedchem.4c00835 (2024).

15 Yamaura, K., Kiyonaka, S., Numata, T., Inoue, R. & Hamachi, I. Discovery of allosteric modulators for GABAA receptors by ligand-directed chemistry. Nature Chemical Biology 12, 822–830, doi:10.1038/nchembio.2150 (2016).

16 Wakayama, S. et al. Chemical labelling for visualizing native AMPA receptors in live neurons. Nature Communications 8, 14850, doi:10.1038/ncomms14850 (2017).

17 Verdier-Pinard, P. et al. Tubulin proteomics: towards breaking the code. Anal Biochem 384, 197–206, doi:10.1016/j.ab.2008.09.020 (2009).

18 Fourel, G. & Boscheron, C. Tubulin mutations in neurodevelopmental disorders as a tool to decipher microtubule function. FEBS Letters 594, 3409–3438, 10.1002/1873-3468.13958 (2020).

19 Song, Y. & Brady, S. T. Post-translational modifications of tubulin: pathways to functional diversity of microtubules. Trends Cell Biol 25, 125–136, doi:10.1016/j.tcb.2014.10.004 (2015).

20 Wang, R. et al. Low-Toxicity Sulfonium-Based Probes for Cysteine-Specific Profiling in Live Cells. Anal Chem 94, 4366–4372, doi:10.1021/acs.analchem.1c05129 (2022).

21 Wang, Y. et al. A Peptide-Based Ligand-Directed Chemistry Enables Protein Functionalization. Org Lett 24, 7205–7209, doi:10.1021/acs.orglett.2c02974 (2022).

22 Lukinavičius, G. et al. A near-infrared fluorophore for live-cell super-resolution microscopy of cellular proteins. Nature Chemistry 5, 132–139, doi:10.1038/nchem.1546 (2013).

23 Bucevicius, J., Kostiuk, G., Gerasimaite, R., Gilat, T. & Lukinavicius, G. Enhancing the biocompatibility of rhodamine fluorescent probes by a neighbouring group effect. Chem Sci 11, 7313–7323, doi:10.1039/d0sc02154g (2020).

24 Lukinavičius, G. et al. Fluorescent dyes and probes for super-resolution microscopy of microtubules and tracheoles in living cells and tissues. Chem Sci 9, 3324–3334, doi:10.1039/c7sc05334g (2018).

25 Deng, F. et al. Multiple Factors Regulate the Spirocyclization Equilibrium of Si-Rhodamines. The Journal of Physical Chemistry B 124, 7467–7474, doi:10.1021/acs.jpcb.0c05642 (2020).

26 Grimm, J. B. et al. Optimized Red-Absorbing Dyes for Imaging and Sensing. J Am Chem Soc 145, 23000–23013, doi:10.1021/jacs.3c05273 (2023).

27 Tamura, T. & Hamachi, I. Chemistry for Covalent Modification of Endogenous/Native Proteins: From Test Tubes to Complex Biological Systems. Journal of the American Chemical Society 141, 2782–2799, doi:10.1021/jacs.8b11747 (2019).

28 Nsamba, E. T. & Gupta, M. L., Jr. Tubulin isotypes – functional insights from model organisms. Journal of Cell Science 135, jcs259539, doi:10.1242/jcs.259539 (2022).

29 Abal, M., Andreu, J. M. & Barasoain, I. Taxanes: Microtubule and Centrosome Targets, and Cell Cycle Dependent Mechanisms of Action. Current Cancer Drug Targets 3, 193–203, doi:10.2174/1568009033481967 (2003).

30 Gerasimaitė, R. et al. Efflux pump insensitive rhodamine–jasplakinolide conjugates for G- and F-actin imaging in living cells. Organic & Biomolecular Chemistry 18, 2929–2937, doi:10.1039/D0OB00369G (2020).

31 Gerasimaitė, R. et al. Blinking Fluorescent Probes for Tubulin Nanoscopy in Living and Fixed Cells. ACS Chem Biol 16, 2130–2136, doi:10.1021/acschembio.1c00538 (2021).

32 Kellogg, E. H. et al. Insights into the Distinct Mechanisms of Action of Taxane and Non-Taxane Microtubule Stabilizers from Cryo-EM Structures. J Mol Biol 429, 633–646, doi:10.1016/j.jmb.2017.01.001 (2017).

33 Ti, S. C., Alushin, G. M. & Kapoor, T. M. Human beta-Tubulin Isotypes Can Regulate Microtubule Protofilament Number and Stability. Dev Cell 47, 175–190 e175, doi:10.1016/j.devcel.2018.08.014 (2018).

34 Goddard, T. D. et al. UCSF ChimeraX: Meeting modern challenges in visualization and analysis. Protein Sci 27, 14–25, doi:10.1002/pro.3235 (2018).

35 Schindelin, J. et al. Fiji: an open-source platform for biological-image analysis. Nature Methods 9, 676–682, doi:10.1038/nmeth.2019 (2012).

