## Supplementary information for "Biocompatible sulfonium-based covalent probes for endogenous tubulin fluorescence nanoscopy in live and fixed cells"

### Supplementary figures

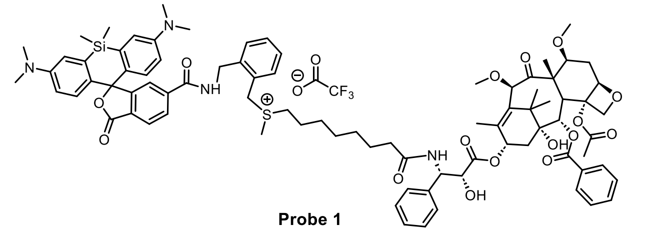

**Probe 1**

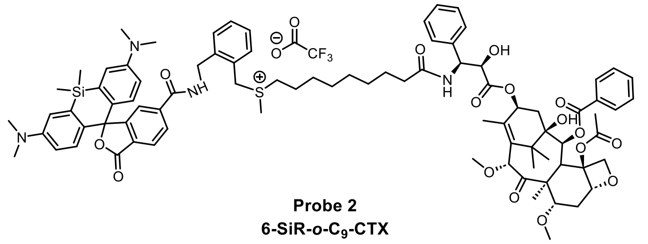

**Probe 2**
**6-SiR-*o*-C$_9$-CTX**

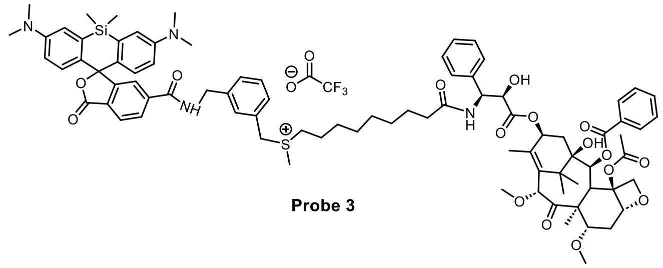

**Probe 3**

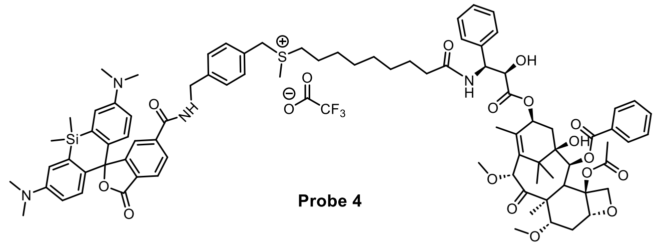

**Probe 4**

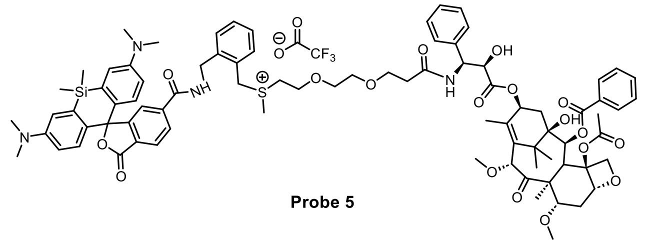

**Probe 5**

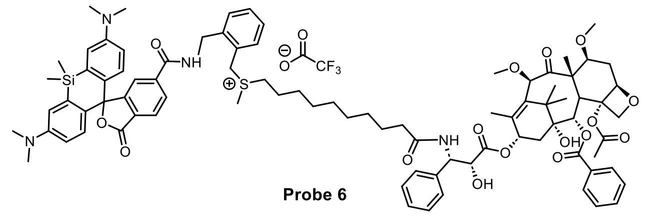

**Probe 6**

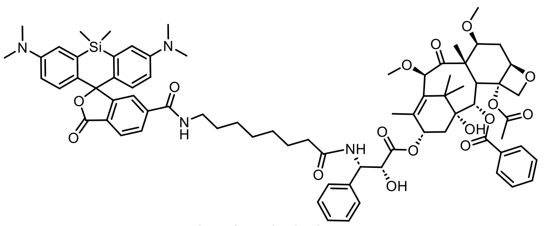

**SiR-CTX (ref. 1)**

**Figure S1. Detailed structures of the covalent probes and non-covalent tubulin probe SiR-CTX[1]**

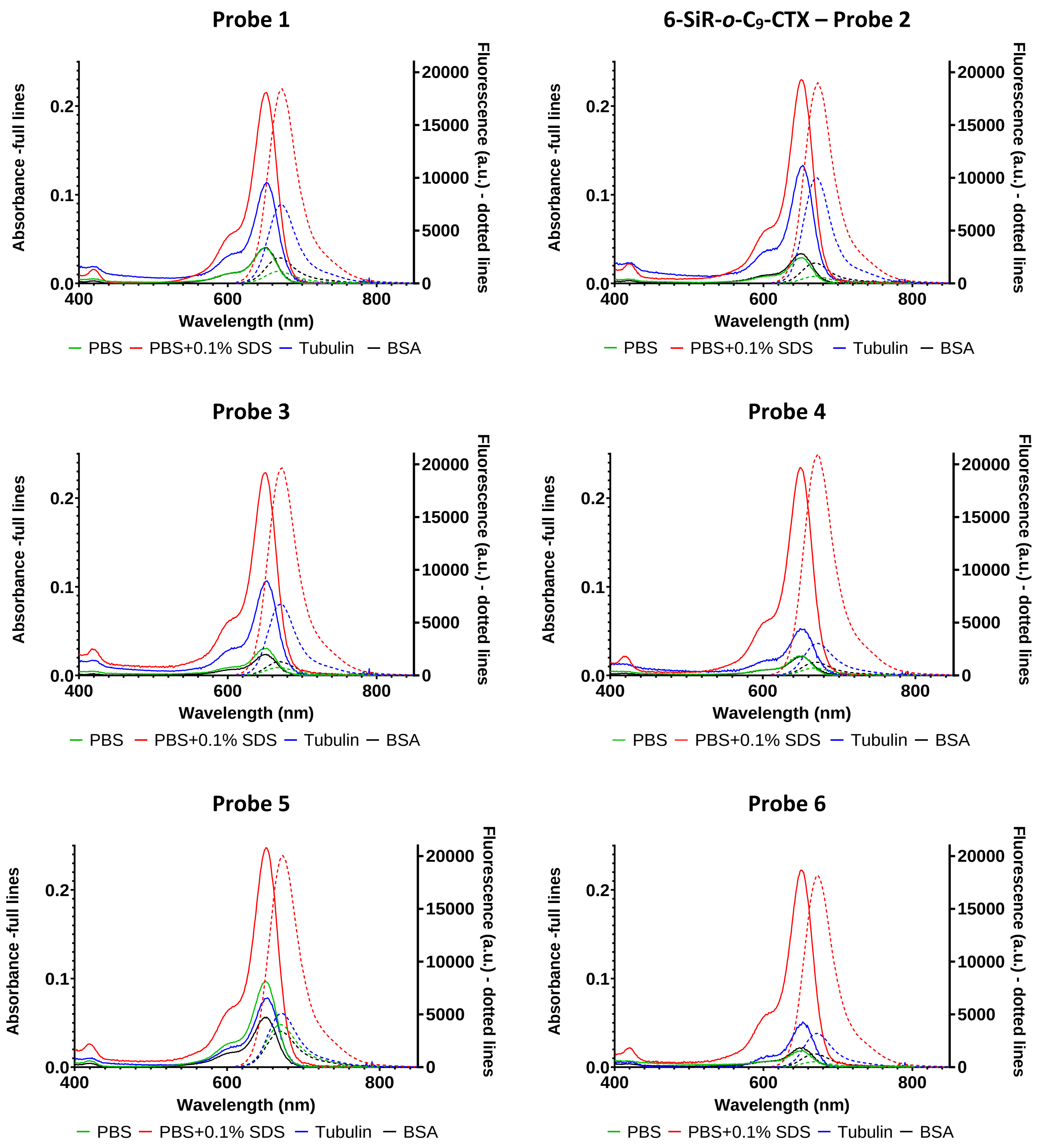

**Figure S2. Absorption and emission spectra of covalent tubulin probes.** Spectra were recorded after incubating 5 µM probes with 0.5 mg/mL tubulin in General Tubulin Buffer (Blue), 0.5 mg/mL BSA in PBS (Black), in PBS (Green), or in PBS+0.1% SDS (Red) at 37°C for 4h. Spectra are represented as averages of three independently repeated experiments (N=3).

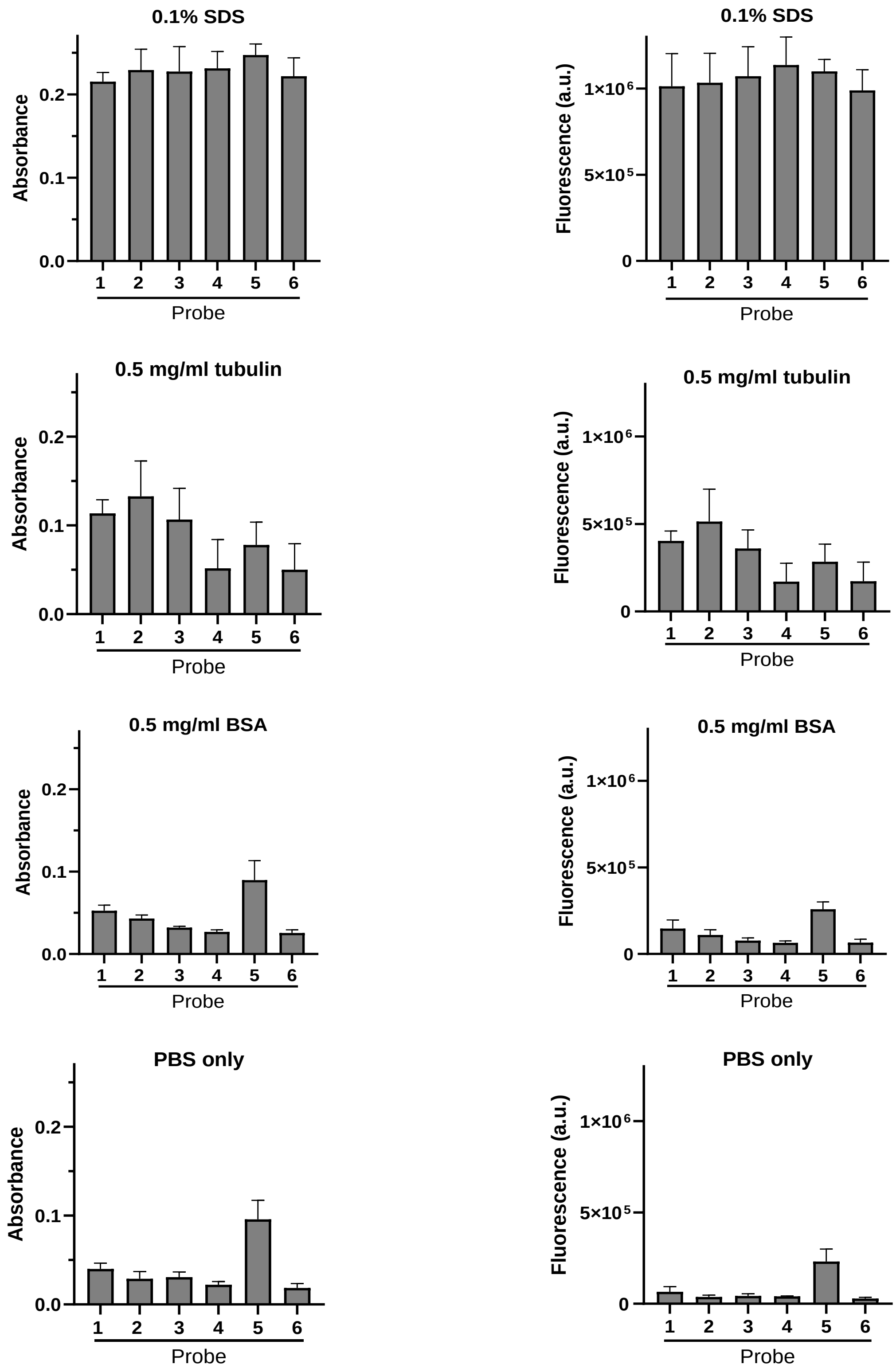

**Figure S3. Absorbance and fluorescence emission of covalent tubulin probes in different conditions.** Spectra were recorded after incubating 5 µM probes with 0.5 mg/mL tubulin in General Tubulin Buffer, 0.5 mg/mL BSA in PBS and in 0.1% SDS in PBS, only in PBS at 37°C for 4h. Absorbance at the maximum (652 nm) and integration of fluorescence spectra (between 610 and 850 nm) are represented as averages of three independently repeated experiments (N=3) with standard deviations.

A)

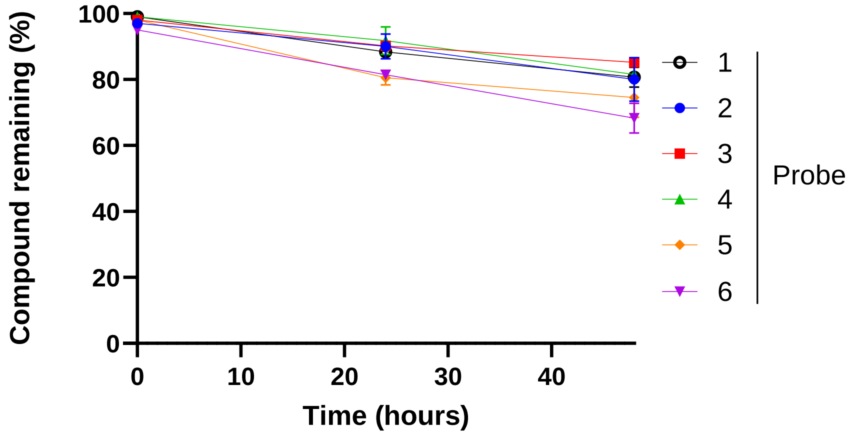

B)

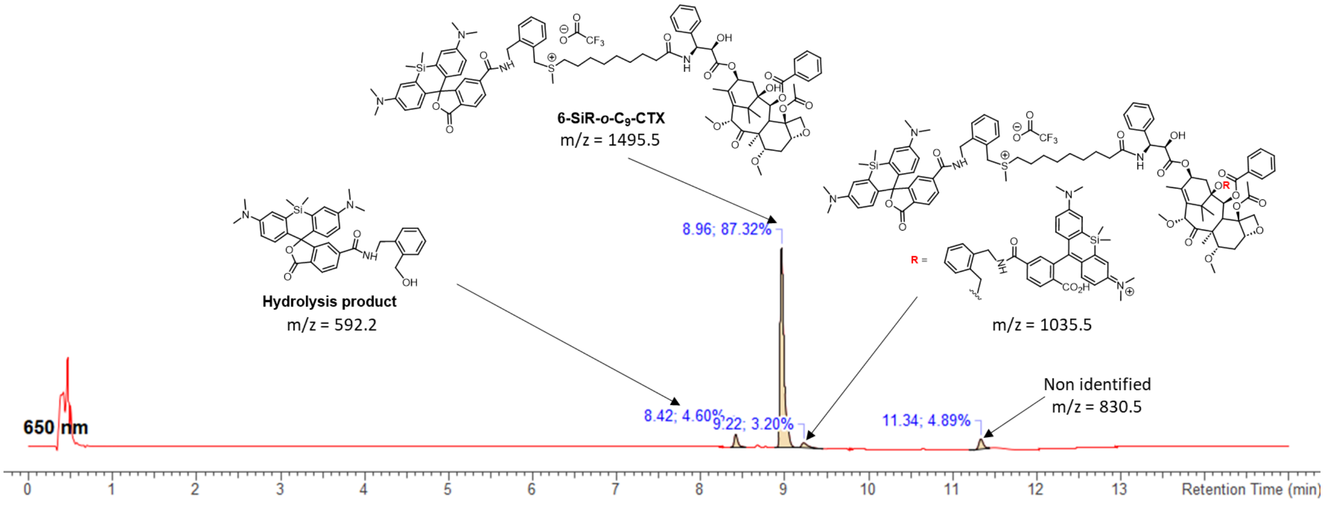

**Figure S4. Stability of covalent tubulin probes in PBS pH=7.4 at 37°C.** Solution of covalent probes (20 µM in PBS) were heated at 37°C in the dark. Solutions were analyzed after 0, 24h and 48h by LC/MS. A) The percentage of compound remaining over time according is plotted. B) Examples of degradation products observed after 48h.

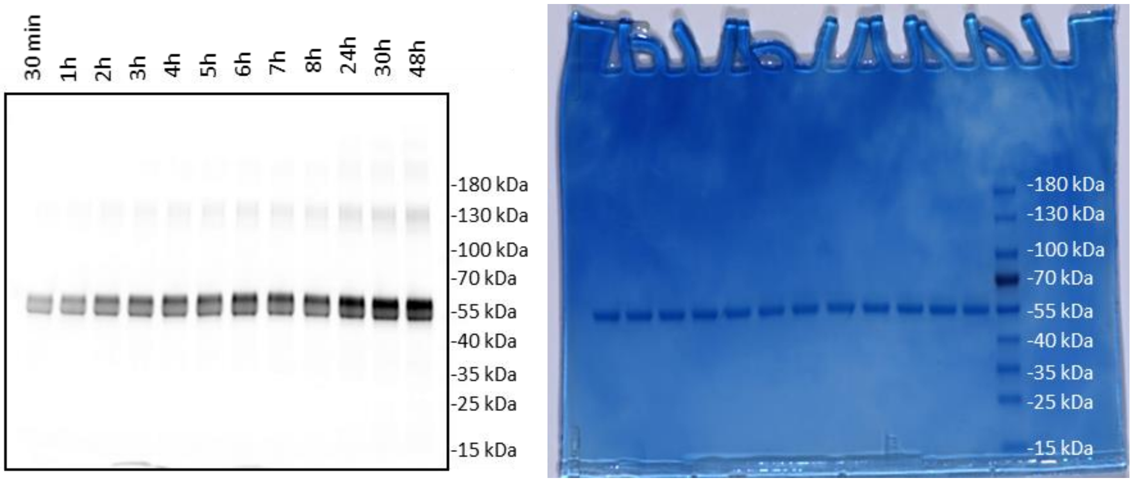

**Figure S5. Full representative *in-gel* fluorescence and Coomassie staining for kinetic (Figure 4A&B).**

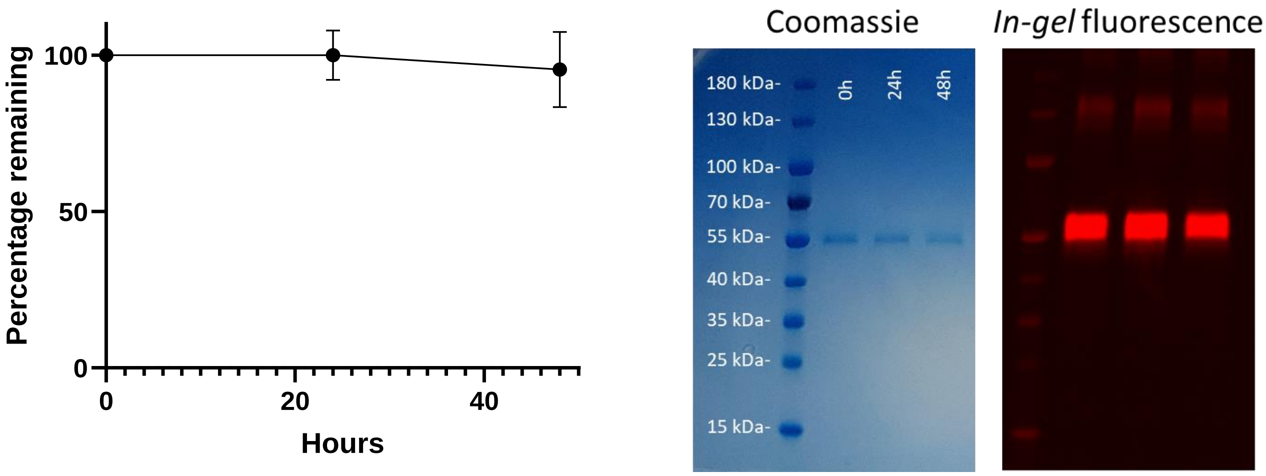

**Figure S6. Stability of the covalent bond between the dye and tubulin in PBS pH=7.4 at 37°C.** The experiment was performed in triplicate (N=3) to obtain mean and standard deviation values (shown as error bars).

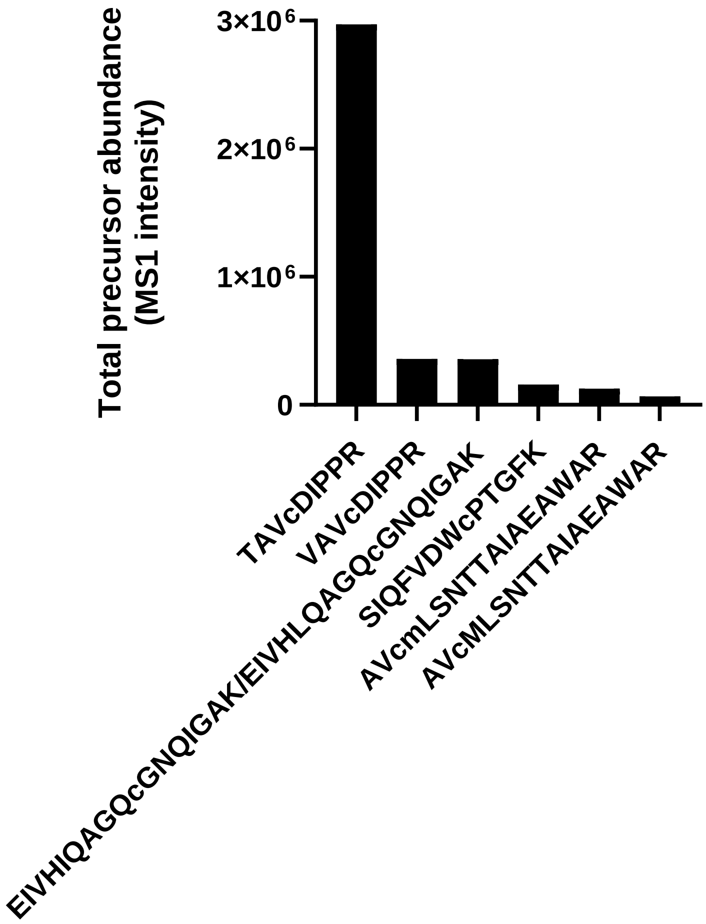

**Figure S7. Identified labeled peptides after *in-gel* analysis of SiR-labeled tubulin.** c is the modified cysteine. m is an oxidized methionine.

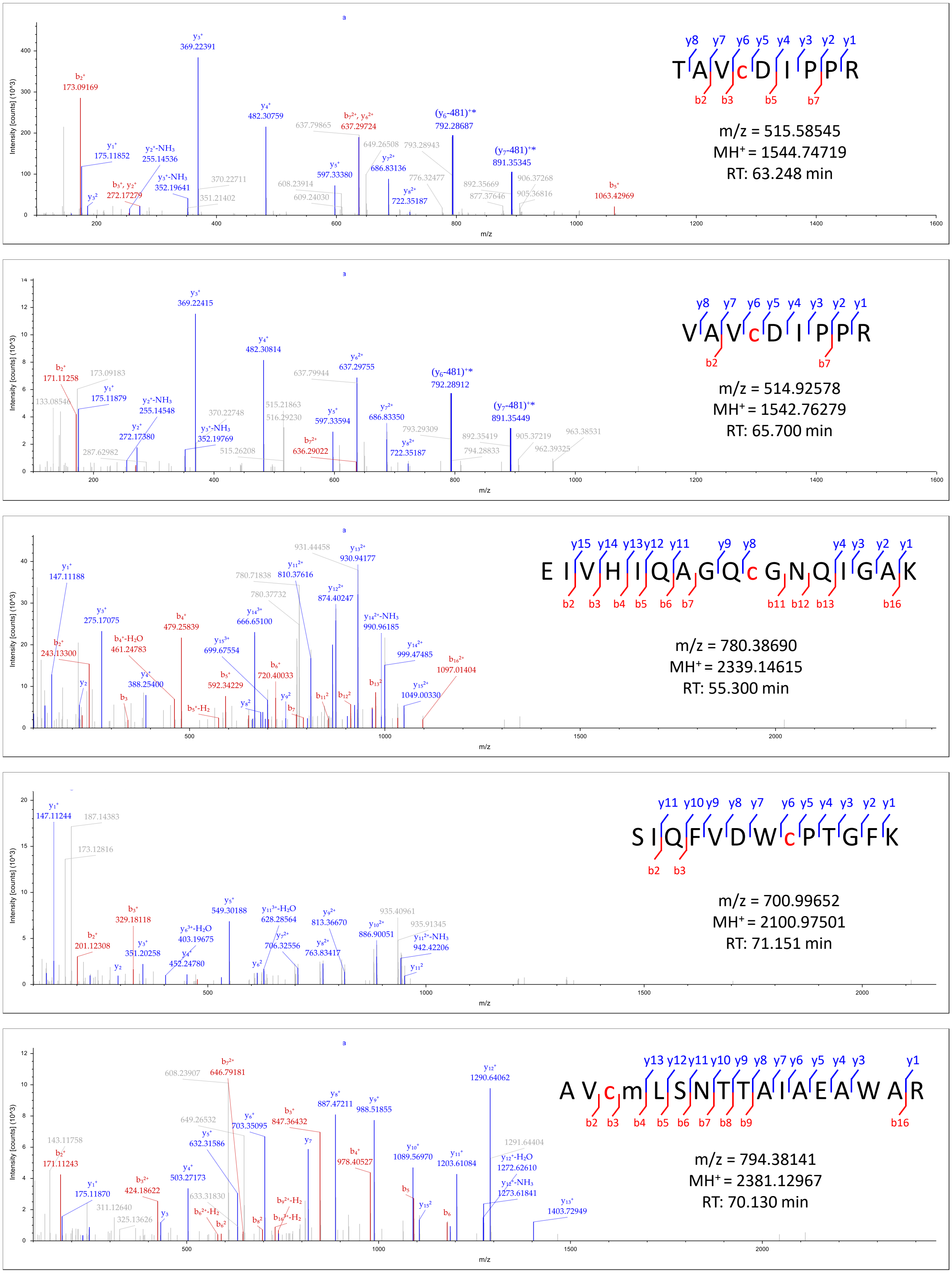

**Figure S8. MS spectra of identified labeled peptides.** c is the modified cysteine. m is an oxidized methionine. Ions marked with a star (*) correspond to fragment ions with a mass loss of 481 Da, which may be caused by fragmentation of the label.

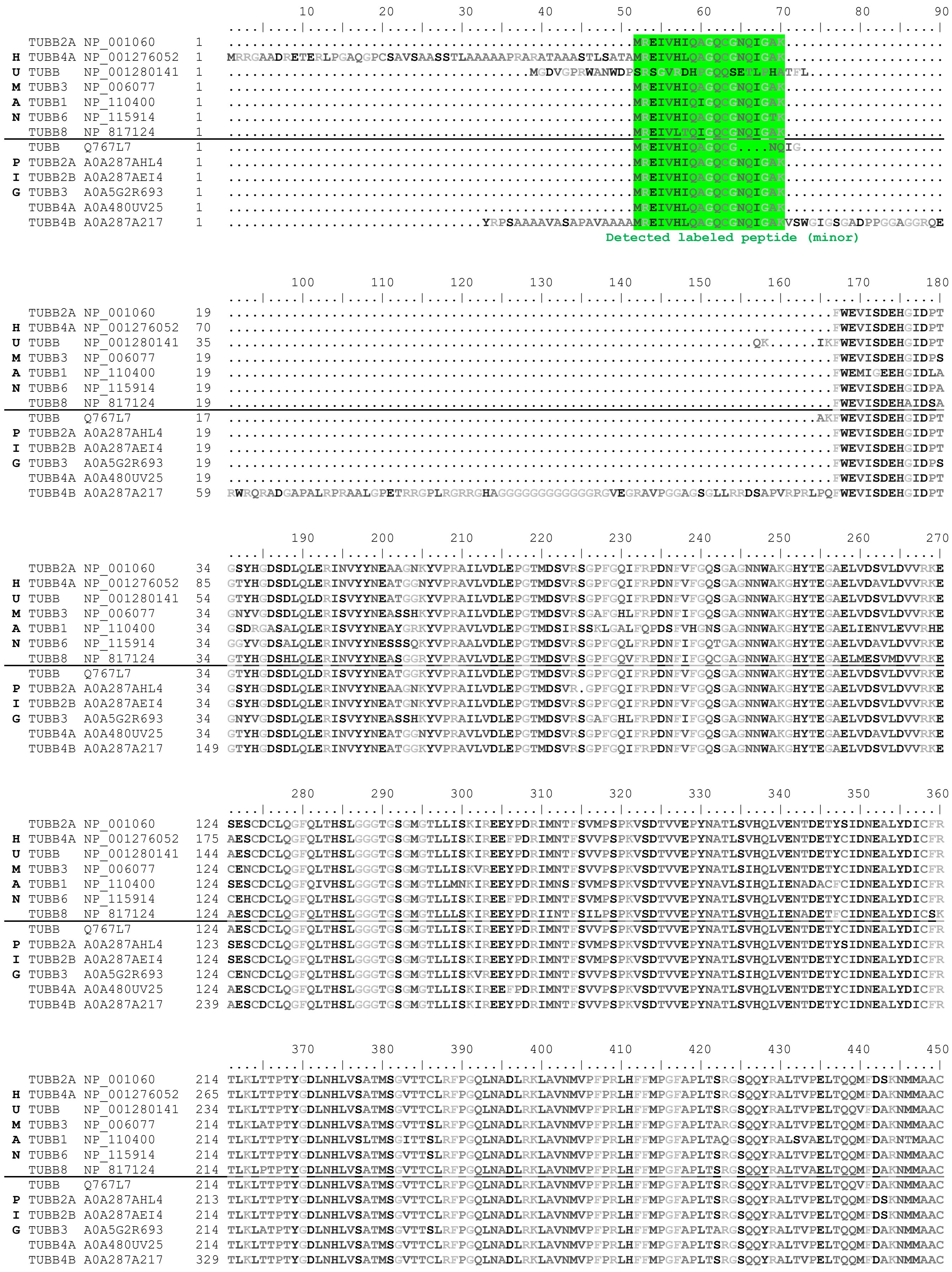
10 20 30 40 50 60 70 80 90
H U M A N
TUBB2A NP_001060
TUBB4A NP_001276052
TUBB NP_001280141
TUBB3 NP_006077
TUBB1 NP_110400
TUBB6 NP_115914
TUBB8 NP_817124
P I G
TUBB Q767L7
TUBB2A A0A287AHL4
TUBB2B A0A287AEI4
TUBB3 A0A5G2R693
TUBB4A A0A480UV25
TUBB4B A0A287A217
Detected labeled peptide (minor)
100 110 120 130 140 150 160 170 180
190 200 210 220 230 240 250 260 270
280 290 300 310 320 330 340 350 360
370 380 390 400 410 420 430 440 450

```
                                             460       470       480       490       500       510       520       530       540
                                    ....|....|....|....|....|....|....|....|....|....|....|....|....|....|....|....|....|....|
  TUBB2A NP_001060             304 DPRHGRYLTVAAIFRGRMSMKEVDEQMLNVQNKNSSYFVEWIPNNVKTAVCDIPPRGLKMSATFIGNSTAIQELFKRISEQFTAMFRRKA
H TUBB4A NP_001276052          355 DPRHGRYLTVAAVFRGRMSMKEVDEQMLSVQSKNSSYFVEWIPNNVKTAVCDIPPRGLKMAATFIGNSTAIQELFKRISEQFTAMFRRKA
U TUBB   NP_001280141          324 DPRHGRYLTVAAVFRGRMSMKEVDEQMLNVQNKNSSYFVEWIPNNVKTAVCDIPPRGLKMAVTFIGNSTAIQELFKRISEQFTAMFRRKA
M TUBB3  NP_006077             304 DPRHGRYLTVATVFRGRMSMKEVDEQMLAIQSKNSSYFVEWIPNNVKVAVCDIPPRGLKMSSTFIGNSTAIQELFKRISEQFTAMFRRKA
A TUBB1  NP_110400             304 DLRRGRYLTVACIFRGKMSTKEVDQQLLSVQTRNSSCFVEWIPNNVKVAVCDIPPRGLSMAATFIGNNTAIQEIFNRVSEHFSAMFKRKA
N TUBB6  NP_115914             304 DPRHGRYLTVATVFRGPMSMKEVDEQMLAIQSKNSSYFVEWIPNNVKVAVCDIPPRGLKMASTFIGNSTAIQELFKRISEQFSAMFRRKA
  TUBB8  NP_817124             304 DPRHGRYLTAAAIFRGRMPMREVDEQMFNIQDKNSSYFADWLPNNVKTAVCDIPPRGLKMSATFIGNNTAIQELFKRVSEQFTAMFRRKA
---------------------------------------------------------------------------------------------------------------------------
  TUBB   Q767L7                304 DPRHGRYLTVAAVFRGRMSMKEVDEQMLNVQNKNSSYFVEWIPNNVKTAVCDIPPRGLKMAVTFIGNSTAIQELFKRISEQFTAMFRRKA
P TUBB2A A0A287AHL4            303 DPRHGRYLTVAAIFRGRMSMKEVDEQMLNVQNKNSSYFVEWIPNNVKTAVCDIPPRGLKMSATFIGNSTAIQELFKRISEQFTAMFRRKA
I TUBB2B A0A287AEI4            304 DPRHGRYLTVAAIFRGRMSMKEVDEQMLNVQNKNSSYFVEWIPNNVKTAVCDIPPRGLKMSATFIGNSTAIQELFKRISEQFTAMFRRKA
G TUBB3  A0A5G2R693            304 DPRHGRYLTVATVFRGRMSMKEVDEQMLAIQSKNSSYFVEWIPNNVKVAVCDIPPRGLPALQHERPGVRVPAVPGRHGRRGGRDVRGRRG
  TUBB4A A0A480UV25            304 DPRHGRYLTVAAVFRGRMSMKEVDEQMLSVQSKNSSYFVEWIPNNVKTAVCDIPPRGLKMAATFIGNSTAIQELFKRISEQFTAMFRRKA
  TUBB4B A0A287A217            419 DPRHGRYLTVAAVFRGRMSMKEVDEQMLNVQNKNSSYFVEWIPNNVKTAVCDIPPRGLKMSATFIGNSTAIQELFKRISEQFTAMFRRKA
                                                                   Detected labeled peptide (major)

                                             550       560       570       580       590       600       610       620       630
                                    ....|....|....|....|....|....|....|....|....|....|....|....|....|....|....|....|....|....|
  TUBB2A NP_001060             394 FLHWYT........GEGMDEMEFTEAESNMNDLVSEYQQYQDATADEQGEFEEEEGEDEA..............................
H TUBB4A NP_001276052          445 FLHWYT........GEGMDEMEFTEAESNMNDLVSEYQQYQDATAEE.GEFEEEAEEEVA..............................
U TUBB   NP_001280141          414 FLHWYT........GEGMDEMEFTEAESNMNDLVSEYQQYQDATAEEEEDFGEEAEEEA...............................
M TUBB3  NP_006077             394 FLHWYT........GEGMDEMEFTEAESNMNDLVSEYQQYQDATAEEEGEMYEDDEEESEAQGPK.........................
A TUBB1  NP_110400             394 FVHWYT........SEGMDINEFGEAENNIHDLVSEYQQFQDAKAVLEEDEEVTEEAEMEPEDKGH........................
N TUBB6  NP_115914             394 FLHWFT........GEGMDEMEFTEAESNMNDLVSEYQQYQDATANDGEEAFEDEEEEIDG.............................
  TUBB8  NP_817124             394 FLHWYT........GEGMDEMEFTEAESNMNDLVSEYQQYQDATAEEEEDEEYAEEEVA...............................
---------------------------------------------------------------------------------------------------------------------------
  TUBB   Q767L7                394 FLHWYT........GEGMDEMEFTEAESNMNDLVSEYQQYQDATAEEEEDFGEEAEEEA...............................
P TUBB2A A0A287AHL4            393 FLHWYT........GEGMDEMEFTEAESNMNDLVSEYQQYQDATADEQGEFEEEEGEDEA..............................
I TUBB2B A0A287AEI4            394 FLHWYT........GEGMDEMEFTEAESNMNDLVSEEWKASSCVPSSYIFLPP..GQPF...............................
G TUBB3  A0A5G2R693            394 GVRGPGPQVRRPGGGRGTARTQRSAPSRVAPDTARPRPTRPERVAPGPPSVRPAVFAASLPRPSPPSHPPPSVISVVCGPLCLLYCPLQA
  TUBB4A A0A480UV25            394 FLHWYT........GEGMDEMEFTEAESNMNDLVSEYQQYQDATAEE.GEFEEEAEEEVA..............................
  TUBB4B A0A287A217            509 FLHWYT........GEGMDEMEFTEAESNMNDLVSEYQQYQDATAEEEGEFEEEAEEEVA..............................
```

**Figure S9. Sequence alignment of β-tubulin isotypes from human (Homo sapiens) and pig (Sus scrofa).** Labeled peptides detected in the tubulin sample are marked green.

```
                                   10        20        30        40        50        60        70        80        90
                          ....|....|....|....|....|....|....|....|....|....|....|....|....|....|....|....|....|....|
H TUBA2  ABD72607       1 ......................................................................MRECISIHVGQAGVQIGNAC
U TUBA1A NP_001257328   1 ......................................................................MRECISIHVGQAGVQIGNAC
M TUBA1C NP_001290043   1 MTAWFNGVEELGRDECFVCLELDPSNGSFLNAKLYISNWILIYFTRTLQLPAPFNTCTSSKSRGEPGCDSKRECISIHVGQAGVQIGNAC
A TUBA4A NP_005991      1 ......................................................................MRECISVHVGQAGVQMGNAC
N TUBA1B NP_006073      1 ......................................................................MRECISIHVGQAGVQIGNAC
  TUBA8  NP_061816      1 ......................................................................MRECISVHVGQAGVQIGNAC
  TUBA1A P02550         1 ......................................................................MRECISIHVGQAGVQIGNAC
P TUBA3C F1RK98         1 ...........................................................LFWASHICSGPQRECISIHVGQAGVQIGNAC
I TUBA4A A0A286ZNY1     1 ......................................................................MRECISVHVGQAGVQMGNAC
G TUBA8A A0A287BMB7     1 ......................................................................MRECISVHVGQAGVQIGNAC

                                  100       110       120       130       140       150       160       170       180
                          ....|....|....|....|....|....|....|....|....|....|....|....|....|....|....|....|....|....|
H TUBA2  ABD72607      21 WELYCLEHGIQPDGQMPSDKTIGGGDDSFNTFFSETGAGKHVPRAVFVDLEPTVVDEVRTGTYRQLFHPEQLITGKEDAANNYARGHYTI
U TUBA1A NP_001257328  21 WELYCLEHGIQPDGQMPSDKTIGGGDDSFNTFFSETGAGKHVPRAVFVDLEPTVIDEVRTGTYRQLFHPEQLITGKEDAANNYARGHYTI
M TUBA1C NP_001290043  91 WELYCLEHGIQPDGQMPSDKTIGGGDDSFNTFFSETGAGKHVPRAVFVDLEPTVIDEVRTGTYRQLFHPEQLITGKEDAANNYARGHYTI
A TUBA4A NP_005991     21 WELYCLEHGIQPDGQMPSDKTIGGGDDSFTTFFCETGAGKHVPRAVFVDLEPTVIDEIRNGPYRQLFHPEQLITGKEDAANNYARGHYTI
N TUBA1B NP_006073     21 WELYCLEHGIQPDGQMPSDKTIGGGDDSFNTFFSETGAGKHVPRAVFVDLEPTVIDEVRTGTYRQLFHPEQLITGKEDAANNYARGHYTI
  TUBA8  NP_061816     21 WELFCLEHGIQADGTFDAQASKINDDDSFTTFFSETGNGKHVPRAVMIDLEPTVVDEVRAGTYRQLFHPEQLITGKEDAANNYARGHYTV
  TUBA1A P02550        21 WELYCLEHGIQPDGQMPSDKTIGGGDDSFNTFFSETGAGKHVPRAVFVDLEPTVIDEVRTGTYRQLFHPEQLITGKEDAANNYARGHYTI
P TUBA3C F1RK98        32 WELYCLEHGIQPDGQMPSDKTIGGGDDSFNTFFSETGAGKHVPRAVFVDLEPTVVDEVRTGTYRQLFHPEQLITGKEDAANNYARGHYTI
I TUBA4A A0A286ZNY1    21 WELYCLEHGIQPDGQMPSDKTIGGGDDSFTTFFCETGAGKHVPRAVFVDLEPTVIDEIRNGPYRQLFHPEQLITGKEDAANNYARGHYTI
G TUBA8A A0A287BMB7    21 WELFCLEHGIQADGTFGAQASKLHDDDSFTTFFSETGNGKHVPRAVMVDLEPTVVDEVRAGTYRQLFHPEQLITGKEDAANNYARGHYTV

                                  190       200       210       220       230       240       250       260       270
                          ....|....|....|....|....|....|....|....|....|....|....|....|....|....|....|....|....|....|
H TUBA2  ABD72607     111 GKEIVDLVLDRIRKLADLCTGLQGFLIFHSFGGGTGSGFASLLMERLSVDYGKKSKLEFAIYPAPQVSTAVVEPYNSILTTHTTLEHSDC
U TUBA1A NP_001257328 111 GKEIIDLVLDRIRKLADQCTGLQGFLVFHSFGGGTGSGFTSLLMERLSVDYGKKSKLEFSIYPAPQVSTAVVEPYNSILTTHTTLEHSDC
M TUBA1C NP_001290043 181 GKEIIDLVLDRIRKLADQCTGLQGFLVFHSFGGGTGSGFTSLLMERLSVDYGKKSKLEFSIYPAPQVSTAVVEPYNSILTTHTTLEHSDC
A TUBA4A NP_005991    111 GKEIIDPVLDRIRKLSDQCTGLQGFLVFHSFGGGTGSGFTSLLMERLSVDYGKKSKLEFSIYPAPQVSTAVVEPYNSILTTHTTLEHSDC
N TUBA1B NP_006073    111 GKEIIDLVLDRIRKLADQCTGLQGFLVFHSFGGGTGSGFTSLLMERLSVDYGKKSKLEFSIYPAPQVSTAVVEPYNSILTTHTTLEHSDC
  TUBA8  NP_061816    111 GKESIDLVLDRIRKLTDACSGLQGFLIFHSFGGGTGSGFTSLLMERLSLDYGKKSKLEFAIYPAPQVSTAVVEPYNSILTTHTTLEHSDC
  TUBA1A P02550       111 GKEIIDLVLDRIRKLADQCTGLQGFSVFHSFGGGTGSGFTSLLMERLSVDYGKKSKLEFSIYPAPQVSTAVVEPYNSILTTHTTLEHSDC
P TUBA3C F1RK98       122 GKEIVDLVLDRIRKLADLCTGLQGFLIFHSFGGGTGSGFASLLMERLSVDYGKKSKLEFAIYPAPQVSTAVVEPYNSILTTHTTLEHSDC
I TUBA4A A0A286ZNY1   111 GKEIIDPVLDRIRKLSDQCTGLQGFLVFHSFGGGTGSGFTSLLMERLSVDYGKKSKLEFSIYPAPQVSTAVVEPYNSILTTHTTLEHSDC
G TUBA8A A0A287BMB7   111 GKESIDLVLDRIRKLTDACSGLQGFLIFHSFGGGTGSGFTSLLMERLSLDYGKKSKLEFAIYPAPQVSTAVVEPYNSILTTHTTLEHSDC

                                  280       290       300       310       320       330       340       350       360
                          ....|....|....|....|....|....|....|....|....|....|....|....|....|....|....|....|....|....|
H TUBA2  ABD72607     201 AFMVDNEAIYDICRRNLDIERPTYTNLNRLIGQIVSSITASLRFDGALNVDLTEFQTNLVPYPRIHFPLATYAPVISAEKAYHEQLSVAE
U TUBA1A NP_001257328 201 AFMVDNEAIYDICRRNLDIERPTYTNLNRLIGQIVSSITASLRFDGALNVDLTEFQTNLVPYPRIHFPLATYAPVISAEKAYHEQLSVAE
M TUBA1C NP_001290043 271 AFMVDNEAIYDICRRNLDIERPTYTNLNRLISQIVSSITASLRFDGALNVDLTEFQTNLVPYPRIHFPLATYAPVISAEKAYHEQLTVAE
A TUBA4A NP_005991    201 AFMVDNEAIYDICRRNLDIERPTYTNLNRLISQIVSSITASLRFDGALNVDLTEFQTNLVPYPRIHFPLATYAPVISAEKAYHEQLSVAE
N TUBA1B NP_006073    201 AFMVDNEAIYDICRRNLDIERPTYTNLNRLISQIVSSITASLRFDGALNVDLTEFQTNLVPYPRIHFPLATYAPVISAEKAYHEQLSVAE
  TUBA8  NP_061816    201 AFMVDNEAIYDICRRNLDIERPTYTNLNRLISQIVSSITASLRFDGALNVDLTEFQTNLVPYPRIHFPLVTYAPIISAEKAYHEQLSVAE
  TUBA1A P02550       201 AFMVDNEAIYDICRRNLDIERPTYTNLNRLIGQIVSSITASLRFDGALNVDLTEFQTNLVPYPRAHFPLATYAPVISAEKAYHEQLSVAE
P TUBA3C F1RK98       212 AFMVDNEAIYDICRRNLDIERPTYTNLNRLIGQIVSSITASLRFDGALNVDLTEFQTNLVPYPRIHFPLATYAPVISAEKAYHEQLSVAE
I TUBA4A A0A286ZNY1   201 AFMVDNEAIYDICRRNLDIERPTYTNLNRLISQIVSSITASLRFDGALNVDLTEFQTNLVPYPRIHFPLATYAPVISAEKAYHEQLSVAE
G TUBA8A A0A287BMB7   201 AFMVDNEAIYDICRRNLDIERPTYTNLNRLISQIVSSITASLRFDGALNVDLTEFQTNLVPYPRIHFPLVTYAPIISAEKAYHEQLSVAE

                                  370       380       390       400       410       420       430       440       450
                          ....|....|....|....|....|....|....|....|....|....|....|....|....|....|....|....|....|....|
H TUBA2  ABD72607     291 ITNACFEPANQMVKCDPRHGKYMACCMLYRGDVVPKDVNAAIATIKTKRTIQFVDWCPTGFKVGINYQPPTVVPGGDLAKVQRAVCMLSN
U TUBA1A NP_001257328 291 ITNACFEPANQMVKCDPRHGKYMACCLLYRGDVVPKDVNAAIATIKTKRTIQFVDWCPTGFKVGINYQPPTVVPGGDLAKVQRAVCMLSN
M TUBA1C NP_001290043 361 ITNACFEPANQMVKCDPRHGKYMACCLLYRGDVVPKDVNAAIATIKTKRTIQFVDWCPTGFKVGINYQPPTVVPGGDLAKVQRAVCMLSN
A TUBA4A NP_005991    291 ITNACFEPANQMVKCDPRHGKYMACCLLYRGDVVPKDVNAAIAAIKTKRSIQFVDWCPTGFKVGINYQPPTVVPGGDLAKVQRAVCMLSN
N TUBA1B NP_006073    291 ITNACFEPANQMVKCDPRHGKYMACCLLYRGDVVPKDVNAAIATIKTKRSIQFVDWCPTGFKVGINYQPPTVVPGGDLAKVQRAVCMLSN
  TUBA8  NP_061816    291 ITSSCFEPNSQMVKCDPRHGKYMACCMLYRGDVVPKDVNVAIAAIKTKRTIQFVDWCPTGFKVGINYQPPTVVPGGDLAKVQRAVCMLSN
  TUBA1A P02550       291 ITNACFEPANQMVKCDPRHGKYMACCLLYRGDVVPKDVNAAIATIKTKRTIQFVDWCPTGFKVGINYEPPTVVPGGDLAKVQRAVCMLSN
P TUBA3C F1RK98       302 ITNACFEPANQMVKCDPRHGKYMACCMLYRGDVVPKDVNAAIATIKTKRTIQFVDWCPTGFKVGINYQPPTVVPGGDLAKVQRAVCMLSN
I TUBA4A A0A286ZNY1   291 ITNACFEPANQMVKCDPRHGKYMACCLLYRGDVVPKDVNAAIAAIKTKRSIQFVDWCPTGFKVGINYQPPTVVPGGDLAKVQRAVCMLSN
G TUBA8A A0A287BMB7   291 ITSSCFEPNSQMVKCDPRHGKYMACCMLYRGDVVPKDVNVAIAAIKTKRTIQFVDWCPTGFKVGINYQPPTVVPGGDLAKVQRAVCMLSN
                                                                                Detected labeled peptides (minor)

                                  460       470       480       490       500       510       520
                          ....|....|....|....|....|....|....|....|....|....|....|....|....|....|.
H TUBA2  ABD72607     381 TTAIAEAWARLDHKFDLMYAKRAFVHWYVGEGMEEGEFSEAREDLAALEKDYEEVGVDSVEAEAE.EGEEY
U TUBA1A NP_001257328 381 TTAIAEAWARLDHKFDLMYAKRAFVHWYVGEGMEEGEFSEAREDMAALEKDYEEVGVDSVEGEGEEEGEEY
M TUBA1C NP_001290043 451 TTAVAEAWARLDHKFDLMYAKRAFVHWYVGEGMEEGEFSEAREDMAALEKDYEEVGADSADGEDE  GEEY
A TUBA4A NP_005991    381 TTAIAEAWARLDHKFDLMYAKRAFVHWYVGEGMEEGEFSEAREDMAALEKDYEEVGIDSYEDEDE  GEE
N TUBA1B NP_006073    381 TTAIAEAWARLDHKFDLMYAKRAFVHWYVGEGMEEGEFSEAREDMAALEKDYEEVGVDSVEGEGEEEGEEY
  TUBA8  NP_061816    381 TTAIAEAWARLDHKFDLMYAKRAFVHWYVGEGMEEGEFSEAREDLAALEKDYEEVGTDSFEEENE..GEEF
  TUBA1A P02550       381 TTAIAEAWARLDHKFDLMYAKRAFVHWYVGEGMEEGEFSEAREDMAALEKDYEEVGVDSVEGEGEEEGEEY
P TUBA3C F1RK98       392 TTAIAEAWARLDHKFDLMYAKRAFVHWYVGEGMEEGEFSEAREDLAALEKDYEEVGVDSVEAEAE.EGEEY
I TUBA4A A0A286ZNY1   381 TTAIAEAWARLDHKFDLMYAKRAFVHWYVGEGMEEGEFSEAREDMAALEKDYEEVGIDSYEDEDE  GEE.
G TUBA8A A0A287BMB7   381 TTAIAEAWARLDHKFDLMYAKRAFVHWYVGEGMEEGEFSEAREDLAALEKDYEEVGTDSFEEENE  GEEF
```

**Figure S10. Sequence alignment of α-tubulin isotypes from human (Homo sapiens) and pig (Sus scrofa).** Labeled peptides detected in the tubulin sample are marked green.

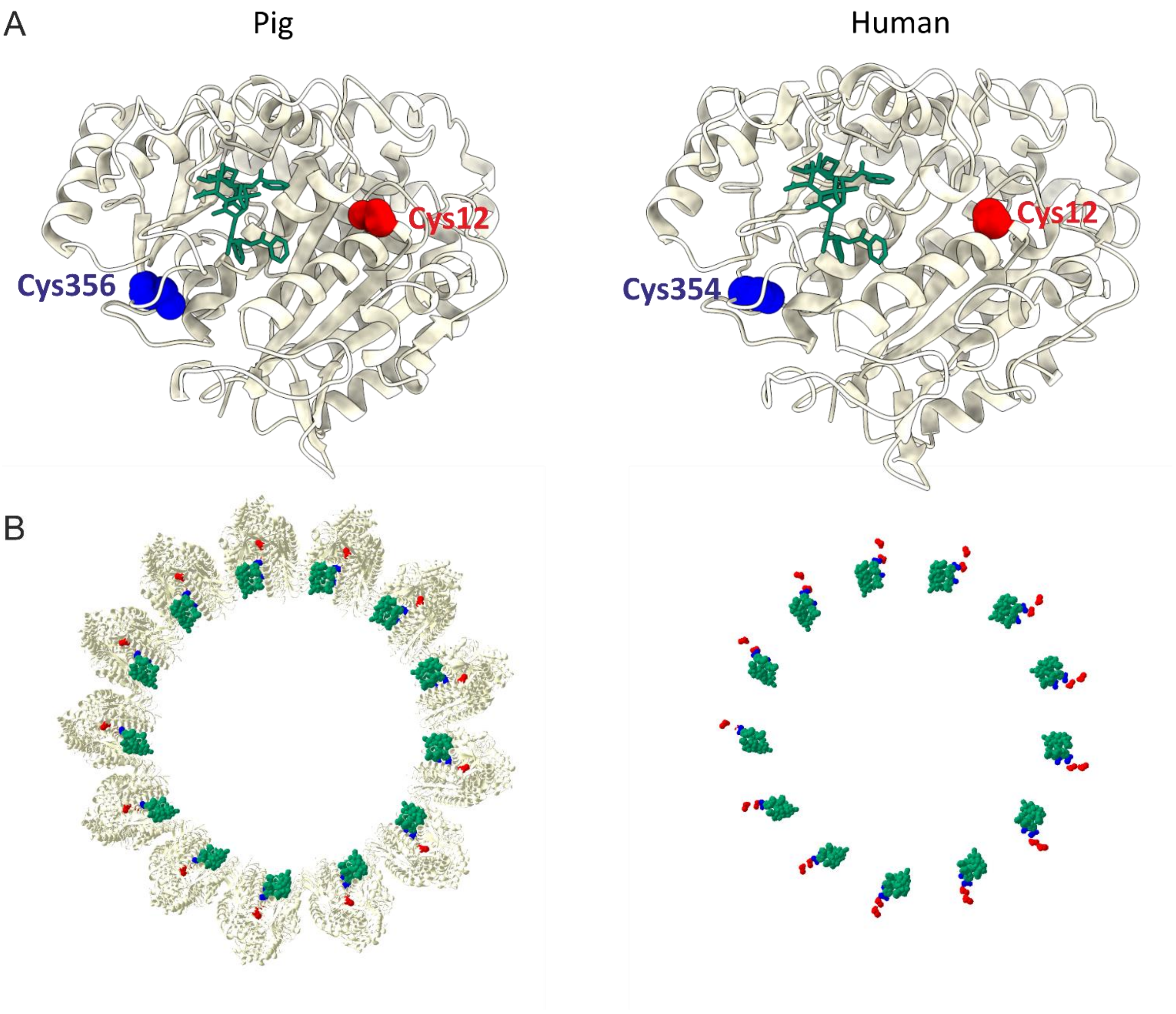

**Figure S11. Structural models of tubulin taxane binding sites and labeled Cys residues.** (A) Comparison of cryo-EM β-tubulin complex with taxol from pig (PDB: 5SYF) and model of human β-tubulin complex with taxol (model is based on cryo-EM structure PDB: 6E7C). (B) Microtubule-taxol complex model (PDB: 5SYF). Left panel shows top view of microtubule composed of tubulin dimers, taxol and the main labeling site (Cys356). Right panel shows same view as on left, but protein molecules are omitted. Protein molecule is shown in white, Taxol is green, the main labeling site (Cys356 or Cys354) is highlighted in blue, and the minor labeling site (Cys12) in red. GTP, GDP and $Mg^{2+}$ molecules are omitted for clarity.

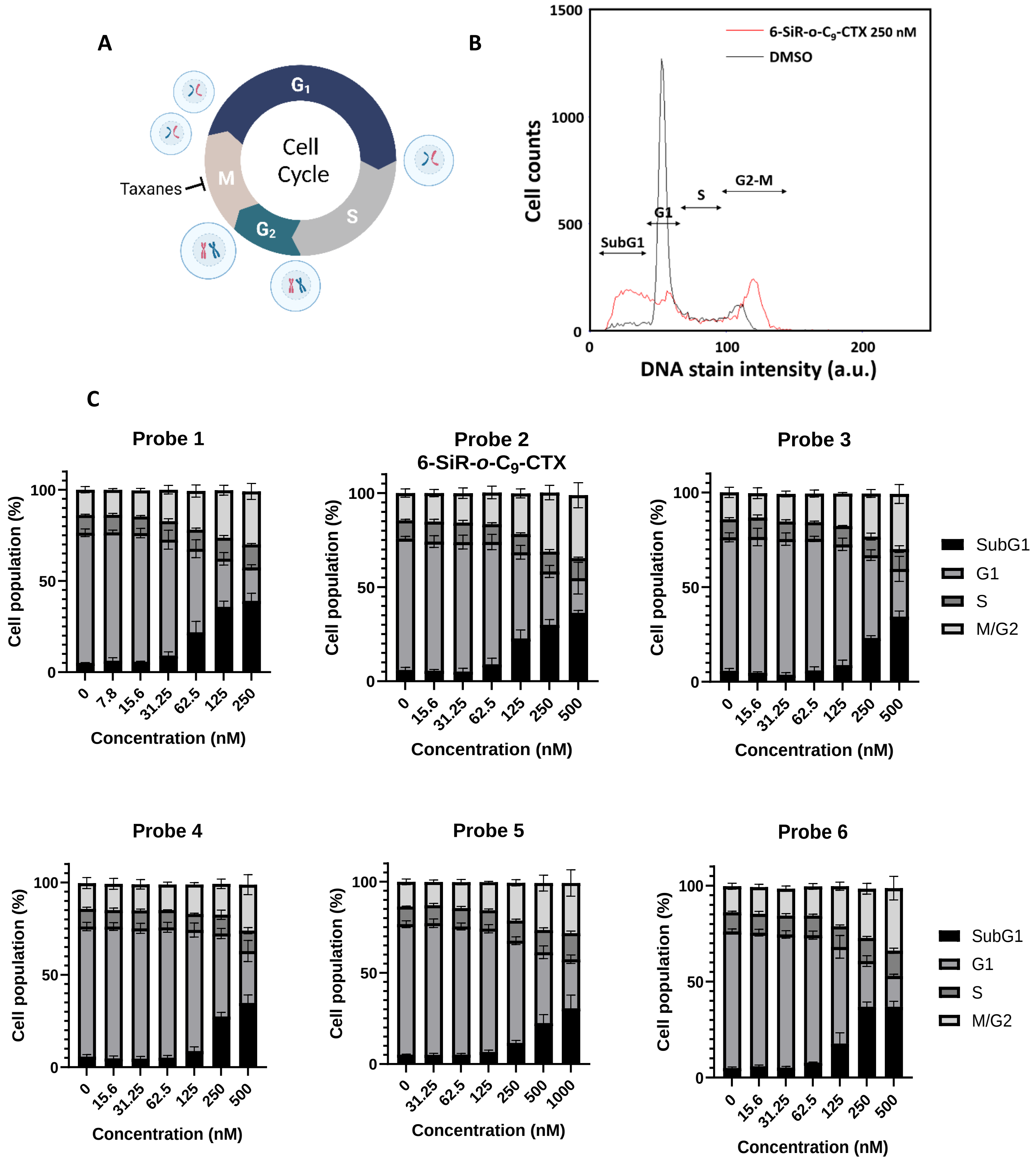

**Figure S12. Cell cycle perturbation induced by covalent tubulin probes.** (A) Cytotoxicity of taxanes results from the inhibition of the cycle at the stage of mitosis. Created with BioRender.com (B). Representative histogram of DNA content distribution in HeLa cells treated with DMSO or 250 nM of **6-SiR-*o*-$C_9$-CTX** for 24h. The cell cycle phases are identified by the amount of DNA per cell. (C) Cytotoxicity measurements of tubulin probes. Experimental data are averages of three independent experiments (N=3) and presented as means with standard deviations.

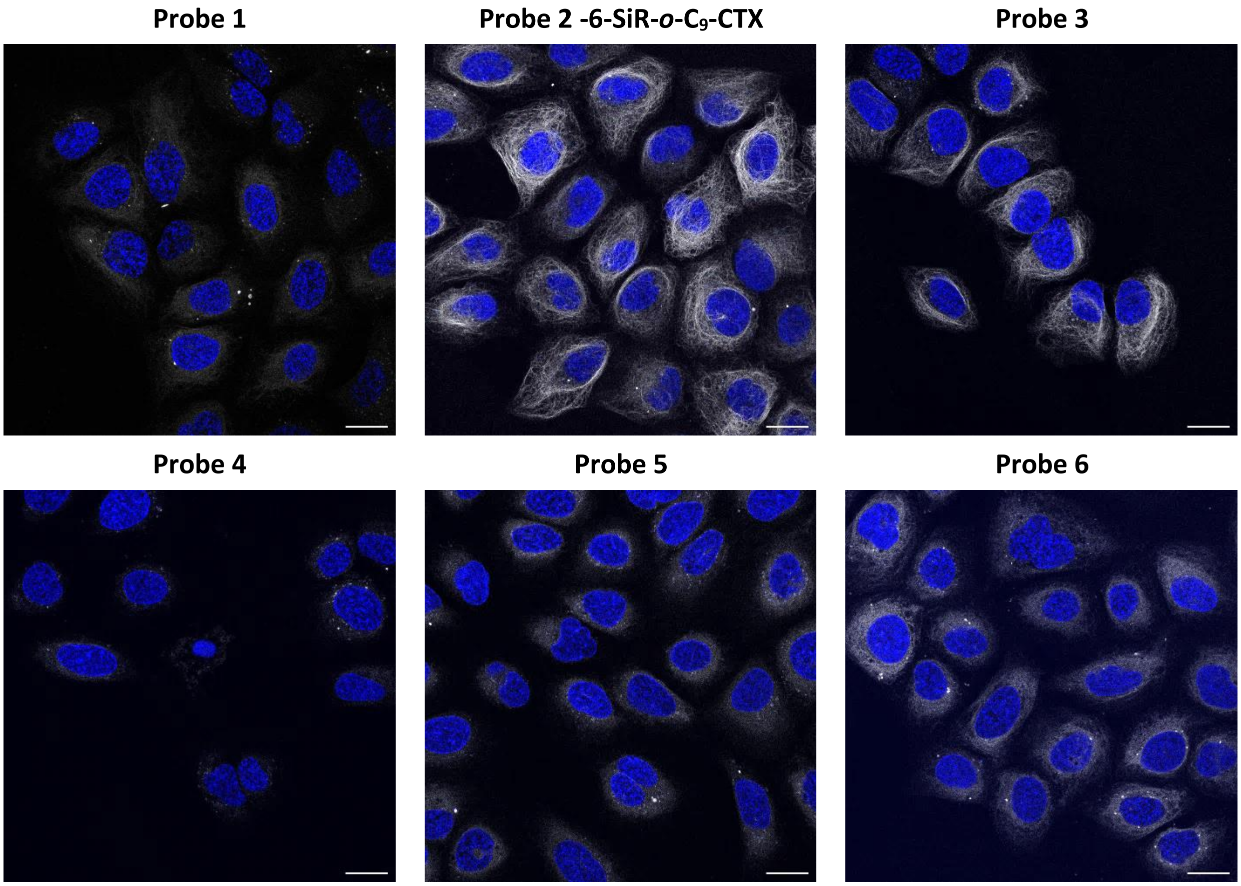

**Figure S13. Live U-2 OS cells stained with probes (1 µM in OptiMEM), verapamil (10 µM) and Hoechst 33342 (1 µg/mL) for 4h.** Confocal microscopy images were acquired using LEICA SP8. Gray channel ($\lambda_{ex}$= 633 nm, $\lambda_{em}$= 650-710 nm) corresponds to probe staining and blue channel ($\lambda_{ex}$= 405 nm, $\lambda_{em}$= 415-480 nm) corresponds to Hoechst 33342 staining. Scale bar = 20 µm.

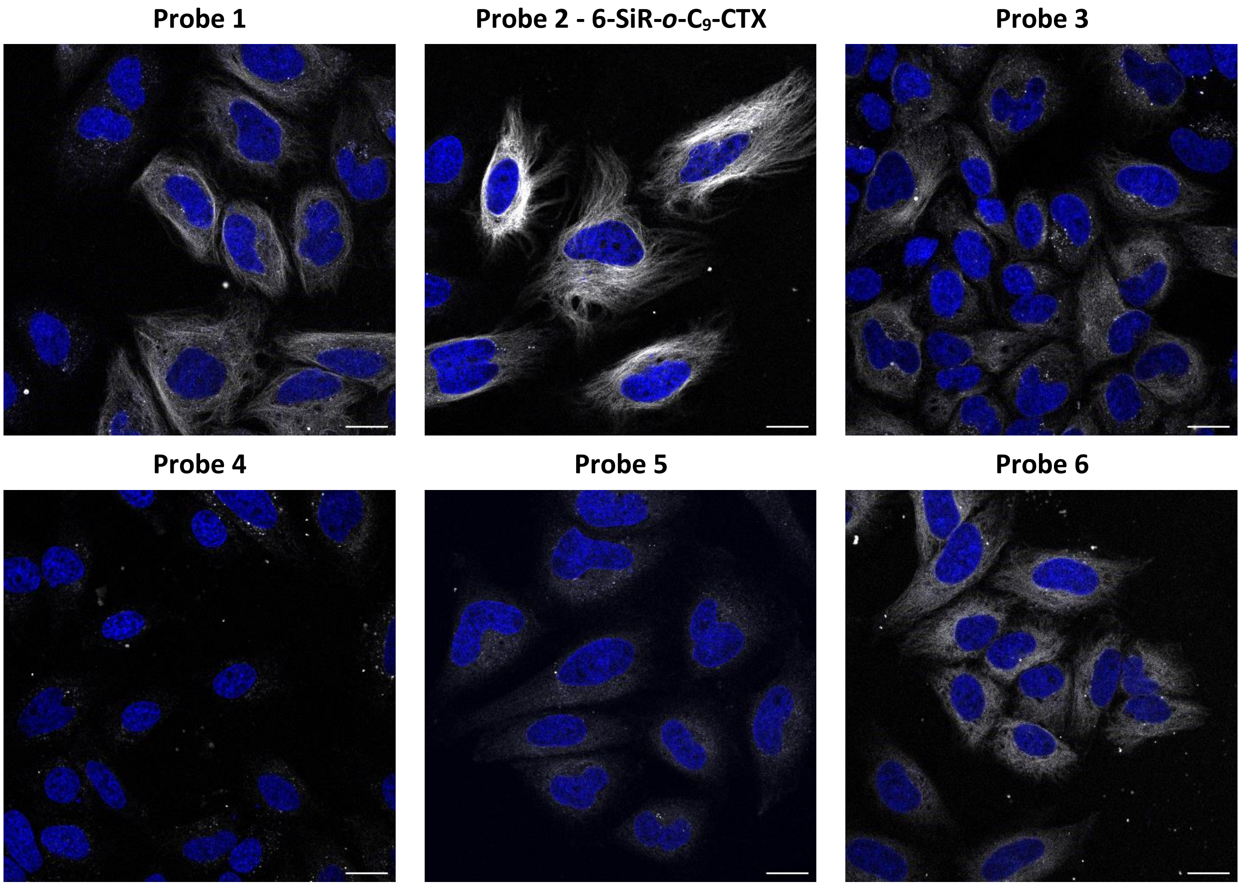

**Figure S14. Live HeLa CCL cells stained with probes (1 µM in OptiMEM) and Hoechst 33342 (1 µg/mL) for 4h.** Confocal microscopy images were acquired using LEICA SP8. Gray channel ($\lambda_{ex}$= 633 nm, $\lambda_{em}$= 650-710 nm) corresponds to probe staining and blue channel ($\lambda_{ex}$= 405 nm, $\lambda_{em}$= 415-480 nm) corresponds to Hoechst 33342 staining. Scale bar = 20 µm.

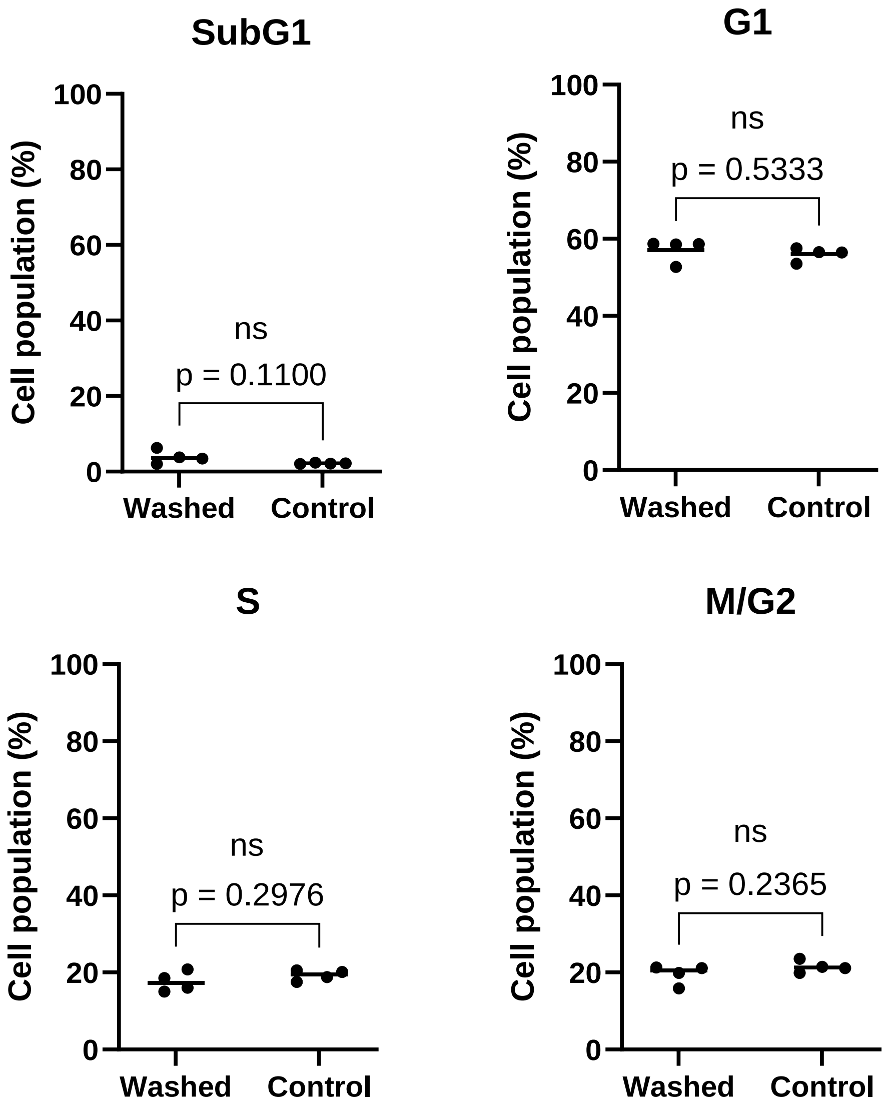

**Figure S15. Statistical analysis associated with the cell cycle analysis of washing experiment.** Data from four different experiments (N=4). The line corresponds to the mean and each dot corresponds to the result of one single experiment. Unpaired t test was performed between the 2 conditions.

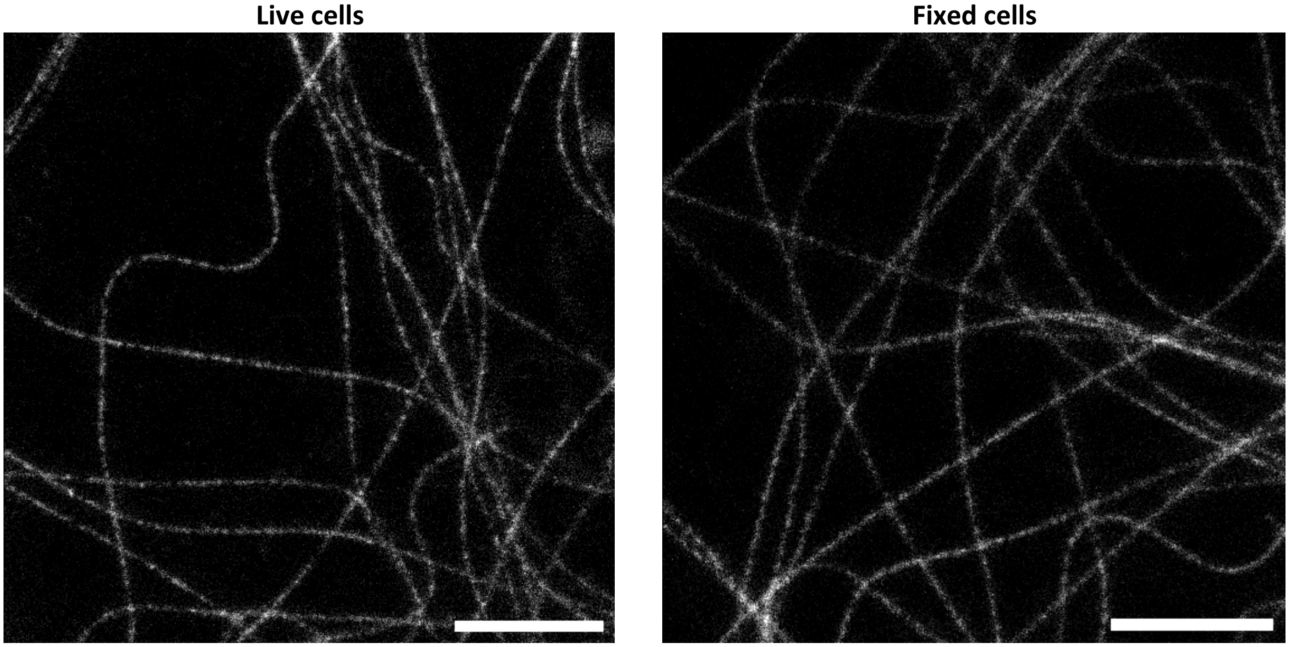

**Figure S16. Representative images used to measure apparent microtubule FWHM.** Human dermal fibroblasts cells incubated with **6-SiR-*o*-$C_9$-CTX** (1 µM in OptiMEM) for 4h, imaged live or after fixation with glutaraldehyde. Images acquired with Abberior Expert Line. Scale bar = 2 µm.

### Supplementary Tables

| | $\lambda_{abs,max}$ (nm) | $\lambda_{em,max}$ (nm) | QY PBS+0.1 % SDS (%) | Fluorescence lifetime PBS+0.1 % SDS (ns) | Percentage dye open Tubulin/PBS+0.1% SDS (%) | Fl increase Tubulin/PBS |
|---|---|---|---|---|---|---|
| **Probe 1** | 652 | 672 | 59.1 ± 0.5 | 3.94 ± 0.01 | 40.0 ± 2.6 | 7.0 ± 2.2 |
| **Probe 2 – 6-SiR-*o*-$C_9$-CTX** | | | 59.3 ± 1.4 | 3.93 ± 0.01 | 49.0 ± 10.8 | 13.7 ± 1.3 |
| **Probe 3** | | | 60.3 ± 3.2 | 3.94 ± 0.00 | 33.3 ± 5.0 | 8.8 ± 2.8 |
| **Probe 4** | | | 59.4 ± 1.3 | 3.95 ± 0.01 | 14.3 ± 8.0 | 4.1 ± 2.4 |
| **Probe 5** | | | 59.4 ± 1.1 | 3.93 ± 0.02 | 25.7 ± 9.0 | 1.3 ± 0.5 |
| **Probe 6** | | | 58.9 ± 1.2 | 3.90 ± 0.01 | 17.0 ± 10.1 | 5.6 ± 2.9 |

**Table S1. Photophysical properties of the probes (1-6).** Experimental data are averages of three independent experiments (N=3) and presented as means with standard deviations.

| Labeled peptide | Tubulin isotypes and isoforms | Total precursor abundance (MS1 intensity) |
|---|---|---|
| TAVcDIPPR | TUBB2A, TUBB2B, TUBB4A, TUBB4B, TUBB | $2{,}97.10^{6}$ |
| VAVcDIPPR | TUBB3 | $3{,}59.10^{5}$ |
| EIVHIQAGQcGNQIGAK | TUBB2A, TUBB2B, TUBB, TUBB3 | $3{,}57.10^{5}$ |
| EIVHLQAGQcGNQIGAK | TUBB4A, TUBB4B | |
| SIQFVDWcPTGFK | LOC100158003, LOC100127131, TUBA4A | $1{,}59.10^{5}$ |
| AVcmLSNTTAIAEAWAR | LOC100158003, LOC100127131, TUBA3C, TUBA1A, TUBA4A, TUBA8 | $1{,}25.10^{5}$ |
| AVcMLSNTTAIAEAWAR | LOC100158003, LOC100127131, TUBA3C, TUBA1A, TUBA4A, TUBA8 | $6{,}63.10^{4}$ |

**Table S2. Identified labeled peptides in the sample.** Reacted Cys residue is marked in red. m is an oxidized methionine.

| Probe name | Cytotoxicity threshold (nM) |
|---|---|
| **Probe 1** | 31.25 |
| **Probe 2 – 6-SiR-*o*-$C_9$-CTX** | 125 |
| **Probe 3** | 250 |
| **Probe 4** | 250 |
| **Probe 5** | 250 |
| **Probe 6** | 125 |

**Table S3. Cytotoxicity threshold of the probes (1-6).** The indicated probe concentration represents the lowest concentration tested, at which a cytotoxicity effect was observed.

| Figure | Probe | Cell line | Microscope | Objective | Excitation (%)/ STED (%) | Pixel dwell time (µs) | Pixel size (nm) | Emission (nm) | Comment |
|---|---|---|---|---|---|---|---|---|---|
| 3A | 1,2,3,4,5,6 | H. fibro | Leica TCS SP8 | 63x 1.40 | 633 (1) | 0.6 | 90 | 650-710 | |
| S13 | 1,2,3,4,5,6 | U-2 OS | | | | | | | |
| S14 | 1,2,3,4,5,6 | HeLa CCL | | | | | | | |
| 3B | 2 | U-2 OS<br>HeLa CCL | | | | | | | |
| 4B | 2, SiR-CTX | U-2 OS | | | | | | | |
| 5A | 2 | HeLa CCL (live) | Abberior Expert Line | 100x 1.40 | 640 (10)/775(25) | 10 | 20 | 650-720 | |
| | | HeLa CCL (fixed) | | | 640 (5)/775(60) | 4 | 20 | 650-720 | Line accum.: 2 |
| | | U-2 OS (live) | | | 640 (10)/775(25) | 10 | 20 | 650-720 | |
| | | U-2 OS (fixed) | | | 640 (10)/775(70) | 4 | 20 | 650-720 | Line accum.: 2 |
| | | H. fibro (live) | | | 640 (17)/ 775(40) | 3 | 20 | 650-720 | |
| 5B | 2 | H. fibro (fixed) | Abberior Expert Line | 100x 1.40 | 640 (10)/775(70) | 4 | 20 | 650-720 | Line accum.: 2 |
| 5D<br>S16 | 2 | H. fibro (fixed) | Abberior Expert Line | 100x 1.40 | 640 (10)/775(75) | 1 | 15 | 650-720 | Line accum.: 3 |
| | | H. fibro (live) | | | 640 (15)/775(75) | 1 | 15 | 650-720 | Line accum.: 3 |
| Movie 2 | 2 | H. fibro (live) | Abberior Facility Line | 60x 1.40 | 640 (5)/775(10) | 5 | 30 | 660-750 | One frame every 25 sec |

**Table S4. Acquisition parameters for microscopy images**

### Supplementary methods

##### Determination of absolute quantum yields

All reported absolute fluorescence quantum yield values (Φ) were measured using a Quantaurus-QY spectrometer (model C11374-01, Hamamatsu Photonics). This instrument uses an integrating sphere to determine photons absorbed and emitted by a sample. Measurements were carried out using dilute samples in air saturated solvents at 25°C at concentrations ranging from $10^{-6}$ to $10^{-7}$ M (A < 0.1 as indicated in the user's manual) and by using 3 mL quartz cuvettes (Hamamatsu Photonics Art. No. A10095-02) provided by the instrument supplier. The fluorescence quantum yields were measured in PBS buffer containing 0.1% SDS. Reported values are averages (N = 3) with standard deviation.

##### Determination of fluorescence lifetimes

The fluorescence decay characteristics of the solution samples in PBS containing 0.1% SDS at concentrations ranging from $10^{-6}$ to $10^{-7}$ M were recorded using a fluorescence lifetime measurement system (Quantaurus-Tau, Hamamatsu Photonics) in 3 mL high performance quartz glass cuvettes (Hellma Analytics Art. No. 101-10-K-40). The decay profile was registered for 53 ns interval after excitation and the experiment was continued until 10 000 peak count was reached. The instrument response function was obtained by using diluted LUDOX® TM-50 colloidal silica (Sigma Aldrich, # 420778). The analysis of the obtained fluorescence decay profile was performed using the instrument software. Reported values are averages (N = 3) with standard deviation.

##### Maintenance and preparation of the cells

Human primary dermal fibroblasts (Lonza, #CC-2511) were cultured in high-glucose DMEM (Thermo Fisher, #31053044) with 10% FBS (Thermo Fisher, #10082147) supplemented with 1 mM Sodium pyruvate (Sigma, #S8636), 1% GlutaMax (Thermo Fisher, #35050038) and 1% Penicillin-Streptomycin (Sigma, #P0781) in a humidified 5% $CO_2$ incubator at 37 °C. The cells were split every 3-4 days or at confluence.

HeLa (ATCC, CCL-2) cells were cultured in high-glucose DMEM (Thermo Fisher, #31966047) with 10% FBS (BioSELL, #S0615) supplemented with 1% Penicillin-Streptomycin (Sigma, #P0781) in a humidified 5% $CO_2$ incubator at 37 °C. The cells were split every 3-4 days or at confluence.

U-2 OS cells (ATCC, HTB-96) were cultured in McCoys 5A medium (Thermo Fisher, #16600082) with 10% FBS (BioSELL, #S0615) supplemented with 1 mM Sodium pyruvate (Sigma, #S8636) and 1% of Penicillin-Streptomycin (Sigma #P0781) in a humidified 5% $CO_2$ incubator at 37 °C. The cells were split every 3-4 days or at confluence.

Confocal and STED microscopy experiments were performed using µ-Slide 8 Well Glass Bottom dishes (Ibidi, #80827).

##### Chemical Synthesis of the building blocks

Reagents were purchased as reagent-grade (from Sigma-Aldrich, BLD Pharm, Enamine, abcr, TCI or Aaron Chem) and used without further purification. 6-SiR-$CO_2$H was synthesized according to published procedure.[2] Flash chromatographies were carried out on Biotage® Selekt with Biotage® Sfär Silica HC (20 µm) columns.

NMR spectra were recorded at 25 °C with an Agilent 400-MR spectrometer at 400 MHz ($^1$H) and 101 MHz ($^{13}$C). Chemical shifts (δ), which are expressed in part per million (ppm), were determined relative to residual non-deuterated solvent as an internal reference: $CDCl_3$ ($^{13}$C NMR: δ = 77.16 ppm; $^1$H NMR: δ = 7.26 ppm) and $(CD_3)_2SO$ ($^{13}$C NMR: δ = 39.52 ppm; $^1$H NMR: δ = 2.50 ppm) or with an external reference ($^{19}$F: δ = 0 ppm for $CFCl_3$). Multiplicities of signals are described as follows: s = singlet, d = doublet, t = triplet, q = quartet, m = multiplet or overlap of non-equivalent resonances; br = broad signal. Coupling constants (*J*) are given in Hz.

ESI-MS were recorded on a Varian 500-MS spectrometer (Agilent). ESI-HRMS were recorded on a MICROTOF spectrometer (Bruker) equipped with ESI ion source (Apollo) and direct injector with LC autosampler Agilent RR 1200.

Analytical LC–MS analysis was performed on an Agilent 1260 Infinity II LC/MS system equipped with an Autosampler (G7129A), Binary Pump (G7112B), Diode Array Detector WR (G7115A), Fluorescence Detector Spectra and Single Quadrupole MSD XT (G6135B). Analysis was done by using a Ascentis® Express AQ-C18 UHPLC Column 2 µm, 5 cm x 2.1 mm with Mobile phase A: 25 mM $HCOONH_4$ (pH = 3.5) aqueous buffer/ Mobile Phase B: MeOH and the following gradient (Flow: 0.4 mL/min, 40°C):

| Time (min) | %B |
|---|---|
| **0** | 40 |
| **2** | 40 |
| **8** | 100 |
| **9** | 100 |
| **10** | 40 |
| **15** | 40 |

Preparative HPLC was performed on a combined Agilent1260/1290 Infinity II preparative system equipped with a 1290 Infinity II open-bed sampler (G7169B)/fraction collector (G7159B), 1260 Infinity II multiple wavelength detector (G7165A) and with:

Device A: 1260 Infinity II preparative binary pump (G7161A) and Agilent 5 Prep-$C_{18}$, 5 µm, 100 x 50 mm preparative column.

Device B: 1290 Infinity II preparative binary pump (G7161B) and Agilent Pursuit 10 $C_{18}$, 10 µm, 250 X 50 mm preparative column.

**Compound *p*-7, *tert*-butyl (4-(mercaptomethyl)benzyl)carbamate :**

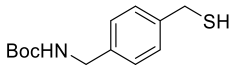

Chemical Formula: $C_{13}H_{19}NO_2S$
Molecular Weight: 253,36

***tert*-Butyl-4-(bromomethyl)benzylcarbamate (2.5 g, 8.32 mmol, 1.0 eq.)** and **thiourea (634 mg, 8.32 mmol, 1.0 eq.)** were refluxed together in **EtOH (21 mL, 0.4M)** for 2h.Water (1 mL) and **NaOH (664 mg, 16.6 mmol, 2.0 eq.)** were added. After 1h at 80°C, the mixture was diluted in water, the pH was lower to 2-3 using aqueous HCl 1M. The aqueous layer was extracted twice with AcOEt. Organic layers were combined, washed with brine and dried over $Na_2SO_4$. The crude was purified by flash chromatography on silica gel (120g, Hexanes/AcOEt from 90:10 to 60:40) to give the expected product as **a white solid (1.63 g, 6.44 mmol, η = 77 %**).

**$^1$H NMR (400 MHz, $CDCl_3$):** δ (ppm) = 7.28 (d, *J* = 8.0 Hz, 2H), 7.23 (d, *J* = 8.0 Hz, 2H), 4.81 (br s, 1H), 4.29 (br s, 2H), 3.72 (d, *J* = 7.5 Hz, 2H), 1.74 (t, *J* = 7.5 Hz, 1H), 1.46 (s, 9H).

**$^{13}$C NMR (101 MHz, $CDCl_3$):** δ (ppm) = 156.0 ($C_q$), 140.4 ($C_q$), 137.9 ($C_q$), 128.4 (CH), 127.9 (CH), 79.7 ($C_q$), 44.5 ($CH_2$), 28.8 ($CH_2$), 28.6 ($CH_3$).

**ESI-MS, positive mode** : m/z = 276.1 $[M+Na]^+$

**ESI-MS, negative mode** : m/z = 252.1 $[M-H]^-$

**HRMS (ESI)** calculated for $C_{13}H_{19}NO_2SNa^+$ $[M+Na]^+$ : 276.1029, found: 276.1031.

**Compound *m*-7, *tert*-butyl (3-(mercaptomethyl)benzyl)carbamate:**

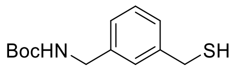

Chemical Formula: $C_{13}H_{19}NO_2S$
Molecular Weight: 253,36

***tert*-Butyl-3-(bromomethyl)benzylcarbamate (2.0 g, 6.66 mmol, 1.0 eq.)** and **thiourea (507 mg, 6.66 mmol, 1.0 eq.)** were refluxed together in **EtOH (17 mL, 0.4 M)** for 2h. **Water (1 mL)** and **NaOH (533 mg, 13.3 mmol, 2.0 eq.)** were added. After 1h at 80°C, the mixture was diluted in water, the pH was lower to 2-3 using aqueous HCl 1M. The aqueous layer was extracted twice with AcOEt. Organic layers were combined, washed with brine and dried over $Na_2SO_4$. The crude was purified by flash chromatography on silica gel (50 g, Hexanes/AcOEt from 100:0 to 80:20) to give the expected product as **white crystals (1450 mg, 5.72 mmol, η=86%).**

**$^{1}$H NMR (400 MHz, $CDCl_3$):** δ (ppm) = 7.35 – 7.17 (m, 3H), 7.17 – 7.13 (m, 1H), 4.90 (s, 1H), 4.27 (s, 2H), 3.70 (d, *J* = 7.5 Hz, 2H), 1.75 (t, *J* = 7.5 Hz, 1H), 1.45 (s, 9H).

**$^{13}$C NMR (101 MHz, $CDCl_3$):** δ (ppm) = 156.0 ($C_q$), 141.5 ($C_q$), 139.5 ($C_q$), 129.0 (CH), 127.1 (CH), 127.0 (CH), 126.2 (CH), 79.6 ($C_q$), 44.7 ($CH_2$), 28.9 ($CH_2$), 28.5 ($CH_3$).

**ESI-MS, positive mode** : m/z = 276.1 $[M+Na]^+$

**ESI-MS, negative mode** : m/z = 252.1 $[M-H]^-$

**HRMS (ESI)** calculated for $C_{13}H_{20}NO_2S^+$ $[M+H]^+$ : 254.1209, found: 254.1213.

#### Compound *o*-7, *tert*-butyl (2-(mercaptomethyl)benzyl)carbamate:

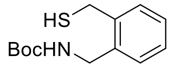

Chemical Formula: $C_{13}H_{19}NO_2S$
Molecular Weight: 253,36

***tert*-Butyl-2-(bromomethyl)benzylcarbamate (2.0 g, 6.66 mmol, 1.0 eq.)** and **thiourea (507 mg, 6.66 mmol, 1.0 eq.)** were refluxed together in **EtOH (17 mL, 0.4M)** for 2h. Water (1 mL) **and NaOH (533 mg, 13.3 mmol, 2.0 eq.)** were added. After 1h at 80°C, the mixture was diluted in water, the pH was lower to 2-3 using aqueous HCl 1M. The aqueous layer was extracted twice with AcOEt. Organic layers were combined, washed with brine and dried over $Na_2SO_4$. The crude was purified by flash chromatography on silica gel (80 g, Hexanes/AcOEt from 100:0 to 80:20) to give the expected product as **white crystals (1500 mg, 5.92 mmol, η=90%).**

**$^{1}$H NMR (400 MHz, $CDCl_3$):** δ (ppm) = 7.46 – 7.15 (m, 4H), 4.96 (s, 1H), 4.42 (s, 2H), 3.78 (d, *J* = 7.0 Hz, 2H), 1.79 (t, *J* = 7.0 Hz, 1H), 1.47 (s, 9H).

**$^{13}$C NMR (101 MHz, $CDCl_3$):** δ (ppm) = 155.8 ($C_q$), 139.2 ($C_q$), 136.3 ($C_q$), 129.5 (CH), 129.2 (CH), 128.1 (CH), 127.8 (CH), 79.7 ($C_q$), 42.1 ($CH_2$), 28.5 ($CH_3$), 26.1 ($CH_2$).

**ESI-MS, positive mode** : m/z = 276.1 $[M+Na]^+$

**ESI-MS, negative mode** : m/z = 252.1 $[M-H]^-$

**HRMS (ESI)** calculated for $C_{13}H_{20}NO_2S^+$ $[M+H]^+$ : 254.1209, found: 254.1219.

#### Compound 8:

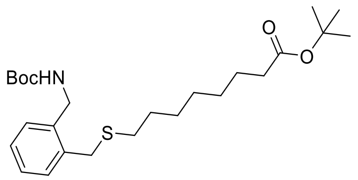

Chemical Formula: $C_{25}H_{41}NO_4S$
Molecular Weight: 451,66

Under Ar atmosphere, ***o*-7 (300 mg, 1.18 mmol, 1.0 eq.)** and **potassium carbonate (546 mg, 2.37 mmol, 2.0 eq.)** were mixed together in **dry DMF (12 mL, 0.1M).** ***Tert*-butyl 8-bromooctanoate (497 mg, 1.78 mmol, 1.5 eq.)** was added and the solution was stirred at rt. After 18h, water (25 mL) and AcOEt (30 mL) were added. The aqueous layer was extracted twice with AcOEt. Organic layers were combined, was washed with brine and dried over $Na_2SO_4$. The crude was purified by flash column chromatography on silica gel (10g, Hexanes/AcOEt from 100:0 to 85:15) to give the expected product as a colorless oil **(493 mg, 1.09 mmol, η = 92%)**.

**$^1$H NMR (400 MHz, CDCl$_3$):** δ (ppm) = 7.34 – 7.30 (m, 1H), 7.25 – 7.17 (m, 3H), 5.08 (s, 1H), 4.41 (s, 2H), 3.73 (s, 2H), 2.49 – 2.40 (m, 2H), 2.18 (t, *J* = 7.5 Hz, 2H), 1.63 – 1.51 (m, 4H), 1.44 (s, 9H), 1.43 (s, 9H), 1.36 – 1.20 (m, 6H).

**$^{13}$C NMR (101 MHz, CDCl$_3$):** δ (ppm) = 173.3 ($C_q$), 155.9 ($C_q$), 137.2 ($C_q$), 136.1 ($C_q$), 130.3 (CH), 129.3 (CH), 127.7 (CH), 127.5 (CH), 80.0 ($C_q$), 79.5 ($C_q$), 42.1 ($CH_2$), 35.6 ($CH_2$), 33.9 ($CH_2$), 29.3 ($CH_2$), 29.0 ($CH_2$), 29.0 ($CH_2$), 28.8 ($CH_2$), 28.5 ($CH_3$), 28.2 ($CH_3$), 25.1 ($CH_2$).

**ESI-MS, positive mode** : m/z = 452.2 $[M+H]^+$

**HRMS (ESI)** calculated for $C_{25}H_{42}NO_4S^+$ $[M+H]^+$ : 452.2829, found: 452.2841.

**Compound 9:**

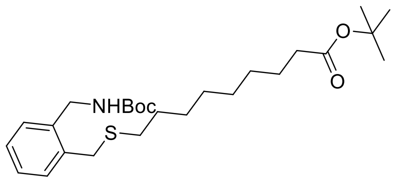

Chemical Formula: $C_{26}H_{43}NO_4S$
Molecular Weight: 465,69

Under Ar atmosphere, ***o*-7 (150 mg, 0.59 mmol, 1.0 eq.)** and **potassium carbonate (163 mg, 1.18 mmol, 2.0 eq.)** were mixed together in **dry DMF (6 mL, 0.1M).** ***Tert*-butyl 9-bromononanoate (249 mg, 0.89 mmol, 1.5 eq.)** was added and the solution was stirred at rt for 24h. AcOEt was added (20 mL). The organic layer was washed once with brine and dried over $Na_2SO_4$. The crude was purified by flash column chromatography on silica gel (10g, Hexanes/AcOEt from 100:0 to 85:15) to give the expected product as a colorless oil **(217 mg, 0.592 mmol, η = 79%)**.

**$^1$H NMR (400 MHz, CDCl$_3$):** δ (ppm) = 7.35 – 7.31 (m, 1H), 7.26 – 7.17 (m, 3H), 5.06 (s, 1H), 4.42 (s, 2H), 3.74 (s, 2H), 2.49 – 2.42 (m, 2H), 2.19 (t, J = 7.5 Hz, 2H), 1.62 – 1.53 (m, 4H), 1.45 (s, 9H), 1.44 (s, 9H), 1.38 – 1.23 (m, 8H).

**$^{13}$C NMR (101 MHz, CDCl$_3$):** δ (ppm) = 173.4 ($C_q$), 155.9 ($C_q$), 137.3 ($C_q$), 136.2 ($C_q$), 130.4 (CH), 129.4 (CH), 127.8 (CH), 127.5 (CH), 80.0 ($C_q$), 79.6 ($C_q$), 42.2 ($CH_2$), 35.7 ($CH_2$), 33.9 ($CH_2$), 32.1 ($CH_2$), 29.4 ($CH_2$), 29.3 ($CH_2$), 29.2 ($CH_2$), 29.1 ($CH_2$), 29.0, ($CH_2$) 28.6 ($CH_3$), 28.3 ($CH_3$), 25.2 ($CH_2$).

**ESI-MS, positive mode** : m/z = 466.2 $[M+H]^+$

**HRMS (ESI)** calculated for $C_{26}H_{44}NO_4S^+$ $[M+H]^+$ : 466.2986, found: 466.2988.

**Compound 10:**

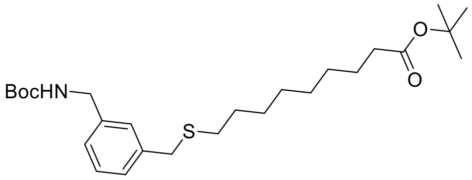

Chemical Formula: $C_{26}H_{43}NO_4S$
Molecular Weight: 465,69

Under Ar atmosphere, ***m*-7 (300 mg, 1.18 mmol, 1.0 eq.)** and **potassium carbonate (163 mg, 1.18 mmol, 2.0 eq.)** were mixed together in **dry DMF (6 mL, 0.1M). Tert-butyl 9-bromononanoate (249 mg, 0.79 mmol, 1.5 eq.)** was added and the solution was stirred at rt for 24h. AcOEt was added (20 mL). The organic layer was washed once with brine and dried over $Na_2SO_4$. The organic layer was washed once with brine and dried over $Na_2SO_4$. The crude was purified by flash column chromatography on silica gel (10g, Hexanes/AcOEt from 100:0 to 85:15) to give the expected product as a colorless oil **(256 mg, η = 93%)**.

**$^1$H NMR (400 MHz, CDCl$_3$):** δ (ppm) = 7.27 – 7.11 (m, 4H), 4.89 (s, 1H), 4.27 (s, 2H), 3.66 (s, 2H), 2.42 – 2.35 (m, 2H), 2.17 (t, J = 7.5 Hz, 2H), 1.60 – 1.49 (m, 4H), 1.44 (s, 9H), 1.42 (s, 9H), 1.38 – 1.21 (m, 8H).

**$^{13}$C NMR (101 MHz, CDCl$_3$):** δ (ppm) = 173.3 (C$_q$), 156.0 (C$_q$), 139.3 (C$_q$), 139.1 (C$_q$), 128.8 (CH), 127.9 (CH), 127.9 (CH), 126.1 (CH), 80.0 (C$_q$), 79.6 (C$_q$), 44.7 (CH$_2$), 36.3 (CH$_2$), 35.7 (CH$_2$), 31.6 (CH$_2$), 29.2 (CH$_2$), 29.2 (CH$_2$), 29.1 (CH$_2$), 29.1 (CH$_2$), 28.9 (CH$_2$), 28.5 (CH$_3$), 28.2 (CH$_3$), 25.1 (CH$_2$).

**ESI-MS, positive mode** : m/z = 466.3 [M+H]$^+$

**HRMS (ESI)** calculated for $C_{26}H_{44}NO_4S^+$ [M+H]$^+$ : 466.2986, found: 466.2992.

**Compound 11:**

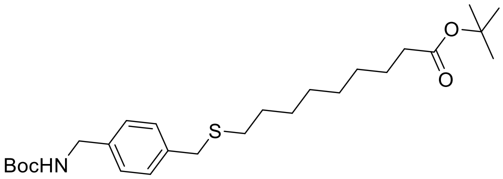

Chemical Formula: $C_{26}H_{43}NO_4S$
Molecular Weight: 465,69

Under Ar atmosphere, ***p*-7 (150 mg, 0.59 mmol, 1.0 eq.)** and **potassium carbonate (163 mg, 1.18 mmol, 2.0 eq.)** were mixed together in **dry DMF (6 mL, 0.1M). Tert-butyl 9-bromononanoate (249 mg, 0.89 mmol, 1.5 eq.)** was added and the solution was stirred at rt for 24h. AcOEt was added (20 mL). The organic layer was washed once with brine and dried over $Na_2SO_4$. The crude was purified by flash column chromatography on silica gel (10g, Hexanes/AcOEt from 100:0 to 80:20) to give the expected product as a colorless oil **(227 mg, 0.59 mmol, η = 82%)**.

**$^1$H NMR (400 MHz, CDCl$_3$):** δ (ppm) = 7.22 (d, *J* = 8.0 Hz, 2H), 7.17 (d, *J* = 8.0 Hz, 2H), 4.95 (s, 1H), 4.24 (s, 2H), 3.63 (s, 2H), 2.38 – 2.31 (m, 2H), 2.15 (t, *J* = 7.5 Hz, 2H), 1.56 – 1.47 (m, 4H), 1.42 (s, 9H), 1.41 (s, 9H), 1.32 – 1.20 (m, 8H).

**$^{13}$C NMR (101 MHz, CDCl$_3$):** δ (ppm) = 173.2 (C$_q$), 155.9 (C$_q$), 137.7 (C$_q$), 137.6 (C$_q$), 129.0 (CH), 127.5 (CH), 79.9 (C$_q$), 79.4 (C$_q$), 44.4 (CH$_2$), 35.9 (CH$_2$), 35.6 (CH$_2$), 31.3 (CH$_2$), 29.1 (CH$_2$), 29.1 (CH$_2$), 29.0 (CH$_2$), 29.0 (CH$_2$), 28.8 (CH$_2$), 28.4 (CH$_3$), 28.1 (CH$_3$), 25.0 (CH$_2$).

**ESI-MS, positive mode** : m/z = 466.2 [M+H]$^+$

**HRMS (ESI)** calculated for $C_{26}H_{43}NO_4SNa^+$ [M+Na]$^+$ : 488.2805, found: 488.2804.

**Compound 12:**

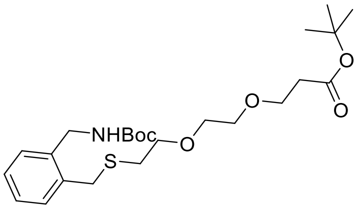

Chemical Formula: $C_{24}H_{39}NO_6S$
Molecular Weight: 469,63

Under Ar atmosphere, ***o*-7 (100 mg, 0.39 mmol, 1.0 eq.)** and **potassium carbonate (108 mg, 0.78 mmol, 2.0 eq.)** were mixed together in **dry DMF (4 mL, 0.1M). Bromo-PEG$_2$- *tert*-butyl ester (129 mg, 0.43 mmol, 1.1 eq.)** was added and the solution was stirred at rt for 18h. AcOEt (20 mL) was added. The organic layer was washed twice with brine and dried over $Na_2SO_4$. The crude was purified by flash column chromatography on silica gel (10g, Hexanes/AcOEt from 70:30 to 0:100) to give the expected product as a colorless oil **(140 mg, 0.30 mmol, η = 76 %)**.

**$^{1}$H NMR (400 MHz, CDCl$_3$):** δ (ppm) = 7.36 – 7.32 (m, 1H), 7.27 – 7.17 (m, 3H), 4.43 (d, *J* = 3.5 Hz, 2H), 3.84 (s, 2H), 3.71 (t, *J* = 6.5 Hz, 2H), 3.65 – 3.56 (m, 6H), 2.66 (t, *J* = 6.5 Hz, 2H), 2.50 (t, *J* = 6.5 Hz, 2H), 1.46 (s, 9H), 1.44 (s, 9H).

**$^{13}$C NMR (101 MHz, CDCl$_3$):** δ (ppm) = 171.0 ($C_q$), 155.9 ($C_q$), 137.4 ($C_q$), 136.0 ($C_q$), 130.5 (CH), 129.5 (CH), 127.9 (CH), 127.6 (CH), 80.7 ($C_q$), 79.7 ($C_q$, from HMBC), 71.2 ($CH_2$), 70.5 ($CH_2$), 70.4 ($CH_2$), 67.0 ($CH_2$), 42.1 ($CH_2$), 36.4 ($CH_2$), 34.3 ($CH_2$), 31.3 ($CH_2$), 28.6 ($CH_3$), 28.2 ($CH_3$).

**ESI-MS, positive mode** : m/z = 470.2 $[M+H]^+$

**HRMS (ESI)** calculated for $C_{24}H_{39}NO_6SNa^+$ $[M+Na]^+$: 492.2390, found: 492.2392.

#### Compound 13:

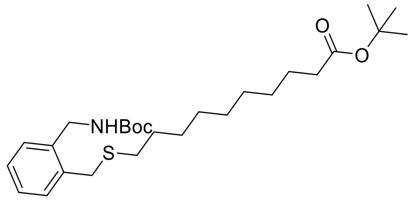

Chemical Formula: $C_{27}H_{45}NO_4S$
Molecular Weight: 479,72

Under Ar atmosphere, ***o*-7 (100 mg, 0.40 mmol, 1.0 eq.)** and **potassium carbonate (109 mg, 0.79 mmol, 2.0 eq.)** were mixed together in **dry DMF (4 mL, 0.1M). Tert-butyl 10-bromodecanoate (147 mg, 0.47 mmol, 1.2 eq.)** was added and the solution was stirred at rt for 24h. AcOEt (20 mL) was added. The organic layer was washed once with brine and dried over $Na_2SO_4$. The crude was purified by flash column chromatography on silica gel (10g, Hexanes/AcOEt from 100:0 to 80:20) to give the expected product as a colorless oil **(133 mg, 0.27 mmol, η = 70%)**.

**$^{1}$H NMR (400 MHz, CDCl$_3$):** δ (ppm) = 7.34 – 7.31 (m, 1H), 7.25 – 7.16 (m, 3H), 5.05 (s, 1H), 4.42 (s, 2H), 3.74 (s, 2H), 2.48 – 2.42 (m, 2H), 2.21 – 2.15 (m, 2H), 1.61 – 1.51 (m, 4H), 1.45 (s, 9H), 1.43 (s, 9H), 1.36 – 1.23 (m, 10H).

**$^{13}$C NMR (101 MHz, CDCl$_3$):** δ (ppm) = 173.4 ($C_q$), 155.9 ($C_q$), 137.2 ($C_q$), 136.2 ($C_q$), 130.4 (CH), 129.4 (CH), 127.8 (CH), 127.5 (CH), 80.0 ($C_q$), 79.6 ($C_q$), 42.2 ($CH_2$), 35.7 ($CH_2$), 33.9 ($CH_2$), 32.1 ($CH_2$), 29.4 ($CH_2$), 29.3 ($CH_2$), 29.3 ($CH_2$), 29.2 ($CH_2$), 29.0 ($CH_2$), 28.6 ($CH_3$), 28.2 ($CH_3$), 25.2 ($CH_2$).

**ESI-MS, positive mode** : m/z = 480.3 $[M+H]^+$

**HRMS (ESI)** calculated for $C_{27}H_{46}NO_4S^+$ $[M+H]^+$ : 480.3142, found: 480.3142.

#### General procedure for sulfonium synthesis:

**The thioether (1.0 eq)** was solubilized in **a HCOOH/AcOH mixture (1:1, 0.5M)**. **Methyl trifluoromethanesulfonate (3.0 eq.)** was added and the solution was stirred at rt for 1h. The mixture was diluted in acetonitrile and purified by reverse-phase HPLC (Device A, $H_2O$+0.1%TFA/ACN, linear gradient from 95:5 to 0:100, 40 mL/min). Solvents were removed under vacuum. As this type of compounds is highly hygroscopic, the final product was dissolved in a precise amount of **DMSO-d$_6$.** Concentration of the samples and therefore yields were determined by $^{1}$H-NMR with an external standard. The products were kept in DMSO-d$_6$ (stored with preactivated powdered 4Å molecular sieves) and use in the final step without any further treatments.

#### Compound 14:

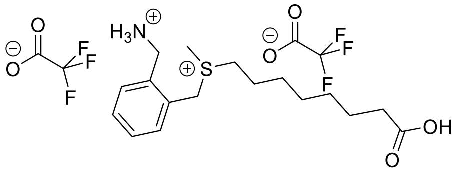

Chemical Formula: $C_{21}H_{29}F_6NO_6S$
Molecular Weight: 537,51

The general procedure starting with **8 (70 mg, 0.155 mmol, 1.0 eq.)** gave the expected product as **a colorless oil (0.140 mmol, η = 90 %).**

**$^{1}$H NMR (400 MHz, DMSO-d$_6$):** δ(ppm) = 8.53 (s, 3H), 7.63 (dd, *J* = 7.5, 1.5 Hz, 1H), 7.58 – 7.44 (m, 3H), 4.92 (d, *J* = 13.0 Hz, 1H), 4.83 (d, *J* = 13.0 Hz, 1H), 4.27 – 4.23 (m, 2H), 3.40 – 3.29 (m, 2H), 2.87 (s, 3H), 2.18 (t, *J* = 7.5 Hz, 2H), 1.74 – 1.63 (m, 1H), 1.60 – 1.44 (m, 3H), 1.36 – 1.20 (m, 6H).

**$^{13}$C NMR (101 MHz, DMSO-d$_6$):** δ(ppm) = 174.5 ($C_q$), 158.8 (q, $J_F$ = 34.2 Hz), 134.4 ($C_q$), 132.3 (CH), 131.0 (CH), 130.2 (CH), 129.5 (CH), 127.8 ($C_q$), 116.4 (q, $J_F$ = 294.5 Hz), 42.8 ($CH_2$), 40.9 ($CH_2$), 38.9 ($CH_2$), 33.7 ($CH_2$), 28.2 ($CH_2$), 28.0 ($CH_2$), 27.7 ($CH_2$), 24.4 ($CH_2$), 23.4 ($CH_2$), 22.0 ($CH_3$).

**$^{19}$F NMR (376 MHz, DMSO-d$_6$):** δ(ppm) = -74.5.

**ESI-MS, positive mode**: m/z = 310.1 $[M-H-2CF_3CO_2]^+$

**ESI-MS, negative mode**: m/z = 308.1 $[M-2H-2CF_3CO_2]^-$

**HRMS (ESI)** calculated for $C_{17}H_{28}NO_2S^+$ $[M-H-2CF_3CO_2]^+$: 310.1835, found: 310.1836.

**Compound 15:**

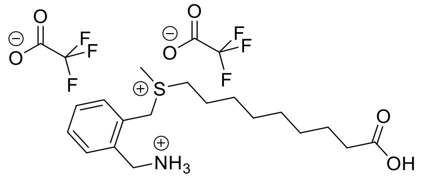

Chemical Formula: $C_{22}H_{31}F_6NO_6S$
Molecular Weight: 551,54

The general procedure starting with **9 (100 mg, 0.215 mmol, 1.0 eq.)** gave the expected product as **a colorless oil (0.210 mmol, η = 98 %).**

**$^{1}$H NMR (400 MHz, DMSO-d$_6$):** δ(ppm) = 8.55 (s, 3H, NH), 7.62 (dd, *J* = 7.5, 1.5 Hz, 1H), 7.57 – 7.44 (m, 3H), 4.92 (d, *J* = 13.0 Hz, 1H), 4.83 (d, *J* = 13.0 Hz, 1H), 4.24 (t, *J* = 5.5 Hz, 2H), 3.38 – 3.29 (m, 2H), 2.86 (s, 3H), 2.18 (t, *J* = 7.5 Hz, 2H), 1.76 – 1.60 (m, 1H), 1.59 - 1.43 (m, 3H), 1.34 – 1.19 (m, 8H).

**$^{13}$C NMR (101 MHz, DMSO-d$_6$):** δ(ppm) = 174.6 ($C_q$), 158.8 (q, $J_F$ = 34.3 Hz), 134.5 ($C_q$), 132.3 (CH), 131.0 (CH), 130.2 (CH), 129.5 (CH), 127.8 ($C_q$), 116.5 (q, $J_F$ = 294.7 Hz), 42.9 ($CH_2$), 40.9 ($CH_2$), 38.9 ($CH_2$), 33.8 ($CH_2$), 28.5 ($CH_2$), 28.4 ($CH_2$), 28.2 ($CH_2$), 27.8 ($CH_2$), 24.5 ($CH_2$), 23.4 ($CH_2$), 22.0 ($CH_3$).

**$^{19}$F NMR (376 MHz, DMSO-d$_6$):** δ(ppm) = -74.5.

**ESI-MS, positive mode**: m/z = 324.2 $[M-H-2CF_3CO_2]^+$

**ESI-MS, negative mode**: m/z = 322.2 $[M-2H-2CF_3CO_2]^-$

**HRMS (ESI)** calculated for $C_{18}H_{30}NO_2S^+$ $[M-H-2CF_3CO_2]^+$: 324.1992, found: 324.1993.

**Compound 16:**

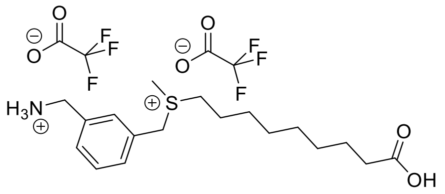

The general procedure starting with **10 (100 mg, 0.215 mmol, 1.0 eq.)** gave the expected product as **a colorless oil (0.210 mmol, η = 98 %).**

**$^{1}$H NMR (400 MHz, DMSO-d$_6$):** δ(ppm) = 8.49 (s, 3H), 7.61 – 7.48 (m, 4H), 4.78 (d, *J* = 13.0 Hz, 1H), 4.69 (d, *J* = 13.0 Hz, 1H), 4.08 (q, *J* = 5.5 Hz, 2H), 3.33 – 3.20 (m, 2H), 2.81 (s, 3H), 2.19 (t, *J* = 7.5 Hz, 2H), 1.78 – 1.59 (m, 2H), 1.55 – 1.42 (m, 2H), 1.40 – 1.20 (m, 8H).

**$^{13}$C NMR (101 MHz, DMSO-d$_6$):** δ(ppm) = 174.6 ($C_q$), 158.8 (q, $J_F$ = 35.1 Hz), 135.4 ($C_q$), 131.2 (CH), 130.9 (CH), 130.3 (CH), 129.7 (CH), 128.8 ($C_q$), 116.2 (q, $J_F$ = 293.1 Hz), 44.4 ($CH_2$), 42.1 ($CH_2$), 40.6 ($CH_2$), 33.8 ($CH_2$), 28.5 ($CH_2$), 28.5 ($CH_2$), 28.2 ($CH_2$), 27.8 ($CH_2$), 24.6 ($CH_2$), 23.4 ($CH_2$), 21.6 ($CH_3$).

**$^{19}$F NMR (376 MHz, DMSO-d$_6$):** δ(ppm) = -74.7.

**ESI-MS, positive mode** : m/z = 324.2 $[M\text{-}H\text{-}2CF_3CO_2]^+$

**ESI-MS, negative mode** : m/z = 322.2 $[M\text{-}2H\text{-}2CF_3CO_2]^-$

**HRMS (ESI)** calculated for $C_{18}H_{30}NO_2S^+$ $[M\text{-}H\text{-}2CF_3CO_2]^+$: 324.1992, found: 324.1992.

**Compound 17:**

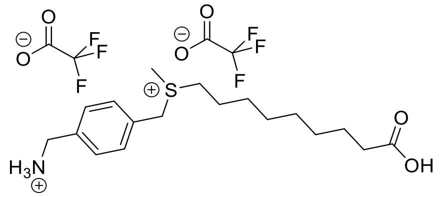

The general procedure starting with **11 (100 mg, 0.215 mmol, 1.0 eq.)** gave the expected product as **a colorless oil (0.196 mmol, η = 91 %).**

**$^{1}$H NMR (400 MHz, DMSO-d$_6$):** δ(ppm) = 8.49 (s, 3H), 7.62 – 7.48 (m, 4H), 4.78 (d, *J* = 13.0 Hz, 1H), 4.71 (d, *J* = 13.0 Hz, 1H), 4.08 (q, *J* = 5.5 Hz, 2H), 3.32 – 3.17 (m, 2H), 2.80 (s, 3H), 2.19 (t, *J* = 7.5 Hz, 2H), 1.78 – 1.59 (m, 2H), 1.52 – 1.45 (m, 2H), 1.38 – 1.17 (m, 8H).

**$^{13}$C NMR (101 MHz, DMSO-d$_6$):** δ(ppm) = 174.6 ($C_q$), 158.8 (q, $J_F$ = 34.7 Hz), 135.7 ($C_q$), 130.9 (CH), 129.8 (CH), 128.7 ($C_q$), 116.3 (q, $J_F$ = 293.8 Hz), 44.3 ($CH_2$), 42.0 ($CH_2$), 40.6 ($CH_2$), 33.8 ($CH_2$), 28.5 ($CH_2$), 28.5 ($CH_2$), 28.2 ($CH_2$), 27.8 ($CH_2$), 24.6 ($CH_2$), 23.3 ($CH_2$), 21.7 ($CH_3$).

**$^{19}$F NMR (376 MHz, DMSO-d$_6$):** δ(ppm) = -74,7.

**ESI-MS, positive mode**: m/z = 324.1 $[M\text{-}H\text{-}2CF_3CO_2]^+$

**ESI-MS, negative mode**: m/z = 322.1 $[M\text{-}2H\text{-}2CF_3CO_2]^-$

**HRMS (ESI)** calculated for $C_{18}H_{30}NO_2S^+$ $[M\text{-}H\text{-}2CF_3CO_2]^+$: 324.1992, found: 324.1999.

**Compound 18:**

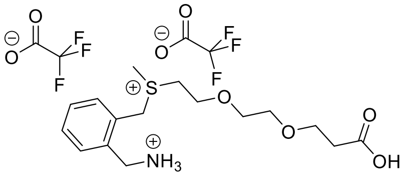

Chemical Formula: $C_{20}H_{27}F_6NO_8S$
Molecular Weight: 555,49

The general procedure starting with **12 (75 mg, 0.160 mmol, 1.0 eq.)** gave the expected product as **a colorless oil (0.088 mmol, η = 55 %).**

**$^1$H NMR (400 MHz, DMSO-d$_6$):** δ(ppm) = 8.47 (s, 3H), 7.65 – 7.60 (m, 1H), 7.59 – 7.44 (m, 3H), 4.93 (d, *J* = 13.0 Hz, 1H), 4.83 (d, *J* = 13.0 Hz, 1H), 4.23 (q, *J* = 5.5 Hz, 2H), 3.91 – 3.78 (m, 2H), 3.71 – 3.43 (m, 8H), 2.88 (s, 3H), 2.43 (t, *J* = 6.0 Hz, 2H).

**$^{13}$C NMR (101 MHz, DMSO-d$_6$):** δ(ppm) = 172.7 ($C_q$), 158.6 (q, $J_F$ = 35.0 Hz, $C_q$), 134.5 ($C_q$), 132.4 (CH), 130.8 (CH), 130.3 (CH), 129.5 (CH), 127.6 ($C_q$), 116.1 (q, $J_F$ = 294.9 Hz, $C_q$), 69.7 ($CH_2$), 69.3 ($CH_2$), 66.3 ($CH_2$), 64.4 ($CH_2$), 43.1 ($CH_2$), 41.6 ($CH_2$), 38.8 ($CH_2$), 34.8 ($CH_2$), 22.5 ($CH_3$).

**$^{19}$F NMR (376 MHz, DMSO-d$_6$):** δ(ppm) = -74.6.

**ESI-MS, positive mode**: m/z = 328.1 $[M-H-2CF_3CO_2]^+$

**ESI-MS, negative mode**: m/z = 326.1 $[M-2H-2CF_3CO_2]^-$

**HRMS (ESI)** calculated for $C_{16}H_{26}NO_4S^+$ $[M-H-2CF_3CO_2]^+$: 325.1577, found: 325.1582.

**Compound 19:**

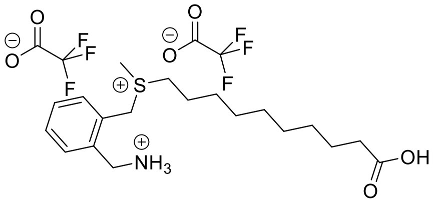

Chemical Formula: $C_{23}H_{33}F_6NO_6S$
Molecular Weight: 565,56

The general procedure starting with **13 (40 mg, 0.083 mmol, 1.0 eq.)** gave the expected product as **a colorless oil (0.080 mmol, η = 96 %).**

**$^1$H NMR (400 MHz, DMSO-d$_6$):** δ(ppm) = 8.52 (s, 3H), 7.66 – 7.59 (m, 1H), 7.59 – 7.42 (m, 3H), 4.90 (d, *J* = 13.0 Hz, 1H), 4.82 (d, *J* = 13.0 Hz, 1H), 4.24 (s, 2H), 3.39 – 3.28 (m, 2H), 2.86 (s, 3H), 2.18 (t, *J* = 7.5 Hz, 2H), 1.73 – 1.63 (m, 1H), 1.60 – 1.42 (m, 3H), 1.36 – 1.28 (m, 2H), 1.27 – 1.20 (m, 8H).

**$^{13}$C NMR (101 MHz, DMSO-d$_6$):** δ(ppm) = 174.5 ($C_q$), 158.6 (q, $J_F$ = 34.9 Hz), 134.3 ($C_q$), 132.3 (CH), 130.9 (CH), 130.2 (CH), 129.4 (CH), 127.7 ($C_q$), 116.2 (q, $J_F$ = 293.6 Hz), 42.7 ($CH_2$), 40.8 ($CH_2$), 38.8 ($CH_2$), 33.7 ($CH_2$), 28.6 ($CH_2$), 28.5 ($CH_2$), 28.5 ($CH_2$), 28.2 ($CH_2$), 27.8 ($CH_2$), 24.5 ($CH_2$), 23.3 ($CH_2$), 21.9 ($CH_3$).

**$^{19}$F NMR (376 MHz, DMSO-d$_6$):** δ(ppm) = -74.6.

**ESI-MS, positive mode**: m/z = 338.2 $[M-H-2CF_3CO_2]^+$

**ESI-MS, negative mode**: m/z = 336.2 $[M-2H-2CF_3CO_2]^-$

**HRMS (ESI)** calculated for $C_{19}H_{32}NO_2S^+$ $[M-H-2CF_3CO_2]^+$: 338.2148, found: 338.2147.

#### CTX-$NH_2$.$CF_3OCO_2H$:

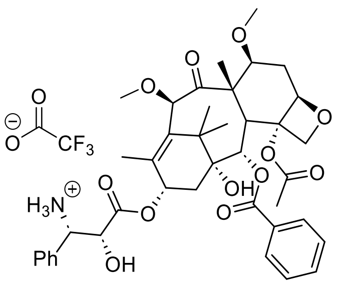

Chemical Formula: $C_{42}H_{50}F_3NO_{14}$
Molecular Weight: 849,85

A solution of **cabazitaxel (1 g, 1.20 mmol, 1.0 eq.)** in **95% formic acid (5 mL)** was stirred at room temperature for 4h. Reaction progress was monitored by analytical HPLC. Once reaction was complete, formic acid was evaporated on rotary evaporator. The residue was solubilized in a $H_2O$/ACN mixture and purified by reverse-phase HPLC (Device B, $H_2O$+0.1%TFA/ACN, linear gradient 70:30 to 0:100, 150 mL/min). The fractions containing the product were lyophilized to obtain the expected compound as a white powder **(660 mg, 0.78 mmol, η = 65 %)**.

**$^1$H NMR (400 MHz, DMSO-d$_6$):** δ(ppm) = 8.71 (s, 3H, $NH_3$), 7.94 – 7.89 (m, 2H), 7.76 – 7.71 (m, 1H), 7.67 – 7.62 (m, 2H), 7.53 – 7.42 (m, 4H), 7.35 – 7.30 (m, 1H), 5.84 (td, *J* = 9.0, 2.0 Hz, 1H), 5.34 (d, *J* = 7.0 Hz, 1H), 4.93 (dd, *J* = 9.5, 2.0 Hz, 1H), 4.67 (s, 1H), 4.49 – 4.38 (m, 2H), 4.01 – 3.96 (m, 2H), 3.72 (dd, *J* = 10.5, 6.5 Hz, 1H), 3.55 (d, *J* = 7.0 Hz, 1H), 3.29 (s, 3H), 3.20 (s, 3H), 2.67 – 2.60 (m, 1H), 2.08 (s, 3H), 1.80 (s, 3H), 1.68 (dd, *J* = 15.5, 9.0 Hz, 1H), 1.57 – 1.42 (m, 5H), 0.97 (s, 3H), 0.94 (s, 3H).

**$^{13}$C NMR (101 MHz, DMSO-d$_6$):** δ(ppm) = 204.7 ($C_q$), 171.2 ($C_q$), 169.8 ($C_q$), 165.1 ($C_q$), 158.3 (q, $J_F$ = 32.0 Hz), 137.9 ($C_q$), 135.1 ($C_q$), 133.5 (CH), 133.4 ($C_q$), 129.9 ($C_q$), 129.5 (CH), 129.3 (CH), 128.9 (CH), 128.7 (CH), 128.0 (CH), 117.0 (q, $J_F$ = 296.5 Hz), 83.2 (CH), 82.0 (CH), 80.3 ($C_q$), 80.2 (CH), 76.8 ($C_q$), 75.2 ($CH_2$), 74.3 (CH), 73.0 (CH), 70.2 (CH), 57.0 (CH), 56.7 ($CH_3$), 56.7 ($CH_3$), 56.0 ($C_q$), 46.4 (CH), 42.9 ($C_q$), 34.7 ($CH_2$), 31.7 ($CH_2$), 26.7 ($CH_3$), 22.5 ($CH_3$), 21.2 ($CH_3$), 14.0 ($CH_3$), 10.2 ($CH_3$).

**$^{19}$F NMR (376 MHz, DMSO-d$_6$):** δ(ppm) = -73.9.

**ESI-MS, positive mode**: m/z = 736.2 [M-$CF_3OCO_2^-$]

**ESI-MS, negative mode**: m/z = 780.30 [M-$CF_3OCO_2H$ + $HCOO^-$]

**HRMS (ESI)** calculated for $C_{40}H_{50}NO_{12}^+$ [M-$CF_3CO_2$]$^+$: 736.3328, found: 736.3347.

##### Characterization of the final probes

#### Probe 1 :

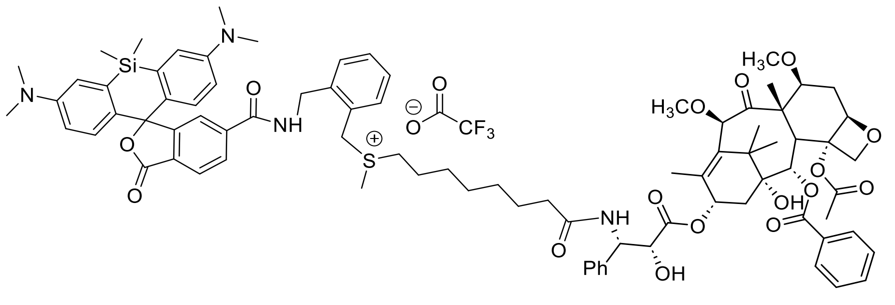

Exact Mass: 1481,67

According to the general procedure, **6-SiR-$CO_2H$** (2.0 mg, 4.23 µmol), sulfonium **14** (13.8 µL -of a 0.4M solution in DMSO-,5.50 µmol) and **CTX-$NH_2$.$CF_3CO_2H$** (10.8 mg, 12.7 µmol) led to the expected product as a blue powder (2.46 µmol, **η = 58 %**).

**$^{1}$H NMR (400 MHz, DMSO-$d_6$):** δ = 9.33 (t, *J* = 5.5 Hz, 1H, NH), 8.36 (d, *J* = 9.0 Hz, 1H, NH), 8.14 (dd, *J* = 8.0, 1.5 Hz, 1H, $H_{Ar}$), 8.07 (d, *J* = 8.0 Hz, 1H, $H_{Ar}$), 8.00 – 7.96 (m, 2H, $H_{Ar}$), 7.72 – 7.66 (m, 2H, $H_{Ar}$), 7.62 – 7.57 (m, 2H, $H_{Ar}$), 7.46 – 7.31 (m, 8H, $H_{Ar}$), 7.24 – 7.20 (m, 1H, $H_{Ar}$), 7.04 (s, 2H, $H_{Ar}$), 6.67 -6.62 (m, 4H, $H_{Ar}$), 5.96 – 5.91 (m, 1H, CH), 5.39 (d, *J* = 7.0 Hz, 1H, CH), 5.32 – 5.27 (m, 1H, CH), 4.98 – 4.94 (m, 1H, CH), 4.86 (d, *J* = 13.0 Hz, 1H, 1H from $CH_2$), 4.76 (d, *J* = 13.0 Hz, 1H, 1H from $CH_2$), 4.71 (s, 1H, CH), 4.56 (d, *J* = 5.5 Hz, 2H, $CH_2$), 4.42 (d, *J* = 5.5 Hz, 1H, CH), 4.03 (s, 2H, $CH_2$), 3.76 – 3.73 (dd, *J* = 10.5, 6.5 Hz, 1H, CH), 3.63 (d, *J* = 7.0 Hz, 1H, CH), 3.33 – 3.26 (m, 5H, $CH_3$ + $CH_2$), 3.21 (s, 3H, $CH_3$), 2.93 (s, 12H, 4*$CH_3$), 2.84 (s, 3H, $CH_3$), 2.68 – 2.63 (m, 1H, 1H from $CH_2$), 2.25 (s, 3H, $CH_3$), 2.19 – 2.11 (td, *J* = 7.0, 4.0 Hz, 2H, $CH_2$), 2.00 – 1.94 (m, 1H, 1H from $CH_2$), 1.90 – 1.81 (m, 4H, $CH_3$ +1H from $CH_2$), 1.60 – 1.41 (m, 8H, 1*$CH_3$ + 2*$CH_2$ + 1H from $CH_2$), 1.24 – 1.13 (m, 6H, 3*$CH_2$), 1.02 (s, 3H, $CH_3$), 0.98 (s, 3H, $CH_3$), 0.62 (s, 3H, $CH_3$), 0.52 (s, 3H, $CH_3$).

**$^{19}$F NMR (376 MHz, DMSO-d$_6$):** δ(ppm) = -74,6.

**ESI-MS, positive mode**: m/z = 1481.5 $[M-CF_3CO_2^-]^+$

**HRMS (ESI)** calculated for $C_{84}H_{101}N_4O_{16}SSi^+$ $[M-CF_3CO_2^-]^+$ : 1481.6697, found: 1481.6697.

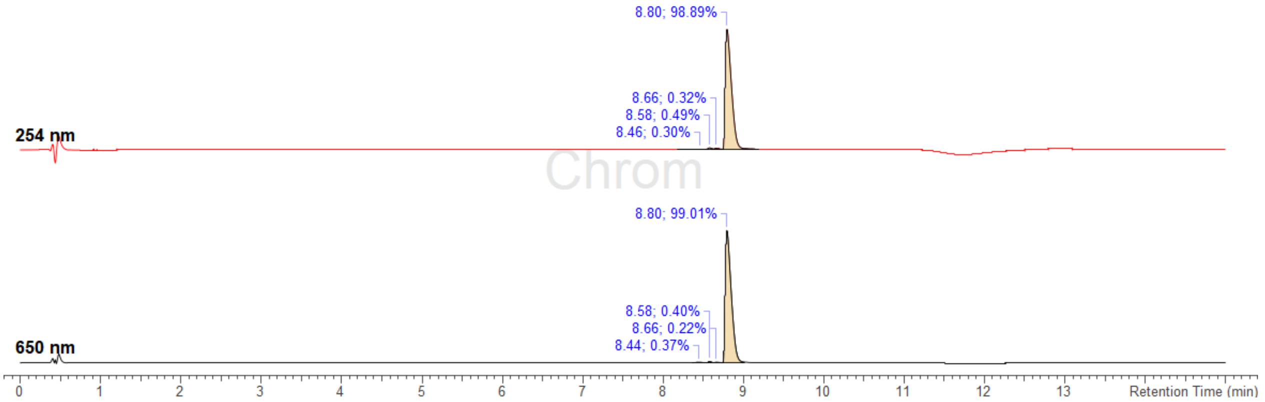

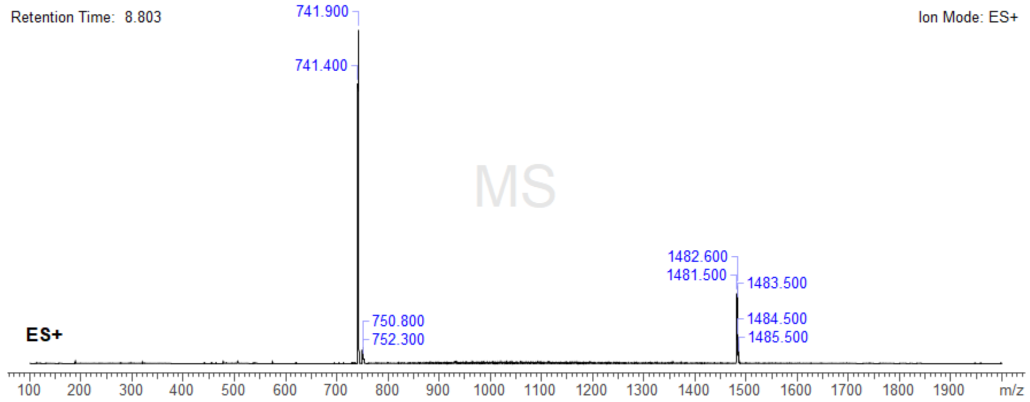

**Analysis Info**

| | | | |
|---|---|---|---|
| Analysis Name | Z:\Data\2024\2409\sam040924\MA230_30_01_127436.d | Acquisition Date | 04.09.2024 23:01:49 |
| Method | hystar_p.m | Operator | BDAL@DE |
| Sample Name | MA230 | Instrument / Ser# | micrOTOF 10237 |
| Comment | | | |

**Acquisition Parameter**

| | | | | | |
|---|---|---|---|---|---|
| Source Type | ESI | Ion Polarity | Positive | Set Nebulizer | 0.4 Bar |
| Focus | Not active | | | Set Dry Heater | 180 °C |
| Scan Begin | 50 m/z | Set Capillary | 4500 V | Set Dry Gas | 4.0 l/min |
| Scan End | 3000 m/z | Set End Plate Offset | -500 V | Set Divert Valve | Source |

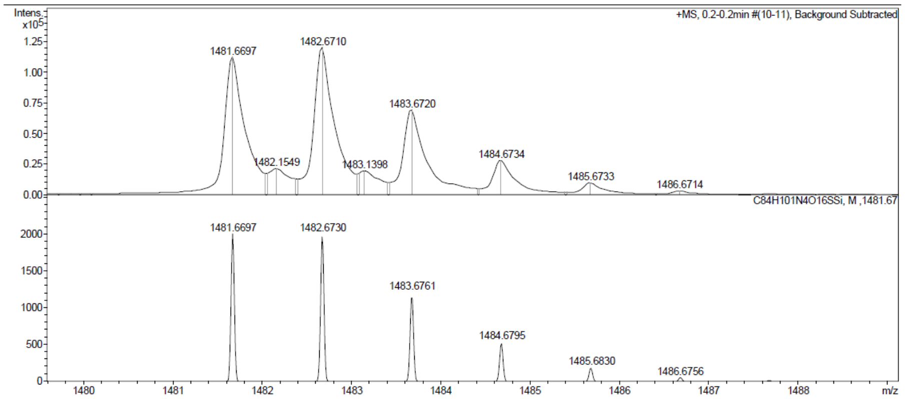

#### Probe 2 - 6-SiR-*o*-$C_9$-CTX:

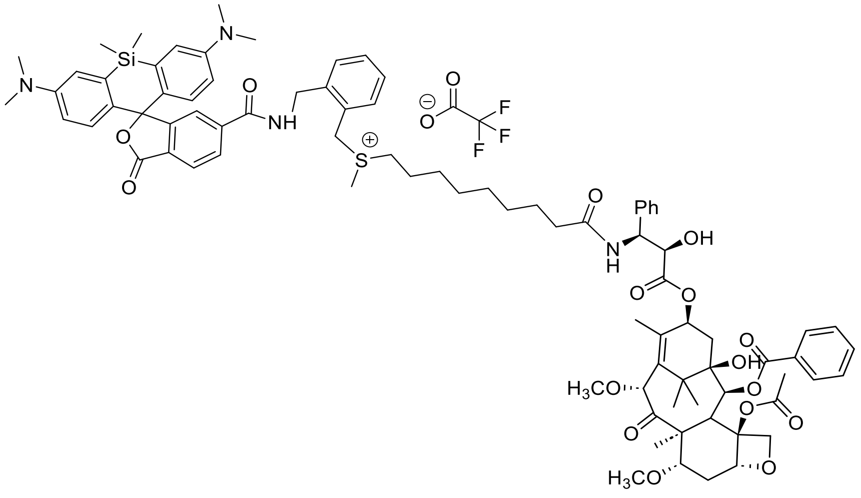

Exact Mass: 1495,69

According to the general procedure, **6-SiR-$CO_2H$** (3.0 mg, 6.35 µmol), sulfonium **15** (27.5 µL -of a 0.3M solution in DMSO-, 8.26 µmol) and **CTX-$NH_2$.$CF_3CO_2H$** (16.2 mg, 19.0 µmol) led to the expected product as a blue powder (3.24 µmol, **η = 51 %**).

**$^1$H NMR (400 MHz, DMSO-*d*$_6$):** δ = 9.34 (t, *J* = 5.5 Hz, 1H, NH), 8.36 (d, *J* = 9.0 Hz, 1H, NH), 8.14 (dd, *J* = 8.0, 1.5 Hz, 1H, $H_{Ar}$), 8.07 (d, *J* = 8.0 Hz, 1H, $H_{Ar}$), 8.00 – 7.96 (m, 2H, $H_{Ar}$), 7.72 – 7.66 (m, 2H, $H_{Ar}$), 7.62 – 7.57 (m, 2H, $H_{Ar}$), 7.48 – 7.28 (m, 8H, $H_{Ar}$), 7.24 – 7.20 (m, 1H, $H_{Ar}$), 7.04 (s, 2H, $H_{Ar}$), 6.66 -6.64 (m, 4H, $H_{Ar}$), 5.96 – 5.91 (m, 1H, CH), 5.40 (d, *J* = 7.0 Hz, 1H, CH), 5.30 (dd, *J* = 9.0, 5.5 Hz, 1H, CH), 4.96 (dd, *J* = 10.0, 2.0 Hz, 1H, CH), 4.86 (d, *J* = 13.0 Hz, 1H, 1H from $CH_2$), 4.76 (d, *J* = 13.0 Hz, 1H, 1H from $CH_2$), 4.71 (s, 1H, CH), 4.56 (d, *J* = 5.5 Hz, 2H, $CH_2$), 4.43 (d, *J* = 5.5 Hz, 1H, CH), 4.03 (s, 2H, $CH_2$), 3.76 – 3.73 (m, 1H, CH, from HSQC), 3.63 (d, *J* = 7.0 Hz, 1H, CH), 3.33 – 3.27 (m, 5H, $CH_3$ + $CH_2$), 3.21 (s, 3H, $CH_3$), 2.93 (s, 12H, 4*$CH_3$), 2.84 (s, 3H, $CH_3$), 2.71 – 2.61 (m, 1H, 1H from $CH_2$), 2.25 (s, 3H, $CH_3$), 2.19 – 2.11 (m, 2H, $CH_2$), 2.01 – 1.94 (m, 1H, 1H from $CH_2$), 1.91 – 1.81 (m, 4H, $CH_3$ +1H from $CH_2$), 1.64 – 1.42 (m, 8H, 1*$CH_3$ + 2*$CH_2$ + 1H from $CH_2$), 1.24 – 1.13 (m, 8H, 4*$CH_2$), 1.03 (s, 3H, $CH_3$), 0.98 (s, 3H, $CH_3$), 0.62 (s, 3H, $CH_3$), 0.52 (s, 3H, $CH_3$).

**$^{19}$F NMR (376 MHz, DMSO-d$_6$):** δ(ppm) = -74,6.

**ESI-MS, positive mode**: m/z = 1495.5 [M-$CF_3CO_2^-$]$^+$

**HRMS (ESI)** calculated for $C_{85}H_{103}N_4O_{16}SSi^+$ [M-$CF_3CO_2^-$]$^+$: 1495.6854, found: 1495.6874.

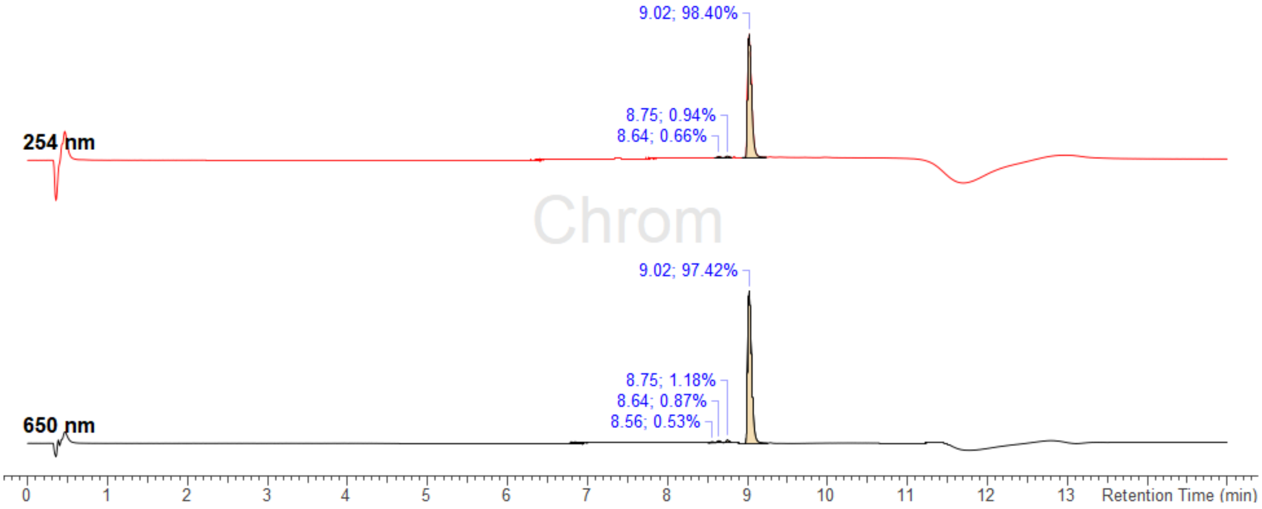

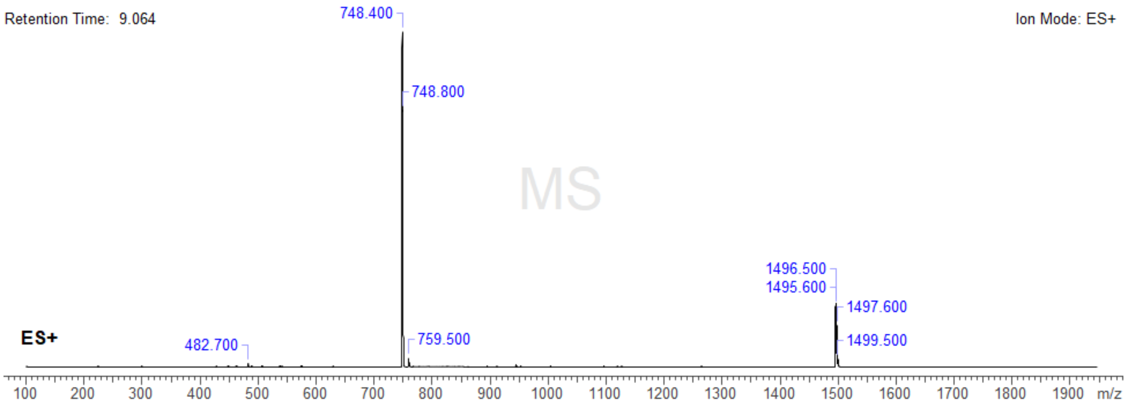

| **Analysis Info** | | | |
| --- | --- | --- | --- |
| Analysis Name | Z:\Data\2024\2409\sam040924\MA280_26_01_127432.d | Acquisition Date | 04.09.2024 22:43:15 |
| Method | hystar_p.m | Operator | BDAL@DE |
| Sample Name | MA280 | Instrument / Ser# | micrOTOF 10237 |
| Comment | | | |

| **Acquisition Parameter** | | | | | |
| --- | --- | --- | --- | --- | --- |
| Source Type | ESI | Ion Polarity | Positive | Set Nebulizer | 0.4 Bar |
| Focus | Not active | | | Set Dry Heater | 180 °C |
| Scan Begin | 50 m/z | Set Capillary | 4500 V | Set Dry Gas | 4.0 l/min |
| Scan End | 3000 m/z | Set End Plate Offset | -500 V | Set Divert Valve | Source |

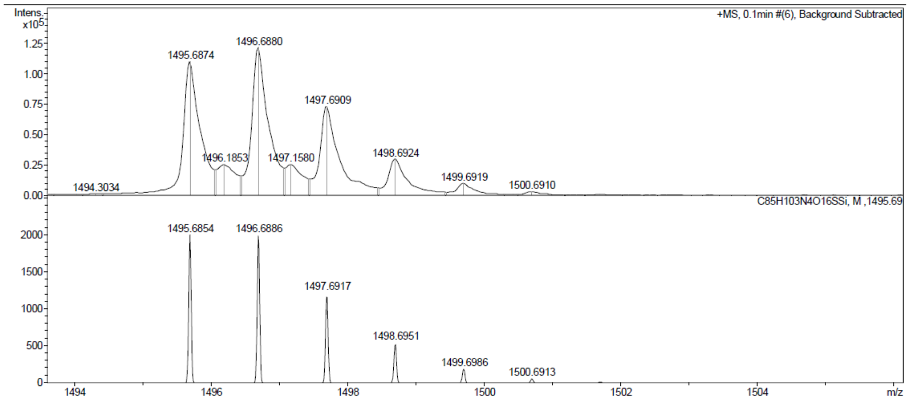

#### Probe 3 :

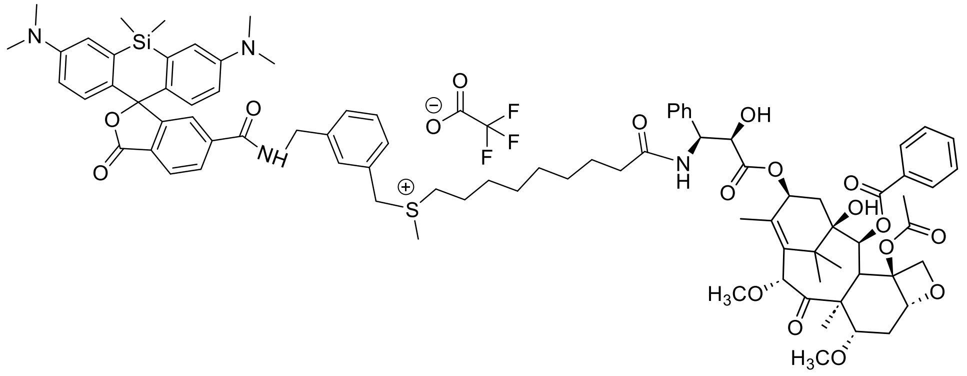

Exact Mass: 1495,69

According to the general procedure, **6-SiR-$CO_2H$** (2.0 mg, 4.23 µmol), sulfonium **16** (18.3 µL -of a 0.3M solution in DMSO-,5.50 µmol) and **CTX-$NH_2$.$CF_3CO_2H$** (10.8 mg, 12.7 µmol) led to the expected product as a blue powder (2.29 µmol, **η = 54 %**).

**$^1$H NMR (400 MHz, DMSO-$d_6$):** δ (ppm) = 9.35 (t, *J* = 6.0 Hz, 1H, NH), 8.36 (d, *J* = 9.0 Hz, 1H, NH), 8.13 (dd, *J* = 8.0, 1.5 Hz, 1H, $H_{Ar}$), 8.06 (d, *J* = 8.0 Hz, 1H, $H_{Ar}$), 8.01 – 7.95 (m, 2H, $H_{Ar}$), 7.72 – 7.65 (m, 2H, $H_{Ar}$), 7.62 – 7.56 (m, 2H, $H_{Ar}$), 7.42 – 7.31 (m, 8H, $H_{Ar}$), 7.24 – 7.20 (m, 1H, $H_{Ar}$), 7.03 (s, 2H, $H_{Ar}$), 6.66 -6.64 (m, 4H, $H_{Ar}$), 5.96 – 5.91 (m, 1H, CH), 5.40 (d, *J* = 7.0 Hz, 1H, CH), 5.30 (dd, *J* = 9.0, 5.5 Hz, 1H, CH), 4.98 – 4.95 (m, 1H, CH), 4.71 (s, 1H, CH), 4.69 – 4.57 (m, 2H, $CH_2$), 4.46 – 4.42 (m, 3H, 1*CH + 1*$CH_2$), 4.03 (s, 2H, $CH_2$), 3.76 (dd, *J* = 10.5, 6.5 Hz, 1H, CH), 3.63 (d, *J* = 7.0 Hz, 1H, CH), 3.31 (s, 3H, $CH_3$), 3.21 (s, 3H, $CH_3$), 3.20 – 3.12 (m, 2H, $CH_2$), 2.92 (s, 12H, 4*$CH_3$), 2.72 (s, 3H, $CH_3$), 2.70 – 2.60 (m, 1H, 1H from $CH_2$), 2.26 (s, 3H, $CH_3$), 2.22 – 2.08 (m, 2H, $CH_2$), 2.02 – 1.95 (m, 1H, $CH_2$, 1H from $CH_2$), 1.90 – 1.83 (m, 4H, 1*$CH_3$+1H from $CH_2$), 1.63 – 1.41 (m, 8H, 1*$CH_3$ + 2*$CH_2$ + 1H from $CH_2$), 1.25 – 1.10 (m, 8H, 4*$CH_2$), 1.03 (s, 3H, $CH_3$), 0.98 (s, 3H, $CH_3$), 0.63 (s, 3H, $CH_3$), 0.52 (s, 3H, $CH_3$).

**$^{19}$F NMR (376 MHz, DMSO-$d_6$):** δ(ppm) = -74,5.

**ESI-MS, positive mode**: m/z = 1495.6 [M-$CF_3CO_2^-$]$^+$

**HRMS (ESI)** calculated for $C_{85}H_{103}N_4O_{16}SSi^+$ [M-$CF_3CO_2^-$]$^+$ : 1495.6854, found: 1495.6861.

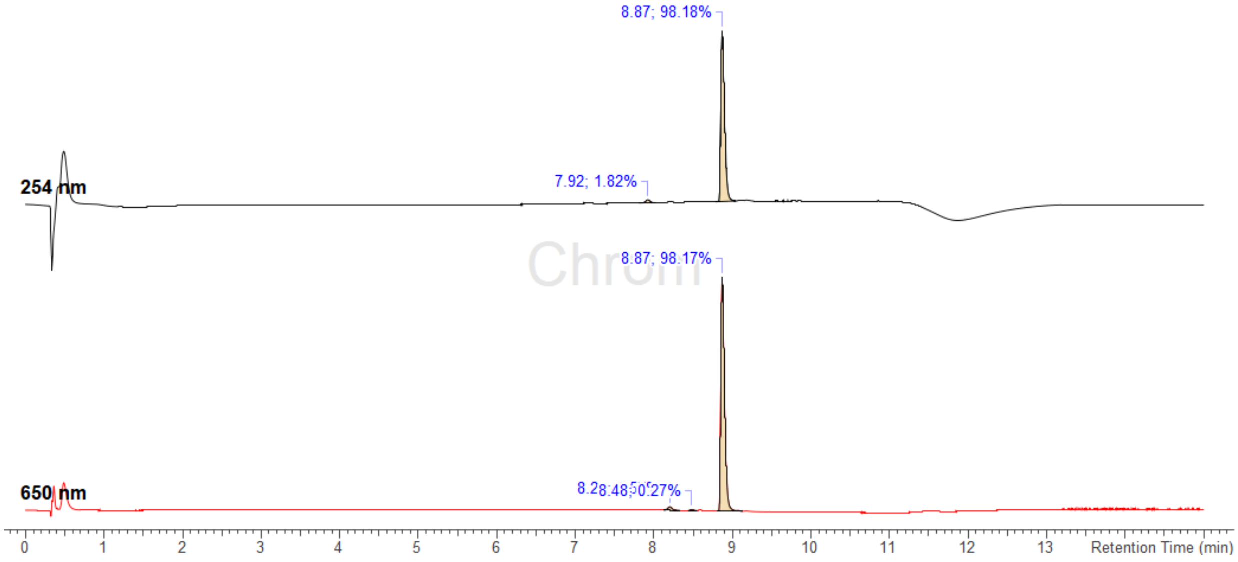

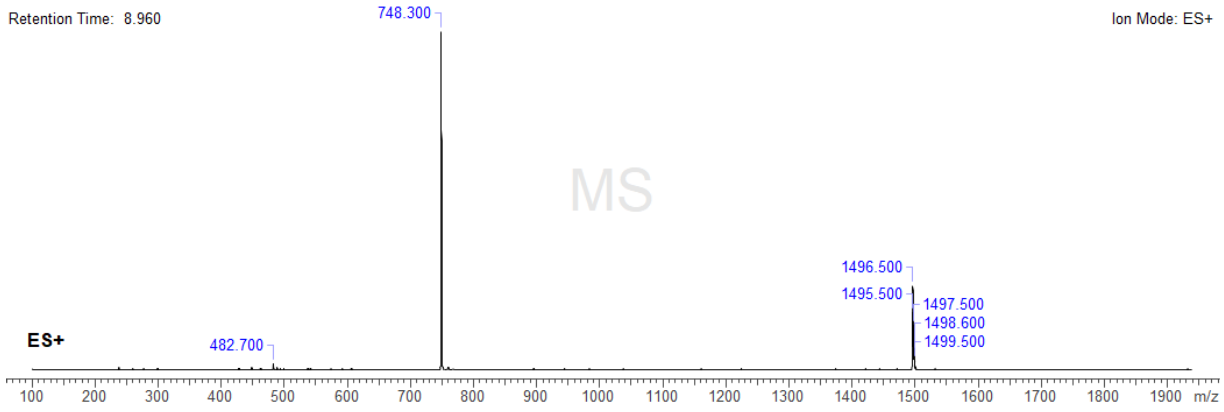

**Analysis Info**

| | | | |
|---|---|---|---|
| Analysis Name | Z:\Data\2024\2409\sam040924\MA278_31_01_127437.d | Acquisition Date | 04.09.2024 23:06:27 |
| Method | hystar_p.m | Operator | BDAL@DE |
| Sample Name | MA278 | Instrument / Ser# | micrOTOF 10237 |
| Comment | | | |

**Acquisition Parameter**

| | | | | | |
|---|---|---|---|---|---|
| Source Type | ESI | Ion Polarity | Positive | Set Nebulizer | 0.4 Bar |
| Focus | Not active | | | Set Dry Heater | 180 °C |
| Scan Begin | 50 m/z | Set Capillary | 4500 V | Set Dry Gas | 4.0 l/min |
| Scan End | 3000 m/z | Set End Plate Offset | -500 V | Set Divert Valve | Source |

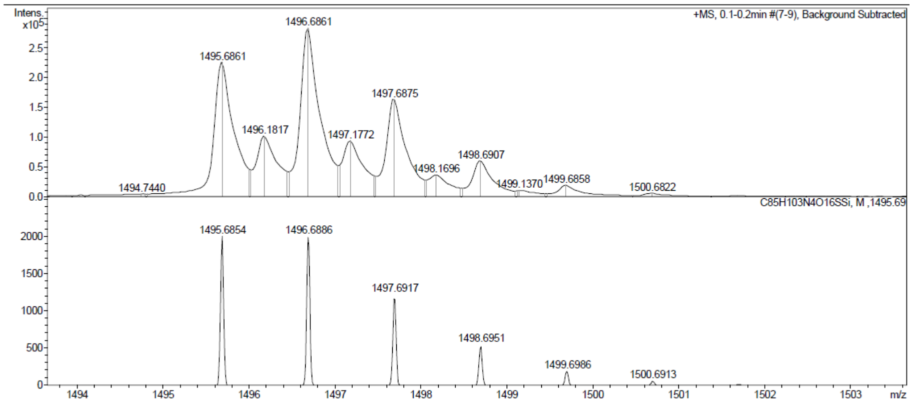

#### Probe 4 :

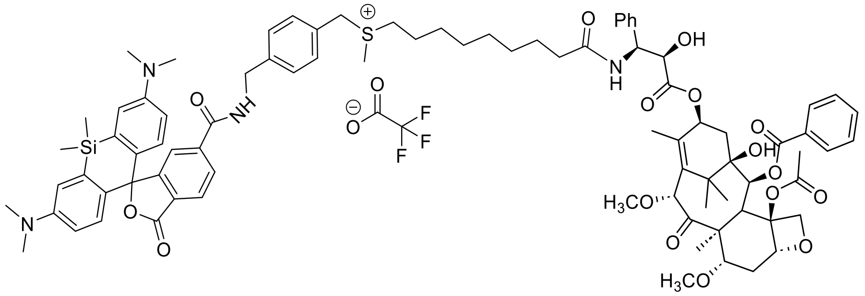

Exact Mass: 1495,68

According to the general procedure, **6-SiR-$CO_2H$** (2.0 mg, 4.23 µmol), sulfonium **17** (19.6 µL -of a 0.28M solution in DMSO-,5.50 µmol) and **CTX-$NH_2$.$CF_3CO_2H$** (10.8 mg, 12.7 µmol) led to the expected product as a blue powder (2.46 µmol, **η = 58 %**).

**$^1$H NMR (400 MHz, DMSO-$d_6$):** δ (ppm) = 9.35 (t, *J* = 6.0 Hz, 1H, NH), 8.36 (d, *J* = 9.0 Hz, 1H, NH), 8.15 – 8.11 (m, 1H, $H_{Ar}$), 8.06 (d, *J* = 8.0 Hz, 1H, $H_{Ar}$), 8.00 – 7.96 (m, 2H, $H_{Ar}$), 7.70 – 7.66 (m, 2H, $H_{Ar}$), 7.62 – 7.56 (m, 2H, $H_{Ar}$), 7.42 – 7.31 (m, 8H, $H_{Ar}$), 7.25 – 7.20 (m, 1H, $H_{Ar}$), 7.03 (s, 2H, $H_{Ar}$), 6.68 -6.63 (m, 4H, $H_{Ar}$), 5.99 – 5.89 (m, 1H, CH),5.40 (d, *J* = 7.0 Hz, 1H, CH), 5.30 (dd, *J* = 9.0, 5.5 Hz, 1H, CH), 4.96 (dd, *J* = 9.5, 2.0 Hz, 1H, CH), 4.71 (s, 1H, CH), 4.67 (d, *J* = 13.0 Hz, 1H, 1H from $CH_2$), 4.59 (d, *J* = 13.0 Hz, 1H, 1H from $CH_2$), 4.46 (d, *J* = 6.0 Hz, 2H, $CH_2$ ), 4.43 (d, *J* = 5.5 Hz, 1H, CH), 4.03 (s, 2H, $CH_2$), 3.76 (dd, *J* = 10.5, 6.5 Hz, 1H, CH), 3.63 (d, *J* = 7.0 Hz, 1H, CH), 3.31 (s, 3H, $CH_3$), 3.23 – 3.13 (m, 5H, 1*$CH_2$ + 1*$CH_3$), 2.93 (s, 12H, 4*$CH_3$), 2.73 (s, 3H, $CH_3$), 2.67 – 2.62 (m, 1H, 1H from $CH_2$), 2.26 (s, 3H, $CH_3$), 2.20 – 2.12 (m, 2H, $CH_2$), 2.00 – 1.95 (m, 1H, 1H from $CH_2$), 1.90 – 1.81 (m, 4H, 1*$CH_3$ + 1H from $CH_2$),1.64 – 1.44 (m, 8H, 1*$CH_3$ + 2*$CH_2$ + 1H from $CH_2$), 1.27 – 1.14 (m, 8H, 4*$CH_2$), 1.03 (s, 3H, $CH_3$), 0.98 (s, 3H, $CH_3$), 0.63 (s, 3H, $CH_3$), 0.52 (s, 3H, $CH_3$).

**$^{19}$F NMR (376 MHz, DMSO-$d_6$):** δ(ppm) = -74,5.

**ESI-MS, positive mode**: m/z = 1495.5 $[M-CF_3CO_2^-]^+$

**HRMS (ESI)** calculated for $C_{85}H_{103}N_4O_{16}SSi^+$ $[M-CF_3CO_2^-]^+$ : 1495.6854, found: 1495.6857.

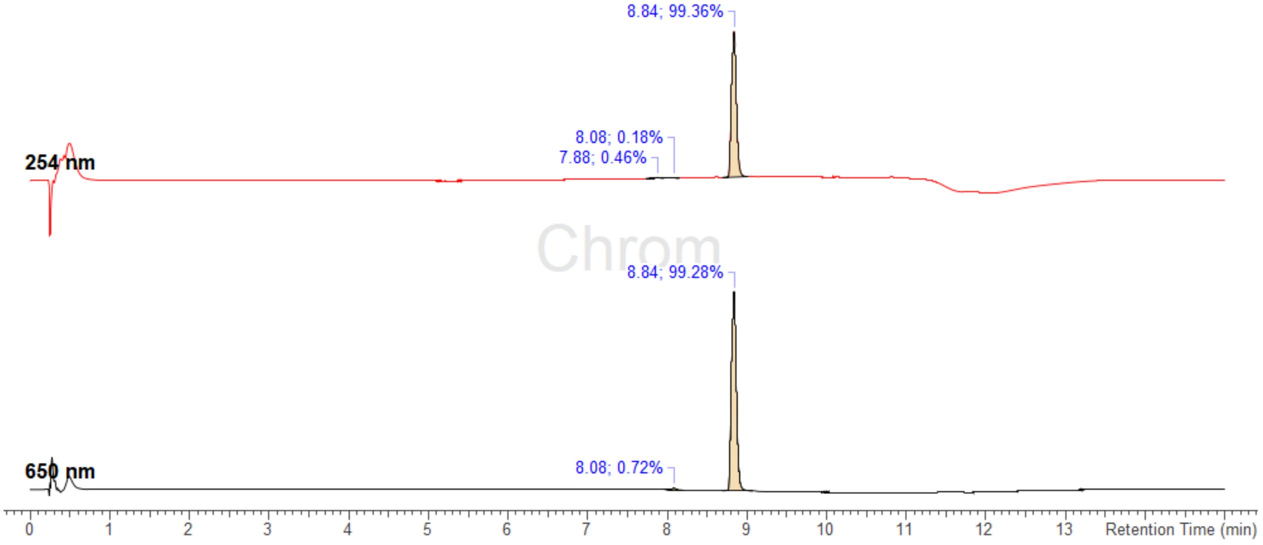

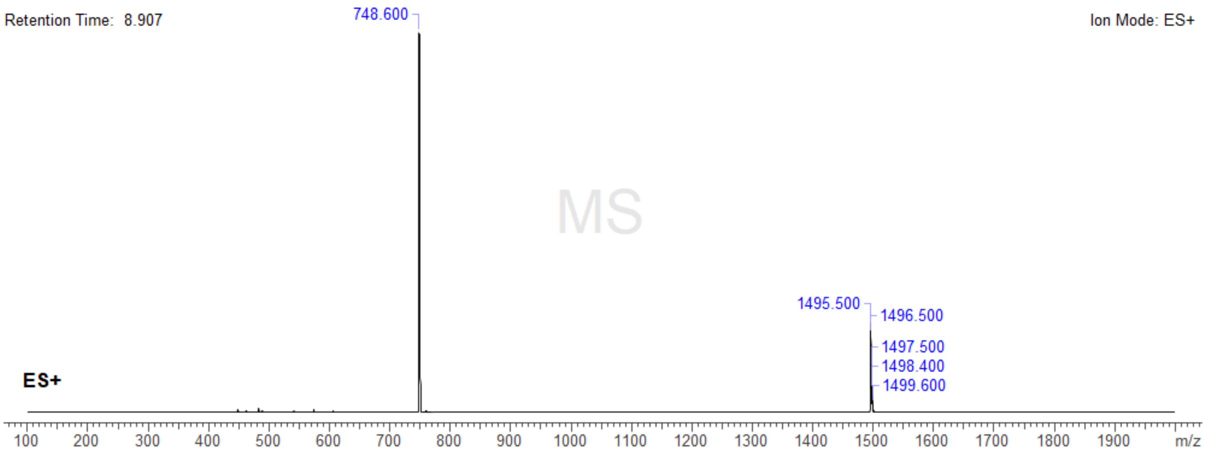

**Analysis Info**

| | | | |
|---|---|---|---|
| Analysis Name | Z:\Data\2024\2409\sam040924\MA279_29_01_127435.d | Acquisition Date | 04.09.2024 22:57:09 |
| Method | hystar_p.m | Operator | BDAL@DE |
| Sample Name | MA279 | Instrument / Ser# | micrOTOF 10237 |
| Comment | | | |

**Acquisition Parameter**

| | | | | | |
|---|---|---|---|---|---|
| Source Type | ESI | Ion Polarity | Positive | Set Nebulizer | 0.4 Bar |
| Focus | Not active | | | Set Dry Heater | 180 °C |
| Scan Begin | 50 m/z | Set Capillary | 4500 V | Set Dry Gas | 4.0 l/min |
| Scan End | 3000 m/z | Set End Plate Offset | -500 V | Set Divert Valve | Source |

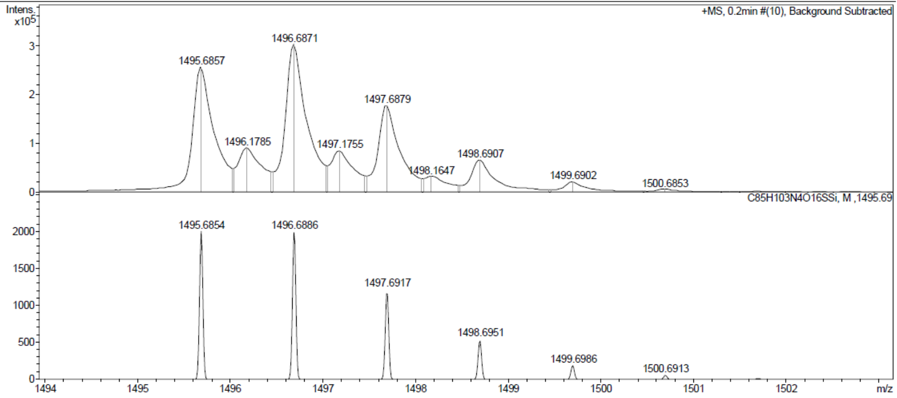

#### Probe 5 :

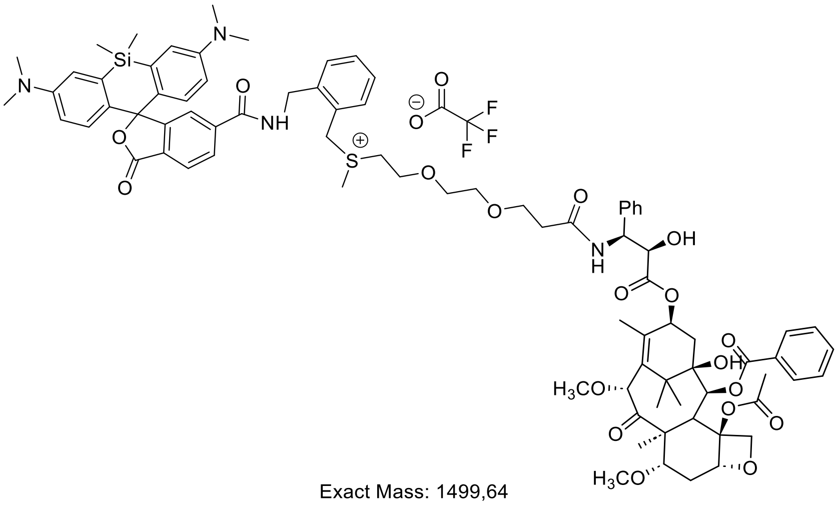

Exact Mass: 1499,64

According to the general procedure, **6-SiR-CO₂H** (2.0 mg, 4.23 µmol), sulfonium **18** (37.7 µL -of a 0.146 M solution in DMSO-,5.50 µmol) and **CTX-NH₂.CF₃CO₂H** (10.8 mg, 12.7 µmol) led to the expected product as a blue powder (2.26 µmol, **η = 53 %**).

**$^1$H NMR (400 MHz, DMSO-d$_6$):** δ (ppm) = 9.34 (t, *J* = 5.5 Hz, 1H, NH), 8.46 (d, *J* = 9.0 Hz, 1H, NH), 8.13 (dd, *J* = 8.0, 1.5 Hz, 1H, $H_{Ar}$), 8.07 (d, *J* = 8.0 Hz, 1H, $H_{Ar}$), 8.00 – 7.96 (m, 2H, $H_{Ar}$), 7.72 – 7.65 (m, 2H, $H_{Ar}$), 7.62 – 7.56 (m, 2H, $H_{Ar}$), 7.45 – 7.40 (m, 3H, $H_{Ar}$), 7.38 – 7.29 (m, 5H, $H_{Ar}$), 7.23 – 7.18 (m, 1H, $H_{Ar}$), 7.03 (s, 2H, $H_{Ar}$), 6.64 (s, 4H, $H_{Ar}$), 5.97 – 5.92 (m, 1H, CH), 5.39 (d, *J* = 7.0 Hz, 1H, CH), 5.28 (dd, *J* = 9.0, 5.5 Hz, 1H, CH), 4.97 – 4.94 (m, 1H, CH), 4.91 (d, *J* = 13.0 Hz, 1H, 1H from $CH_2$), 4.77 (d, *J* = 13.0 Hz, 1H, 1H from $CH_2$), 4.70 (s, 1H, CH), 4.55 (d, *J* = 5.5 Hz, 2H, $CH_2$), 4.43 (d, *J* = 6.0 Hz, 1H, CH), 4.05 – 4.00 (m, 2H, $CH_2$), 3.84 – 3.68 (m, 3H, $CH_2$ + CH), 3.63 (d, *J* = 7.0 Hz, 1H, CH), 3.57 – 3.41 (m, 8H, 4*$CH_2$), 3.30 (s, 3H, $CH_3$), 3.21 (s, 3H, $CH_3$), 2.92 (s, 12H, 4*$CH_3$), 2.83 (s, 3H, $CH_3$), 2.69 – 2.61 (m, 1H, 1H from $CH_2$), 2.45 – 2.38 (m, 2H, $CH_2$), 2.23 (s, 3H, $CH_3$), 1.96 (dd, *J* = 15.0, 9.0 Hz, 1H, 1H from $CH_2$), 1.85 – 1.79 (m, 4H, 1*$CH_3$ + 1H from $CH_2$), 1.54 – 1.46 (m, 4H, 1*$CH_3$ + 1H from $CH_2$), 1.02 (s, 3H, $CH_3$), 0.98 (s, 3H, $CH_3$), 0.62 (s, 3H, $CH_3$), 0.52 (s, 3H, $CH_3$).

**$^{19}$F NMR (376 MHz, DMSO-d$_6$):** δ(ppm) = -74.5.

**ESI-MS, positive mode**: m/z = 1499.6 [M-$CF_3CO_2^-$]$^+$

**HRMS (ESI)** calculated for $C_{83}H_{99}N_4O_{18}SSi^+$ [M-$CF_3CO_2^-$]$^+$ : 1499.6439, found: 1499.6442.

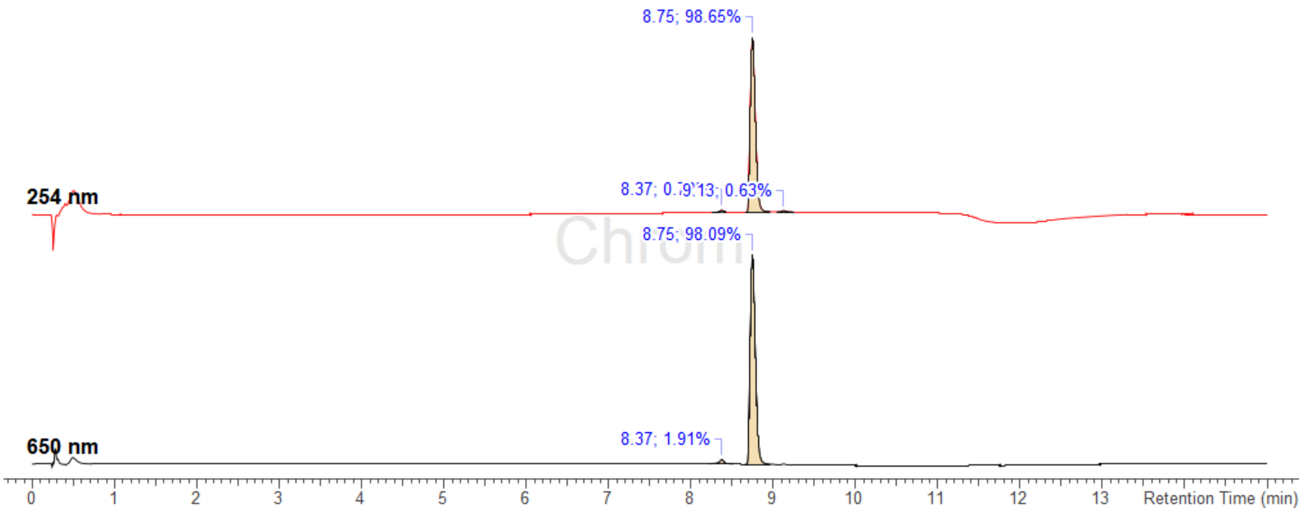

Retention Time: 8.829 Ion Mode: ES+

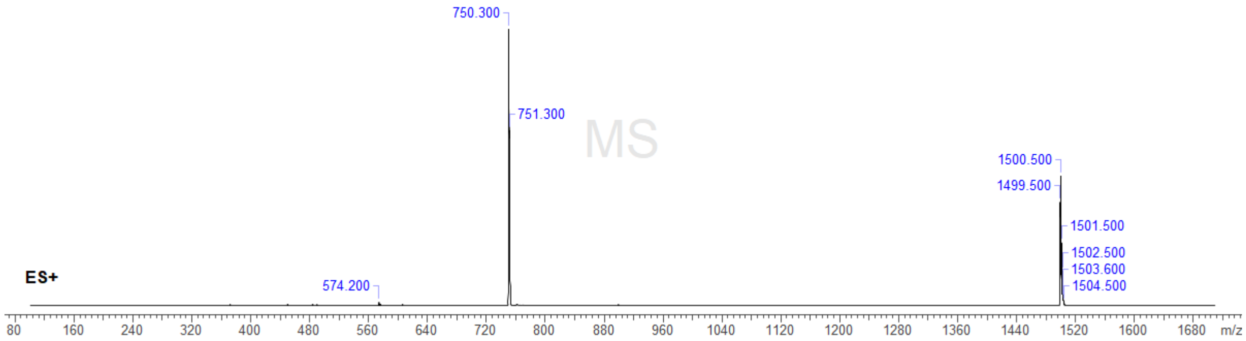

| | | | |
|---|---|---|---|
| **Analysis Info** | | Acquisition Date | 04.09.2024 22:47:54 |
| Analysis Name | Z:\Data\2024\2409\sam040924\MA244_27_01_127433.d | | |
| Method | hystar_p.m | Operator | BDAL@DE |
| Sample Name | MA244 | Instrument / Ser# | microTOF 10237 |
| Comment | | | |

| **Acquisition Parameter** | | | | | |
|---|---|---|---|---|---|
| Source Type | ESI | Ion Polarity | Positive | Set Nebulizer | 0.4 Bar |
| Focus | Not active | | | Set Dry Heater | 180 °C |
| Scan Begin | 50 m/z | Set Capillary | 4500 V | Set Dry Gas | 4.0 l/min |
| Scan End | 3000 m/z | Set End Plate Offset | -500 V | Set Divert Valve | Source |

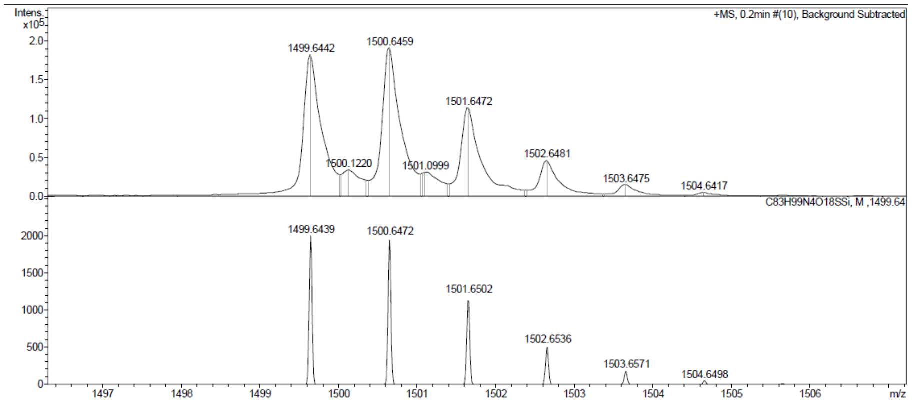

#### **Probe 6 :**

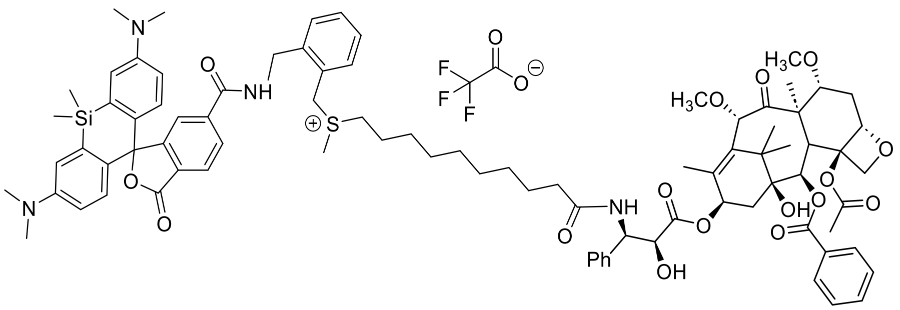

Exact Mass: 1509,70

According to the general procedure, **6-SiR-$CO_2$H** (2.0 mg, 4.23 µmol), sulfonium **18** (71.4 µL -of a 77 mM solution in DMSO- ,5.50 µmol) and **CTX-$NH_2$.$CF_3CO_2H$** (10.8 mg, 12.7 µmol) led to the expected product as a blue powder (1.48 µmol, **η = 35 %**).

**$^1$H NMR (400 MHz, DMSO-d$_6$):** δ (ppm) = 9.33 (t, $J$ = 5.5 Hz, 1H, NH), 8.35 (d, $J$ = 9.0 Hz, 1H, NH), 8.13 (dd, $J$ = 8.0, 1.5 Hz, 1H, , $H_{Ar}$), 8.06 (d, $J$ = 8.0 Hz, 1H, $H_{Ar}$), 8.00 – 7.96 (m, 2H, , $H_{Ar}$), 7.72 – 7.66 (m, 2H, $H_{Ar}$), 7.62 – 7.57 (m, 2H,$H_{Ar}$), 7.46 – 7.40 (m, 3H, $H_{Ar}$), 7.39 – 7.31 (m, 5H, $H_{Ar}$), 7.24 – 7.20 (m, 1H, $H_{Ar}$), 7.02 (s, 2H, $H_{Ar}$), 6.63 (s, 4H, $H_{Ar}$), 5.96 -5.91 (m, 1H, CH), 5.40 (d, $J$ = 7.0 Hz, 1H, CH), 5.32 – 5.28 (m, 1H, CH), 4.98 – 4.94 (m, 1H, CH), 4.86 (d, $J$ = 13.0 Hz, 1H, 1H from $CH_2$), 4.76 (d, $J$ = 13.0 Hz, 1H, 1H from $CH_2$), 4.71 (s, 1H, CH), 4.64 (s, 1H, OH), 4.57 – 4.54 (m, 2H, $CH_2$), 4.43 (d, $J$ = 5.5 Hz, 1H, CH), 4.03 (s, 2H, $CH_2$), 3.76 (dd, $J$ = 10.5, 6.5 Hz, 1H, CH), 3.63 (d, $J$ = 7.0 Hz, 1H, CH), 3.33 – 3.27 (m, 5H, 1*$CH_2$ + 1*$CH_3$), 3.21 (s, 3H, $CH_3$), 2.92 (s, 12H, 4*$CH_3$), 2.84 (s, 3H, $CH_3$), 2.69 – 2.64 (m, 1H, 1H from $CH_2$), 2.26 (s, 3H, $CH_3$), 2.19 – 2.11 (m, 2H, $CH_2$), 2.01 – 1.95 (m, 1H, 1H from $CH_2$), 1.92 – 1.82 (m, 4H, 1*$CH_3$ + 1H from $CH_2$), 1.64 – 1.40 (m, 8H, 1*$CH_3$ + 2*$CH_2$ + 1H from $CH_2$), 1.23 – 1.13 (m, 10H, 5*$CH_2$), 1.03 (s, 3H, $CH_3$), 0.98 (s, 3H, $CH_3$), 0.62 (s, 3H, $CH_3$), 0.52 (s, 3H, $CH_3$).

**$^{19}$F NMR (376 MHz, DMSO-d$_6$):** δ(ppm) = -74,3.

**ESI-MS, positive mode**: m/z = 1509.5 [M-$CF_3CO_2^-$]$^+$

**HRMS (ESI)** calculated for $C_{86}H_{105}N_4O_{16}SSi^+$ [M-$CF_3CO_2^-$]$^+$ : 1509.7010, found: 1509.7014.

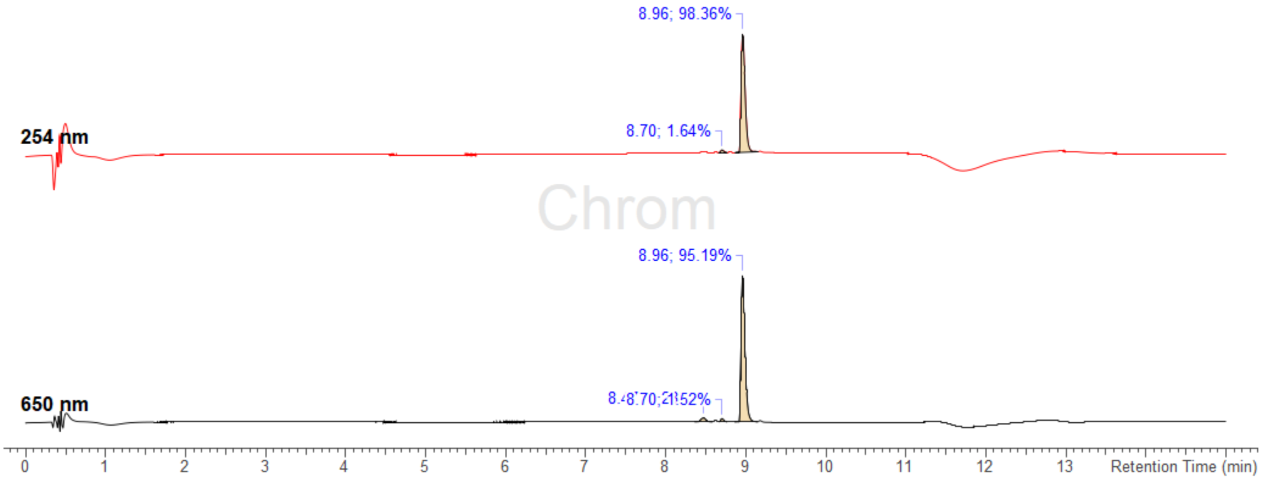

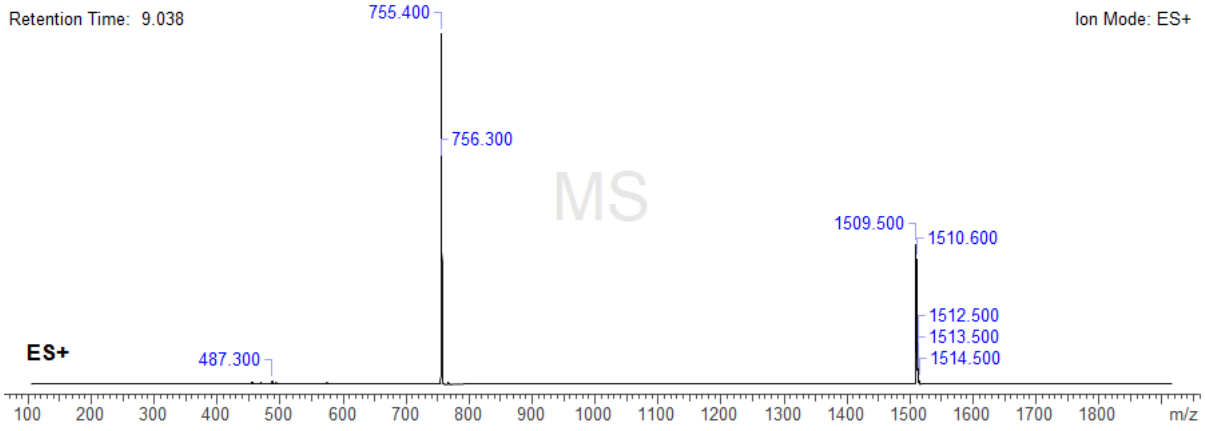

**Analysis Info**

| | | | |
|---|---|---|---|
| Analysis Name | Z:\Data\2024\2409\sam040924\MA282_28_01_127434.d | Acquisition Date | 04.09.2024 22:52:31 |
| Method | hystar_p.m | Operator | BDAL@DE |
| Sample Name | MA282 | Instrument / Ser# | micrOTOF 10237 |
| Comment | | | |

**Acquisition Parameter**

| | | | | | |
|---|---|---|---|---|---|
| Source Type | ESI | Ion Polarity | Positive | Set Nebulizer | 0.4 Bar |
| Focus | Not active | | | Set Dry Heater | 180 °C |
| Scan Begin | 50 m/z | Set Capillary | 4500 V | Set Dry Gas | 4.0 l/min |
| Scan End | 3000 m/z | Set End Plate Offset | -500 V | Set Divert Valve | Source |

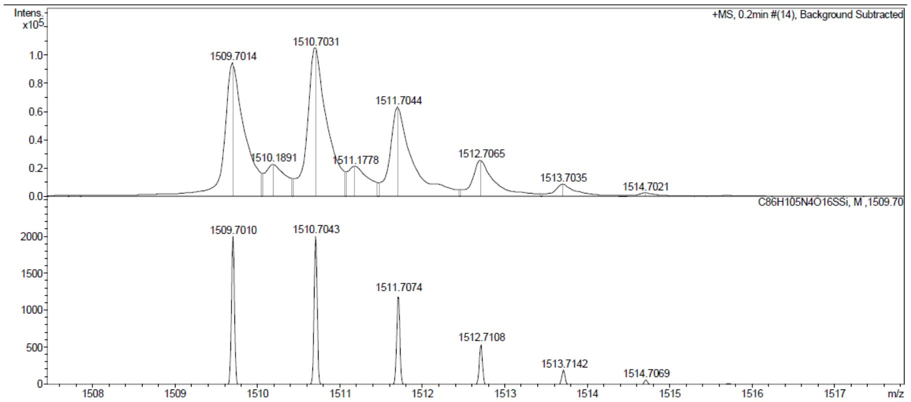

### NMR spectra

$^{1}$H NMR of ***p*-7:**

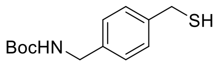

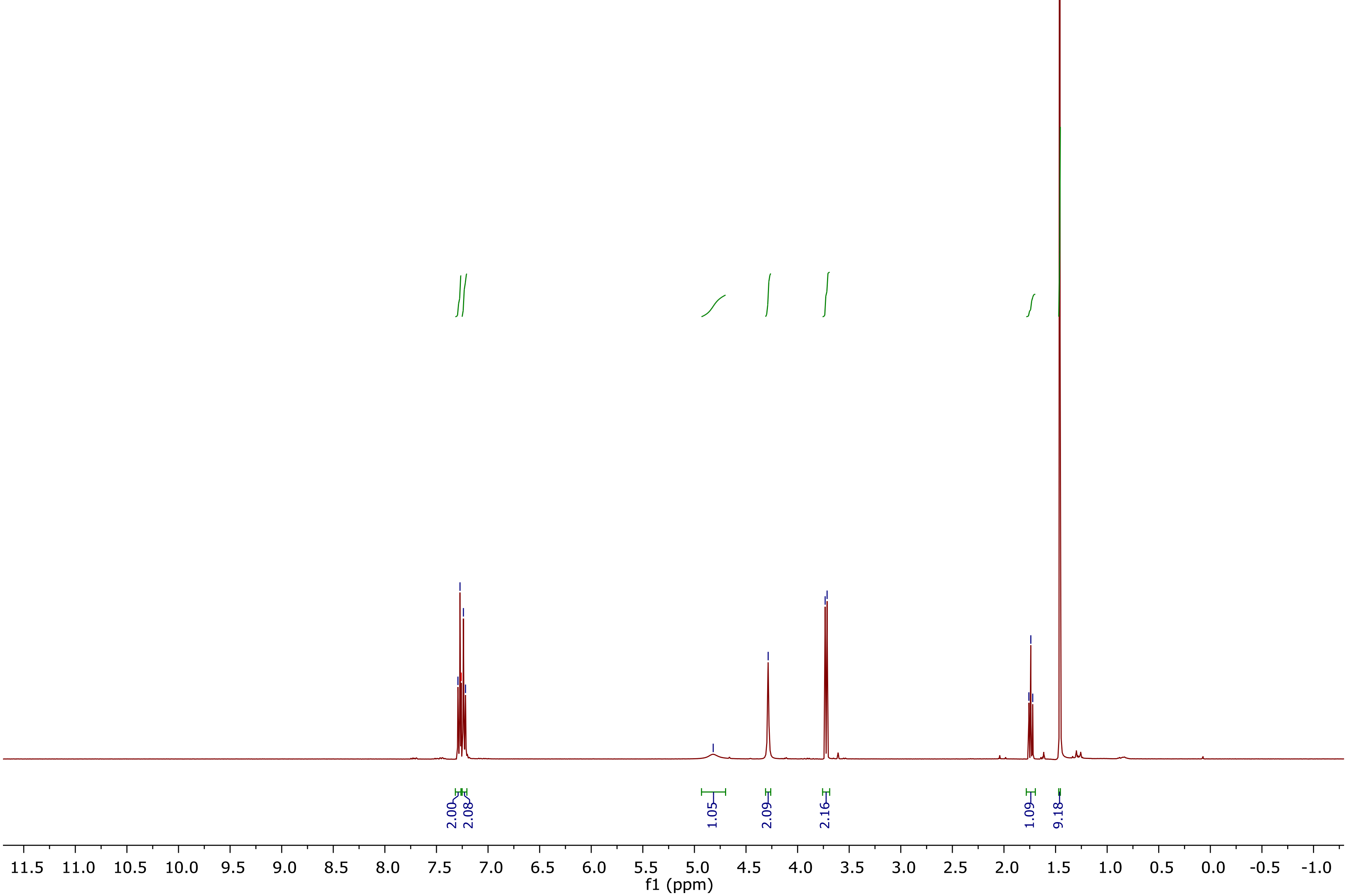

$^{13}$C NMR of ***p*-7**:

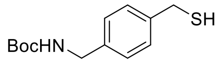

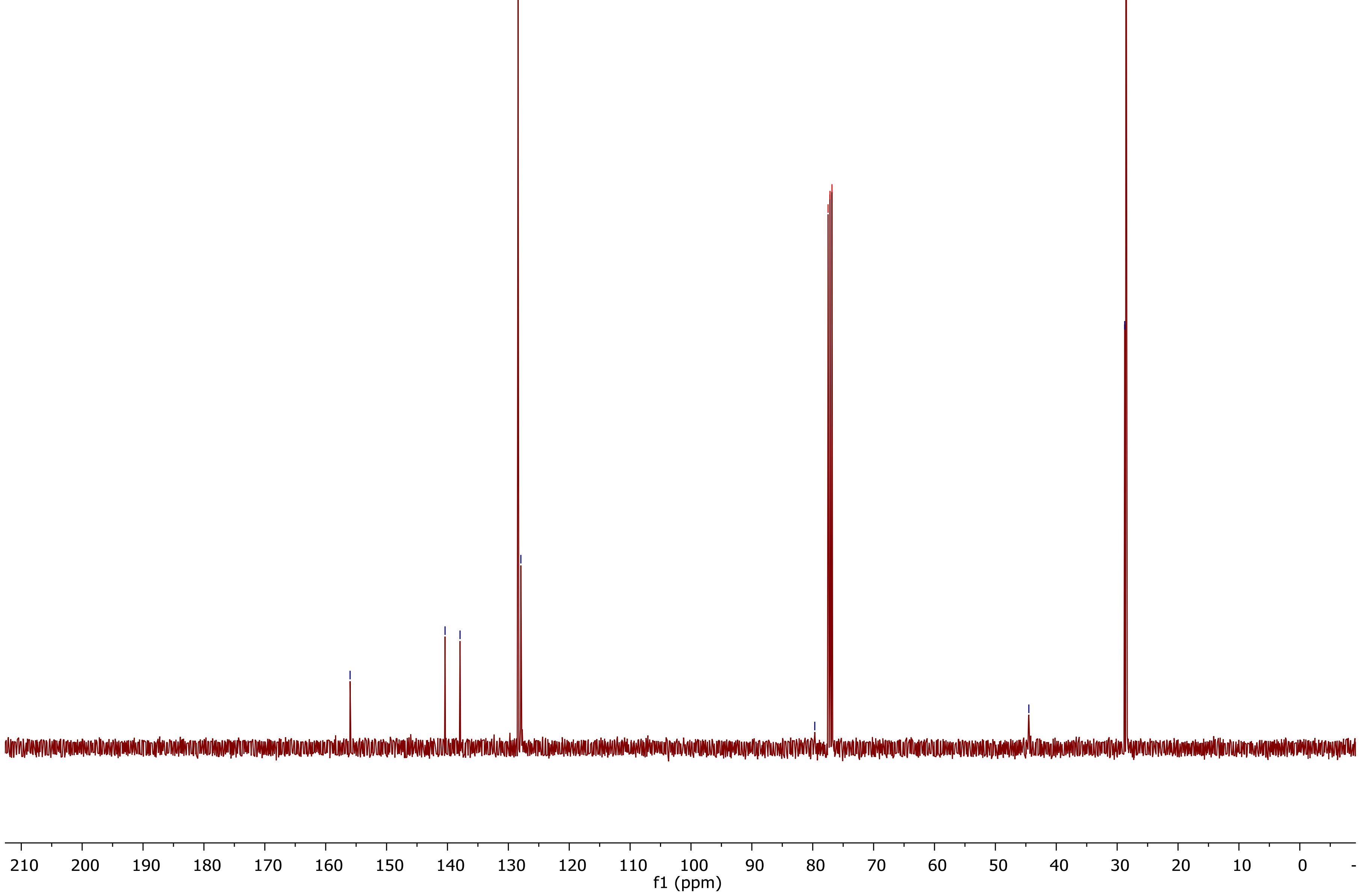

156.0
140.4
137.9
128.4
127.9
79.7
77.5 cdcl3
77.2 cdcl3
76.8 cdcl3
44.5
28.8
28.5
210
200
190
180
170
160
150
140
130
120
110
100
90
80
70
60
50
40
30
20
10
0
f1 (ppm)

[1]H NMR of ***m*-7**:

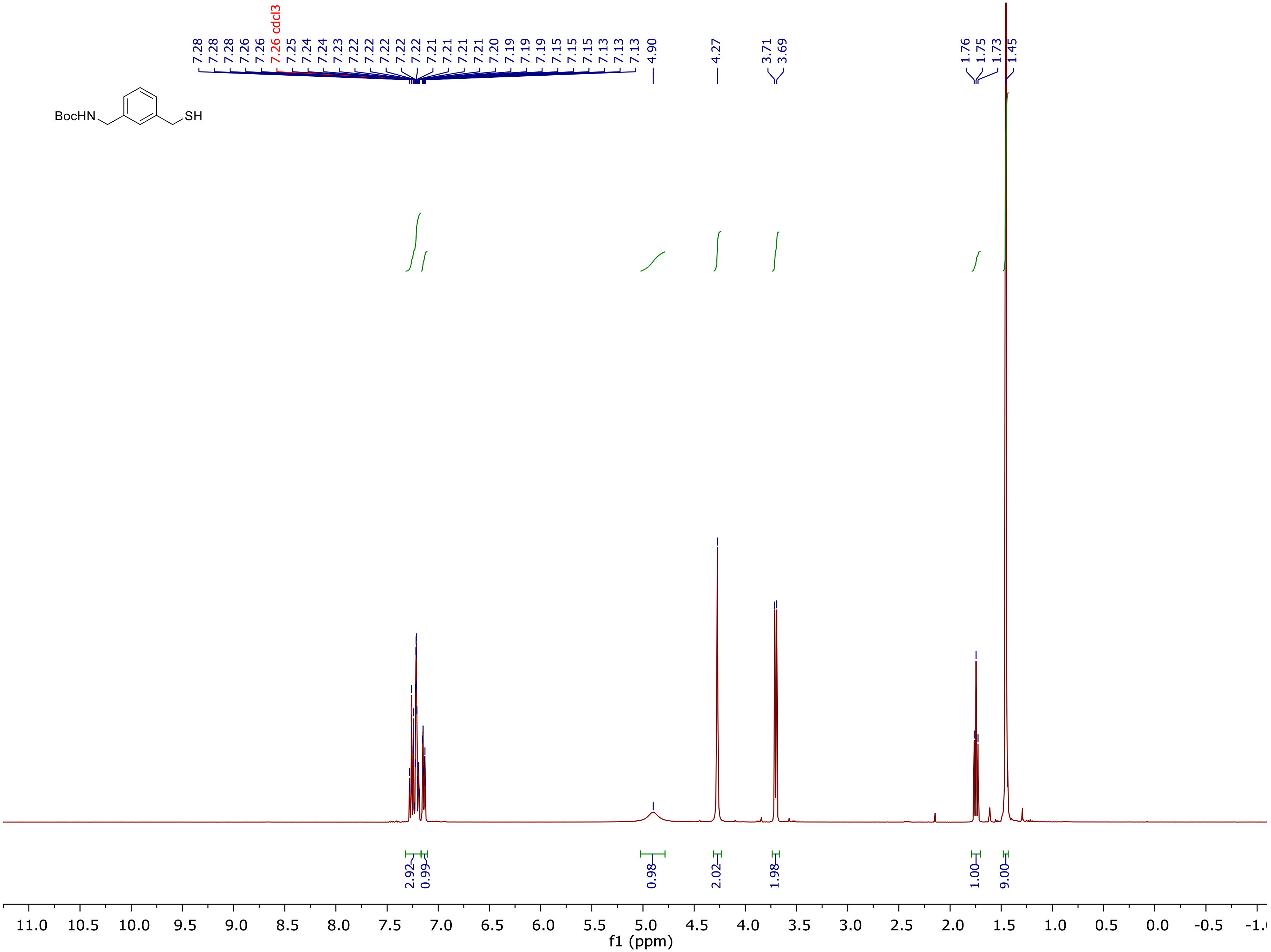

7.28
7.28
7.28
7.26
7.26
7.26 cdcl3
7.25
7.24
7.24
7.23
7.22
7.22
7.22
7.22
7.22
7.21
7.21
7.21
7.20
7.19
7.19
7.19
7.15
7.15
7.15
7.13
7.13
7.13
4.90
4.27
3.71
3.69
1.76
1.75
1.73
1.45
BocHN
SH
2.92
0.99
0.98
2.02
1.98
1.00
9.00
11.0
10.5
10.0
9.5
9.0
8.5
8.0
7.5
7.0
6.5
6.0
5.5
5.0
4.5
4.0
3.5
3.0
2.5
2.0
1.5
1.0
0.5
0.0
-0.5
f1 (ppm)

[13]C NMR of ***m*-7**:

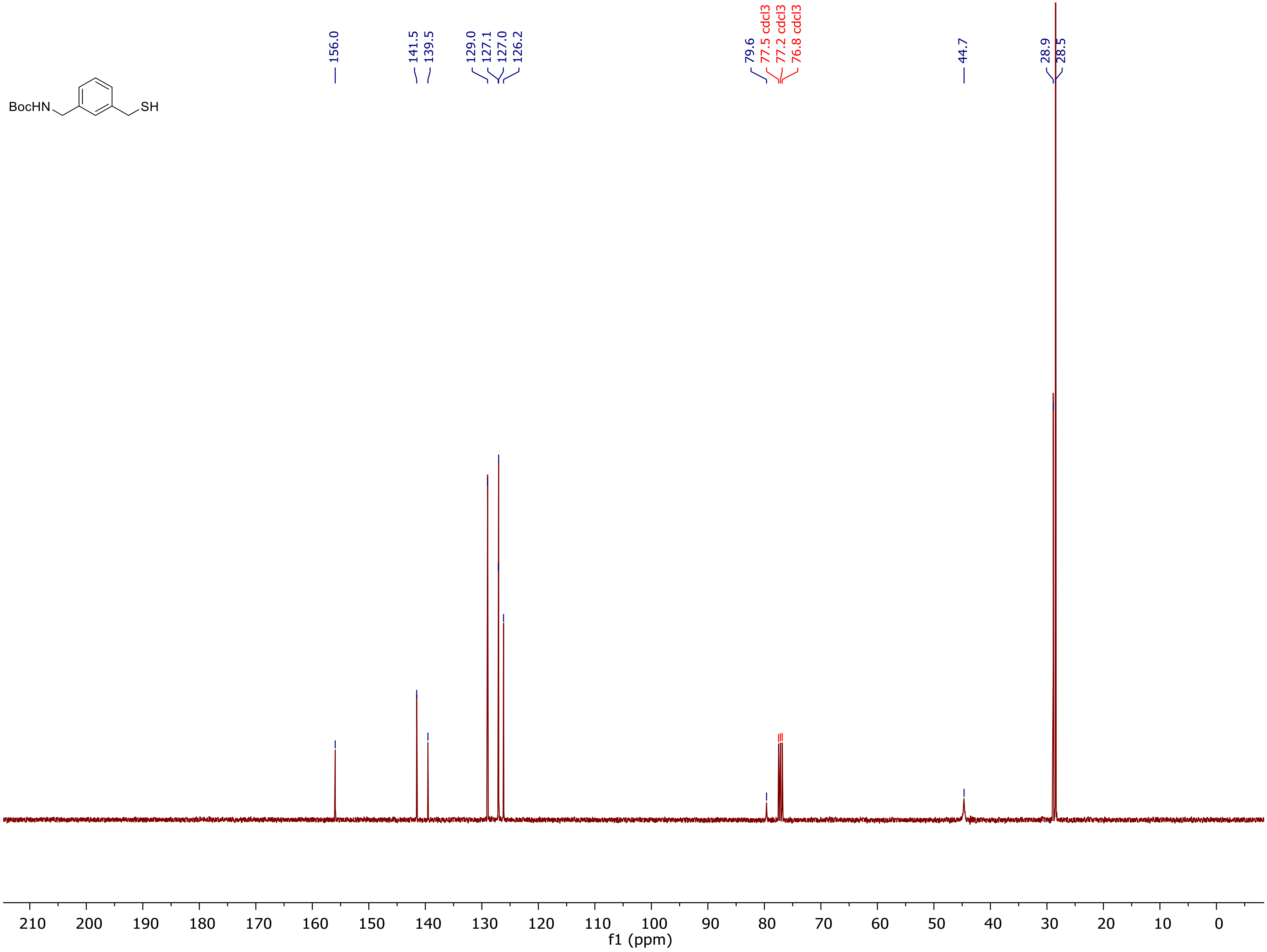

BocHN
SH
156.0
141.5
139.5
129.0
127.1
127.0
126.2
79.6
77.5 cdcl3
77.2 cdcl3
76.8 cdcl3
44.7
28.9
28.5
210
200
190
180
170
160
150
140
130
120
110
100
90
80
70
60
50
40
30
20
10
0
f1 (ppm)

$^1$H NMR of ***o*-7**:

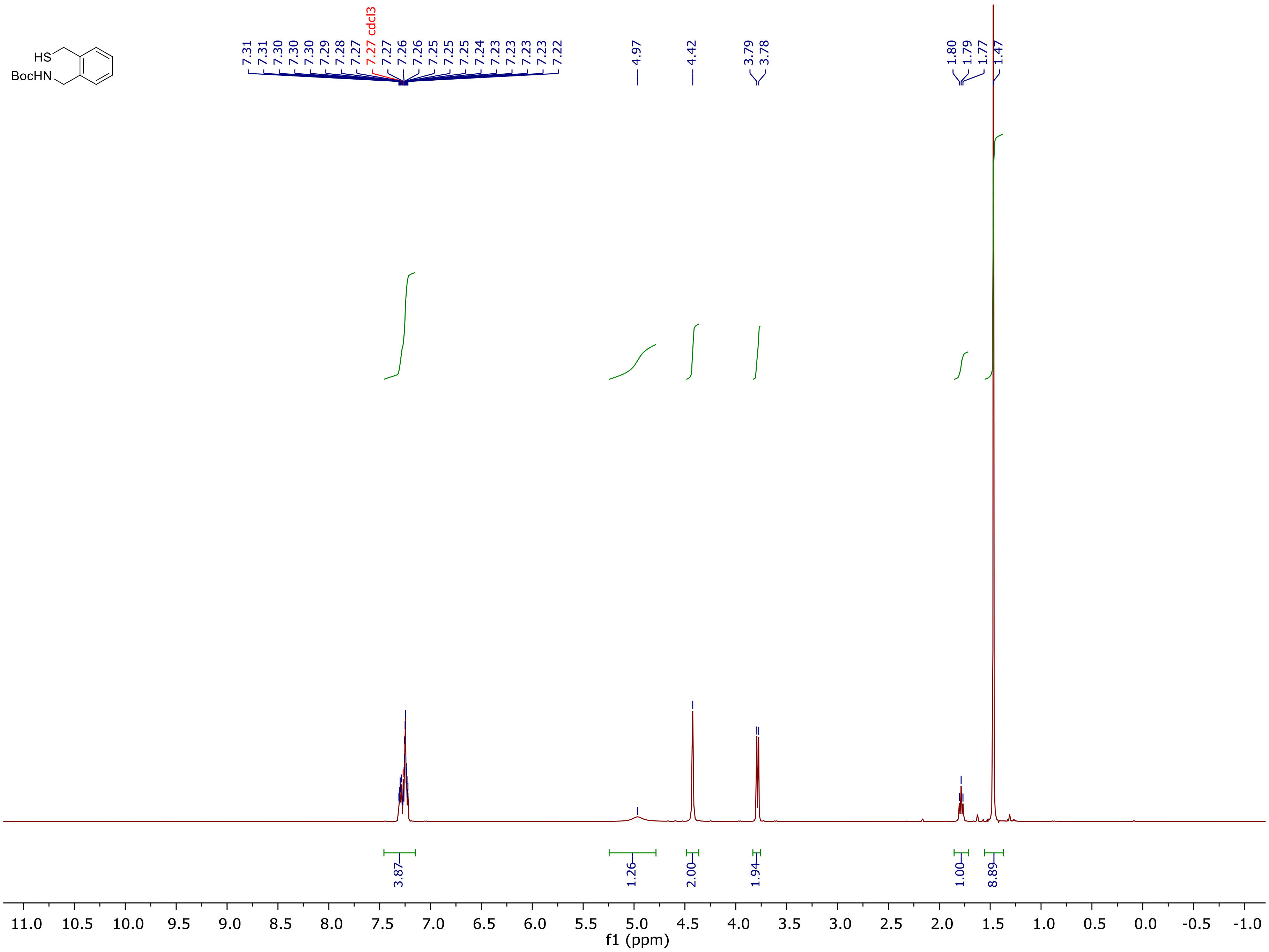
HS
BocHN
7.31
7.31
7.30
7.30
7.29
7.28
7.27
7.27 cdcl3
7.27
7.26
7.26
7.25
7.25
7.25
7.24
7.23
7.23
7.23
7.23
7.22
4.97
4.42
3.79
3.78
1.80
1.79
1.77
1.47
3.87
1.26
2.00
1.94
1.00
8.89
11.0
10.5
10.0
9.5
9.0
8.5
8.0
7.5
7.0
6.5
6.0
5.5
5.0
4.5
4.0
3.5
3.0
2.5
2.0
1.5
1.0
0.5
0.0
-0.5
-1.0
f1 (ppm)

$^{13}$C NMR of ***o*-7:**

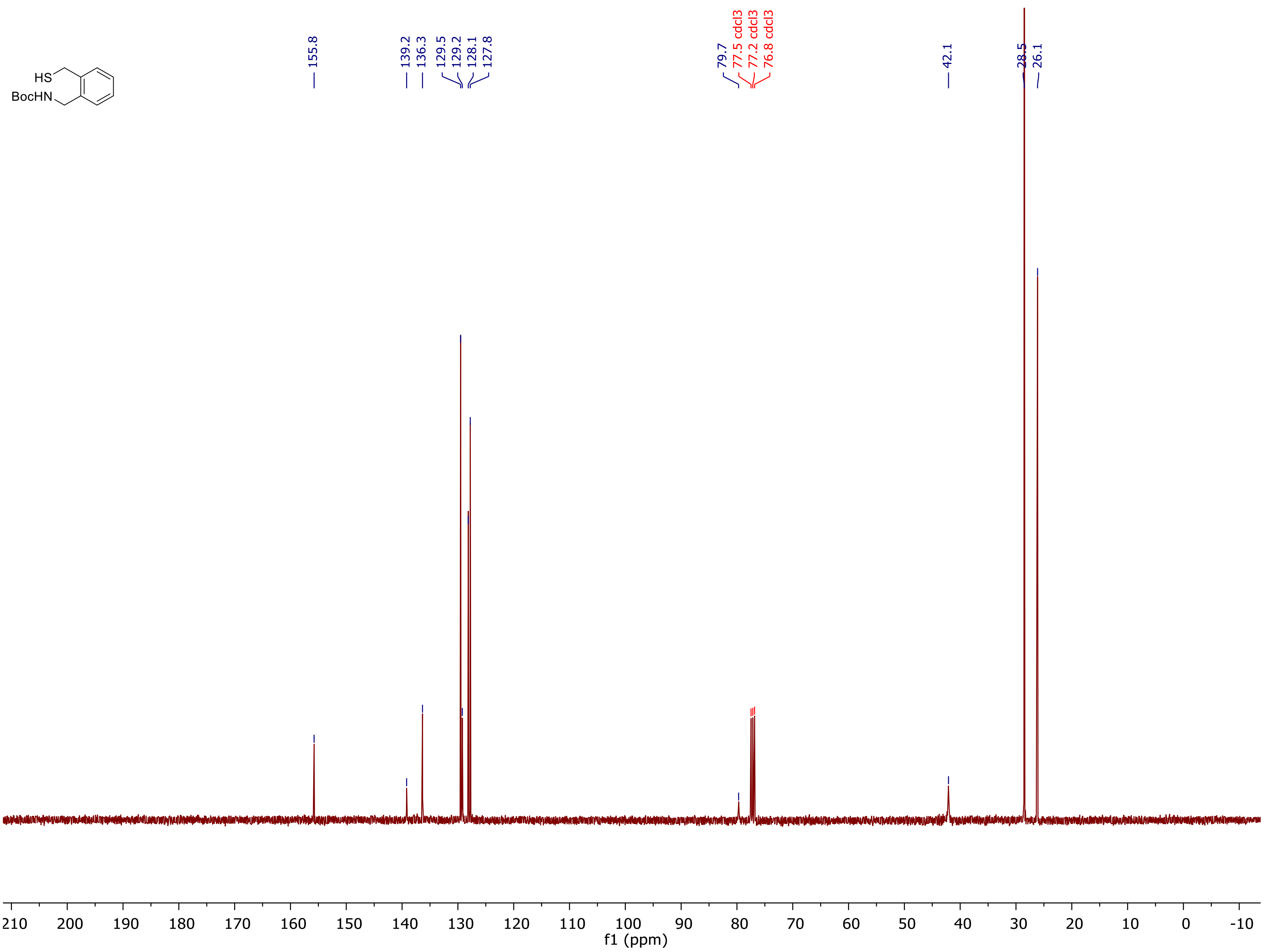
HS
BocHN
155.8
139.2
136.3
129.5
129.2
128.1
127.8
79.7
77.5 cdcl3
77.2 cdcl3
76.8 cdcl3
42.1
28.5
26.1
210
200
190
180
170
160
150
140
130
120
110
100
90
80
70
60
50
40
30
20
10
0
-10
f1 (ppm)

[1]H NMR of **8:**

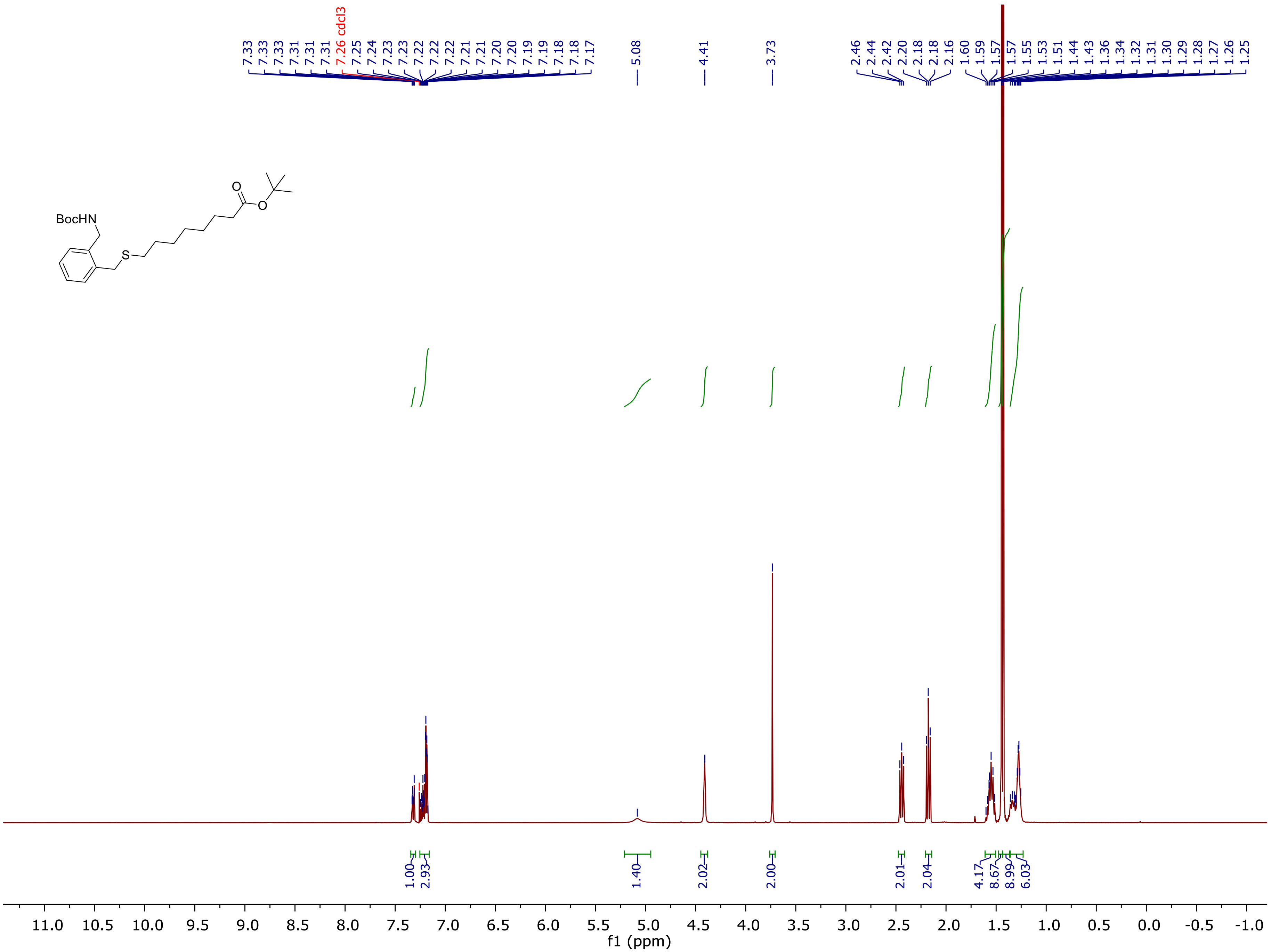

[13]C NMR of **8:**

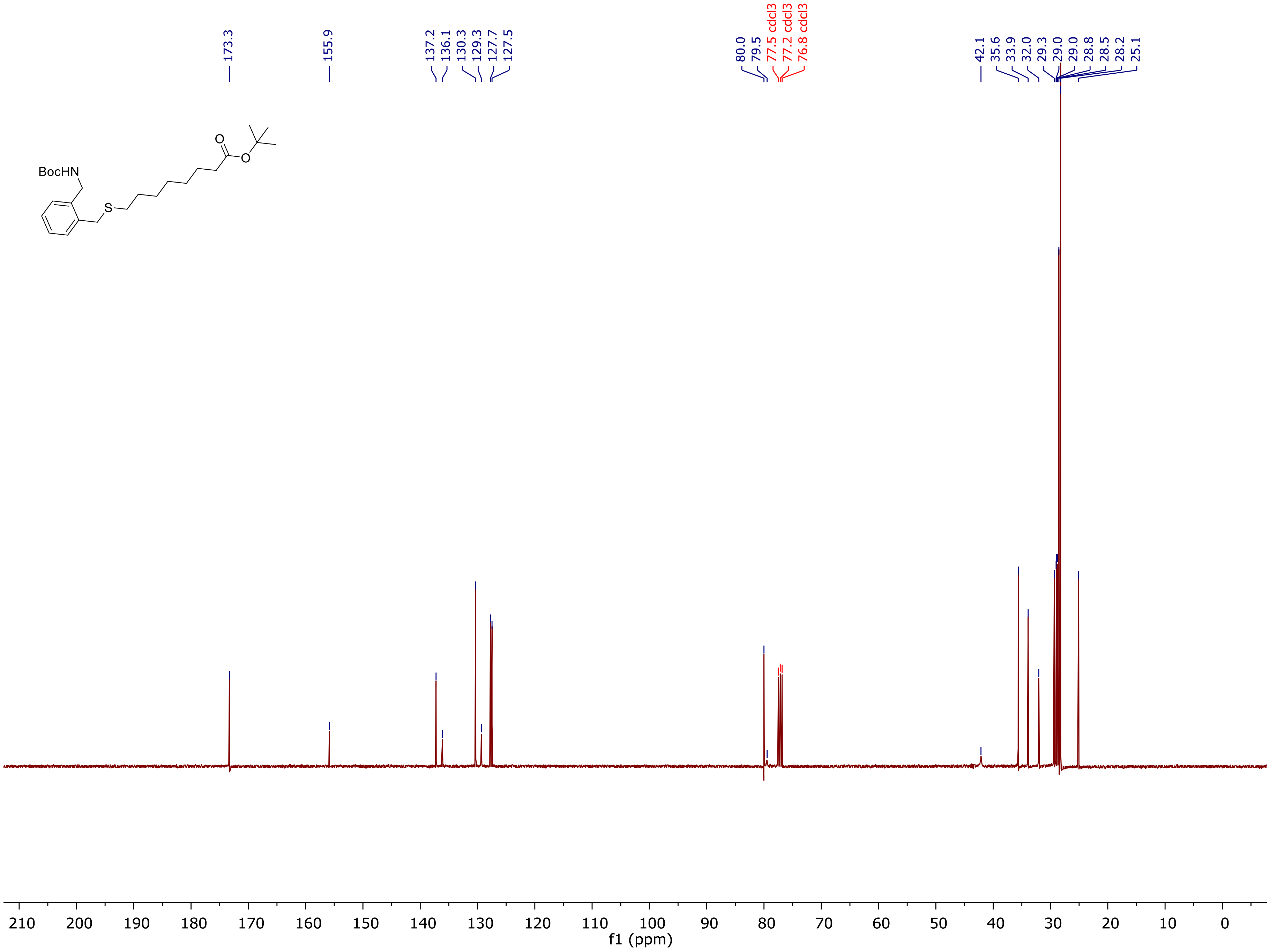
173.3
155.9
137.2
136.1
130.3
129.3
127.7
127.5
80.0
79.5
77.5 cdcl3
77.2 cdcl3
76.8 cdcl3
42.1
35.6
33.9
32.0
29.3
29.0
29.0
28.8
28.5
28.2
25.1
BocHN
S
O
O
210 200 190 180 170 160 150 140 130 120 110 100 90 80 70 60 50 40 30 20 10 0
f1 (ppm)

[1]H NMR of **9:**

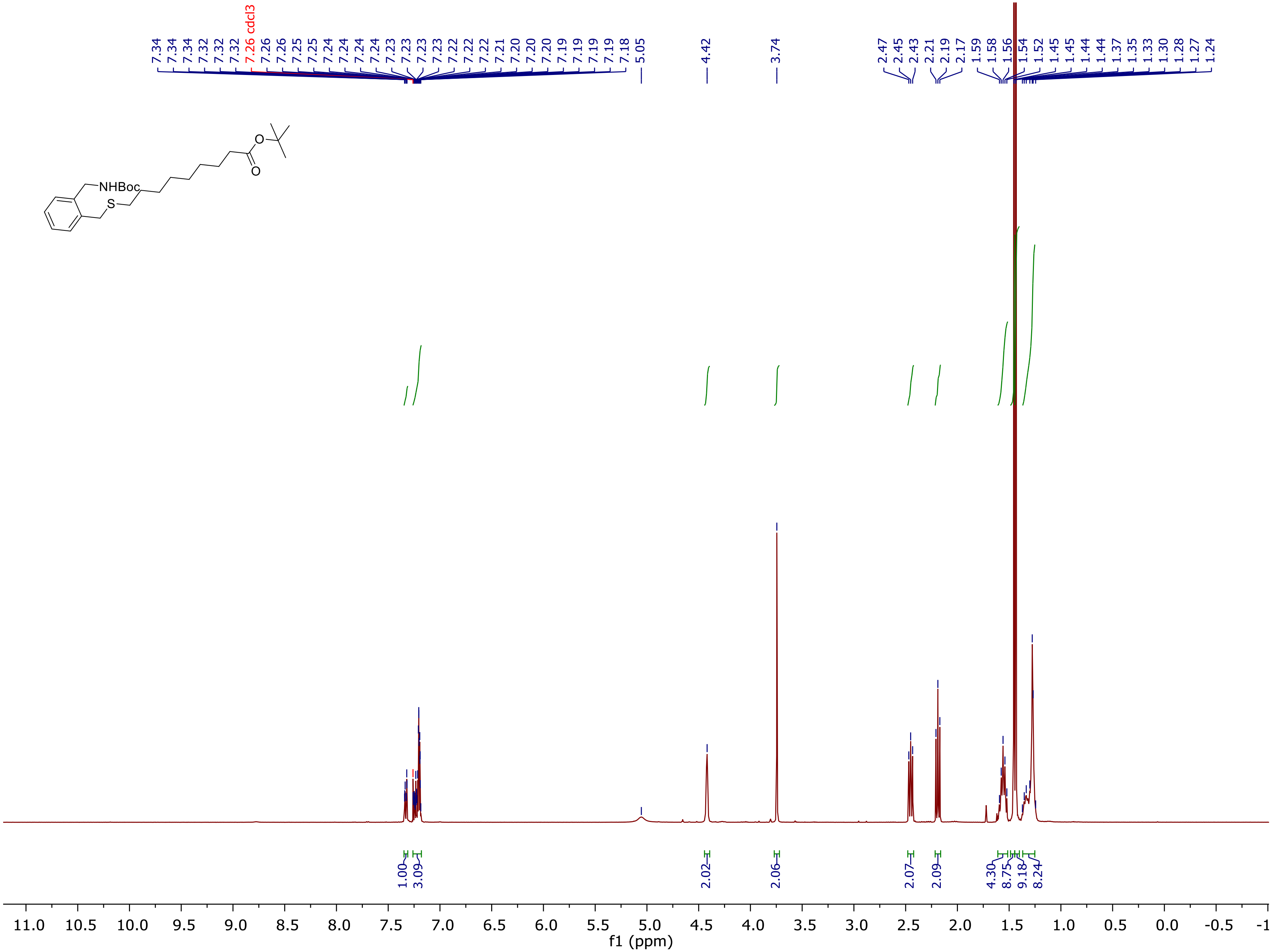

7.34
7.34
7.34
7.32
7.32
7.32
7.26 cdcl3
7.26
7.26
7.25
7.25
7.24
7.24
7.24
7.24
7.23
7.23
7.23
7.23
7.22
7.22
7.22
7.21
7.20
7.20
7.20
7.19
7.19
7.19
7.19
7.18
5.05
4.42
3.74
2.47
2.45
2.43
2.21
2.19
2.17
1.59
1.58
1.56
1.54
1.52
1.45
1.45
1.44
1.44
1.37
1.35
1.33
1.30
1.28
1.27
1.24
NHBoc
S
O
O
1.00
3.09
2.02
2.06
2.07
2.09
4.30
8.75
9.18
8.24
11.0
10.5
10.0
9.5
9.0
8.5
8.0
7.5
7.0
6.5
6.0
5.5
5.0
4.5
4.0
3.5
3.0
2.5
2.0
1.5
1.0
0.5
0.0
-0.5
-1
f1 (ppm)

[13]C NMR of **9:**

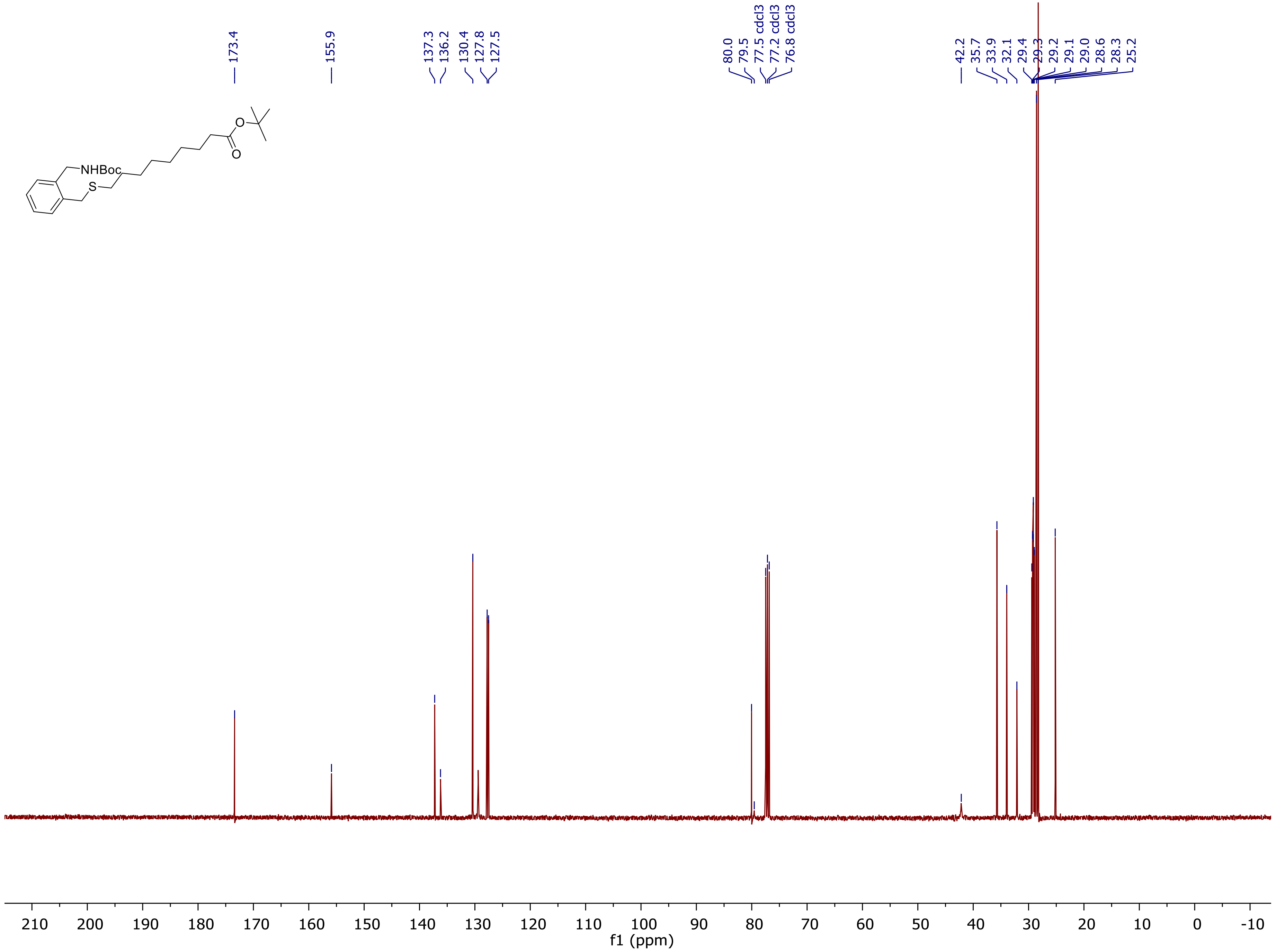
173.4
155.9
137.3
136.2
130.4
127.8
127.5
80.0
79.5
77.5 cdcl3
77.2 cdcl3
76.8 cdcl3
42.2
35.7
33.9
32.1
29.4
29.3
29.2
29.1
29.0
28.6
28.3
25.2
NHBoc
S
O
O
210 200 190 180 170 160 150 140 130 120 110 100 90 80 70 60 50 40 30 20 10 0 -10
f1 (ppm)

$^{1}$H NMR of **10:**

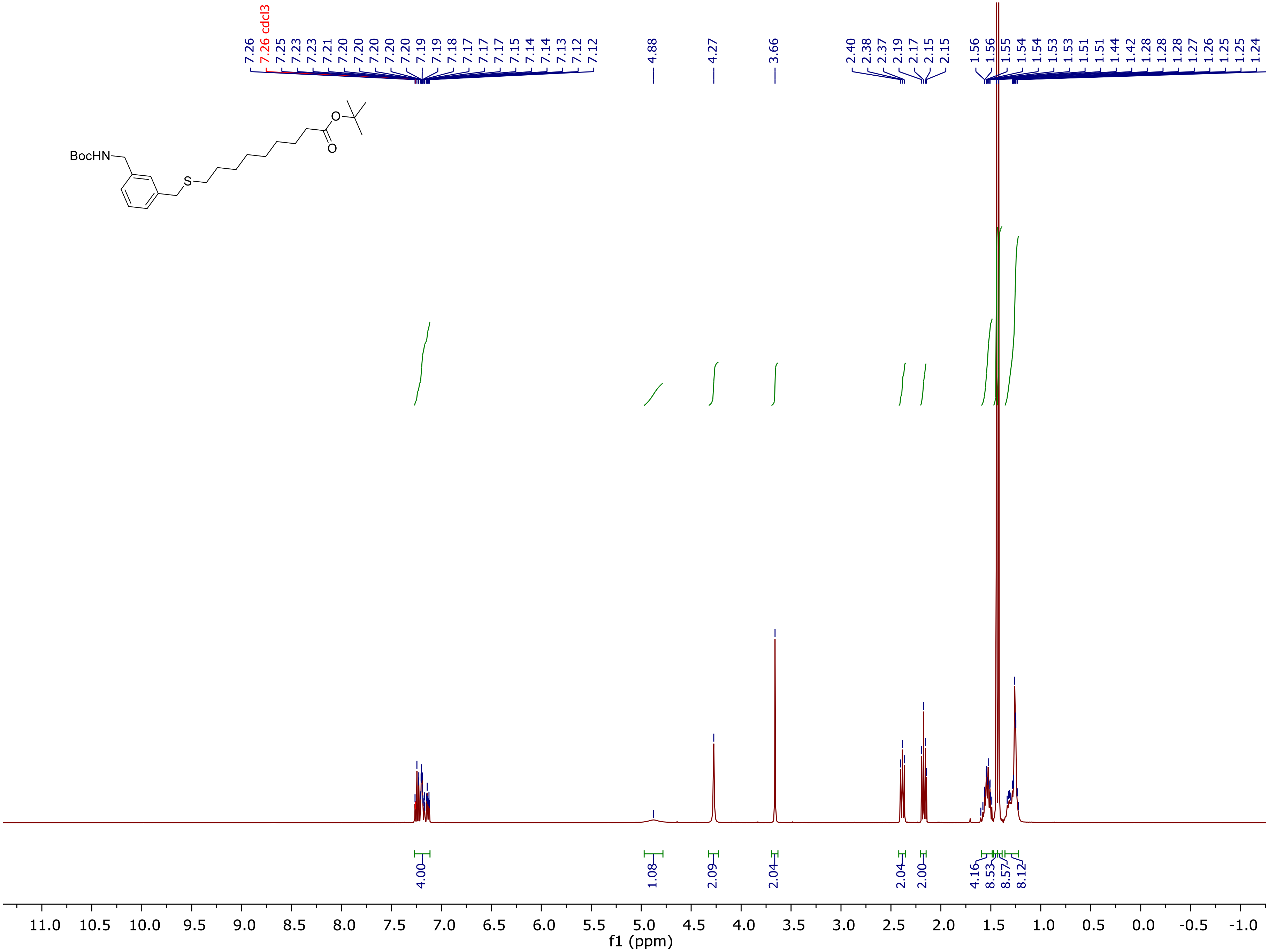
7.26
7.26 cdcl3
7.25
7.23
7.23
7.21
7.20
7.20
7.20
7.20
7.19
7.19
7.18
7.17
7.17
7.17
7.15
7.14
7.14
7.13
7.12
7.12
4.88
4.27
3.66
2.40
2.38
2.37
2.19
2.17
2.15
2.15
1.56
1.56
1.55
1.54
1.54
1.53
1.53
1.51
1.51
1.44
1.42
1.28
1.28
1.28
1.27
1.26
1.25
1.25
1.24
BocHN
S
O
O
4.00
1.08
2.09
2.04
2.04
2.00
4.16
8.53
8.57
8.12
11.0 10.5 10.0 9.5 9.0 8.5 8.0 7.5 7.0 6.5 6.0 5.5 5.0 4.5 4.0 3.5 3.0 2.5 2.0 1.5 1.0 0.5 0.0 -0.5 -1.0
f1 (ppm)

<sup></sup>

[13]C NMR of **10:**

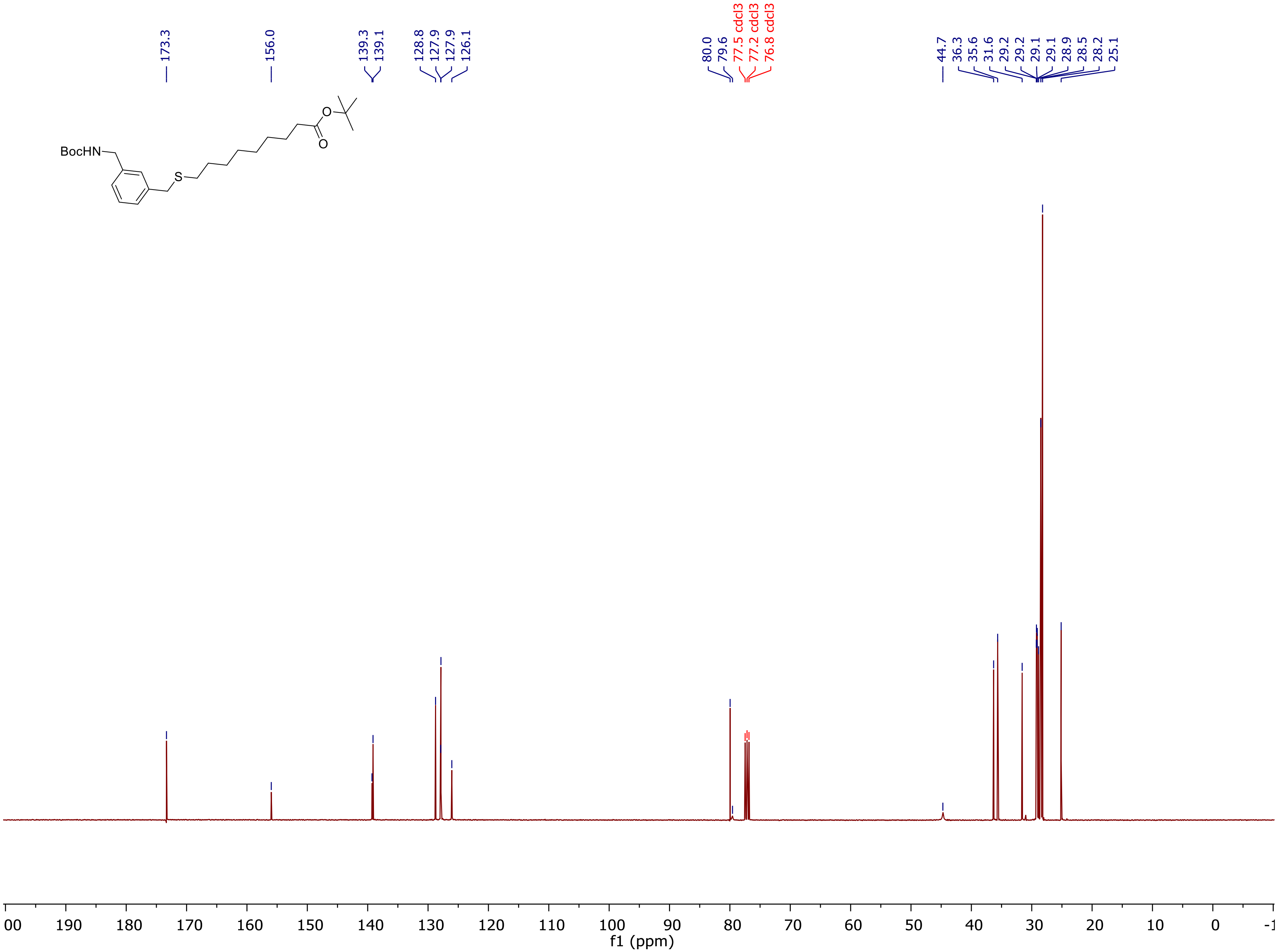

[1]H NMR of **11**:

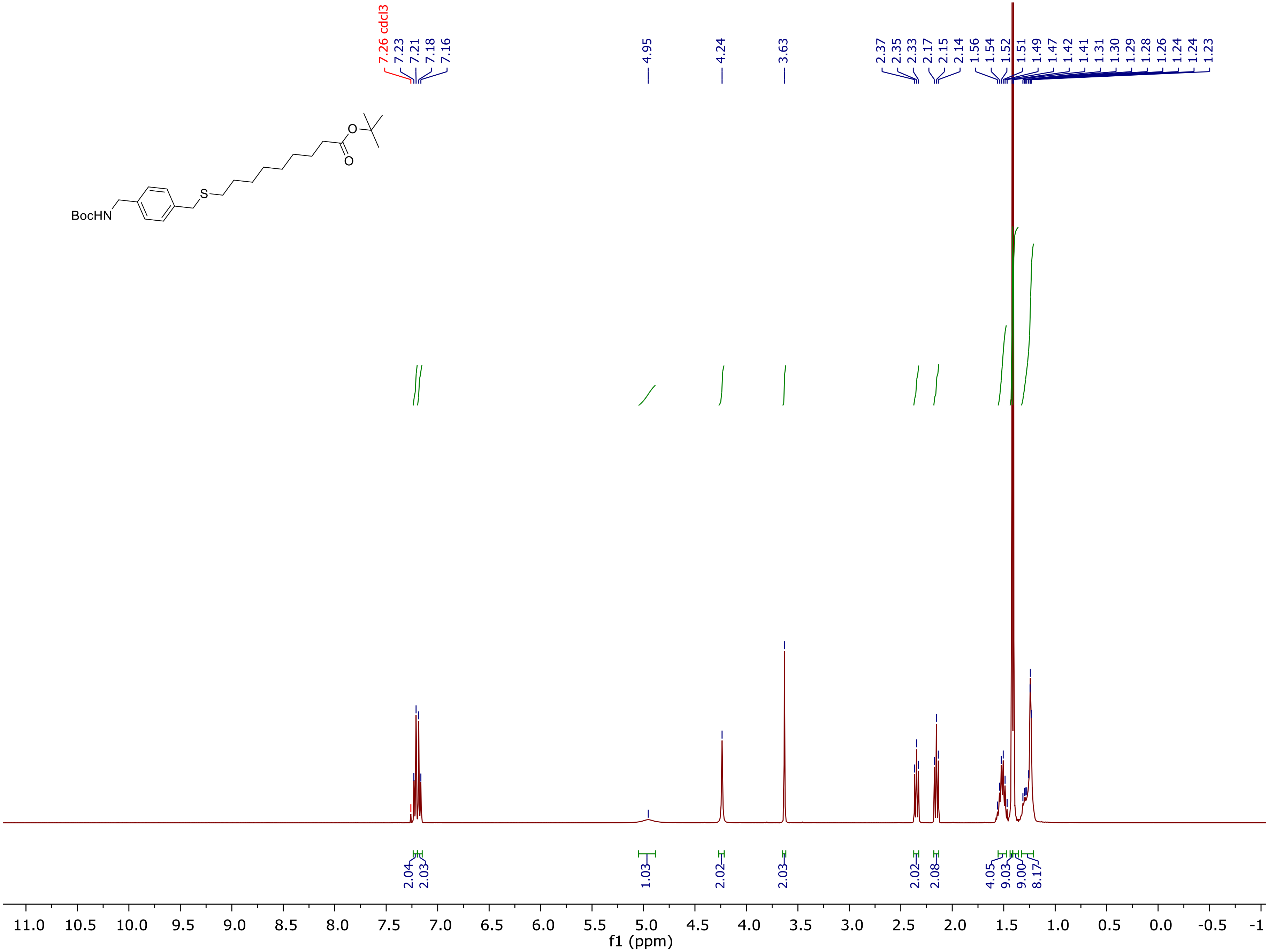
7.26 cdcl3
7.23
7.21
7.18
7.16
4.95
4.24
3.63
2.37
2.35
2.33
2.17
2.15
2.14
1.56
1.54
1.52
1.51
1.49
1.47
1.42
1.41
1.31
1.30
1.29
1.28
1.26
1.24
1.24
1.23
2.04
2.03
1.03
2.02
2.03
2.02
2.08
4.05
9.03
9.00
8.17
f1 (ppm)

$^{13}C$ NMR of **11:**

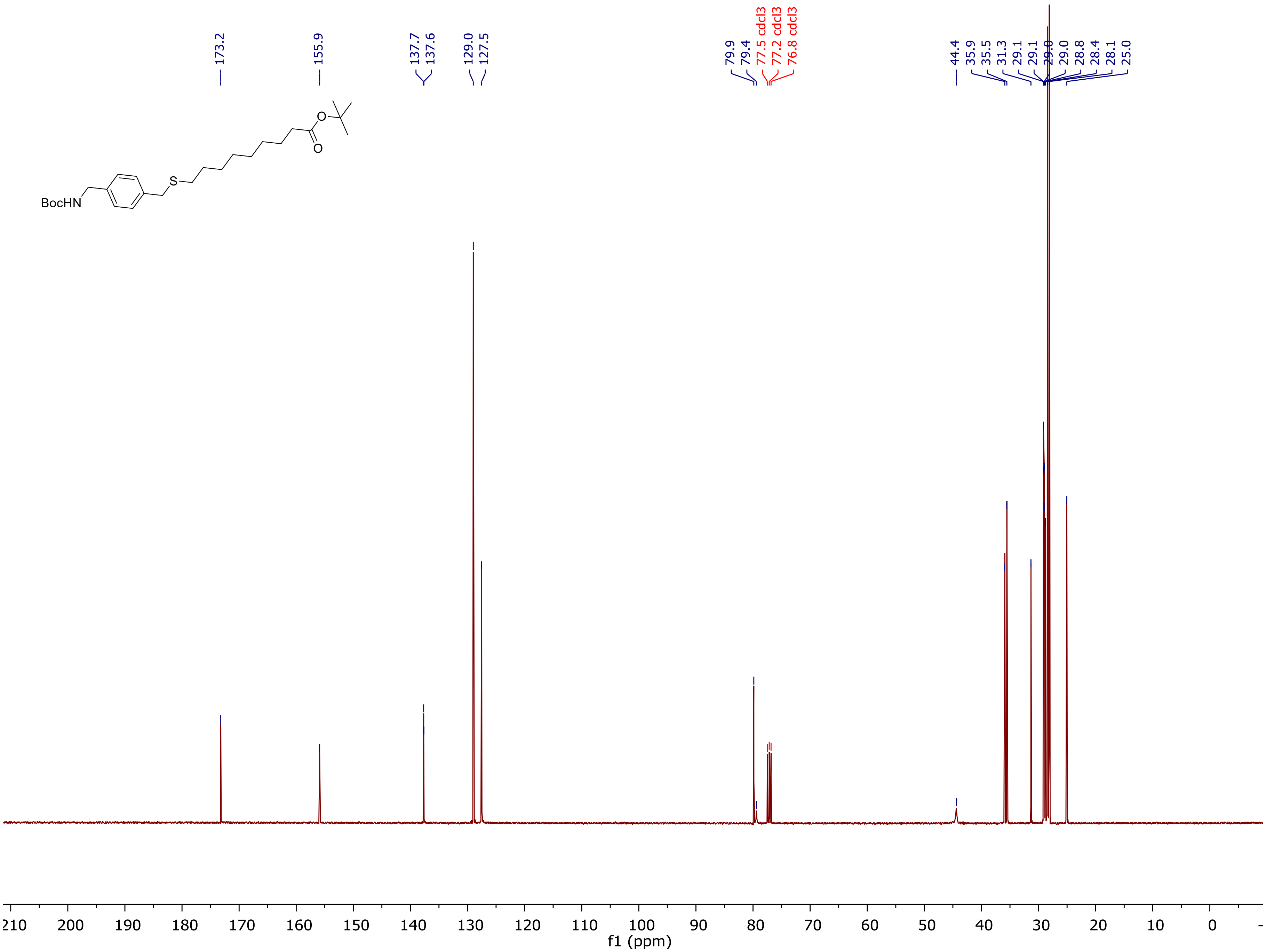
173.2
155.9
137.7
137.6
129.0
127.5
79.9
79.4
77.5 cdcl3
77.2 cdcl3
76.8 cdcl3
44.4
35.9
35.5
31.3
29.1
29.1
29.0
29.0
28.8
28.4
28.1
25.0
BocHN
S
O
O
f1 (ppm)

[1]H NMR of **12:**

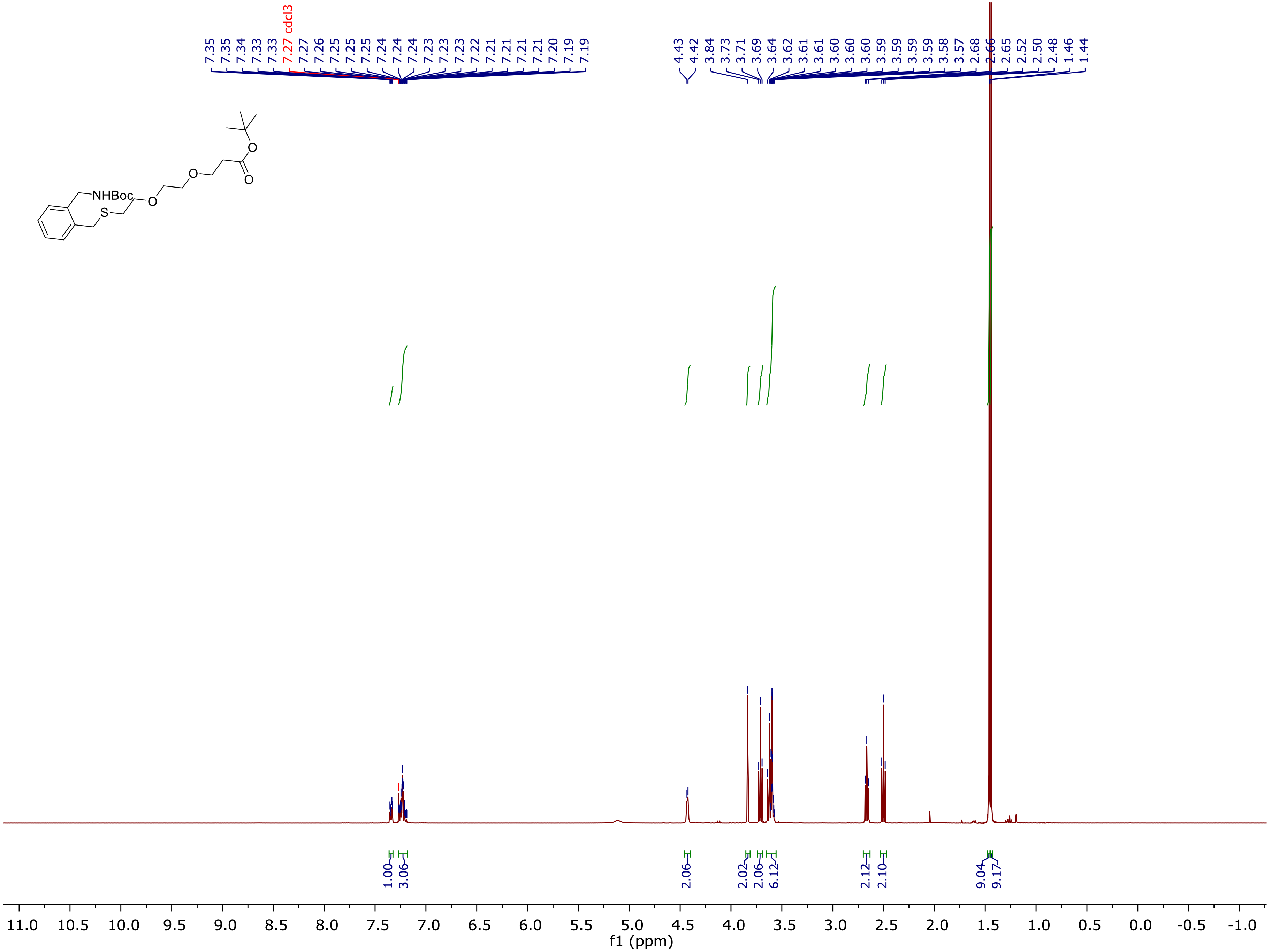

[13]C NMR of **12:**

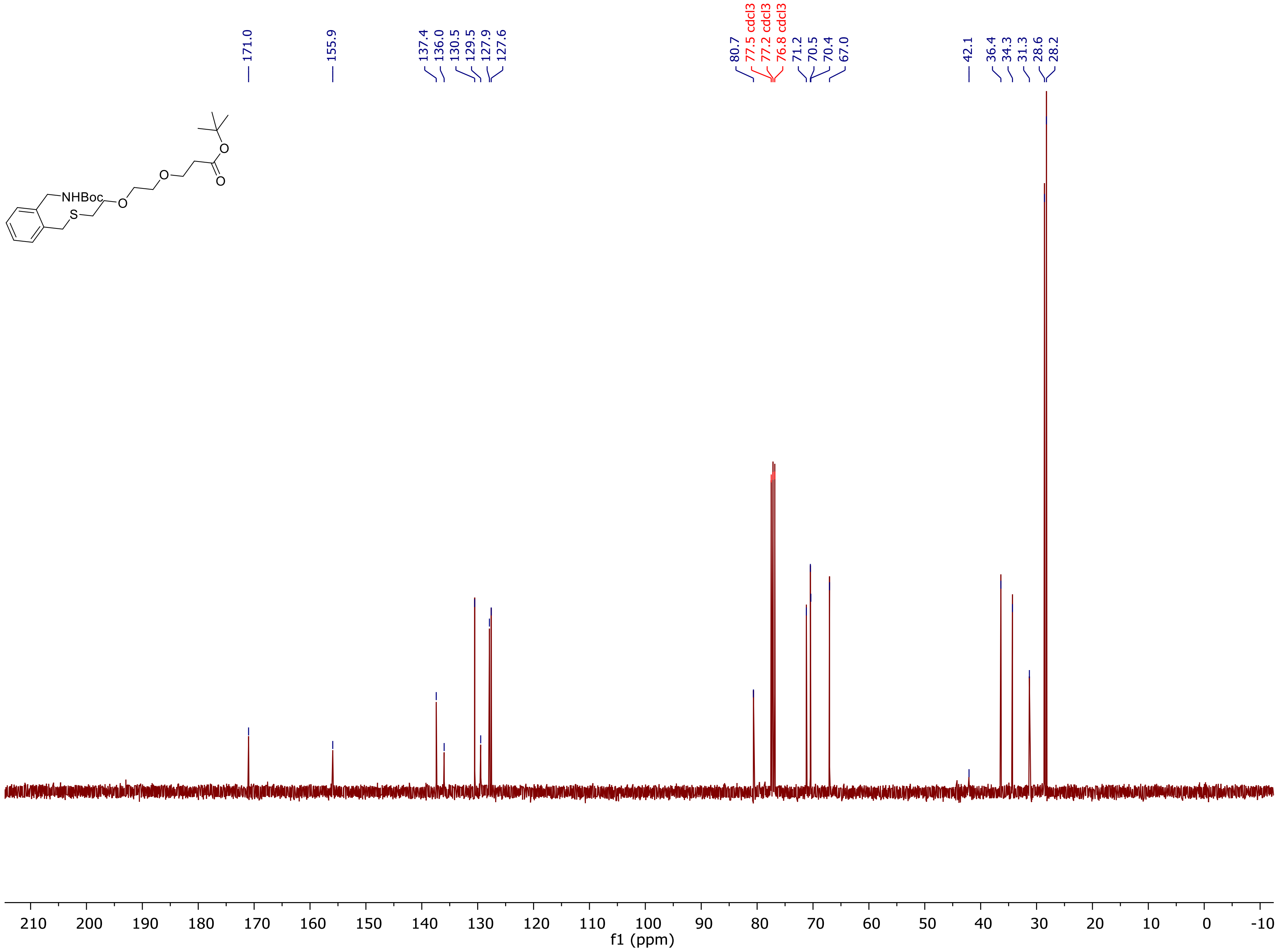

NHBoc
S
O
O
O
O
171.0
155.9
137.4
136.0
130.5
129.5
127.9
127.6
80.7
77.5 cdcl3
77.2 cdcl3
76.8 cdcl3
71.2
70.5
70.4
67.0
42.1
36.4
34.3
31.3
28.6
28.2
210
200
190
180
170
160
150
140
130
120
110
100
90
80
70
60
50
40
30
20
10
0
-10
f1 (ppm)

$^{1}$H NMR of **13:**

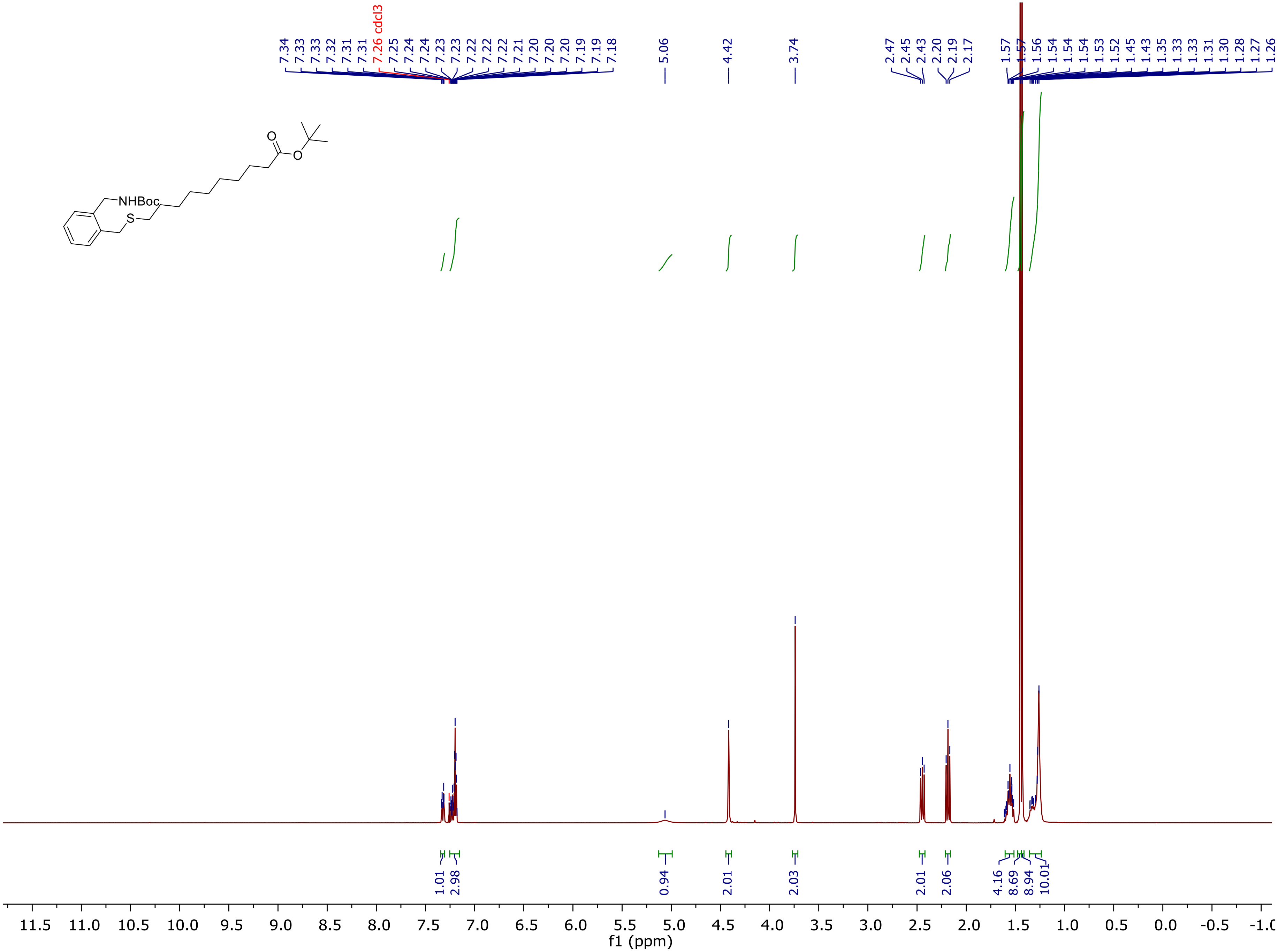
NHBoc
S
O
O
7.34 7.33 7.33 7.32 7.31 7.31 7.26 cdcl3 7.25 7.24 7.24 7.23 7.23 7.22 7.22 7.22 7.21 7.20 7.20 7.20 7.19 7.19 7.18
5.06
4.42
3.74
2.47 2.45 2.43 2.20 2.19 2.17
1.57 1.57 1.56 1.54 1.54 1.53 1.52 1.45 1.43 1.35 1.33 1.33 1.31 1.30 1.28 1.27 1.26
1.01 2.98 0.94 2.01 2.03 2.01 2.06 4.16 8.69 8.94 10.01
11.5 11.0 10.5 10.0 9.5 9.0 8.5 8.0 7.5 7.0 6.5 6.0 5.5 5.0 4.5 4.0 3.5 3.0 2.5 2.0 1.5 1.0 0.5 0.0 -0.5 -1.0
f1 (ppm)

$^{13}$C NMR of **13:**

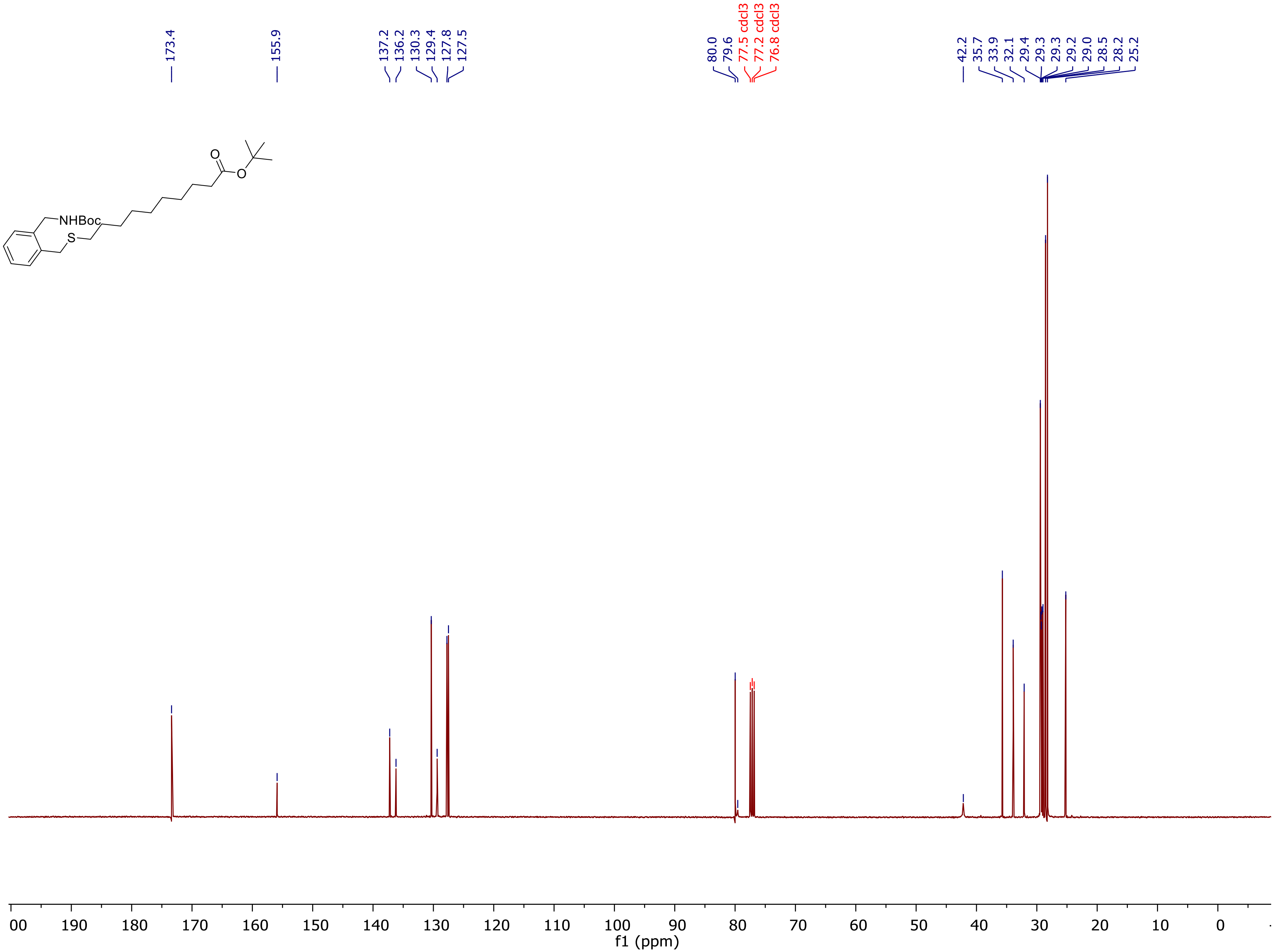

173.4
155.9
137.2
136.2
130.3
129.4
127.8
127.5
80.0
79.6
77.5 cdcl3
77.2 cdcl3
76.8 cdcl3
42.2
35.7
33.9
32.1
29.4
29.3
29.3
29.2
29.0
28.5
28.2
25.2
NHBoc
S
O
O
00
190
180
170
160
150
140
130
120
110
100
90
80
70
60
50
40
30
20
10
0
f1 (ppm)

[1]H NMR of **14:**

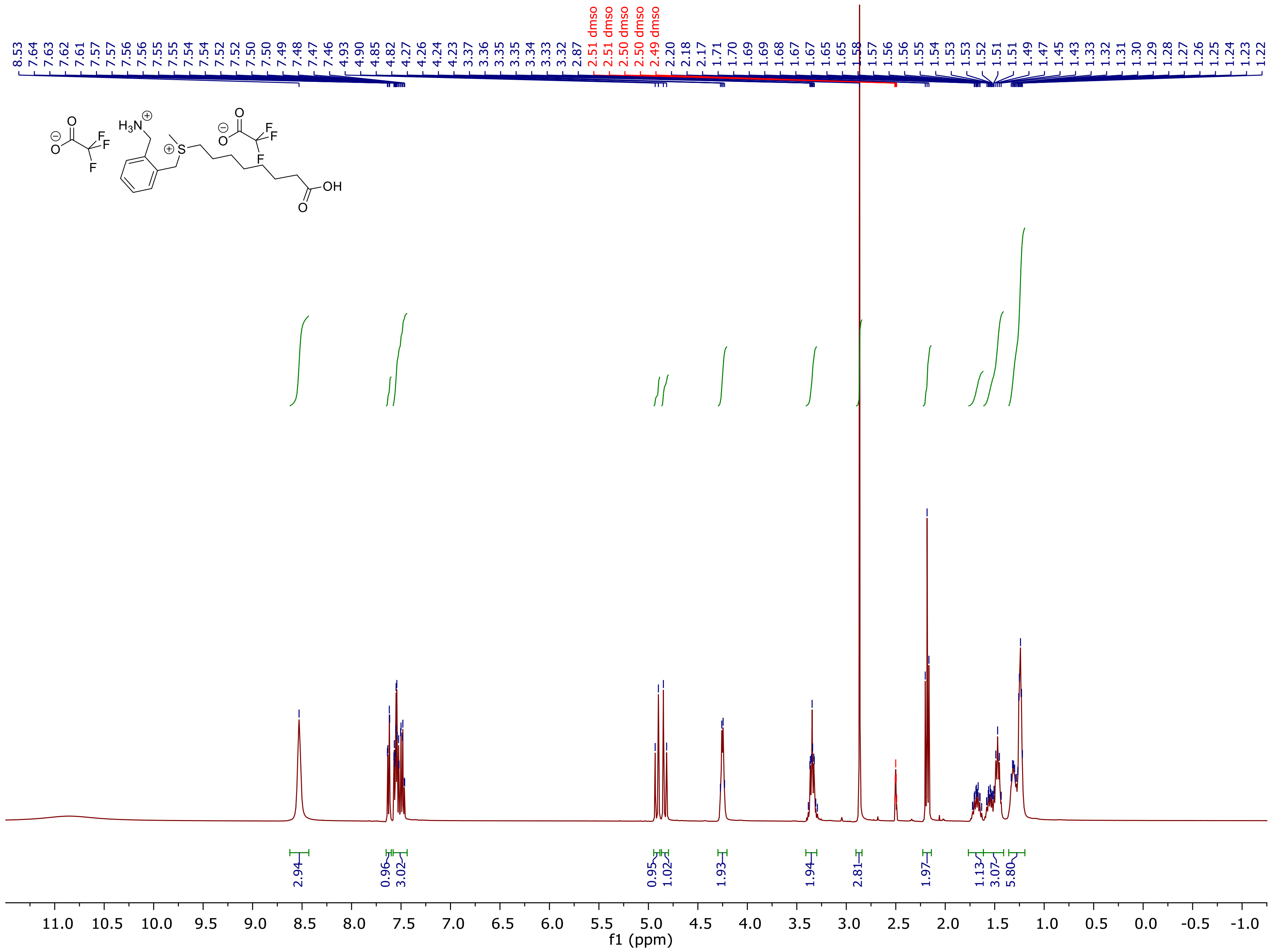

[13]C NMR of **14:**

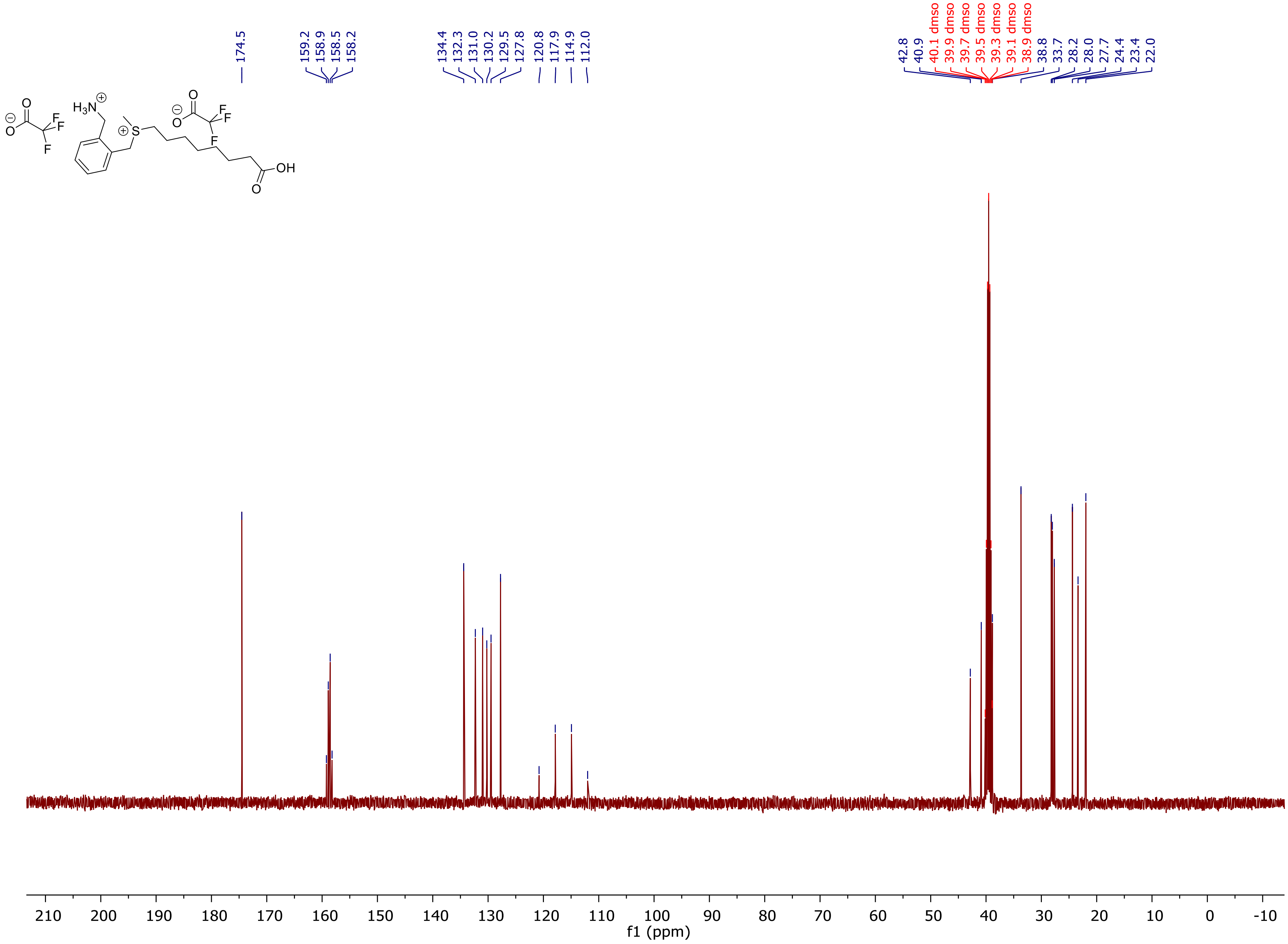

174.5
159.2
158.9
158.5
158.2
134.4
132.3
131.0
130.2
129.5
127.8
120.8
117.9
114.9
112.0
42.8
40.9
40.1 dmso
39.9 dmso
39.7 dmso
39.5 dmso
39.3 dmso
39.1 dmso
38.9 dmso
38.8
33.7
28.2
28.0
27.7
24.4
23.4
22.0
210
200
190
180
170
160
150
140
130
120
110
100
90
80
70
60
50
40
30
20
10
0
-10
f1 (ppm)

$^1$H NMR of **15:**

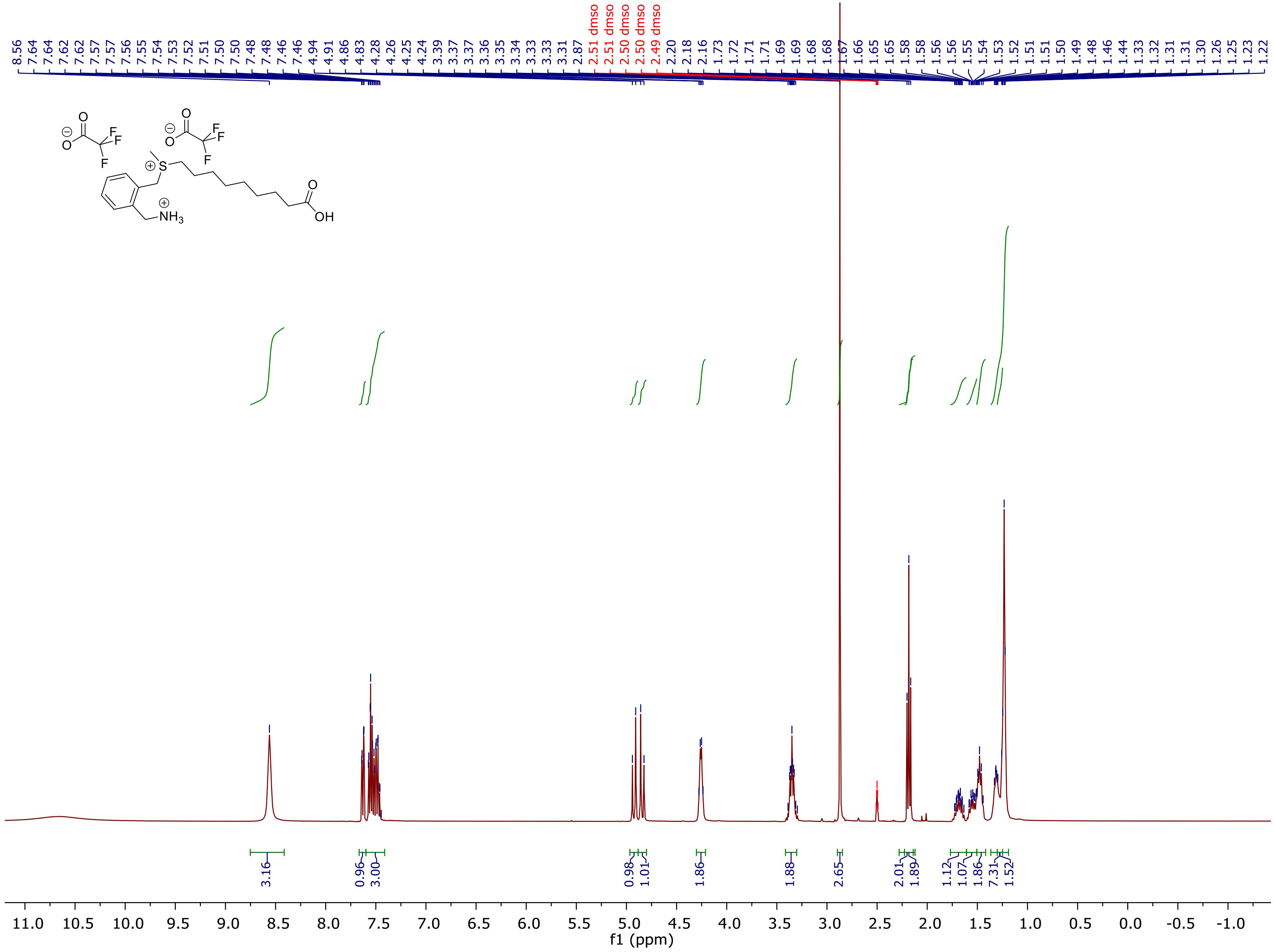

8.56
7.64
7.64
7.62
7.62
7.57
7.57
7.56
7.55
7.54
7.53
7.52
7.51
7.50
7.50
7.48
7.48
7.46
7.46
4.94
4.91
4.86
4.83
4.28
4.26
4.25
4.24
3.39
3.37
3.37
3.36
3.35
3.34
3.33
3.33
3.31
2.87
2.51 dmso
2.51 dmso
2.50 dmso
2.50 dmso
2.49 dmso
2.20
2.18
2.16
1.73
1.72
1.71
1.71
1.69
1.69
1.68
1.68
1.67
1.66
1.65
1.65
1.58
1.58
1.56
1.56
1.55
1.54
1.53
1.52
1.51
1.51
1.50
1.49
1.48
1.46
1.44
1.33
1.32
1.31
1.31
1.30
1.26
1.25
1.23
1.22
O
F
F
F
O
O
F
F
F
O
O
S
NH3
O
OH
3.16
0.96
3.00
0.98
1.01
1.86
1.88
2.65
2.01
1.89
1.12
1.07
1.86
7.31
1.52
11.0
10.5
10.0
9.5
9.0
8.5
8.0
7.5
7.0
6.5
6.0
5.5
5.0
4.5
4.0
3.5
3.0
2.5
2.0
1.5
1.0
0.5
0.0
-0.5
-1.0
f1 (ppm)

$^{13}C$ NMR of **15:**

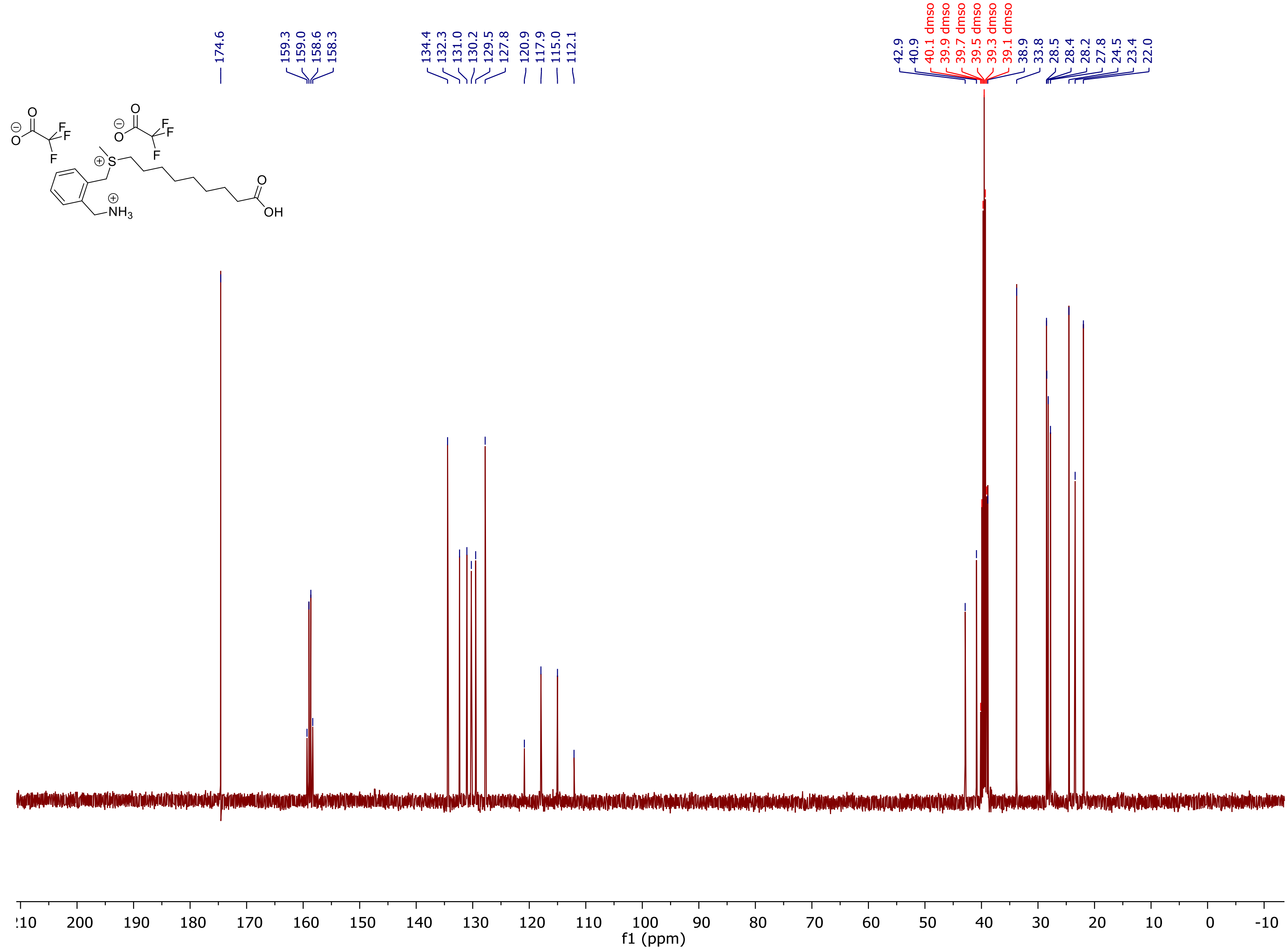
174.6
159.3
159.0
158.6
158.3
134.4
132.3
131.0
130.2
129.5
127.8
120.9
117.9
115.0
112.1
42.9
40.9
40.1 dmso
39.9 dmso
39.7 dmso
39.5 dmso
39.3 dmso
39.1 dmso
38.9
33.8
28.5
28.4
28.2
27.8
24.5
23.4
22.0
f1 (ppm)

[1]H NMR of **16:**

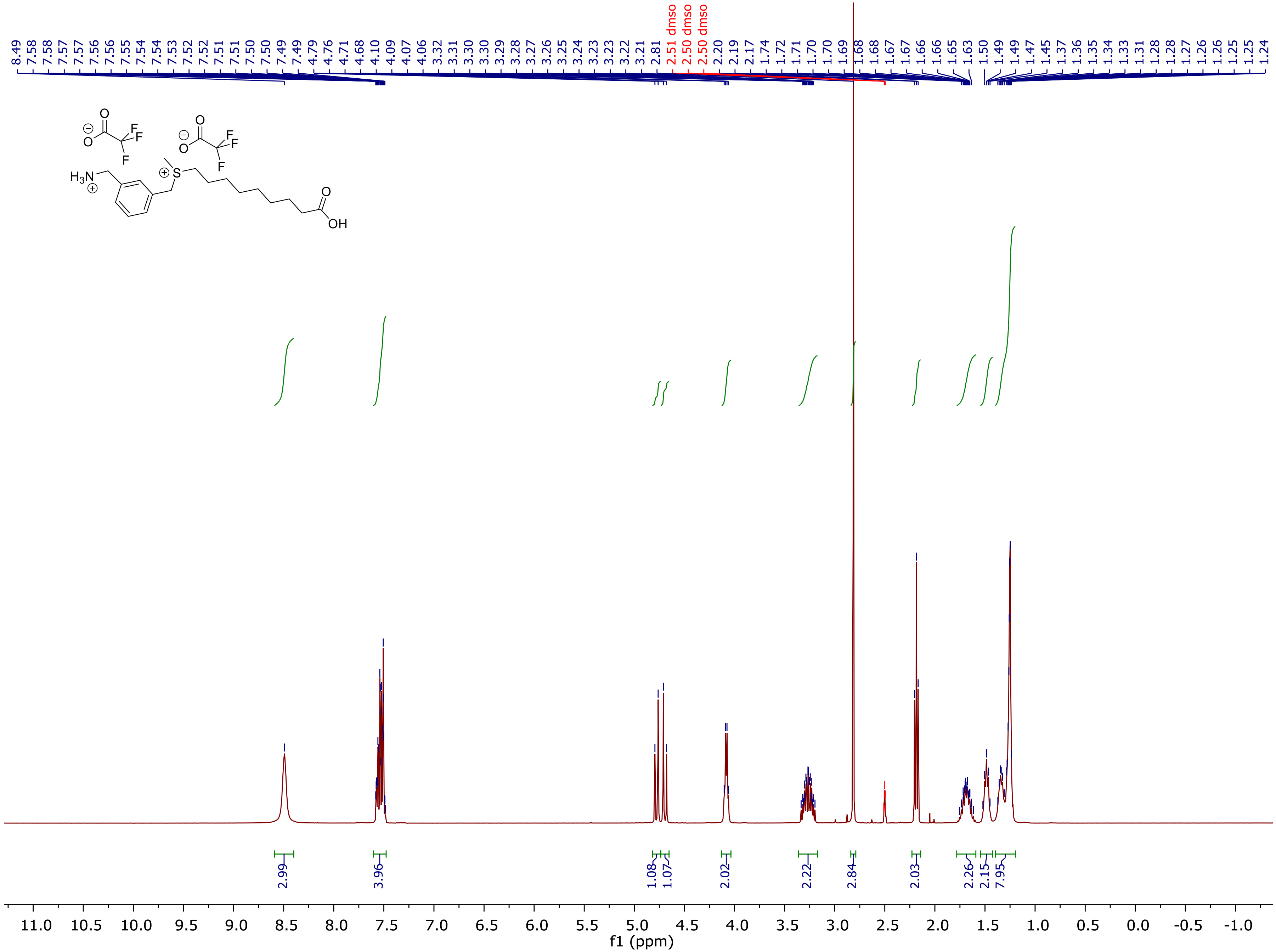

8.49
7.58
7.58
7.57
7.57
7.56
7.56
7.55
7.54
7.54
7.53
7.52
7.52
7.51
7.51
7.50
7.50
7.49
7.49
4.79
4.76
4.71
4.68
4.10
4.09
4.07
4.06
3.32
3.31
3.30
3.30
3.29
3.28
3.27
3.26
3.25
3.24
3.23
3.23
3.22
3.21
2.81
2.51 dmso
2.50 dmso
2.50 dmso
2.20
2.19
2.17
1.74
1.72
1.71
1.70
1.70
1.69
1.68
1.68
1.67
1.67
1.66
1.66
1.65
1.63
1.50
1.49
1.49
1.47
1.45
1.37
1.36
1.35
1.34
1.33
1.31
1.28
1.27
1.26
1.26
1.25
1.24
2.99
3.96
1.08
1.07
2.02
2.22
2.84
2.03
2.26
2.15
7.95
11.0
10.5
10.0
9.5
9.0
8.5
8.0
7.5
7.0
6.5
6.0
5.5
5.0
4.5
4.0
3.5
3.0
2.5
2.0
1.5
1.0
0.5
0.0
-0.5
-1.0
f1 (ppm)

$^{13}C$ NMR of **16:**

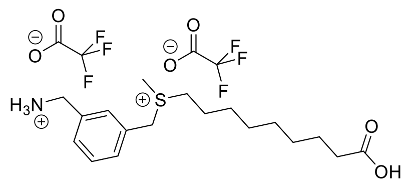

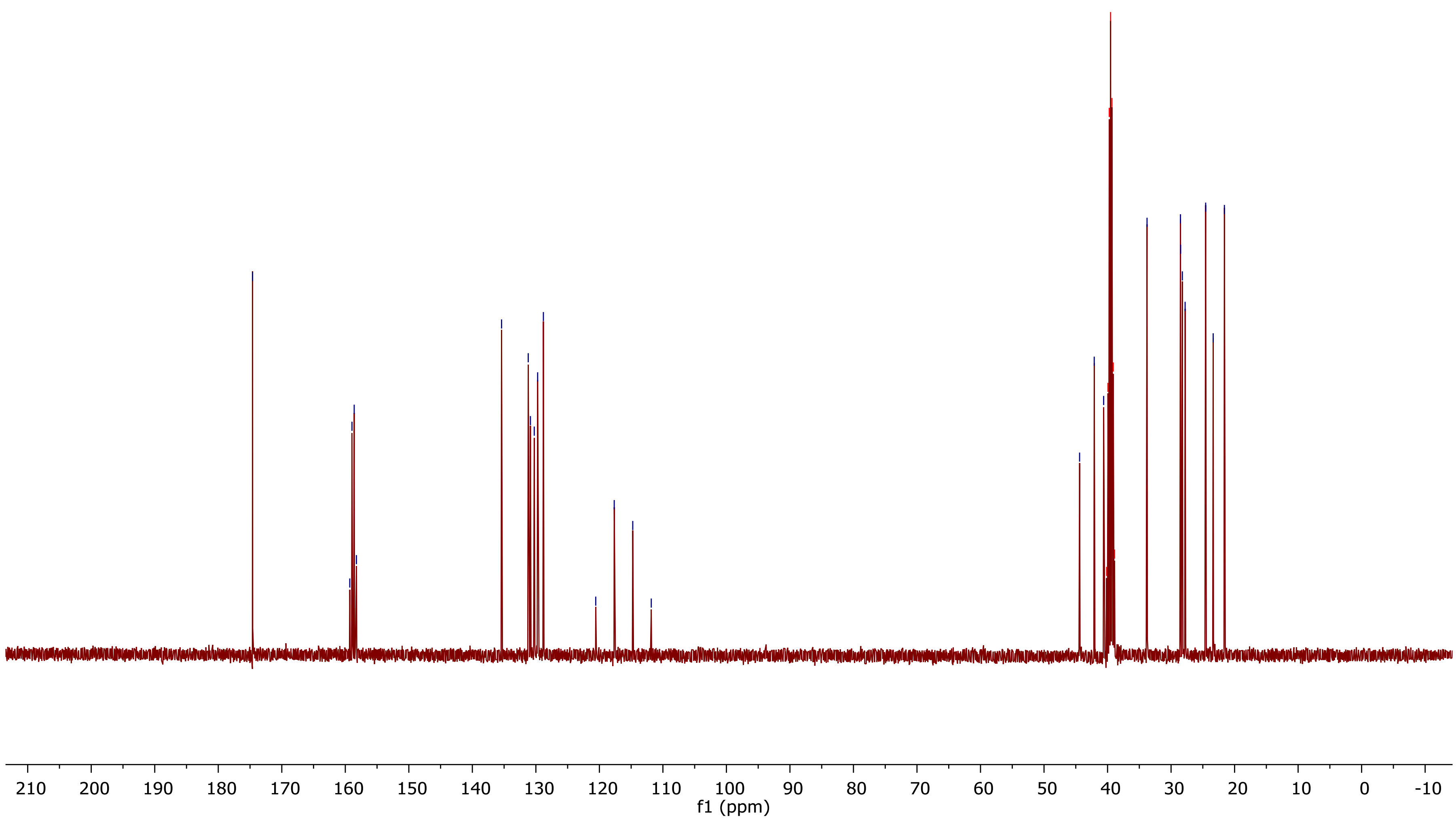
174.6
159.3
158.9
158.6
158.3
135.4
131.2
130.9
130.3
129.7
128.8
120.6
117.7
114.7
111.8
44.4
42.1
40.6
40.1 dmso
39.9 dmso
39.7 dmso
39.5 dmso
39.3 dmso
39.1 dmso
38.9 dmso
33.8
28.5
28.5
28.2
27.8
24.5
23.4
21.6
f1 (ppm)

$^{1}$H NMR of **17:**

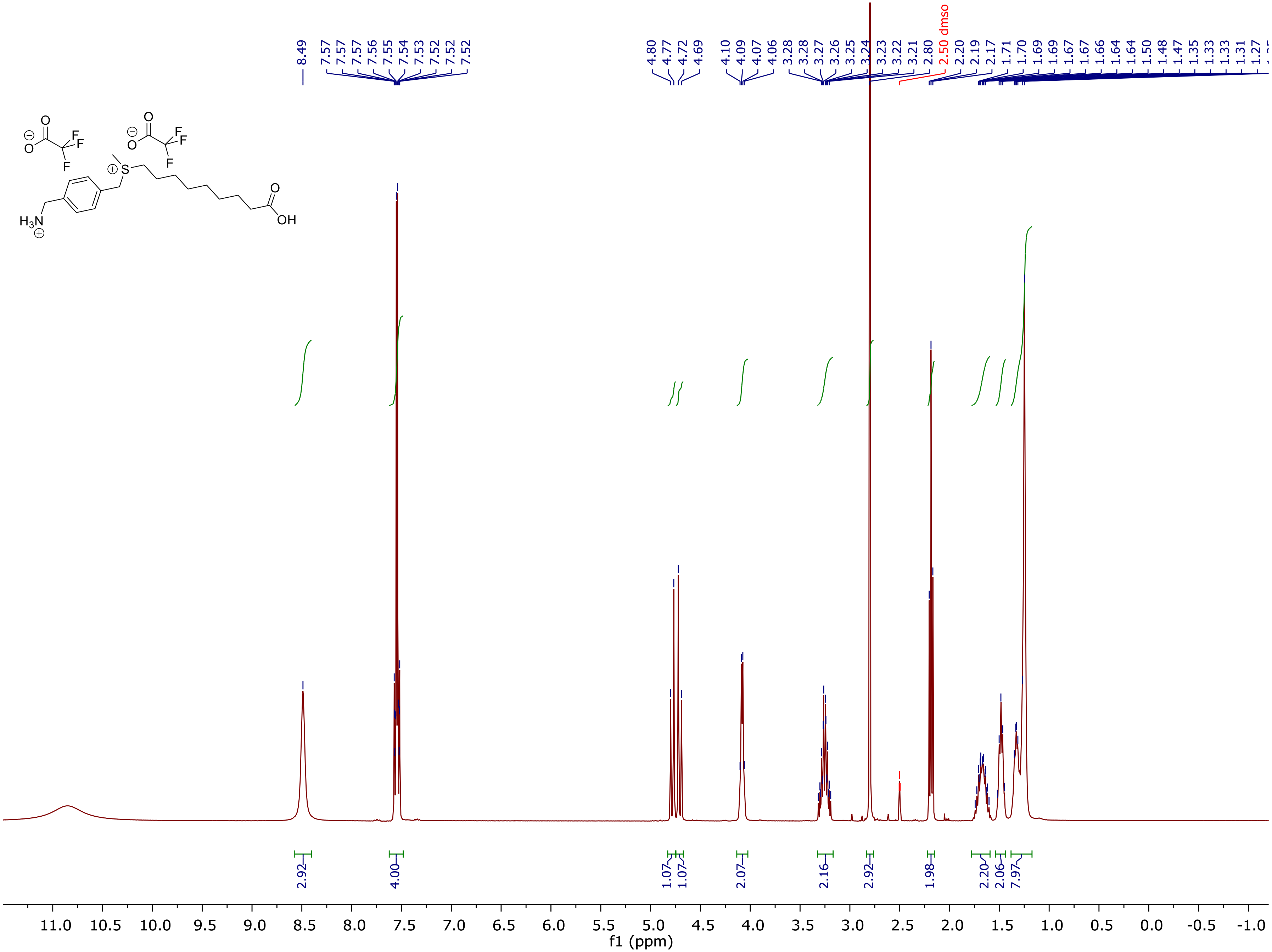

[13]C NMR of **17:**

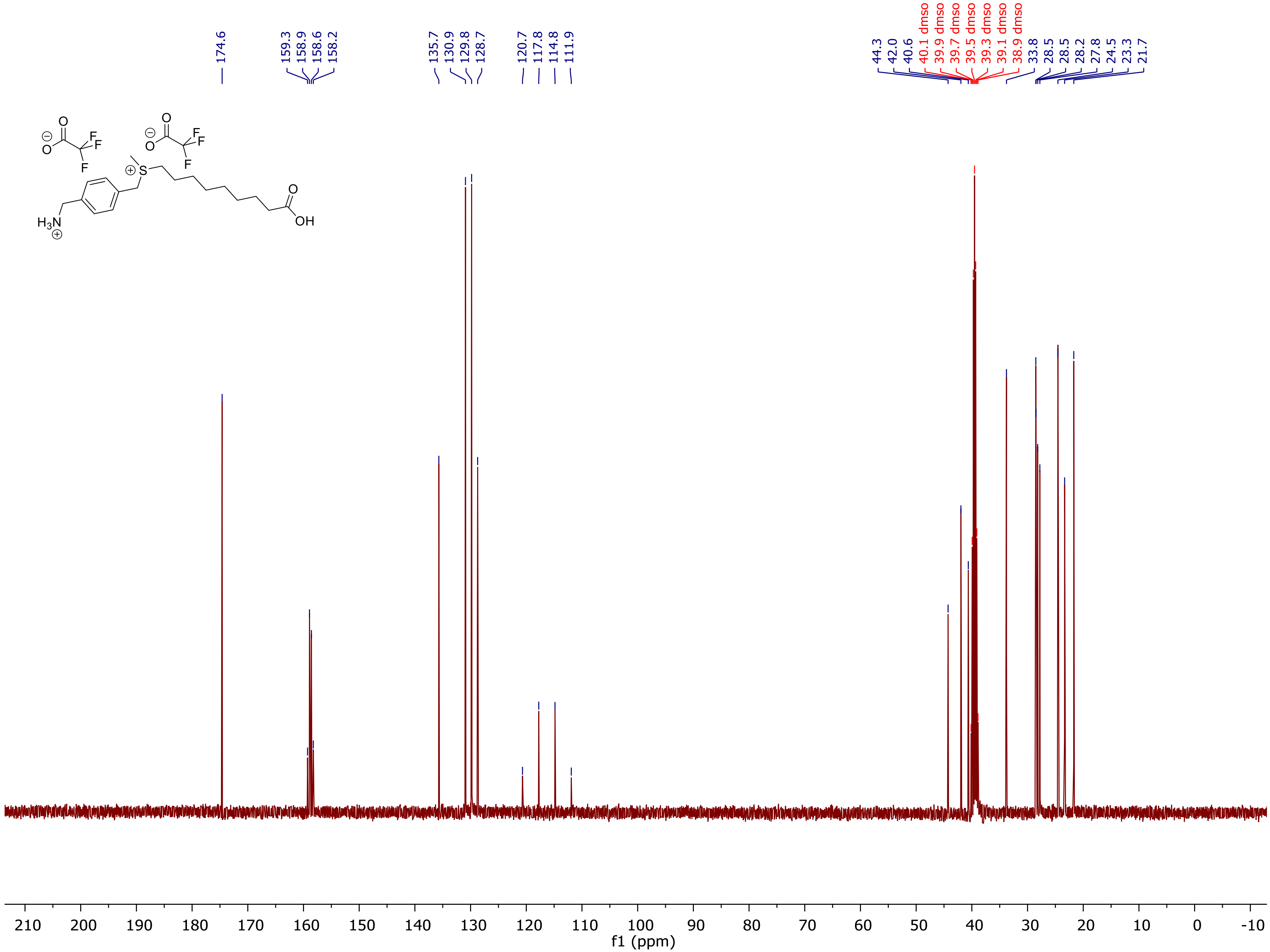

174.6
159.3
158.9
158.6
158.2
135.7
130.9
129.8
128.7
120.7
117.8
114.8
111.9
44.3
42.0
40.6
40.1 dmso
39.9 dmso
39.7 dmso
39.5 dmso
39.3 dmso
39.1 dmso
38.9 dmso
33.8
28.5
28.5
28.2
27.8
24.5
23.3
21.7
f1 (ppm)

[1]H NMR of **18:**

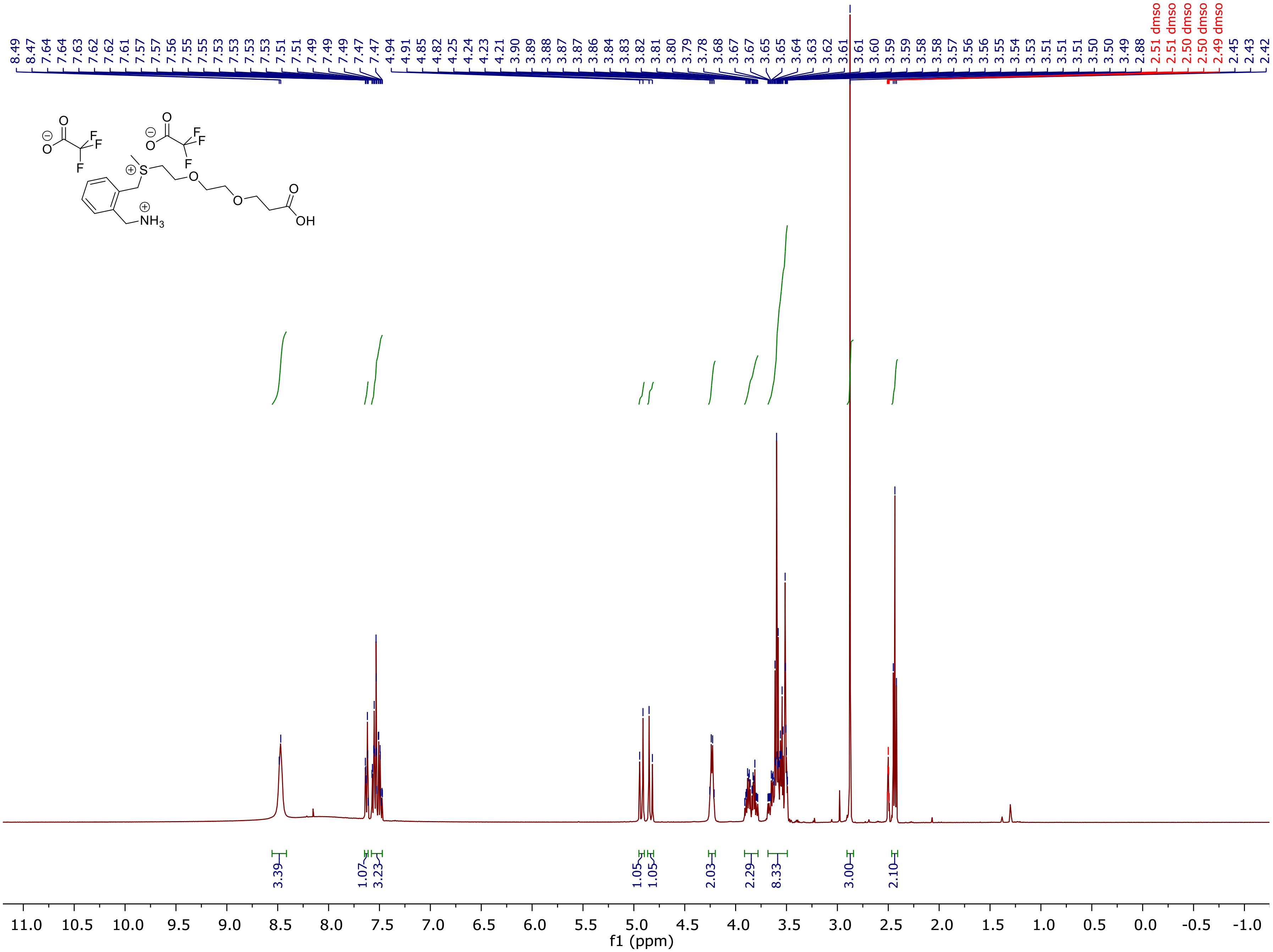

8.49 8.47 7.64 7.64 7.63 7.62 7.62 7.61 7.57 7.57 7.56 7.55 7.55 7.53 7.53 7.53 7.53 7.51 7.51 7.49 7.49 7.47 7.47 4.94 4.91 4.85 4.82 4.25 4.24 4.23 4.21 3.90 3.89 3.88 3.87 3.87 3.86 3.84 3.83 3.82 3.81 3.80 3.79 3.78 3.68 3.67 3.67 3.65 3.65 3.64 3.63 3.62 3.61 3.61 3.60 3.59 3.59 3.58 3.58 3.57 3.56 3.56 3.55 3.54 3.53 3.51 3.51 3.51 3.50 3.49 2.88
2.51 dmso 2.51 dmso 2.50 dmso 2.50 dmso 2.49 dmso
2.45 2.43 2.42
O
F
S
NH3
OH
3.39 1.07 3.23 1.05 1.05 2.03 2.29 8.33 3.00 2.10
11.0 10.5 10.0 9.5 9.0 8.5 8.0 7.5 7.0 6.5 6.0 5.5 5.0 4.5 4.0 3.5 3.0 2.5 2.0 1.5 1.0 0.5 0.0 -0.5 -1.0
f1 (ppm)

[13]C NMR of **18:**

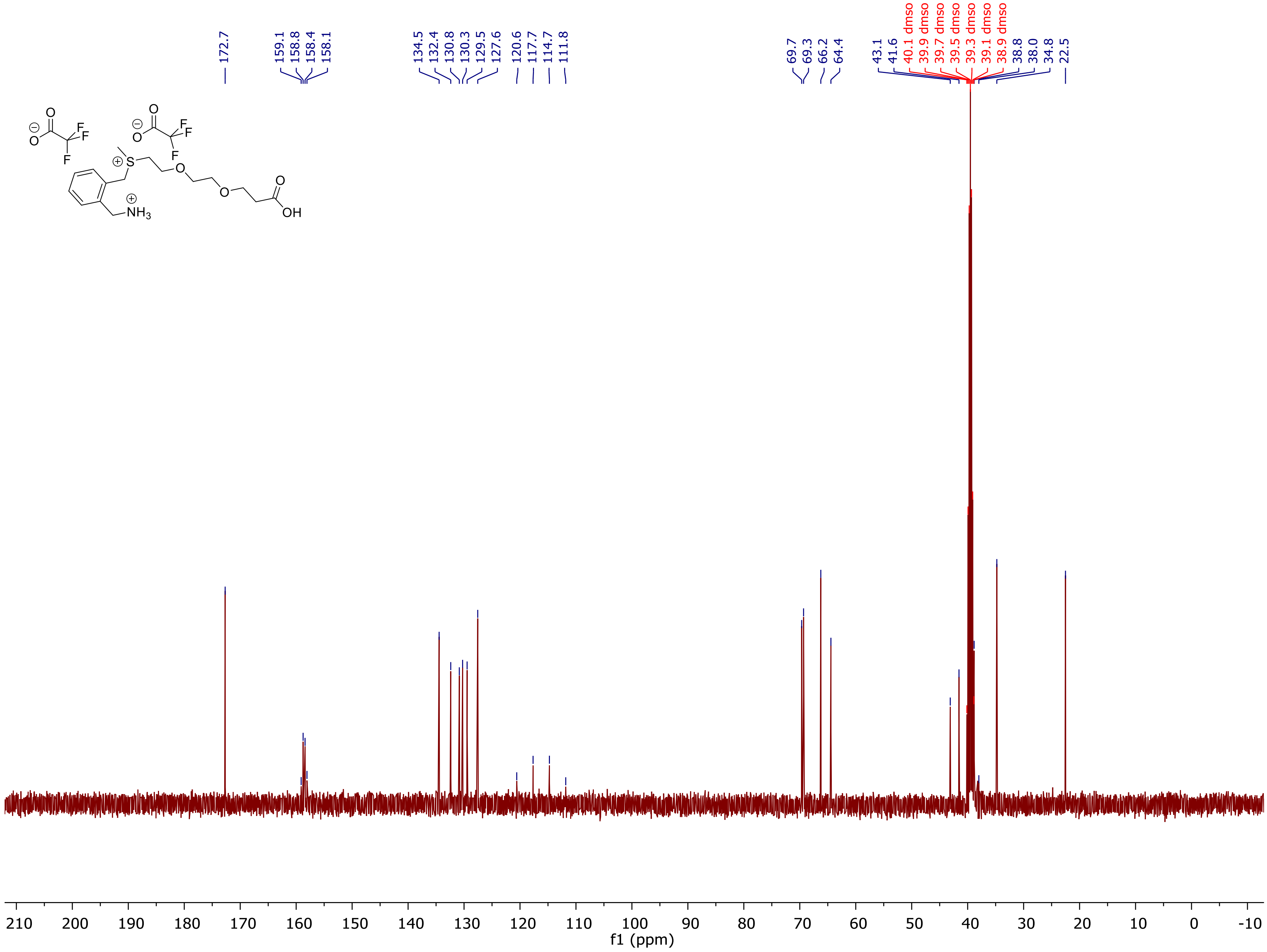
172.7
159.1
158.8
158.4
158.1
134.5
132.4
130.8
130.3
129.5
127.6
120.6
117.7
114.7
111.8
69.7
69.3
66.2
64.4
43.1
41.6
40.1 dmso
39.9 dmso
39.7 dmso
39.5 dmso
39.3 dmso
39.1 dmso
38.9 dmso
38.8
38.0
34.8
22.5
f1 (ppm)

$^1H$ NMR of **19:**

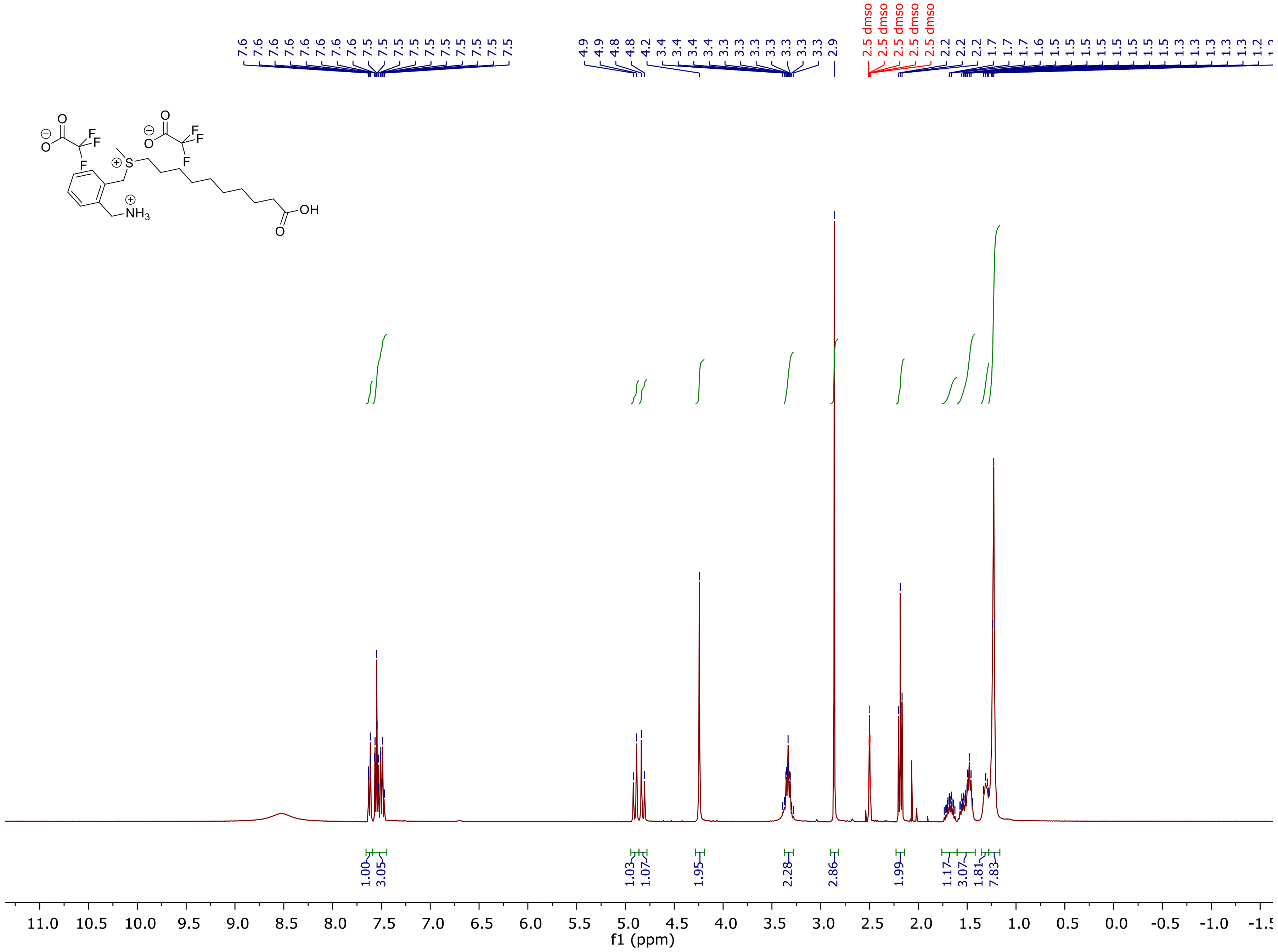

$^{13}$C NMR of **19:**

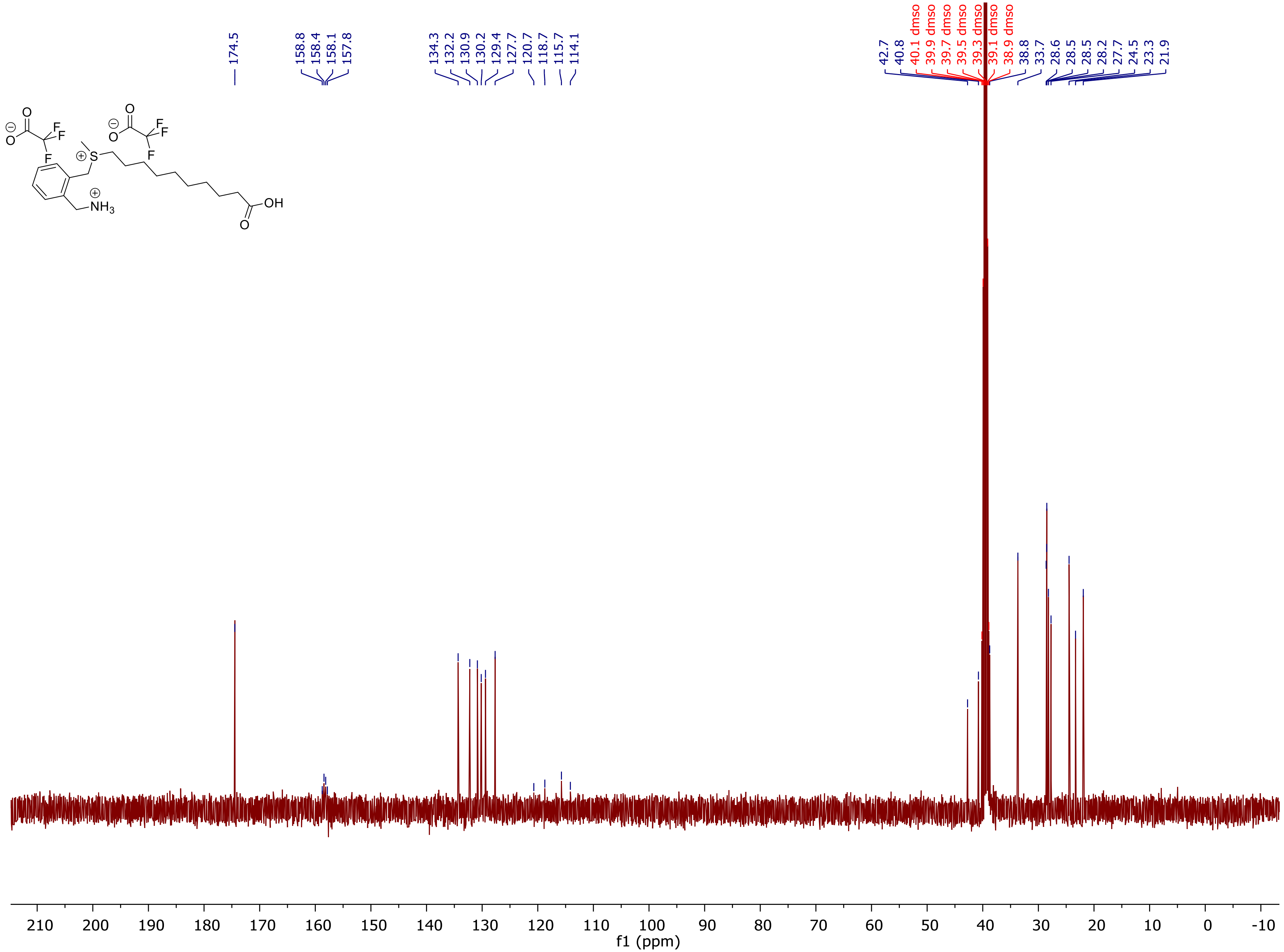

O
F
F
F
S
NH3
OH
174.5
158.8
158.4
158.1
157.8
134.3
132.2
130.9
130.2
129.4
127.7
120.7
118.7
115.7
114.1
42.7
40.8
40.1 dmso
39.9 dmso
39.7 dmso
39.5 dmso
39.3 dmso
39.1 dmso
38.9 dmso
38.8
33.7
28.6
28.5
28.5
28.2
27.7
24.5
23.3
21.9
210 200 190 180 170 160 150 140 130 120 110 100 90 80 70 60 50 40 30 20 10 0 -10
f1 (ppm)

[1]H NMR of ***CTX-NH$_3$-CF$_3$CO$_2$:***

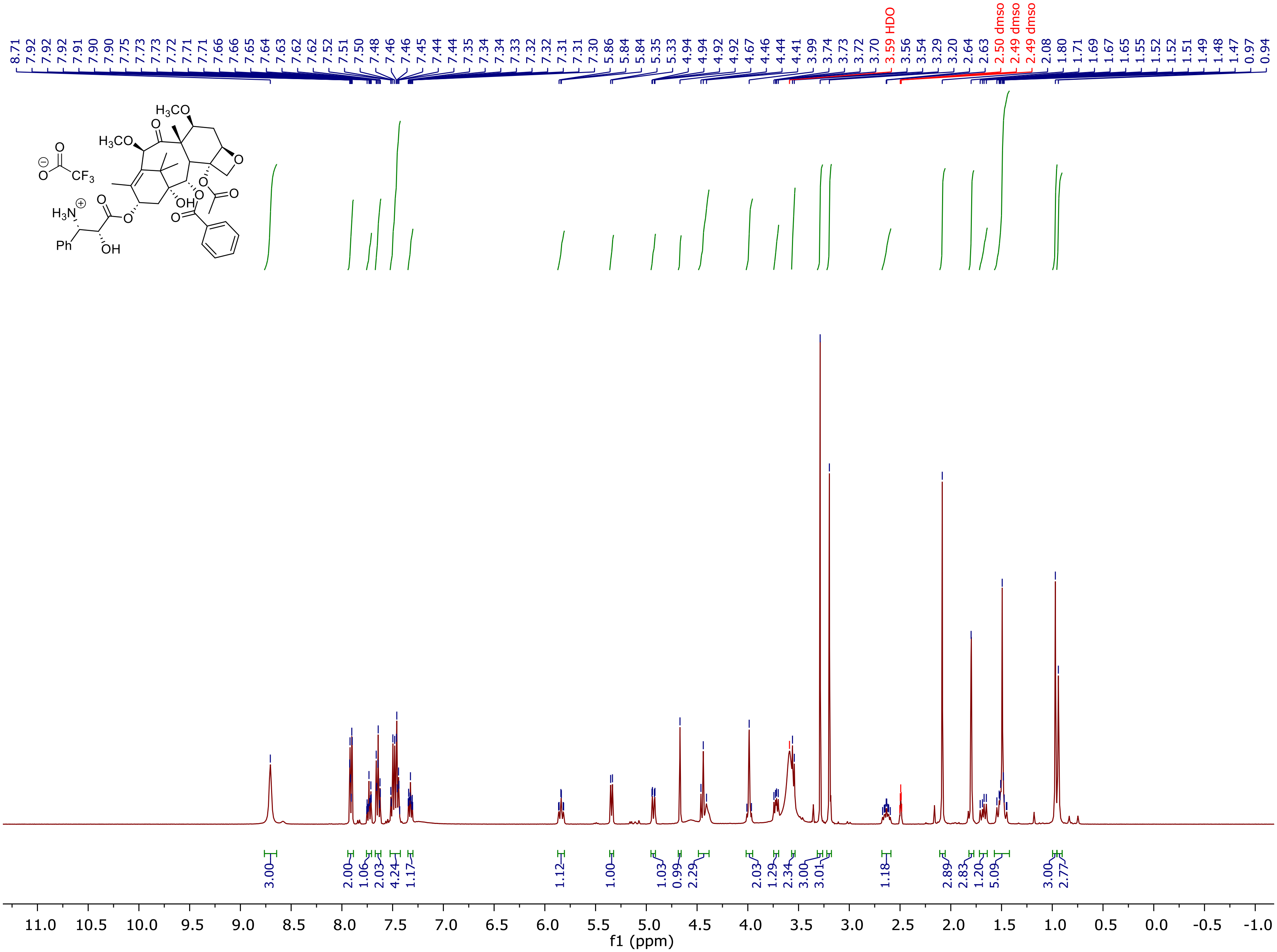

[13]C NMR of ***CTX-NH$_3$-CF$_3$CO$_2$:***

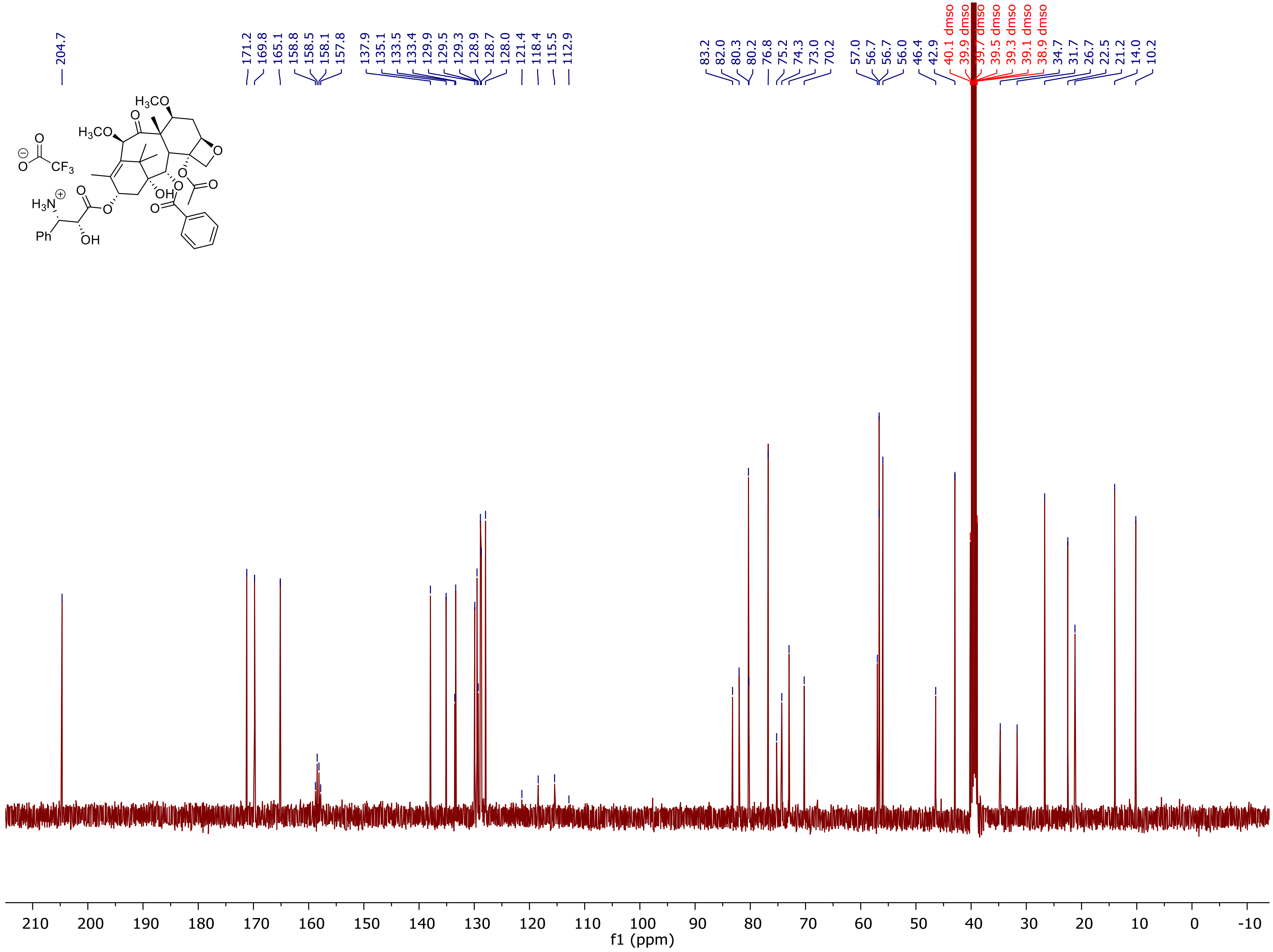

[1]H NMR of **Probe 1:**

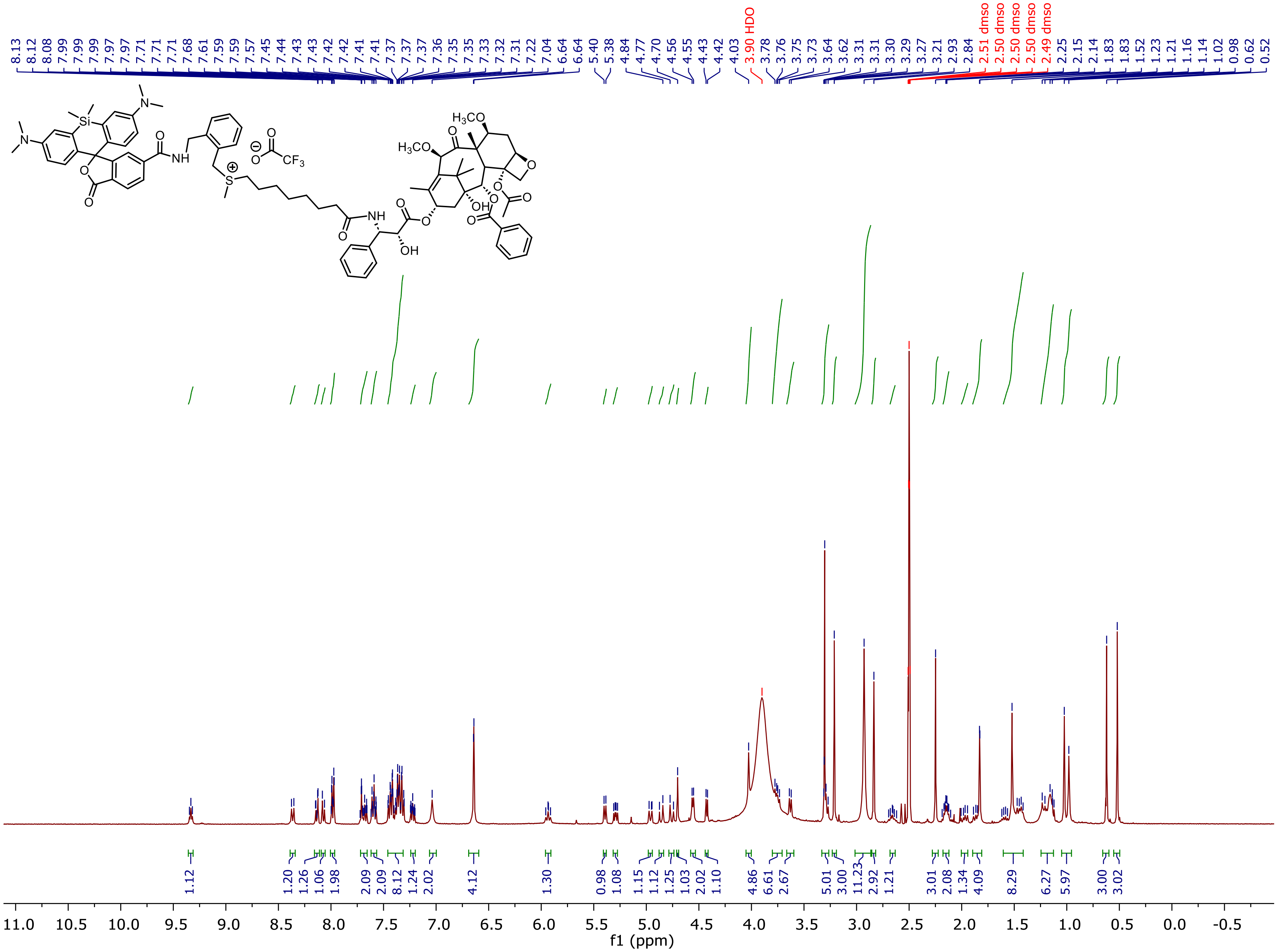

HSQC of **Probe 1:**

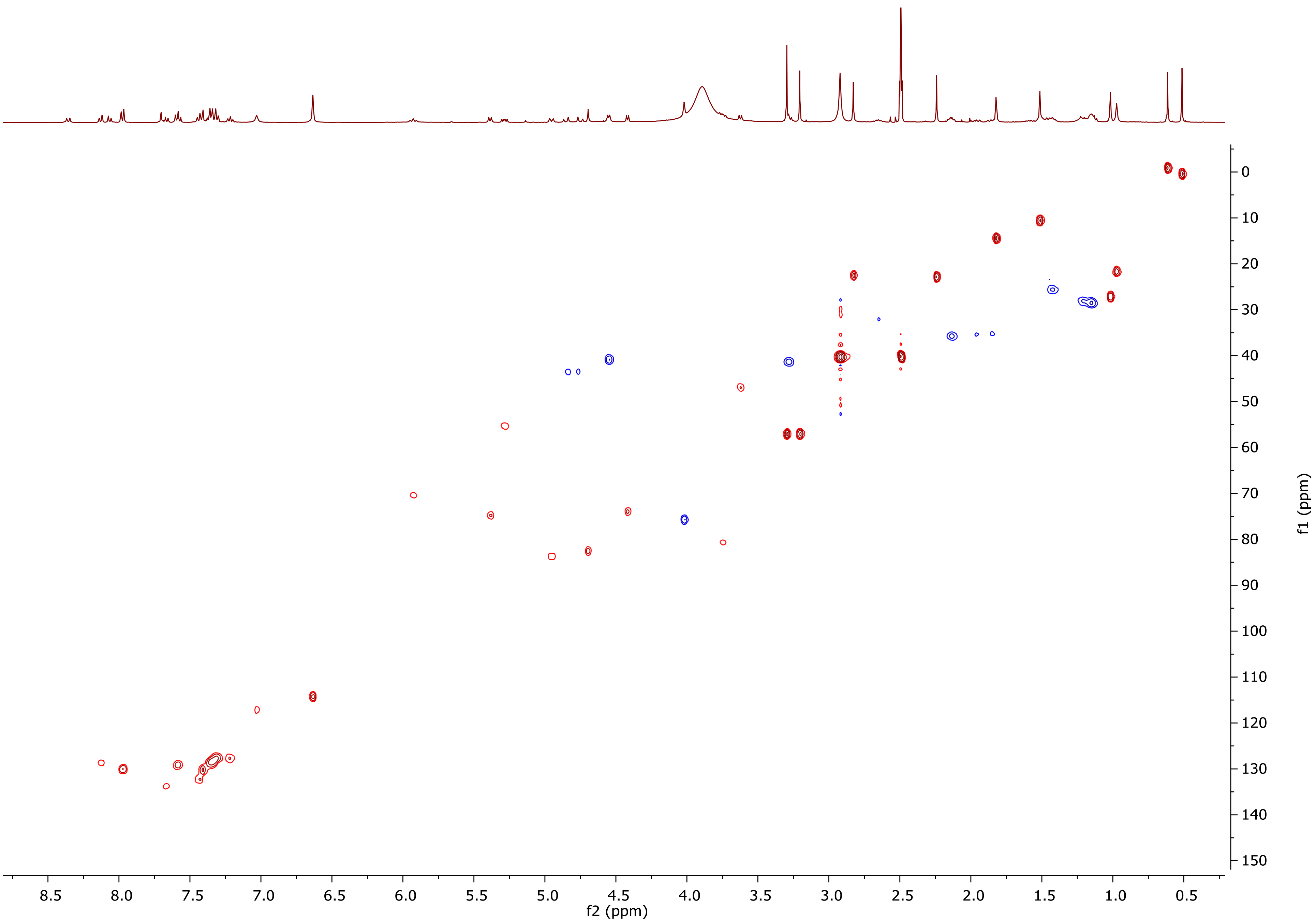

$^{1}H$ NMR of **Probe 2 - 6-SiR-*o*-C$_9$-CTX:**

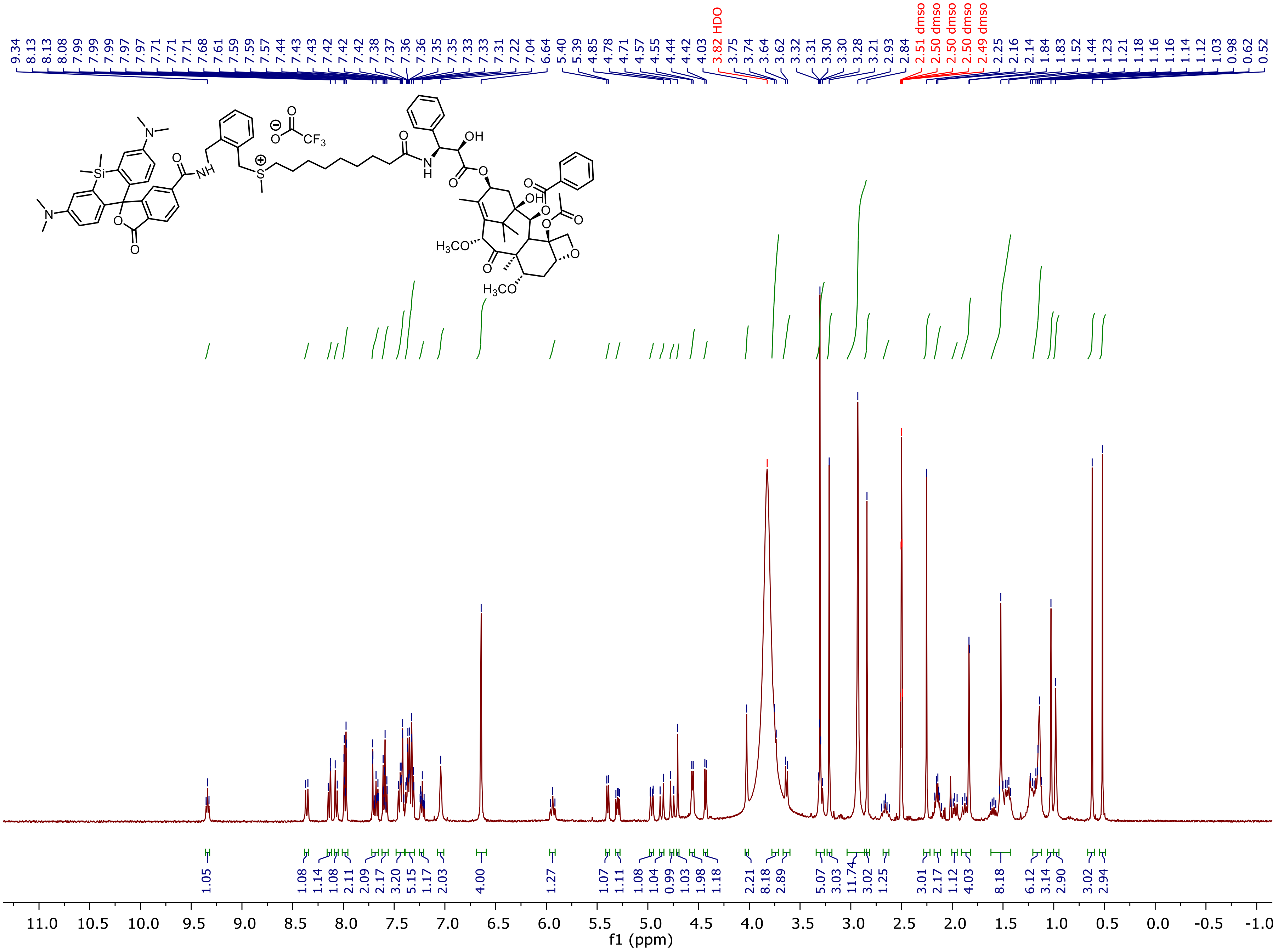

HSQC of **Probe 2 - 6-SiR-*o*-$C_9$-CTX:**

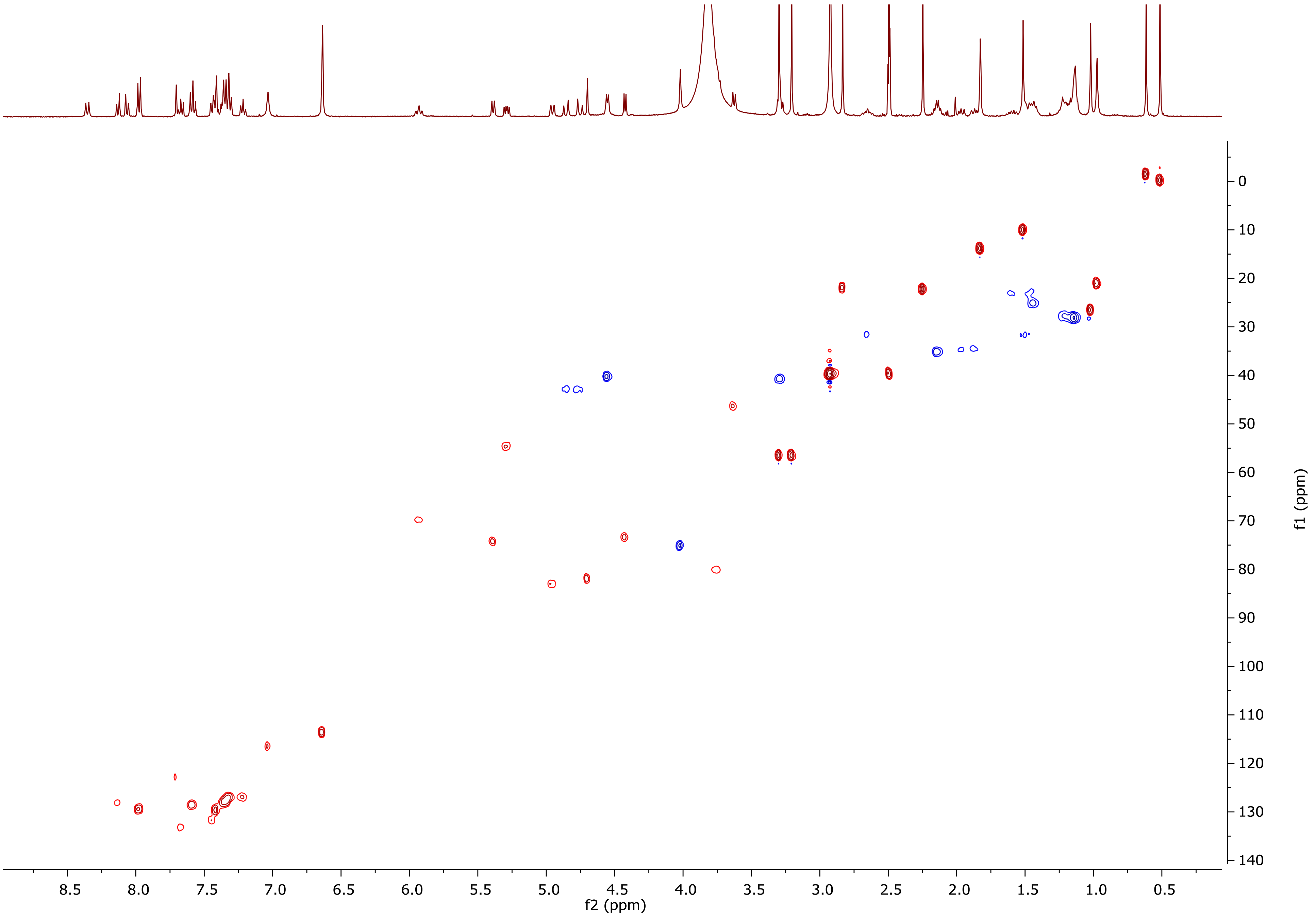

[1]H NMR of **Probe 3:**

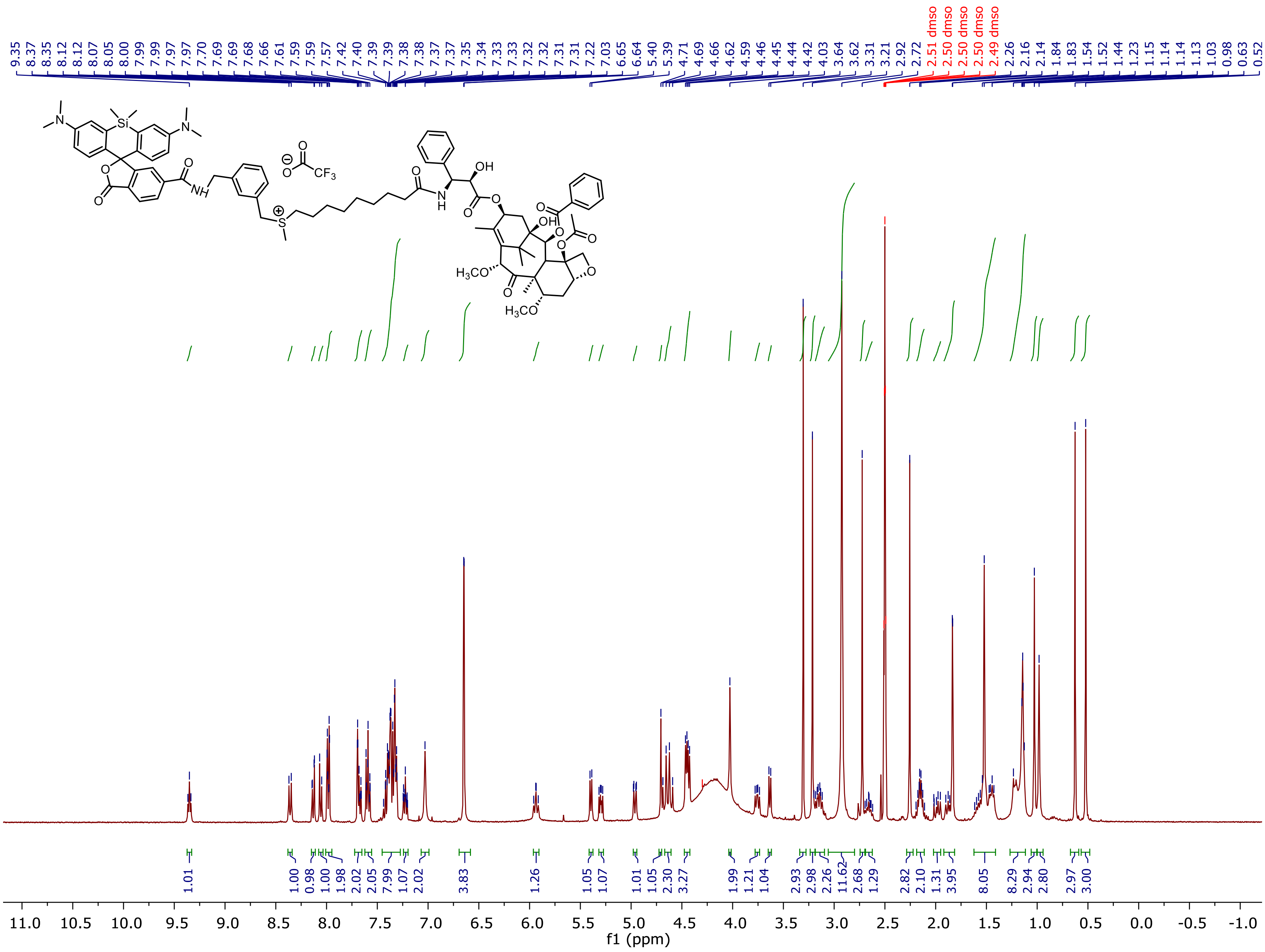

HSQC of **Probe 3:**

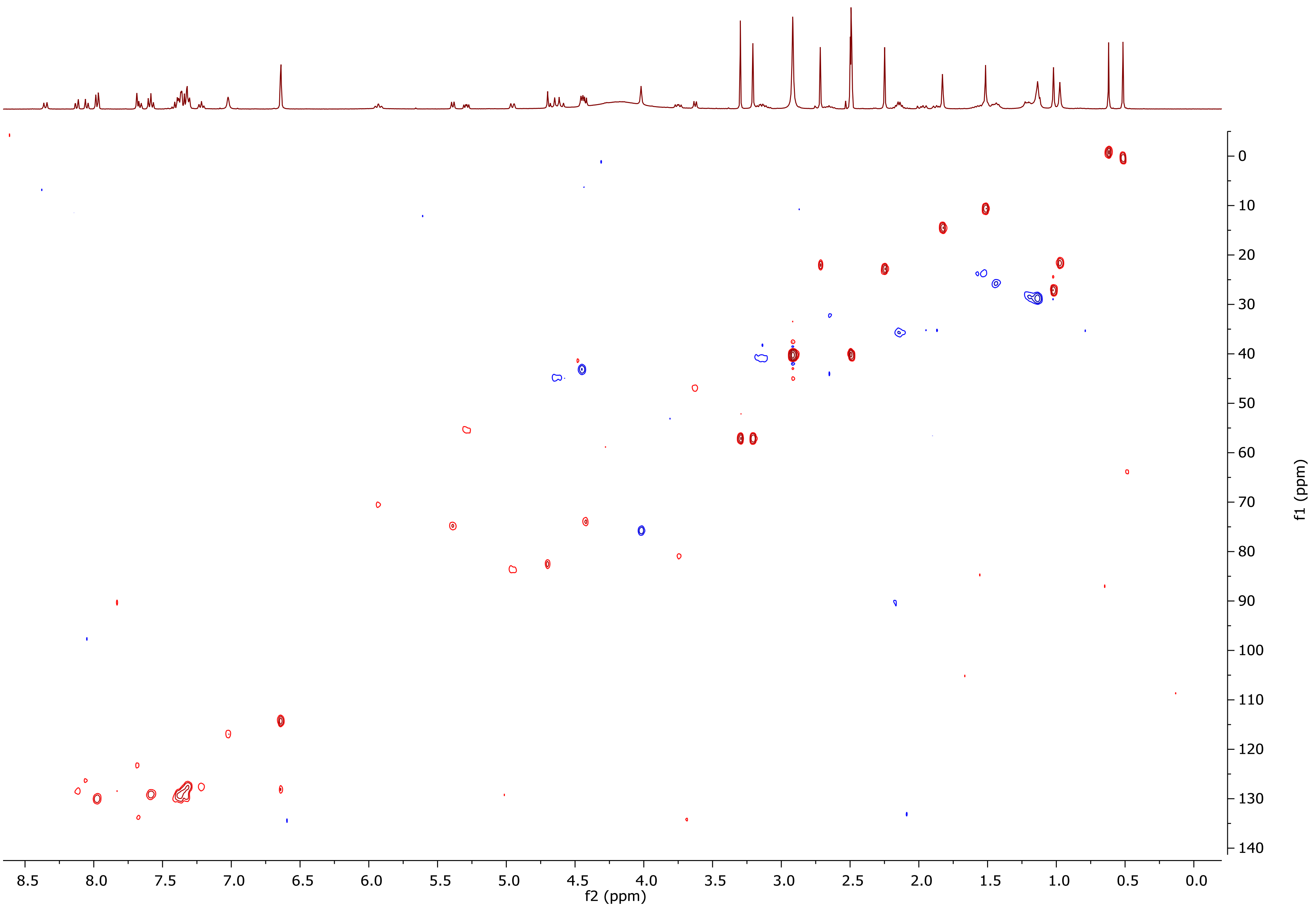

$^{1}$H NMR of **Probe 4:**

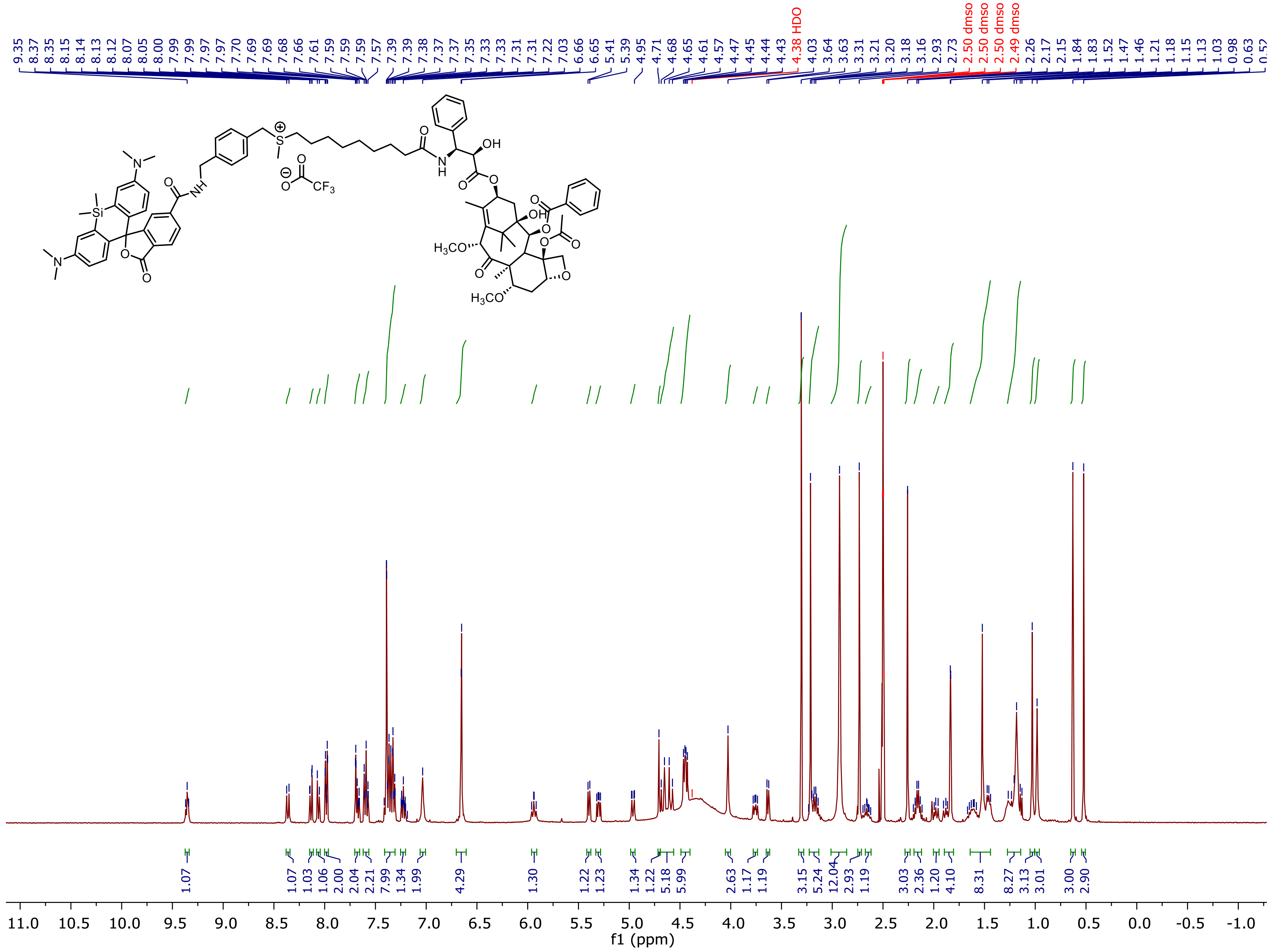

COSY of **Probe 4:**

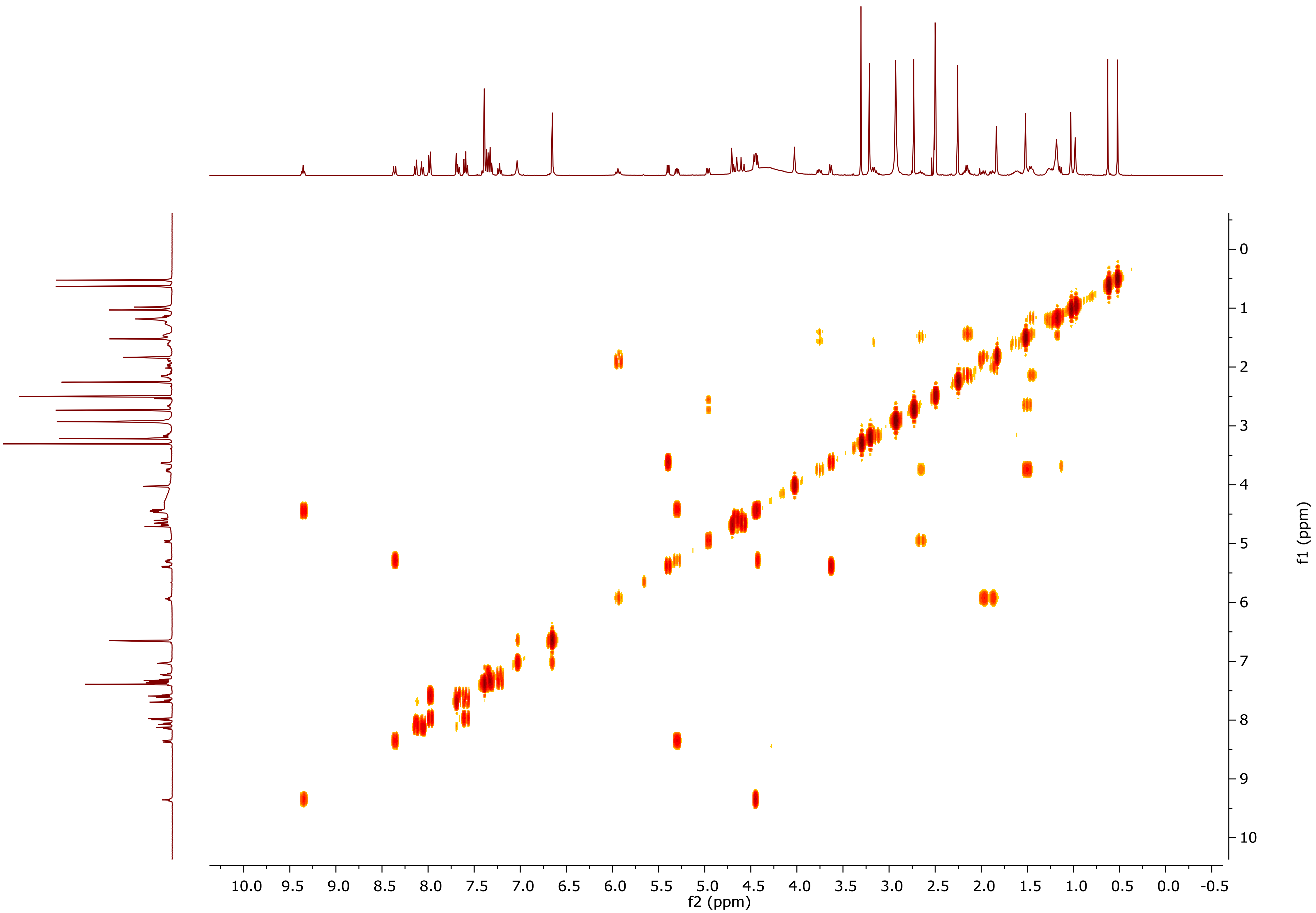

[1]H NMR of **Probe 5:**

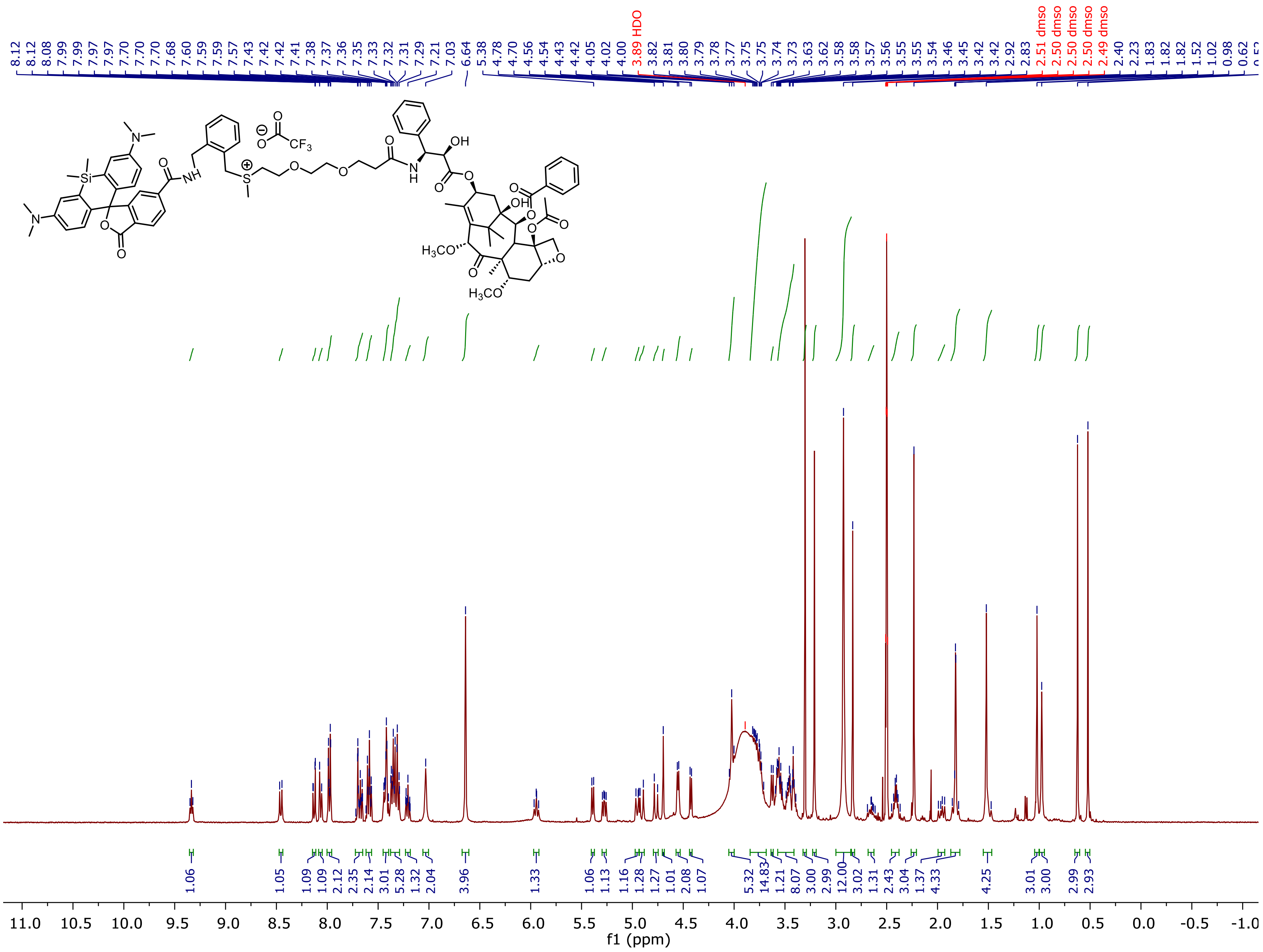

HSQC of **Probe 5:**

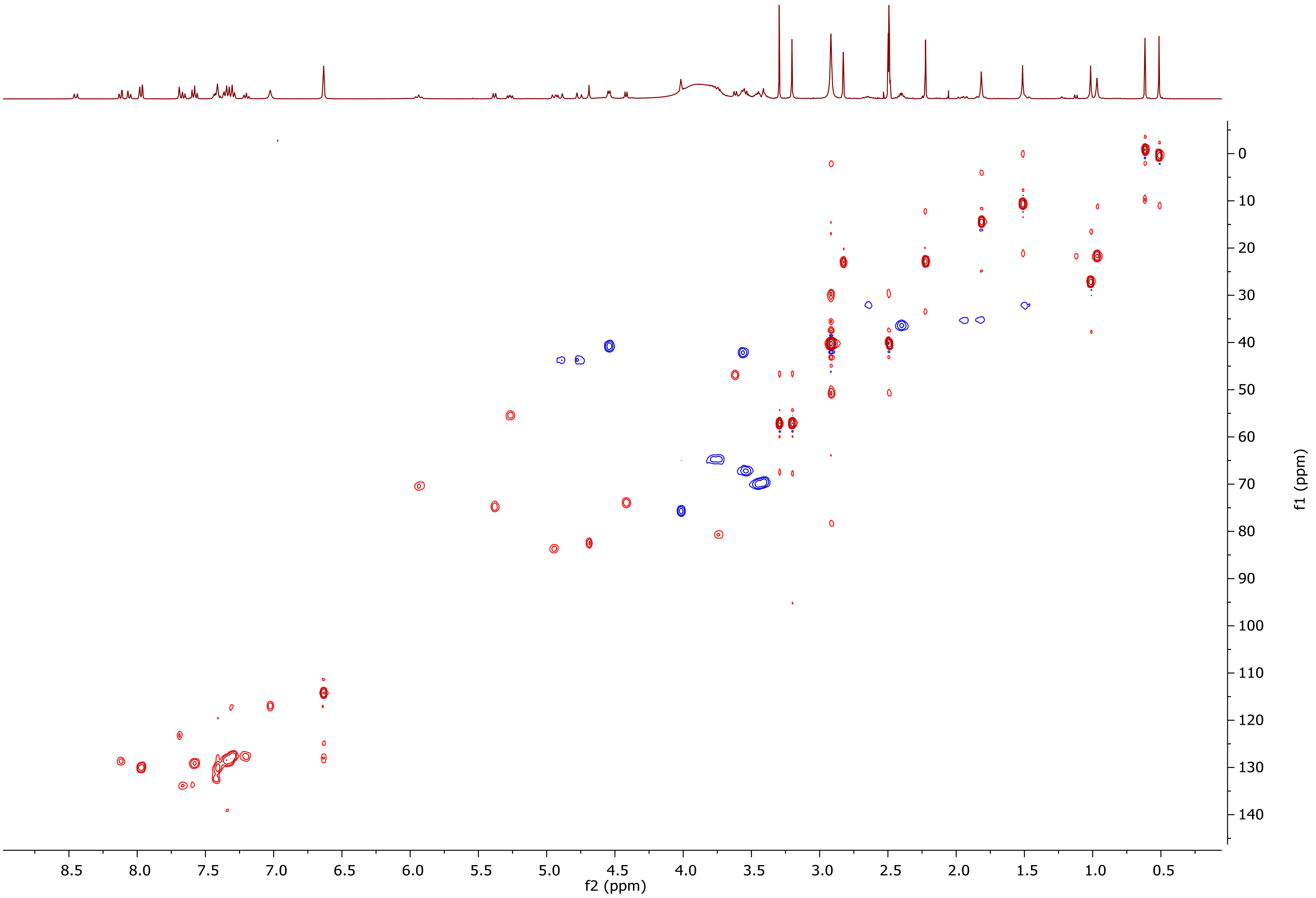

[1]H NMR of **Probe 6:**

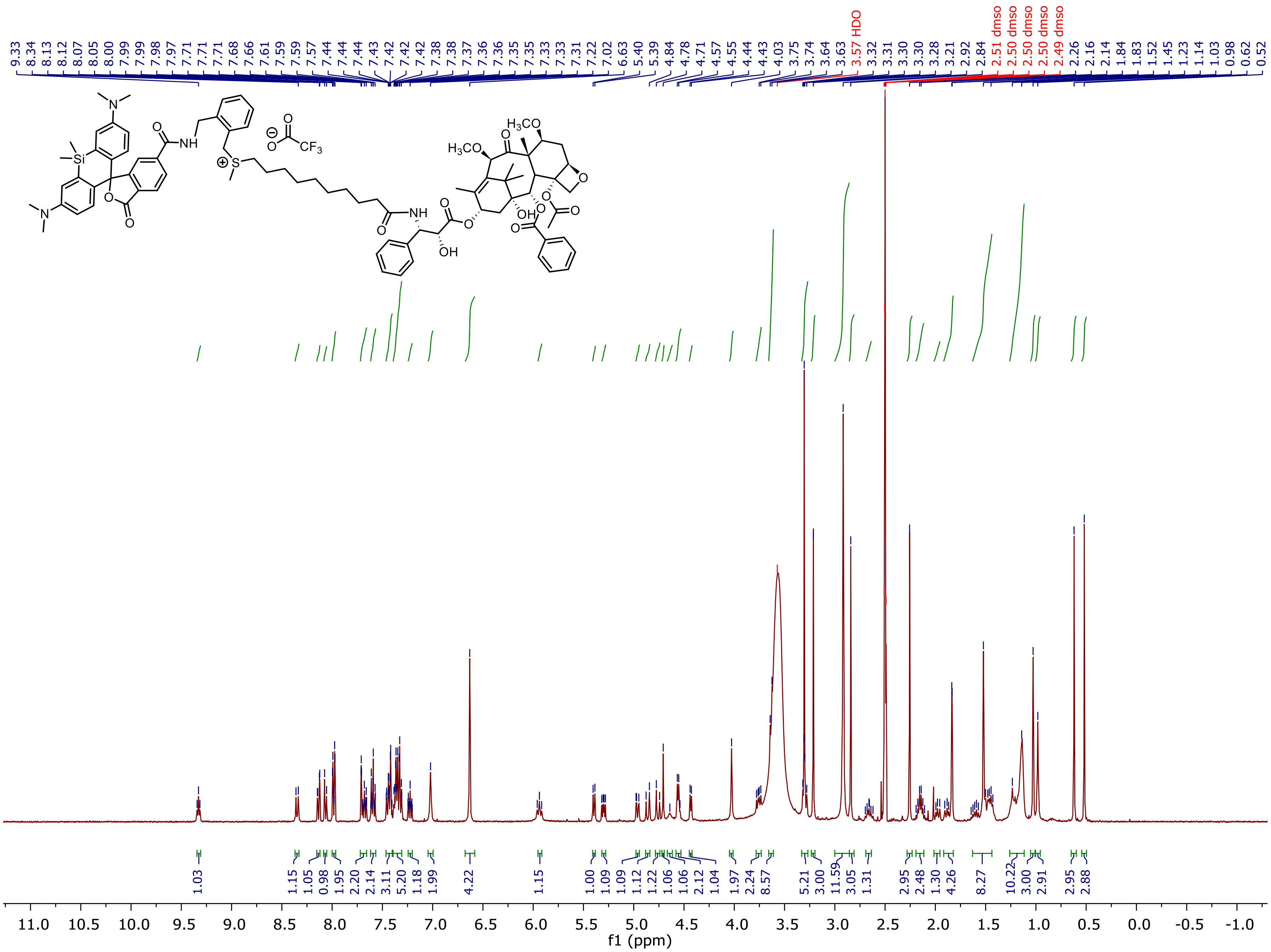

HSQC of **Probe 6:**

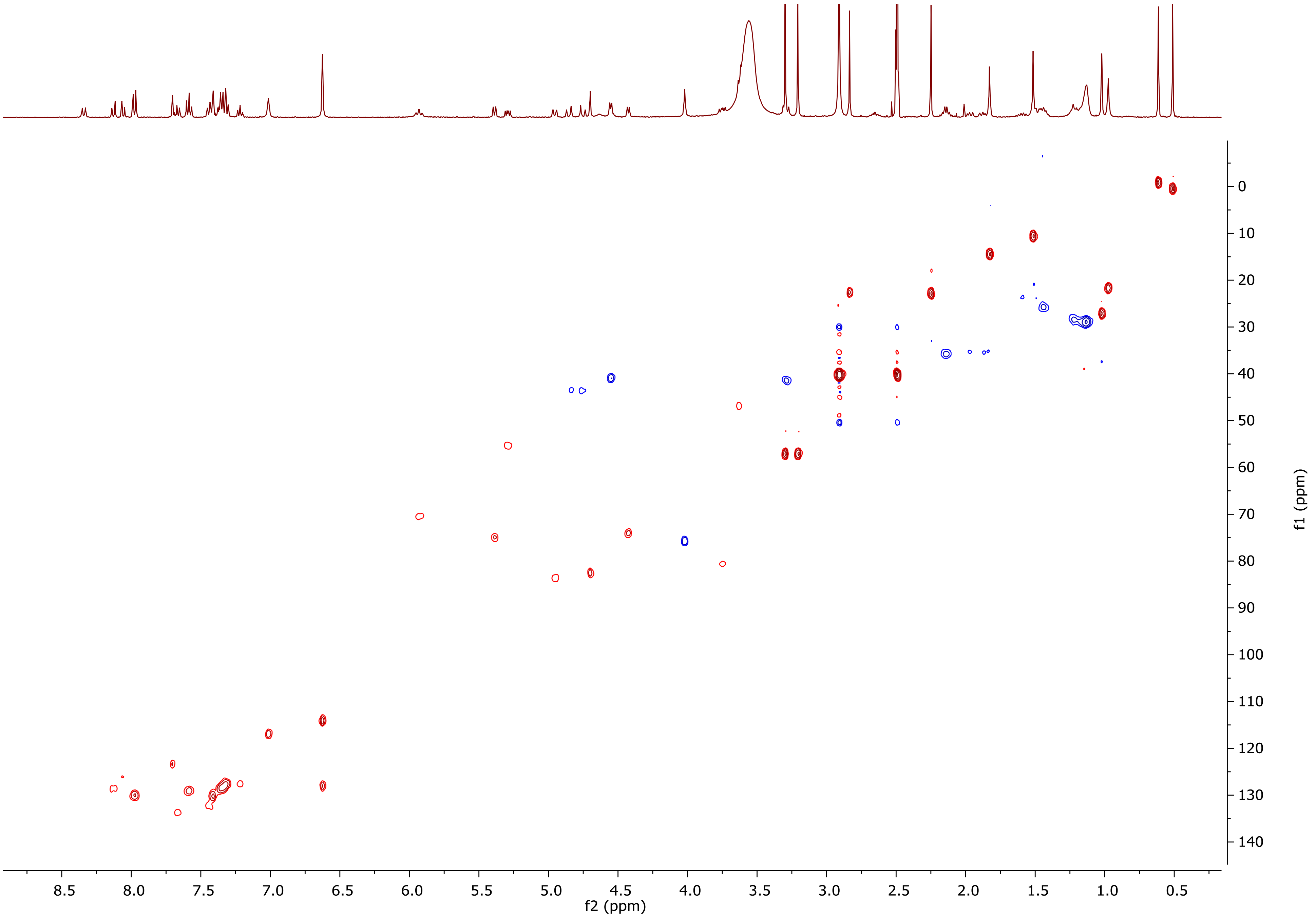

### References

1 Lukinavicius, G. *et al.* Fluorescent dyes and probes for super-resolution microscopy of microtubules and tracheoles in living cells and tissues. *Chem Sci* **9**, 3324-3334, doi:10.1039/c7sc05334g (2018).

2 Bucevicius, J., Keller-Findeisen, J., Gilat, T., Hell, S. W. & Lukinavicius, G. Rhodamine-Hoechst positional isomers for highly efficient staining of heterochromatin. *Chem Sci* **10**, 1962-1970, doi:10.1039/c8sc05082a (2019).

3 Bucevicius, J., Kostiuk, G., Gerasimaite, R., Gilat, T. & Lukinavicius, G. Enhancing the biocompatibility of rhodamine fluorescent probes by a neighbouring group effect. *Chem Sci* **11**, 7313-7323, doi:10.1039/d0sc02154g (2020).
